# Sound from Ultrasound: Characterization of a MEMS-based personal-audio device in human temporal bones

**DOI:** 10.64898/2026.09.10.750660

**Authors:** Sunil Puria, Kevin N. O’Connor, Jeffrey Tao Cheng

## Abstract

Emerging MEMS-based audio devices generate ultrasonic acoustic output, but little is known about long-term biological effects of such exposures, which vary in frequency, intensity, duration, and coupling pathway. In this study, a novel transducer was characterized for its ultrasonic acoustic output, generating amplitude-modulated (AM) pressure pulses at a carrier frequency of approximately 200 kHz and operating frequency of approximately 100 kHz. To understand ultrasonic pressure wave transmission through the human middle and inner ear, sound pressure levels in the ear canal (P_EC_) and mechanical vibration velocities at the stapes (V_ST_) and promontory (V_PRM_) were measured in human cadaveric temporal bones across frequencies up to 240 kHz. This represents the first such measurements at ultrasonic frequencies in human temporal bone specimens. Results demonstrated relatively consistent P_EC_ measurements at ultrasonic frequencies across specimens, with minimal inter-specimen variation. At the operating frequency (∼100 kHz), overall P_EC_ was 82.2±2.3 dB SPL (SNR = 40.6 dB), while at the carrier frequency (∼200 kHz), P_EC_ increased to 90.6 dB SPL (SNR = 34.8 dB). V_ST_ and V_PRM_ measurements at the operating frequency remained relatively independent of audio stimulus-drive frequency and voltage, with comparable magnitudes, suggesting that bone conduction pathways contribute significantly to inner ear ultrasonic exposure. When compared to existing safety guidelines, this device appears to meet specified criteria; however, given the limited scientific basis for these guidelines and unknown long-term effects, cautious application is recommended.

## Introduction

Microelectromechanical systems (MEMS) audio transducers, which typically utilize piezoelectric materials to move a small but highly stiff silicon diaphragm to produce sound, offer numerous benefits over electromagnetic transducers, both in terms of providing high-fidelity performance for end users in the frequency range most suited to the design of the device, and in terms of providing manufacturing advantages such as high precision and scalability. Still, the traditional moving-diaphragm approach of typical MEMS audio transducers also tends to limit performance at lower frequencies due to the typically small sizes and displacements of such transducers.

However, a new class of MEMS device has emerged recently that is claimed to significantly improve low– and mid-frequency acoustic performance, including the ability to produce sufficient low-frequency pressure to drive modern noise-cancelling earbuds, through the application of sound-from-ultrasound techniques [1–4]. By combining the many performance and manufacturing advantages of MEMS devices with the added bandwidth advantages of the sound-from-ultrasound approach, it has now become possible to mass-produce compact and lightweight devices that also feature wide-bandwidth, high-fidelity audio performance—attributes that are very attractive for portable audio applications such as consumer earbuds, virtual-reality headsets, smart watches, and potentially hearing-aid devices [3].

One such device of this class, whose performance and ultrasonic emissions are the focus of this study, operates through the rapid-fire production of precisely modulated air-pressure pulses at an ultrasonic carrier rate of around 200 kHz [4]. The device’s internal moving parts produce ultrasonic motions at half the period of the carrier frequency, which is around 100 kHz. This is called the operating frequency because this has the effect of converting the input audio signal into an audible pressure wave. The Appendix provides further details on the theory of operation for this class of transducers.

In spite of the many apparent advantages of such devices, they nonetheless introduce a new public health concern due to the fact that their widespread adoption would expose a population of people to potentially high levels of ultrasonic emissions, delivered directly into their ears or through skull vibrations, for potentially long periods of time [5, 6].

Depending on the design parameters of a given MEMS audio device, the amount of airborne ultrasonic leakage can correspond to sound pressure levels (expressed in dB SPL) that would substantially exceed comfortable listening levels if they were to occur within audible frequencies. Furthermore, one type of sound-from-ultrasound transducer, the xMEMS Cypress device under consideration in this paper, appears to continuously generate ultrasonic emissions while it is powered on, even in the absence of an input audio signal, which raises further questions about cumulative exposure and potential biological effects that warrant further investigation.

From an official safety perspective, current exposure guidelines establish a general public limit of 100 dB SPL (1/3-octave band) for ultrasonic frequencies in the 25–100 kHz range [7], although this standard was developed well before widespread consumer deployment of ultrasonic transducers in portable audio devices, many of which primarily produce pure-tone ultrasound (for which a safety guideline based on 1/3-octave bands may not be well-suited [6]), and has been criticized for lacking a strong evidential foundation [8]. Potential hazards associated with exposure to airborne ultrasound include cavitation in biological fluids and tissues at frequencies below 500 kHz, although it has been argued that cavitation is unlikely from typical levels of airborne ultrasound [9]. Subjective adverse effects in some individuals may still be possible due to hypothesized mechanisms [6], underscoring the need for empirical characterization of ultrasonic transducer output.

To gauge the degree of any potential safety risk to the human ear from the ultrasonic emissions of one such device, we made pressure and vibrometry measurements up to 240 kHz in three human cadaveric temporal bones with the device coupled to the ear canal. Specifically, we measured the pressure of the acoustic ultrasonic emissions entering the ear canal (in dB SPL), and the 3D velocities of the corresponding middle-ear vibrations (in mm/s), of both the stapes and the cochlear promontory. The stapes is the third and final bone in the ossicular chain connecting the eardrum to the cochlea, and the cochlear promontory forms part of the bony wall encasing the delicate sensory structures of the fluid-filled cochlea.

Mechanical vibrations of the stapes within the “oval window” (i.e., the entrance to the cochlea) are what primarily transmit sound energy from the ear canal to the cochlea through the “air-conduction” (AC) pathway, at least for frequencies within the range of human hearing up to 20 kHz. Another means of transmitting sound energy to the cochlea within the range of human hearing is through various “bone-conduction” (BC) pathways [10–13], of which vibrations of the bony cochlear promontory are a common indicator for some of the BC pathways [13]. Yet another possible pathway is the stimulation of the cochlea through the round window membrane due to vibrations of the eardrum that produces middle ear cavity pressure [14, 15].

Although it is not clear what the primary transmission mechanisms of vibration to the cochlea may be for the ∼100 kHz and ∼200 kHz ultrasonic frequencies emitted by the device being tested in this paper, our assumption is that measuring the 3D vibrations of both the stapes and cochlear promontory will provide an adequate indicator of the level of ultrasonic vibrations reaching the cochlea due to the airborne ultrasonic emissions and vibrations from the device.

Our measurements employed two complementary techniques: an optical microphone for high-frequency measurements of the sound pressure entering the ear canal, and a 3D laser Doppler vibrometer (3D LDV) for measuring ossicular and cochlear-promontory velocities. This dual-measurement approach, combining established 3D LDV methodology for temporal-bone studies [16, 17] with extended-frequency-range pressure characterization, allowed us to distinguish between acoustic transmission through the normal AC pathway to the cochlea via the eardrum and middle-ear ossicles, and BC transmission to the cochlea through alternative mechanical pathways. The results provide new empirical insights into the potential transmission of ultrasonic frequencies to the ear—information critical for assessing the safety profile of sound-from-ultrasound transducers for personal audio and hearing applications.

## Methods

### Equipment and Configuration

Figure 1 illustrates a block diagram of the equipment employed for the measurements. We utilized a National Instruments (NI) USB-6346 multifunction data-acquisition (DAQ) device, which has 8 analog inputs (16-bit bit depth, 500 kS/s sampling rate) and 2 analog outputs (900 kS/s). This device was connected to an NI PXI Windows computer through a USB cable. Pressure was measured using a membrane-free optical microphone with an acoustic frequency bandwidth spanning from 10 Hz to 1 MHz, equipped with a single sensor head and associated electronics (XARION, Eta250 Ultra). A CLV-3D laser Doppler vibrometer (Polytec, Germany) was utilized to acquire 3D velocity measurements with a bandwidth ranging from 0.5 Hz to 250 kHz^1^. In order to reduce any effects of aliasing, the four output signals from the microphone (one channel) and 3D LDV (three channels, representing orthogonal velocities) were subsequently fed through a four-channel low-pass Bessel filter with a 200 kHz bandwidth (Krohn-Hite 3384) on their way to the input channels of the USB-6346. This filter rolled-off with frequency with attenuation of less than 1 dB near the operating frequency and about 6 dB near the carrier frequency. No attempts were made to compensate for this and thus the actual pressures and velocities might be slightly higher than reported here. To minimize environmental vibration artifacts, the equipment and prepared temporal-bone specimens were placed on top of an air-suspended optical table.

**Figure 1:**
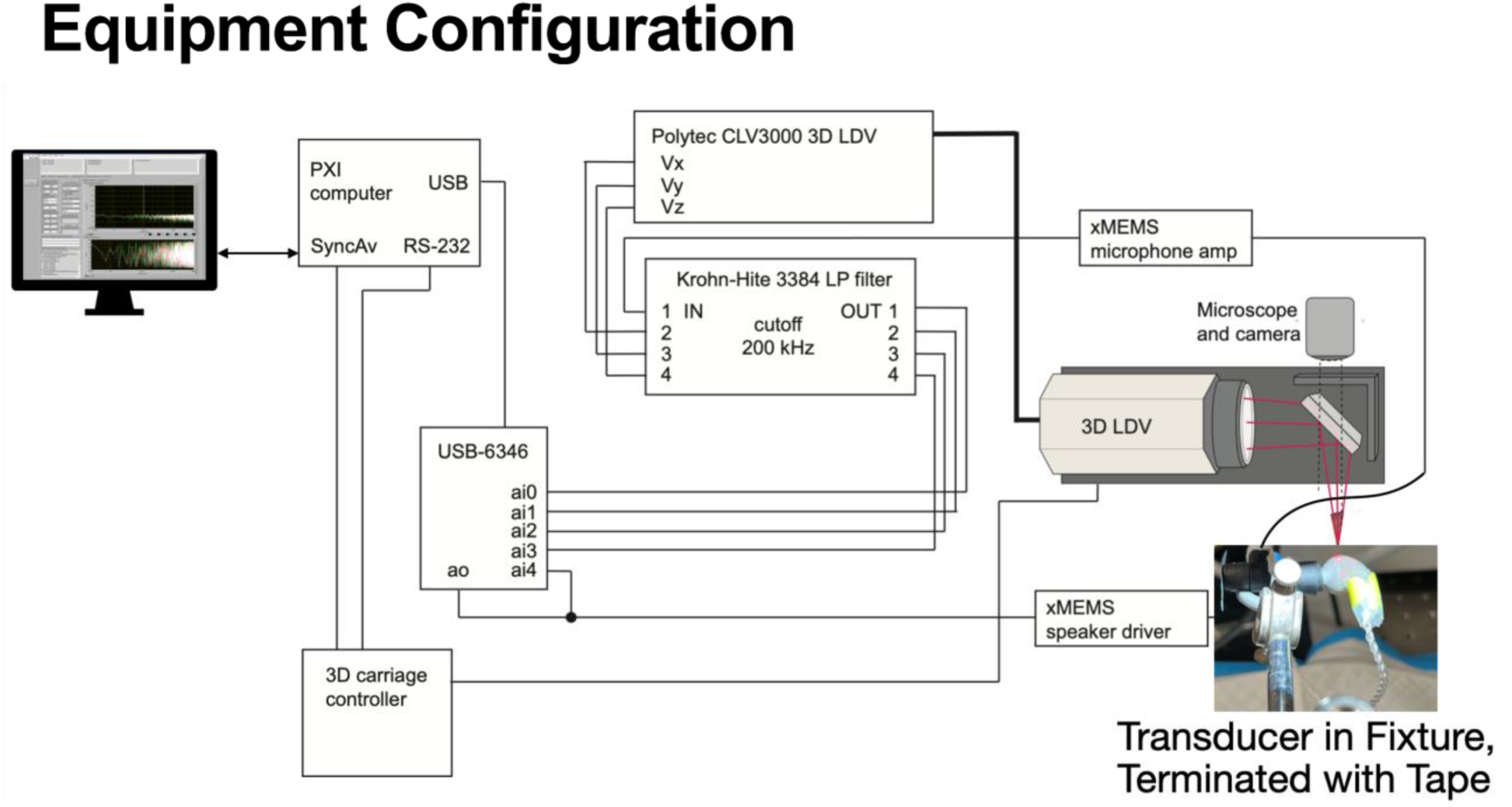
Block diagram of the equipment configuration for the experiment. A Windows PXI computer (NI, USA) running custom LabVIEW-based SyncAv software is connected to a USB-6346 data-acquisition board (NI) through a USB cable, as well as a C142-4 3D carriage controller (Isel-automation, Germany) through an RS-232 serial cable. The USB-6346 produces an analog output signal (“ao”) that provides the audio-band input signal to the xMEMS speaker driver. The ao signal is copied into the “ai4” analog input for recording alongside the other inputs. The xMEMS speaker driver is wired to the “Cypress” transducer under test to provide the power and other signals necessary to acoustically generate the input audio signal, as well as additional ultrasonic leakage artifacts, based on its “sound-from-ultrasound” design. The image in the lower-right corner depicts the earbud with a foam ear tip inserted into the right-hand side of a 3D-printed black coupling tube, which is terminated with another foam ear tip on the left. The second ear tip is terminated with a piece of household tape in this example, but it would be inserted into the ear canal of a prepared human cadaveric temporal-bone specimen for an actual experiment. The Eta259 Ultra optical microphone (XARION, Austria) is inserted into a specially made port in the side of the coupler, and is wired into the xMEMS microphone amp. The CLV-3D 3D laser Doppler vibrometer (3D LDV; Polytec, Germany) is positioned using the 3D carriage controller and motorized stage, and in this example its three laser beams are focused on the earbud housing. In an actual experiment the beams would be focused on retroreflective targets placed on the stapes head or the bony cochlear promontory. The output of the xMEMS microphone amp and the three outputs of the 3D LDV pass through a Krohn-Hite 3384 low-pass filter, with a 200 kHz cutoff frequency, on their way to analog inputs ai0-ai3 of the USB-6346.

A custom application written in LabVIEW (NI) and running in Windows (Microsoft Corporation), called “SyncAv” (ver 0.4) was used to generate output tones and collect input measurements [e.g., 18]. SyncAv is an FFT-based measurement and analysis system for which the output and input channels are synchronized with one another. The A/D and D/A converters were configured with a sampling rate of 480 kS/s and an FFT length of 16,384 samples, corresponding to a minimum frequency (and frequency resolution) of 29.297 Hz and a maximum frequency of 240 kHz.

The sensor head of the 3D LDV was mounted on three X, Y, Z motorized stages (Kugelgewindevorschub LF5, 6; Isel-automation, Germany) that were controlled through commands sent from SyncAv to the 3D-carriage controller (Schrittmotor-Controller C142-4; Isel-automation, Germany) via an RS-232 cable.

### The MEMS-based “Sound-from-Ultrasound” Transducer Under Test

For the present experiments (Fig. 2), a prototype “Cypress” earbud system (xMEMS Labs, San Jose, CA) was used, which consisted of a proprietary MEMS-based “sound-from-ultrasound” driver, specialized drive electronics, associated cabling, and a 3D-printed housing in the shape of a typical “true wireless stereo” (TWS) earbud. The output signal from the USB-6346 (representing the stimulus waveform) was used as the input to the drive electronics, which in turn transformed the stimulus into a pair of specially tailored analog signals that were then used to drive the MEMS device in a manner appropriate for its design (see Appendix for details).

**Figure 2:**
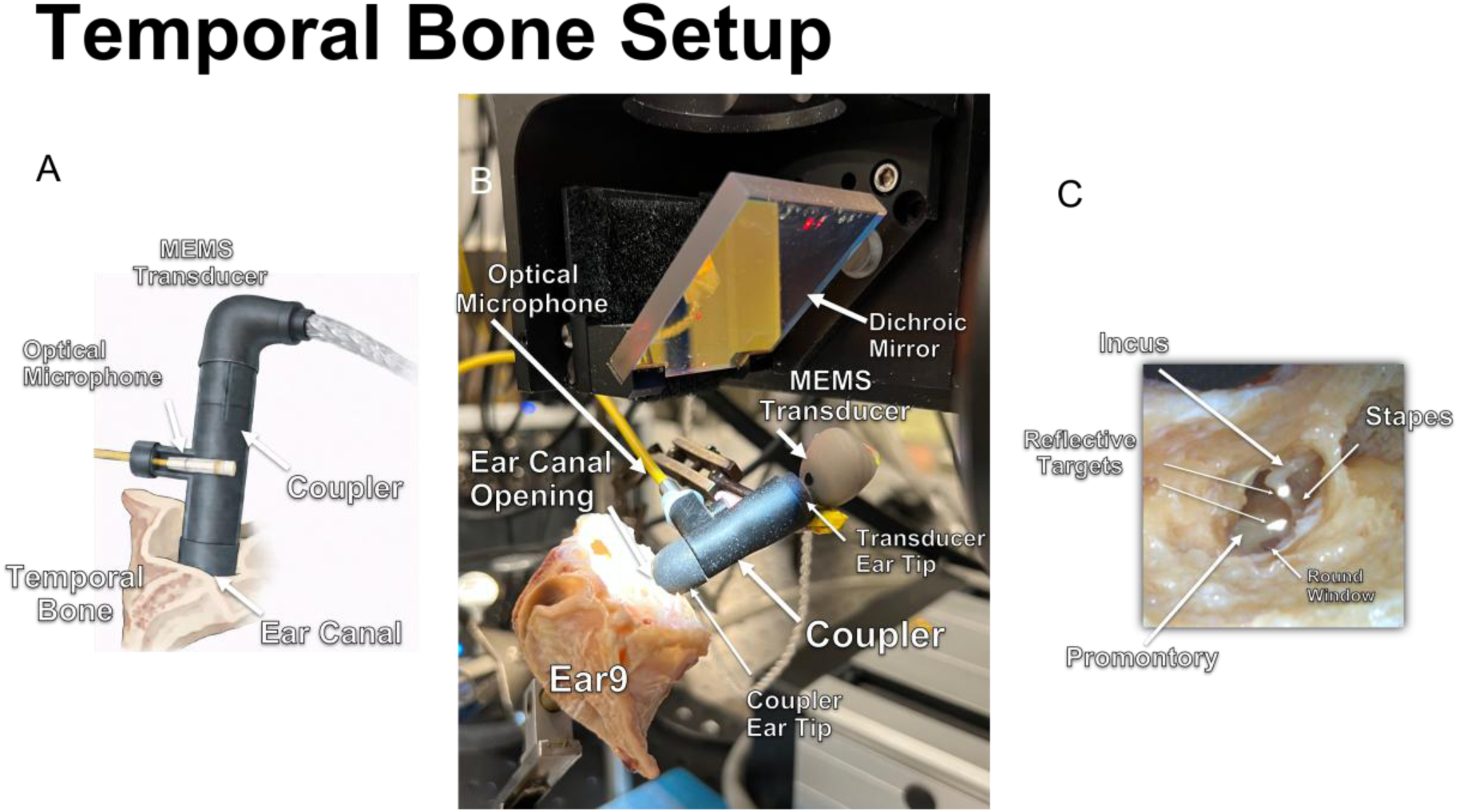
Depictions of the experimental setup used to deliver sound to the temporal bone’s ear canal and measure pressure and velocities. Panel A shows a stylized drawing of the earbud (“MEMS Transducer”) and the 3D-printed coupler (“Coupler”) with a cutout showing the tip of the optical microphone inserted into the body of the coupler through a specially designed side port. Panel B shows the actual assembly inserted into the ear-canal opening of specimen Ear9. In this case the beams of the 3D LDV are focused on the body of the coupler rather than any structures within Ear9. The dichroic mirror enables mounting of the 3D LDV at a 90-degree angle to the specimen, which is mounted on a motorized stage. A microscope allows visualization of the specimen below through the dichroic mirror. Panel C shows a zoomed-in view of the opened middle-ear cavity with retroreflective targets placed on the stapes head and cochlear promontory. In an actual experiment the three beams of the 3D LDV would be aimed at one of these targets to record its vibration.

We made measurements using three different variations of the Cypress earbud system, referred to as Transducer “1”, “1b”, and “2a”. The Cypress systems each produced ultrasonic artifacts at a “carrier frequency” (*f*_c_) and an “operating frequency” (*f*_o_), where *f*_o_ is *f*_c_/2. The carrier and operating frequencies were tied to the inner workings of the device. The audio signal was evidently amplitude modulated (AM) at both *f*_c_ and *f*_o_, as spectral spikes appeared in the wideband acoustic output of the device at these two ultrasonic frequencies, and each had accompanying AM side bands. For the purposes of this study, the ultrasonic spikes at *f*_c_ and *f*_o_ in the device’s acoustic output signature are regarded as ultrasonic leakage from a “black box” device whose inner workings are beyond the scope of this study. The frequencies and amplitudes of these ultrasonic artifacts were designed by the manufacturer to be somewhat different for each variation of the Cypress system (among other differences).

The aim of this study was to characterize the extent to which airborne ultrasonic artifacts produced by the Cypress system appear in measurements of the ear-canal pressure, 3D stapes velocity, and 3D cochlear-promontory velocity. In so doing, we aim to quantify the extents to which an end user of the device might be exposed to 1) direct airborne ultrasound in the ear canal; 2) mechanical ultrasonic vibrations of the stapes head, which relate to ultrasonic exposure in the cochlea through the AC pathway of hearing [19]; and 3) mechanical ultrasonic vibrations of the bone surrounding the cochlea, which relate to ultrasonic exposure in the cochlea through the BC pathway of hearing [12].

This study is primarily focused on quantifying *airborne* ultrasonic emissions from the Cypress earbud systems and the extent to which they induce vibrations of the middle and inner ear, and to this end two sets of mechanically isolating foam ear tips (see Fig. 2) were used in the measurement protocol to reduce any ultrasound transmission to the temporal-bone specimens through direct mechanical coupling.

### Postmortem human temporal bones

A total of nine human postmortem temporal-bone specimens were used in the development and carrying out of this study. The specimens were harvested from the Massachusetts General Hospital (MGH) morgue in accordance with Human Subjects Protocol # 2021P002348 from the Mass Eye and Ear Otopathology Laboratory. Human-subject research in this manuscript complies with the criteria for exemption from the Mass General Brigham Institutional Review Board (IRB) requirements as outlined in the Department of Health and Human Services (DHHS) regulations (45 CFR 46), which necessitates participant consent. The present research falls under Exemption 4: Research involving the collection or study of existing data, documents, records, pathological specimens, or diagnostic specimens, provided these sources are publicly available or if the information is recorded by the investigator in a manner that prevents the identification of subjects, either directly or through identifiers linked to them.

All temporal-bone specimens were obtained from donors with no history of otologic disease. Upon arrival, the specimens were visually screened for outer ear and eardrum pathologies and labeled as “Ear1” through “Ear9”. All specimens were immediately frozen in a –20°C freezer after harvesting until they were ready for the experiment (within 2–3 months of freezing). They were thawed in a refrigerator in saline with a few drops of iodine as an antibacterial agent overnight before preparation, and stored in the refrigerator between measurement sessions. After completion of the experiments, the specimens were wrapped in 0.9% saline-soaked gauze, were further wrapped in plastic food wrap and aluminum foil, and were then placed back in the freezer.

### Ear specimen preparation and setup

During dissection and preparation, each temporal bone was secured within a standard Ear-specimen holder using its included adjustable screws. Using an otologic drill, the middle-ear cavity (MEC) was partially opened through the posterior–inferior facial recess to check the normality of the middle-ear ossicles (malleus, incus, and stapes). The MEC was opened just enough to allow the placement of retroreflective targets on the stapes and cochlear promontory and to provide clear optical access to those targets for the three beams of the 3D LDV. A retroreflective-tape target was placed on the stapes head, just medial to the incudo-stapedial joint. This location was chosen to minimize disturbance to the ossicles. A second retroreflective-tape target was placed on the bony cochlear promontory (see Fig. 2C). The tendon of the stapedius muscle was left intact, as this structure may potentially affect high-frequency sound transmission in the middle ear. The prepared ear specimen was then brought to the optical table for the sound-pressure and 3D LDV measurements, where it was held firmly by a manipulator that was attached to a metal-rod assembly screwed down to the vibration isolation table.

### Sound stimulation and ear-canal sound pressure

Under normal use, the MEMS-based earbud under test would have had its ear tip placed directly into the user’s ear canal. However, such an arrangement for the temporal-bone experiments would not have allowed the ear-canal pressure generated by the transducer to be measured in situ using the optical microphone. To facilitate these pressure measurements, the ‘transducer ear tip’ was inserted into a custom 3D-printed coupler (33 mm in length), and a ‘coupler ear tip’ attached to the far end of the coupler was then inserted into the actual ear canal of the specimen (Fig. 2A, B). The coupler featured a special side port to accommodate the head of the optical microphone, such that the pressure between the transducer and the ear canal could be measured in a way that safeguarded the delicate, sensitive, and expensive sensor head. The distance between the microphone sensor and the ear tip inserted into the ear canal was 10 mm. The distance between that ear tip, when inserted into the ear canal, and the eardrum varied from specimen to specimen and was not measured, but is estimated to be in the 5–10 mm range. Thus, the distance between the eardrum and the microphone sensor was approximately 15–20 mm.

## 3D velocity of the stapes and cochlear promontory

The methods used in this study for 3D-motion measurements have been described previously [17, 20]. Briefly, the three beams of the 3D LDV were aimed at one of the retroreflective targets, using SyncAv to control the position of the sensor head, and the resulting coordinates were recorded. The same procedure was then repeated for the second target. By recording both sets of coordinates in SyncAv, it was possible to easily move the 3D LDV back and forth between the two targets whenever needed. Each time the LDV was repositioned, the locations of the laser beams relative to the target were verified through the scope-mounted camera attached to a computer monitor, and the reflectivity strength of the three laser beams was verified using the LED signal-level indicators on the LDV controller as a further test of correct positioning.

The 3D LDV measured the velocity of each target in three orthogonal directions. The measured velocity components of the retroreflective targets are in terms of the reference frame of the 3D LDV: V_X_ (left–right) and V_Y_ (front–back) in the horizontal plane, and V_Z_ (up–down) in an orthogonal vertical plane. The three LDV beams were reflected by an angled dichroic mirror (Fig. 2B) such that the effective Z-axis remained vertical.

### Measurement Procedure

The following data-collection steps were performed for Ear7, Ear8, and Ear9:

1. Position the 3D LDV at the stapes and measure (P_EC_, V_ST_)
  a. Stepped-tone responses at 0.2 V_PEAK_ followed by 0.05 V_PEAK_ (161 log-spaced tones from 59 Hz to 200 kHz, with 10 averages each)
  b. Stepped-tone noise-floor response (repeat with 1e-6 V_PEAK_ output level, 10 averages)
  c. Chirp responses at 0.2 and 0.05 V_PEAK_ (100 averages)
  d. Chirp noise-floor response (repeat with 1e-6 V_PEAK_ output level, 100 averages)
  e. 100 single-tone measurements (5 frequencies x 2 levels x 10 repetitions, with no averaging)
2. Uncouple and re-couple the transducer from the ear canal, then re-measure (P_EC_, V_ST_)
  a. Repeat the stepped-tone measurement at 0.2 V_PEAK_
  b. Repeat the 100 single-tone measurements (5 frequencies x 2 levels x 10 repetitions)
3. Reposition the 3D LDV to the cochlear promontory and measure (P_EC_, V_PRM_)
  a. Stepped-tone responses at 0.2 and 0.05 V_PEAK_ (10 averages)
  b. Stepped-tone noise-floor response (10 averages)
  c. Chirp responses at 0.2 and 0.05 V_PEAK_ (100 averages)
  d. Chirp noise-floor response (100 averages)
  e. 100 single-tone measurements (5 frequencies x 2 levels x 10 repetitions, with no averaging)
4. Reposition the 3D LDV to the stapes and re-measure (P_EC_, V_ST_)
  a. Repeat the stepped-tone response at 0.2 V_PEAK_ (10 averages)

For the two output levels, the drive voltage was set to either 0.2 V_PEAK_ or 0.05 V_PEAK_ at a given time, for the single-tone, stepped-tone, and chirp measurements.

For the stepped-tone frequency-response measurements, 161 log-spaced stimulus tones ranging from 59 Hz to 200 kHz were presented sequentially and the responses at the stimulus frequencies were recorded and plotted. The number of averages was set to 10 for the stepped-tone measurements, which theoretically resulted in an increase in the signal-to-noise ratio (SNR) by approximately 9 dB (in comparison to no averaging) at the individual tones measured.

For spectral measurements using a single-tone stimulus, the number of averages used for the FFT calculations was set to 1 (i.e., no averaging). For these single-tone measurements, the stimulus frequency for a given measurement was selected from the set of exact FFT frequencies closest to 0.5, 1, 3, 10 and 20 kHz. The above procedure produced 200 pairs of (P_EC_, V_ST_) and 100 pairs of (P_EC_, V_PRM_) spectral measurements for each ear.

We also made additional control and equipment measurements, which are discussed later in the text.

### Analysis

One of the unique characteristics of the present study is that we made velocity measurements of the stapes and cochlear promontory in three dimensions using a 3D LDV. It is known that for frequencies below the ME resonance around 1 kHz, the footplate of the stapes exhibits a piston-like motion that displaces cochlear fluid. However, at higher frequencies within the audible range, the stapes motion also includes non-piston-like rotational components around its long and short axes. At ultrasonic frequencies, it is very likely that there are motions in all three dimensions, and it is possible that other forms of vibrational transmission such as bone compression and bending through the ME structures beyond mere translation and rotation could also come into play. Thus, we did not try to project the stapes or promontory motions onto any particular 1D reference axis as we have done previously [16]. Instead, a resultant velocity magnitude was obtained from the orthogonal velocity components (*Vx*, *Vy*, *Vz*) from the 3D LDV. The resultant velocity magnitude was calculated using the method described in [21]. Since we were interested in the maximum magnitude of the transmitted velocity, particularly at the ultrasonic frequencies, we did not attempt to calculate or report the phase of the 3D motions. All pressure measurements P are reported in dB SPL relative to a 20 µPa reference pressure P_REF_ in air, calculated using the formula P_dB SPL_ = 20*log_10_(P/P_REF_).

The reference “spike-adjacent noise-floor average” (NF) for a given spike in the spectrum of a single-tone measurement was calculated by averaging the magnitudes at the 3 spectral bins below and the 3 spectral bins above the respective target center frequency. Because the frequency bins were 29.297 Hz apart, this corresponds to a NF bandwidth of about 176 Hz around each center frequency. Reference NFs of this type were calculated for each single-tone measurement at the stimulus-drive frequency as well as the *f*_o_ and *f*_c_ ultrasonic-emission frequencies.

For the stepped-tone frequency-response measurements, the computed resultant velocity magnitude of the stapes head or promontory bone was normalized by the simultaneously recorded ear-canal pressure magnitude to compare the V_ST_/P_EC_ and V_PRM_/P_EC_ middle-ear transfer functions across the specimens within the auditory frequency range up to 20 kHz (Fig. 5).

## Results

### Preliminary results of transducer measurements

Measurements on Ear1–Ear6 were conducted to practice specimen preparation, develop the coupler design, and establish the measurement protocol and analysis scripts. These preliminary studies were initially performed using Transducer 1. Measurements were taken at its maximum input voltage of 2.83 V_PEAK_ and at 0.283 V_PEAK_, which is 20 dB below the maximum input voltage. Regardless of the input voltage, the operating frequency for Transducer 1 remained constant at 89.882 kHz, resulting in a sound level of approximately 123 dB SPL. At the carrier frequency (179.795 kHz), the sound level was 92.5 dB SPL. The high sound level generated at the operating frequency necessitated a redesign of the transducer, leading to the introduction of two new versions: Transducers 1b and 2a. These redesigns resulted in a significant reduction, by more than 30 dB, of the sound levels at the operating frequency (as shown in Table 1).

**Table 1:** Operating and carrier frequencies and representative pressures measured with a 1 kHz input tone for the three transducer designs tested.

| Transducer | 1 kHz Drive Level ($V_{PEAK}$) | 1 kHz Drive Pressure (dB SPL) | Operating Frequency (kHz) | Operating Level (dB SPL) | Carrier Frequency (kHz) | Carrier Level (dB SPL) |
| --- | --- | --- | --- | --- | --- | --- |
| **1** | 0.283 | 76.6 | 89.882 | 122.7 | 179.795 | 92.5 |
|  | 2.83 | 96.8 | 89.882 | 123.0 | 179.795 | 92.6 |
| **1b** | 0.1 | 99.5 | 98.847 | 90.8 | 197.695 | <u>90.4</u> |
|  | 1 | 119.6 | 98.847 | 90.8 | 197.695 | 90.1 |
| **2a** | 0.1 | 97.1 | 98.847 | 86.2 | 197.695 | <u>86.0</u> |
|  | 1 | 118.2 | 98.847 | 86.0 | 197.695 | <u>86.1</u> |

Additionally, during the redesign process, the operating frequency was increased from 89.882 kHz to 98.847 kHz, with the corresponding carrier frequency increased from 179.795 kHz to 197.695 kHz, and the maximum allowable input voltage was reduced to 1 V_PEAK_. Notably, the pressure at the carrier frequencies remained relatively unchanged, exhibiting only a slight decrease of a few dB due to the redesigns (Table 1).

At the lower transducer input voltages, such as 0.283 and 0.1 V_PEAK_, the upper and lower AM sidebands were 6 dB or more below the central tone. Consequently, the modulation index was between 10% and 30%. However, when the transducer input levels were increased to 2.82 and 1 V_PEAK_, the sidebands exceeded the level of the central tone (not shown). This indicates that the AM modulation index was in the 170% and 550% range in these cases.

While the levels generated by Transducers 1b and 2a were quite similar, we opted to use Transducer 1b because its output was approximately 2 dB higher at the audio-band drive tones for the same transducer input levels. The results obtained using Transducer 1b are presented next.

### Device behavior when coupled to a simple closed cavity

For our initial assessment of Transducer 1b, we sought to measure the pressure it generated when connected to a simple closed cavity. To accomplish this, we used the custom ear-canal coupling fixture with its ear tip terminated using household tape, in order to roughly approximate a mobile termination akin to the eardrum (see inset photo in Fig. 1). We inserted the transducer ear tip into the other end of the coupler and used the optical microphone to measure the pressure generated by the device in response to stepped-tone inputs (59 Hz to 200 kHz) with tone amplitudes of 0.2 and 0.05 V_PEAK_. We also measured the 3D vibrations of the device housing, the transducer ear tip, and the coupler ear tip. Figure 3 shows the measurement results for the human audio band (up to 20 kHz), for simplicity.

**Figure 3:**
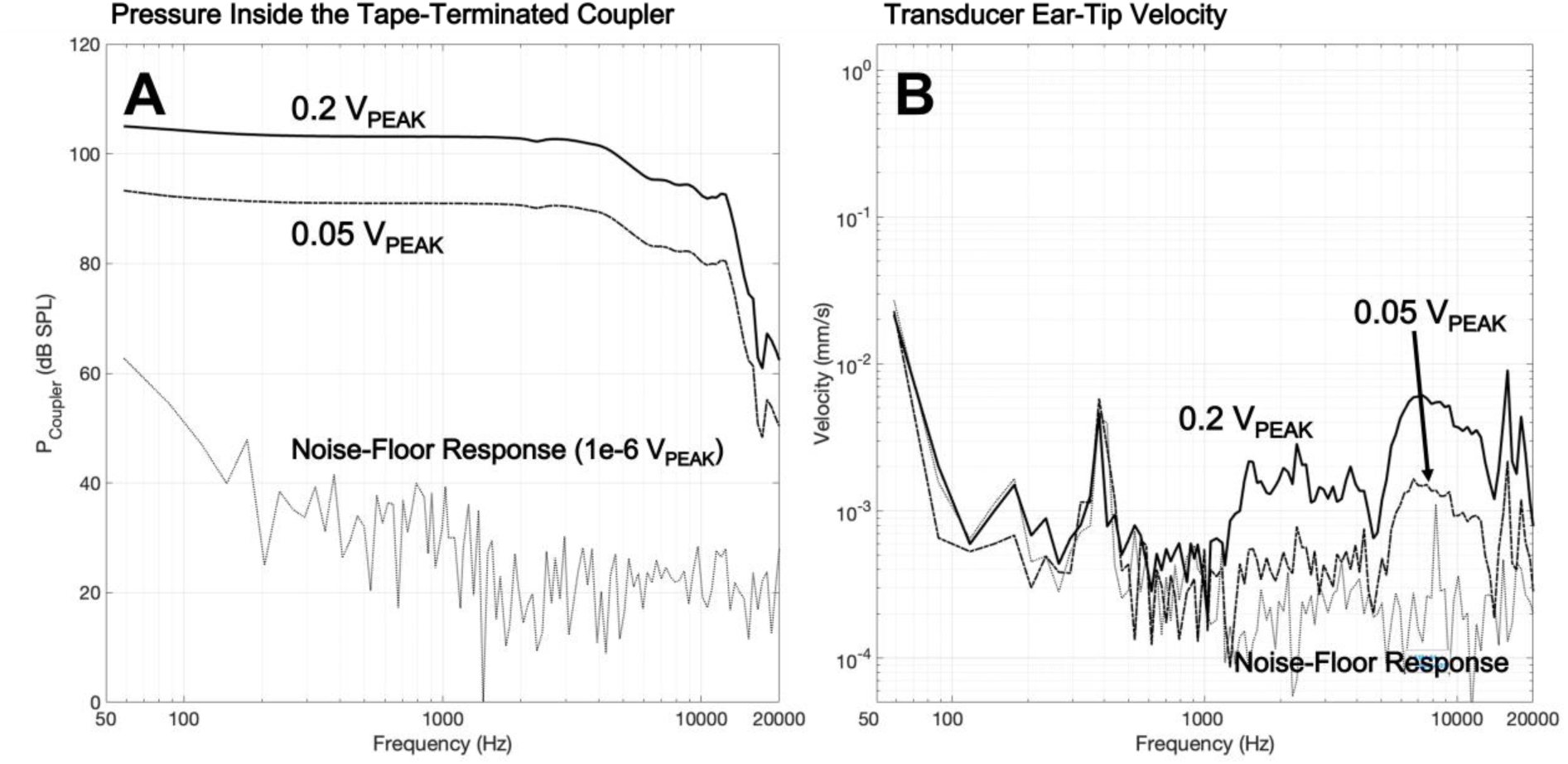
Tape-terminated audio-band sine-sequence measurements of the output pressure in the coupler. (A) and the transducer’s 3D-resultant ear-tip velocity (B), for Cypress transducer 1b with a piece of tape terminating the coupler’s foam ear tip. The transducer was stimulated with a sequence of tones from 59 Hz to 20 kHz with an amplitude of either 0.2 VPEAK or 0.05 VPEAK for the driven responses, or 1e-6 VPEAK to record the noise-floor response. The responses to each tone were synchronously averaged 10 times.

With 0.2 V_PEAK_ tones, the pressure (Fig. 3A) inside the coupler ranged between 105 and 102 dB SPL in the 59 Hz to 4 kHz frequency range, and decreased at higher frequencies, where it dropped to about 93 dB SPL by 12 kHz. Above 12 kHz, the pressure decreased rapidly and fell to about 65 dB SPL near 18 kHz. The pressure measurements with 0.05 V_PEAK_ tones were nearly identical in shape, but were reduced by a factor 4 (about 12 dB lower), indicating that the cavity pressure exhibited a linear relationship to the input voltage. The microphone noise-floor response (measured with an input voltage of 1e-6 V_PEAK_) decreased from about 62 dB SPL at 59 Hz to about 20 dB SPL at 2 kHz and above. The driven pressure responses were well above the microphone noise-floor response at all frequencies measured (by at least 20 dB and typically much more).

Vibration measurements were made at the transducer ear tips and the transducer body. We report the results from the transducer ear tip, because this would be the direct point of contact with the ear canal (without the coupler). The 3D resultant vibrations for the transducer ear tip are shown in Fig. 3B. In comparison to the pressure response within the coupling fixture (Fig. 3A), the vibration response looks very different. Whereas the pressure exhibits a low-pass response in that it passes frequencies below about 12 kHz and attenuates higher frequencies, the vibrations of the coupling fixture exhibit more of a high-pass response. For the 0.2 V_PEAK_ stimulus, the velocity was within the noise-floor response below about 1 kHz, but above that it increased with some fluctuations from about 0.0005 mm/s around 1 kHz to a peak of nearly 0.01 mm/s at 15.9 kHz. The velocity decreased linearly with the change of stimulus level to 0.05 V_PEAK_. The noise-floor response increased below about 120 Hz, reaching a peak near 59 Hz. There is a spike in the noise-floor response around 380 Hz, or 410 Hz in some of the measurements, which may have been related to building vibrations.

As noted above, we also measured vibrations of the transducer body. In the audio band, the velocity of the transducer body is similar to that of the transducer ear tip. However, at the operating frequency (98.847 kHz) there is significant vibration attenuation at the ear tip relative to the transducer body, but the *f*_o_ vibration at the ear tip is not completely eliminated (SI Slides 82–84). At the carrier frequency (197.695 kHz) there was near elimination of vibrations at the ear tip relative to the transducer body. These coupler measurements help us to better interpret the V_PRM_ results reported below, and show the importance of having a well-designed ear tip to reduce the transmission of ultrasonic frequencies to the cochlea through the BC pathway due to mechanical contact with the device. Likewise, they show the importance of minimizing the vibration of the transducer housing in order to reduce ultrasonic transmission into the hands or parts of the external ear that come into contact with the transducer housing.

### Middle-ear transmission in the audio band

Figure 4A presents stepped-tone measurements from 59 Hz to 20 kHz of the ear-canal pressure (P_EC_) produced by the device (Transducer 1b) within the coupler, with the transducer ear tip inserted into one end of the coupler and the coupler ear tip inserted into the ear canal of the specimen (see Fig. 2B), for a stimulus voltage of 0.2 V_PEAK_. These results are comparable to the tape-terminated coupling-fixture pressure measurements of Fig. 3A, but exhibit slightly more variability with frequency. Notably, Ear8 exhibits a nearly flat pressure profile below 1 kHz, whereas the pressure responses for Ear7 and Ear9 decrease at lower frequencies. This difference likely stems from a better seal between the transducer ear tip and the ear-canal opening of Ear8, whereas for the other ears there was likely a slight leak. Despite this variation, the SNR consistently remained above 30 dB throughout the entire plotted frequency range.

**Figure 4:**
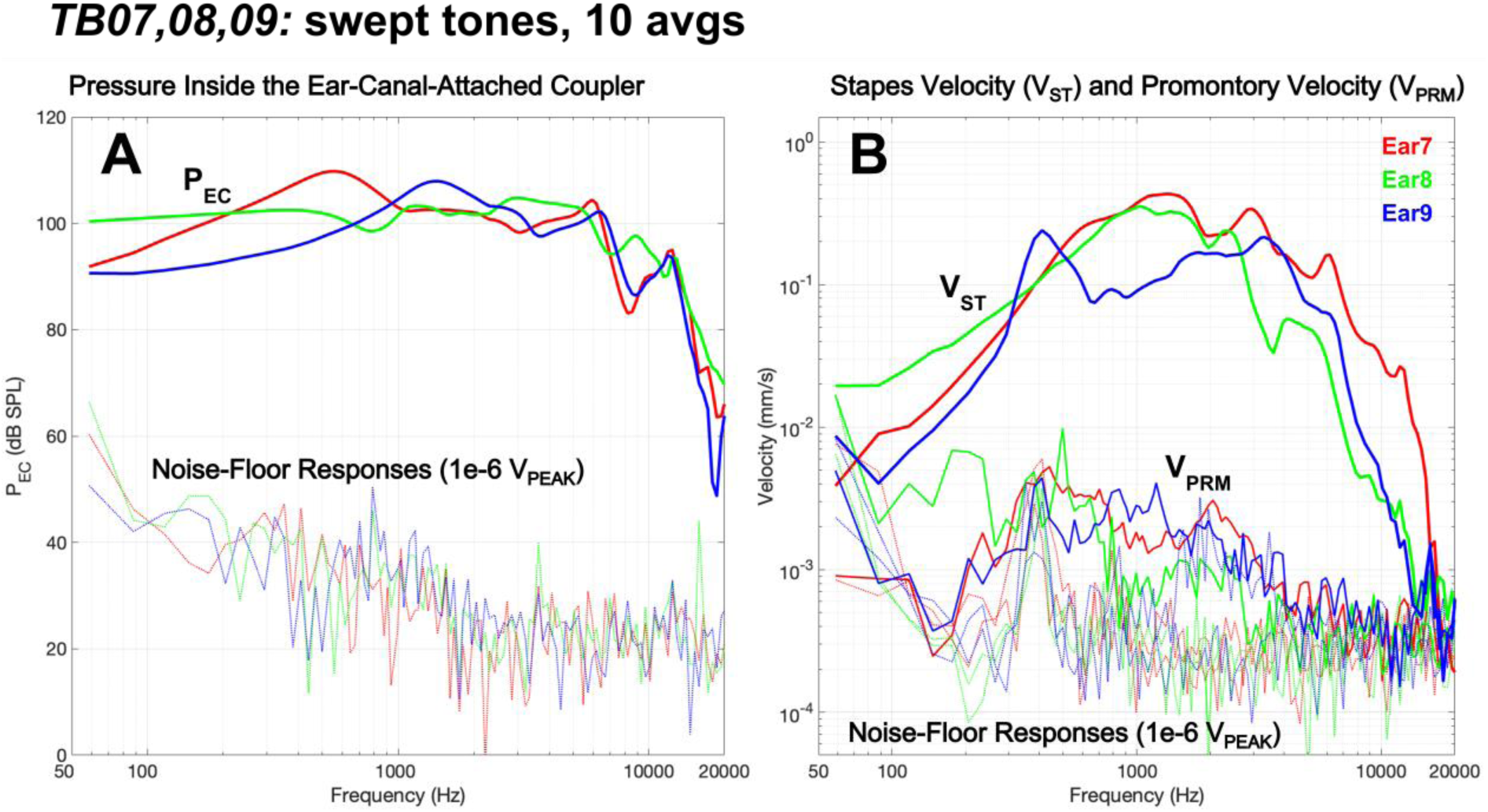
Temporal-bone audio-band sine-sequence measurements of. (A) the pressure inside the coupler which was attached to the ear canal (PEC), and (B) the 3D-resultant velocity of the stapes head (VST) or bony cochlear promontory (VPRM). Responses from Ear7 (red), Ear8 (green) and Ear9 (blue) are shown. The stimulus voltage was 0.2 VPEAK and the responses to each tone were synchronously averaged 10 times. The noise-floor responses are shown with dotted lines.

**Figure 5:**
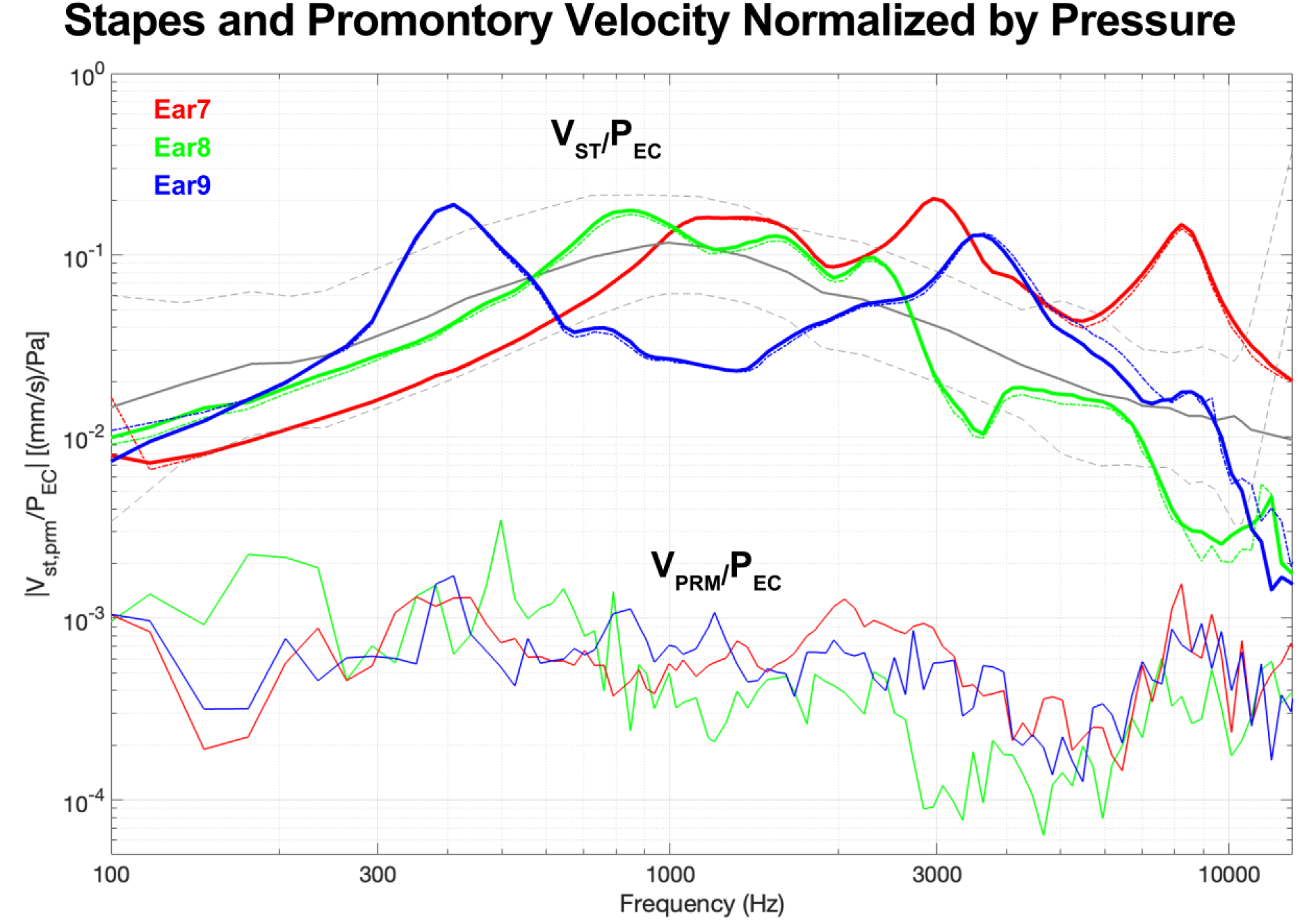
Middle-ear VST/PEC and VPRM/PEC audio-band sine-sequence transfer functions for Ear7–Ear9. The VST/PEC responses are shown for 0.2 VPEAK (solid lines) and 0.05 VPEAK (dash-dot lines) stimuli, whereas for VPRM/PEC only the 0.2 VPEAK responses are shown. The grey lines show, for comparison, the VST/PEC mean (solid) and ± 95% confidence intervals (dashed) published as part of an ASTM standard [22].

Figure 4B illustrates the 3D-resultant velocity of the stapes head (V_ST_) and cochlear promontory (V_PRM_) for Ear7–Ear9, corresponding to the respective P_EC_ responses shown in Fig. 4A. V_ST_ exhibits an upside-down U-shaped pattern where its magnitude increases with frequency, reaches a peak between 0.1 and 0.3 mm/s in the 0.5 to 3 kHz mid-frequency region, and subsequently decreases as the frequency continues to rise. V_PRM_ also exhibits an upside-down U-shaped pattern, but its magnitude is significantly lower than that of V_ST_ by a factor of 10–30 times (20-30 dB) in the same mid-frequency region.

Velocity noise-floor responses are also shown for the stapes head (thinner solid lines) and cochlear promontory (thinner dotted lines). The V_ST_ SNR typically ranges from 30 to 40 dB in the mid-frequency region and at least 10 dB at lower and higher frequencies. In contrast, the V_PRM_ magnitude barely exceeds the noise-floor response in the mid-frequency region. Notably, all velocity noise-floor responses exhibit a local spectral peak in the 400 Hz region.

### Normalized VST/PEC and VPRM/PEC responses

To determine if these temporal bones were normal, we calculated the velocity to ear-canal-pressure transfer functions (V_ST_/P_EC_ and V_PRM_/P_EC_) from 100 Hz to 12 kHz, where the V_ST_ SNR was greater than 10 dB (Fig. 4). For each ear, we calculated V_ST_/P_EC_ for both 0.2 and 0.05 V_PEAK_ drives, and the two results (colored solid and dashed lines) were nearly identical, indicating that both the transducer and the ME responses were linear. For comparison, we included the mean and 95% confidence intervals of the V_ST_/P_EC_ transfer function (gray solid and dashed lines) as part of an ASTM standard, which was derived from measurements conducted in various laboratories [22]. Nine of those laboratories employed LDV measurements, and three utilized video-stroboscopic measurements. Additionally, one laboratory employed a round-window sound-pressure estimate of stapes velocity in human cadaveric ears. A further methodological difference is that the velocity responses of the present study are the 3D-resultant magnitudes calculated from three orthogonal velocity components, which produces the maximum velocity amplitude in 3D space at each frequency regardless of how the motion happens to relate to the measurement angle. Our V_ST_/P_EC_ measurements in the three ears were generally within the 95% confidence intervals of the ASTM standard, which suggests that the baseline transfer-function measurements were obtained from normal ears without ME pathologies.

Figure 5 also illustrates V_PRM_/P_EC_, which was 30-40 dB lower than V_ST_/P_EC_ in the mid-frequency range, and at least 6 dB lower near 100 Hz and above 8 kHz. This suggests that, in the audio band, the primary route for sound transmission to the cochlea for the sound-from-ultrasound device under test was through the ossicular chain (the AC pathway). The relatively low magnitude of V_PRM_/P_EC_ implies poor mechanical coupling to the cochlea via BC pathways, at least in the audio-band frequencies. We next turn our attention to understanding the ultrasonic pressures emitted by the device and how they affect the spectra of P_EC_, V_ST_, and V_PRM_.

### Example of a Spectral Response

To investigate the ultrasonic emissions of the device, multiple repeated FFTs (N_FFT_=16,384 points) were taken of the outputs from the microphone and 3D LDV as the earbud device and its associated electronics responded to a single-tone input signal from SyncAv. Each FFT was converted to a single-sided frequency spectrum from 0 to 240 kHz with a 29.297 Hz frequency resolution, for which the magnitude of each frequency component was scaled by 2/N_FFT_ to match the amplitude of the corresponding real signal. Because the ultrasonic artifacts produced by the device were asynchronous in relation to SyncAv, we were not able to improve the SNR through time-domain synchronous averaging without destructively interfering with the signals of interest. Instead, we collected 10–20 separate repetitions of each measurement and averaged their spectral magnitudes as a way of identifying the consistently repeatable spectral components of the device’s output signature over the noisy components with randomly varying amplitudes. An example set of P_EC_ spectral responses from Ear9 with a 0.2 V_PEAK_, ∼1 kHz drive tone (corresponding to an exact spectral frequency of 0.99609375 kHz) is shown in Fig. 6A, with individual repetitions shown as colored lines and the magnitude average shown in black. To minimize crowding, only 10 repetitions are shown for all measurements. To place greater emphasis on ultrasonic frequencies, the results are plotted on a linear frequency axis.

**Figure 6:**
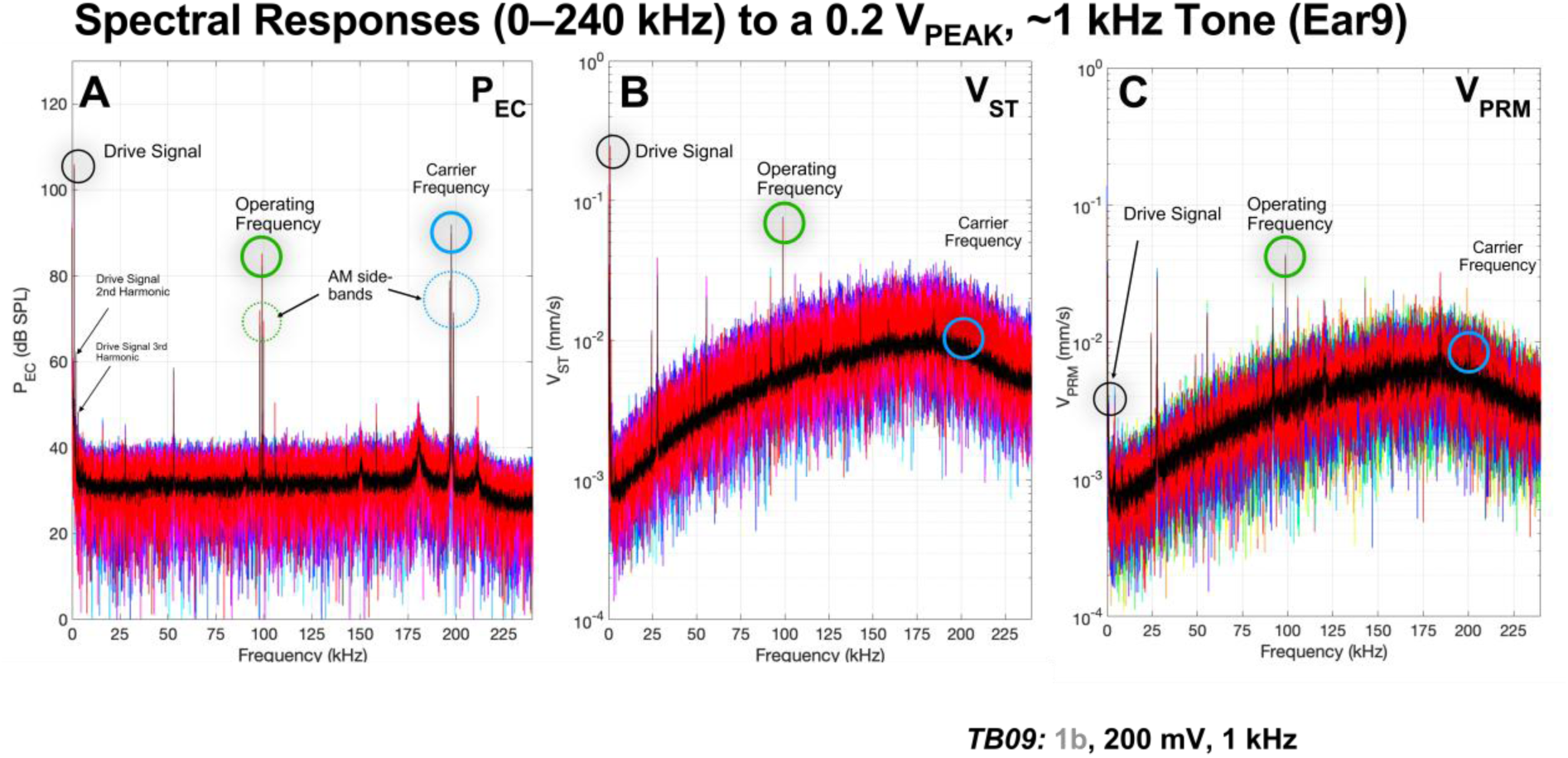
An example set of spectral-response magnitudes when driven by a single tone (0.2 VPEAK, ∼1 kHz; Ear9), showing all spectral frequencies on a linear frequency axis from 0 to 240 kHz, for: (A) PEC, (B) VST, and (C) VPRM. The colored lines show individual repeat measurements (with no averaging) and the black lines in the foreground show the magnitude average of all repeats in the given panel (N=20 for PEC and VST and N=10 for VPRM). The spectral spikes at the drive-tone frequency (∼1 kHz), operating frequency (∼100 kHz), and carrier frequency (∼200 kHz) are circled.

For the 0.2 V_PEAK_, ∼1 kHz stimulus signal, the P_EC_ magnitude from the averaged spectrum (black) at the ∼1 kHz primary stimulus frequency was approximately 105 dB SPL (see also Fig. 4A). The level at the second harmonic (∼2 kHz) was 62 dB SPL, which is about 0.7% of the primary level, and the level at the third harmonic (∼3 kHz) was 48 dB SPL, which is about 0.1% of the primary level.

We also observed two distinct sidebands centered around the 98.8477 kHz operating frequency *f*_o_ of the transducer. The lower sideband occurred at 97.8516 kHz and the upper sideband occurred at 99.8438 kHz. These three peaks collectively form an amplitude modulation (AM) signal: *f*_o_ ± *f*_input_, where *f*_input_ for the ∼1 kHz input in this case is actually 0.99609375 kHz (as mentioned above). The average pressure at the operating frequency was 82 dB SPL.

Similarly, we observed another AM signal centered at 197.695 kHz at the carrier frequency *f*_c_, which had an output level of 92 dB SPL. The carrier signal was accompanied by a lower sideband at 196.699 kHz and an upper sideband at 198.691 kHz (*f*_c_ ± *f*_input_). Both the operating and carrier frequencies differ from their respective sidebands by ∼0.996 kHz, which corresponds to the frequency of the audio-band input signal *f*_input_. It should be noted that the carrier frequency with its side bands as a set are not harmonics of the operating frequency and its side bands. Only the center frequencies are harmonically related.

The (spike-adjacent) NF surrounding each AM-signal spike in the magnitude-averaged P_EC_ spectrum (Fig. 6A, black) was usually around 32 dB SPL (as described earlier, calculated by averaging the 6 spectral bins surrounding a given spike: 3 on either side), which is considerably lower than the levels of the spikes themselves (at the stimulus-drive frequency, operating frequency, carrier frequency, and the four associated sidebands).

For the 0.2 V_PEAK_, ∼1 kHz stimulus tone, the V_ST_ magnitude from the averaged spectrum at the ∼1 kHz primary stimulus frequency was 0.13 mm/s for Ear9 (Fig. 6B, black). The level at the second harmonic was 0.0018 mm/s, which is approximately 1.4% of the primary magnitude, and the level at the third harmonic was near the NF level. At the operating frequency, the V_ST_ spike was above the NF at about 0.049 mm/s, but its AM side bands were ∼0.008 mm/s, just above the measured NF. V_ST_ at the carrier and its two sideband frequencies were within the spectral noise.

The corresponding V_PRM_ magnitude from the averaged spectrum (Fig. 6C, black; based on 10 repetitions rather than 20) was 0.0028 mm/s at the ∼1 kHz primary frequency, and its harmonics were indistinguishable from the surrounding spectral noise. At the operating frequency, the V_PRM_ magnitude was approximately 0.042 mm/s, and its AM side bands were 0.010–0.012 mm/s, which were above the noise level (∼0.004 mm/s). The V_PRM_ magnitudes at the carrier frequency and its two sidebands (*f*_c_ ± *f*_input_) were all in the noise (∼0.006 mm/s).

Note that the spectral noise floor of the velocity measurements generally tends to increase with frequency. With an LDV, only displacement noise can be white noise with a flat frequency spectrum. Consequently, velocity noise, theoretically increases proportionally to frequency. As a result, velocity resolution is frequency-dependent, while displacement resolution remains independent (with the exception of very low frequencies due to 1/f noise) [23]. The SNR therefore degrades as the displacement amplitude decreases with frequency.

### Pressure and velocity artifacts beyond the audio band

As is evident from the pressure and velocity spectral plots, several spurious spectral spikes were present apart from the stimulus-drive frequency, its harmonics, and the known device-related ultrasonic frequencies and their side bands (Fig. 6). Since we had not previously conducted human cadaveric temporal bone measurements beyond 20 kHz in our laboratory, we aimed to identify the sources of these additional spectral spikes. Specifically, we sought to determine whether they were related in any way to the device under test, or if they could be traced to electrical, mechanical, or acoustic artifacts from other equipment or from the measurement environment. To gain insights into this, we systematically turned off key components in the signal chain, as depicted in Fig. 1, and recorded the pressure in the coupler and 3D velocity of the transducer body (see Fig. 1 inset) after each change. A detailed account of these measurements is provided in the supplementary information (SI), but a concise summary is as follows:

• The 3D Carriage Controller generates three spikes that are electrically detected, primarily on the pressure channel. These spikes correspond to a 52.881 kHz fundamental, and its second (105.762 kHz) and third (158.643 kHz) harmonics. Occasionally peaks at similar frequencies may be observed in the velocity response.
• Turning on the 3D LDV introduces several spikes in both the pressure and velocity channels, and also raises the noise floor on the velocity channels.
  ○ The spikes in the pressure channel appear at 16.3, 27.8, 111.8, 143.2, 211.6, and 215.5 kHz
  ○ The spikes in the velocity channels appear at 16.4, 24.4, 27.8, 48.9, 83.5, 92, 143.2, and 184 kHz
• The xMEMS transducer introduces high-level spikes at the operating and carrier frequencies, which is expected. Additionally, it generates shorter, wider peaks around 180 kHz, primarily observed in the pressure measurements. The Ear measurements also reveal these spikes.

In summary, the majority of the spurious spikes were identified as artifacts caused by the 3D Carriage Controller and 3D LDV that have no relation to the device under test, so we ignore those for the purposes of the current study.

### Summarizing the relationships between airborne ultrasonic emissions from the device under test and the resulting middle-ear velocities

One of our primary objectives was to determine the extent to which the ear-canal pressure generated by the MEMS device under test produces ME vibrations that could potentially adversely affect the delicate structures within cochlea, at the: 1) audio-band stimulus frequency, 2) ultrasonic operating frequency, and 3) ultrasonic carrier frequency. While a smaller amplitude of ME vibration is likely to be less harmful, it is unclear how the structures within the cochlea might or might not respond to particular ultrasonic frequencies. Our focus is on identifying the extent to which vibrational inputs to the cochlea are produced due to the airborne ultrasonic sound and bone-conducted vibrations emitted by the device under test. The vibrational inputs being tested consist of V_ST_ and V_PRM_ because of their respective relevance to the AC and BC pathways of sound transmission to the cochlea.

Toward this end, we extracted P_EC_, V_ST_, and V_PRM_ data points at the three frequencies of interest (i.e., the stimulus-signal, operating, and carrier frequencies) across the full set of individual tone-driven spectral measurements made in Ear7–Ear9. For each of the extracted points, we also calculated the NF surrounding the central frequencies.

Figures 7–10 provide illustrative summaries of a subset of the extracted data points for two representative drive frequencies (∼1 kHz: Figs 7, 8; and ∼10 kHz: Figs 9, 10) and both 0.2 V_PEAK_ (Figs 7 and 9) and 0.05 V_PEAK_ (Figs 8 and 10) drive voltages, in a manner that highlights the relationship between each ear-canal (EC) pressure spike and the corresponding middle-ear velocity spike measured at the same time, as well as the degree to which each rises above the surrounding NF. As mentioned previously, the spectral noise tends to increase by 10–25 dB at the ultrasonic frequencies comparing with the audio frequencies.

**Figure 7:**
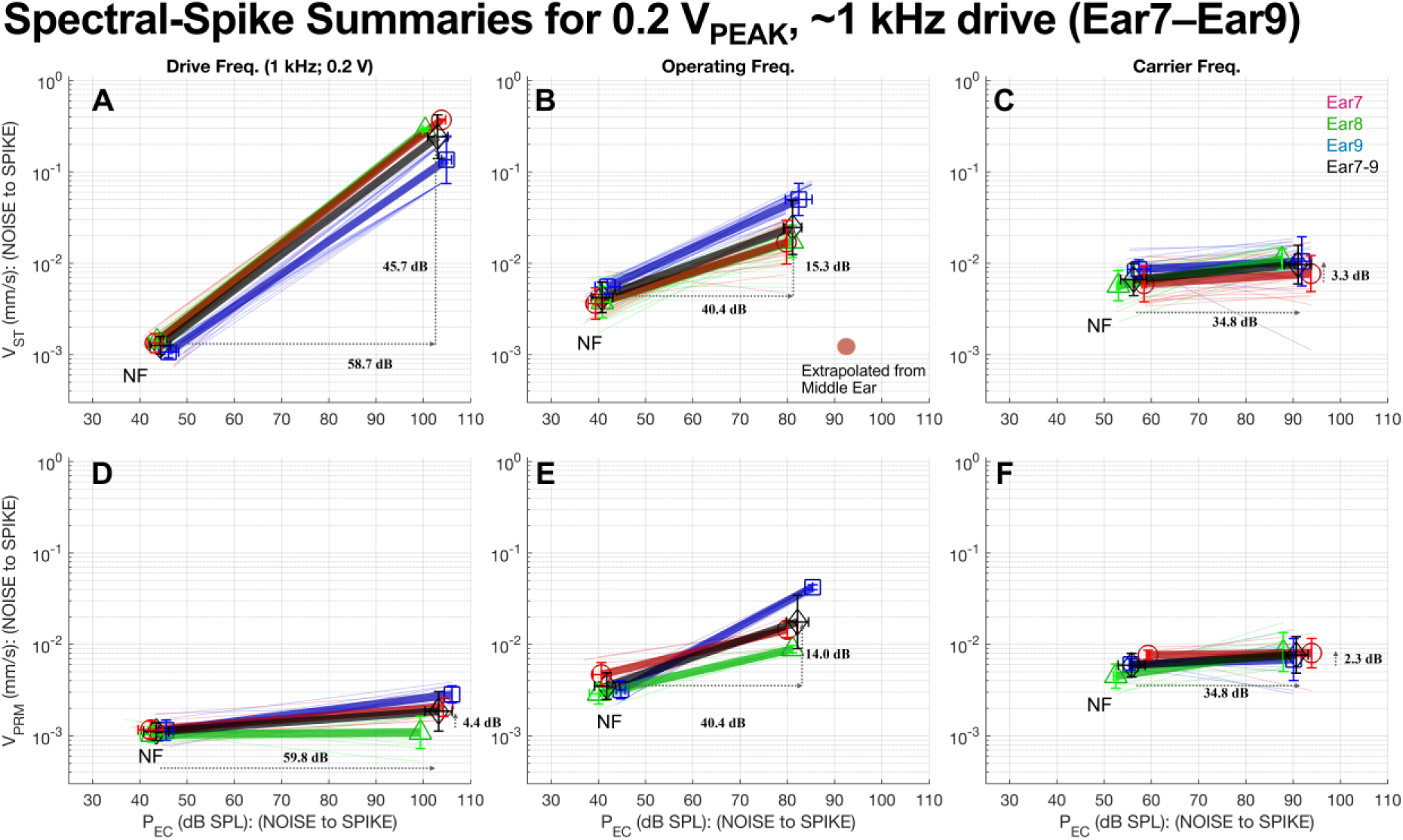
Summaries of the spectral spikes when driven by a single tone at 0.2 V_PEAK_ and ∼1 kHz for Ear7 (red, circles), Ear 8 (green, triangles), and Ear9 (blue, squares), along with the full ensemble combined across the three ears (black, diamonds). Each panel features P_EC_ on the horizontal axis and either V_ST_ (top row: A, B, and C) or V_PRM_ (bottom row: D, E, and F) on the vertical axis, and contains various line segments for which the leftmost (P_EC_, V_ST_)|_NF_ or (P_EC_, V_PRM_)|_NF_ point represents the spike-adjacent noise-floor-magnitude average (NF) and the rightmost (P_EC_, V_ST_)|_SPIKE_ or (P_EC_, V_PRM_)|_SPIKE_ point represents the spectral-spike magnitude. The first column (A and D) summarizes the spikes at the stimulus-drive frequency, the second column (B and E) summarizes the spikes at the ∼100 kHz operating frequency, and the third column (C and F) summarizes the spikes at the ∼200 kHz carrier frequency. Thin colored lines represent individual measurements from a given ensemble and symbols joined by thick translucent lines represent the mean for a given ensemble. Error bars extending from the symbols represent one standard deviation along each axis. Horizontal and vertical projections are shown for the black full-ensemble averages.

**Figure 8:**
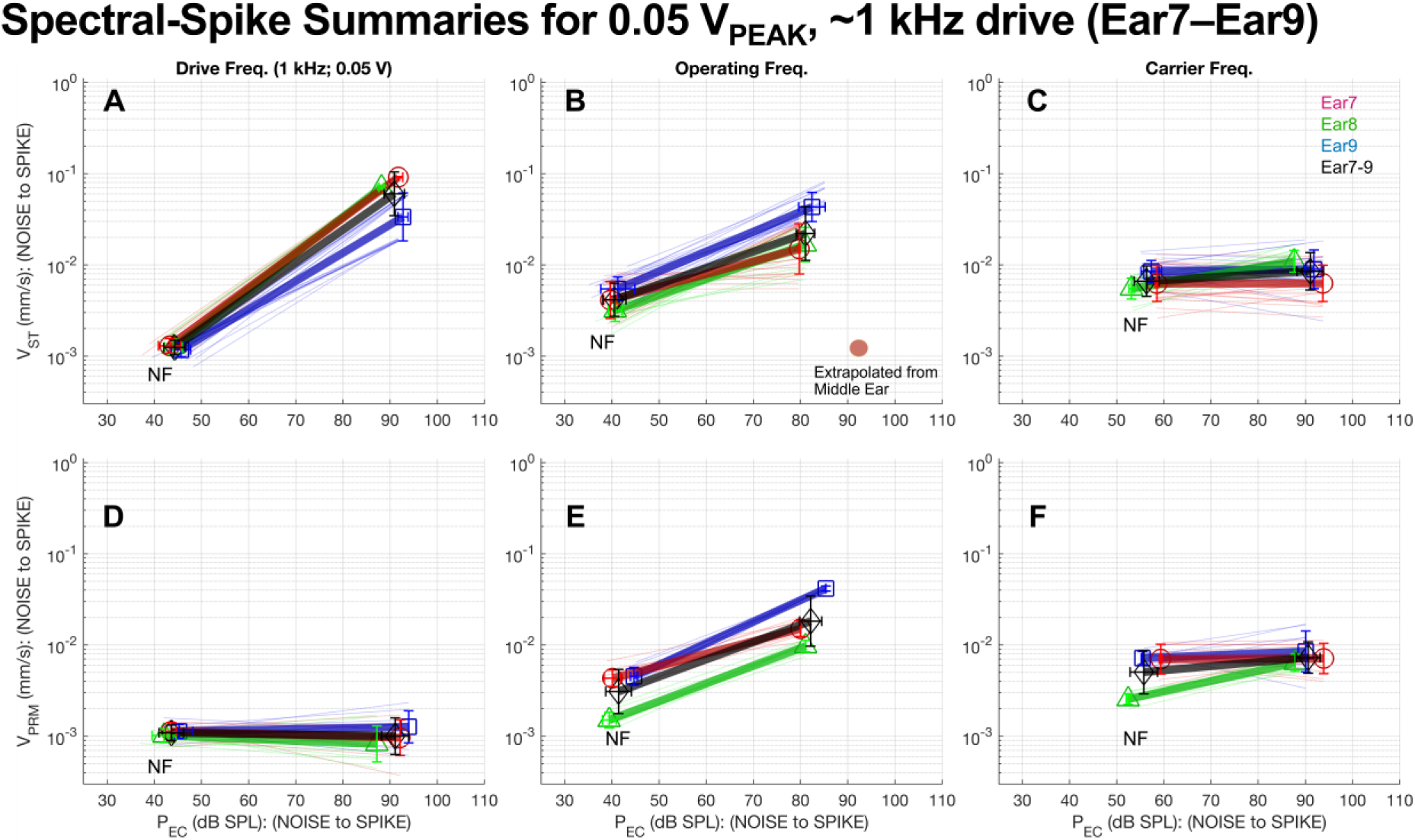
Summaries of the spectral spikes when driven by a single tone at 0.05 VPEAK and ∼1 kHz. See the caption to Fig. 7 for a detailed explanation.

**Figure 9:**
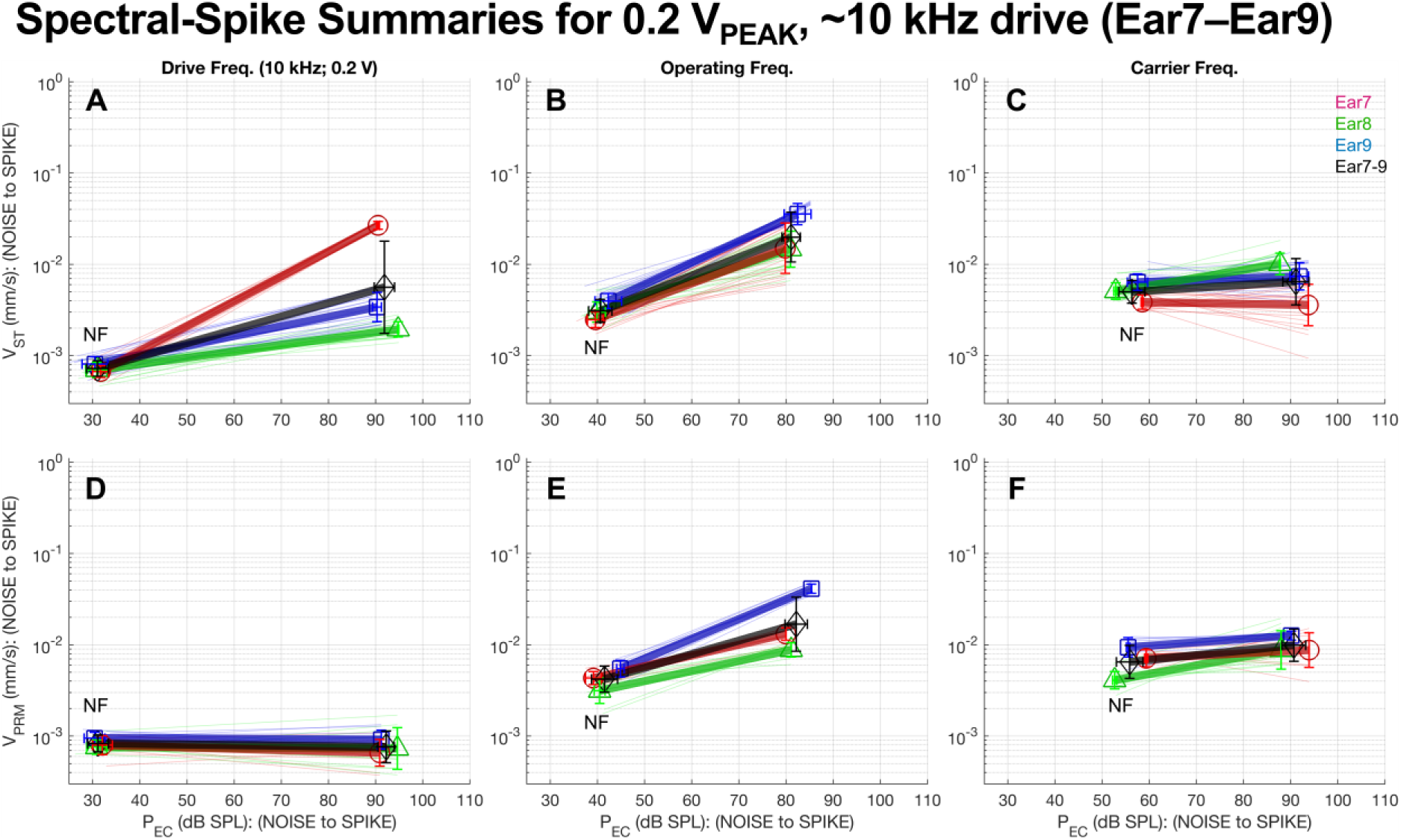
Summaries of the spectral spikes when driven by a single tone at 0.2 VPEAK and ∼10 kHz. See the caption to Fig. 7 for a detailed explanation.

**Figure 10:**
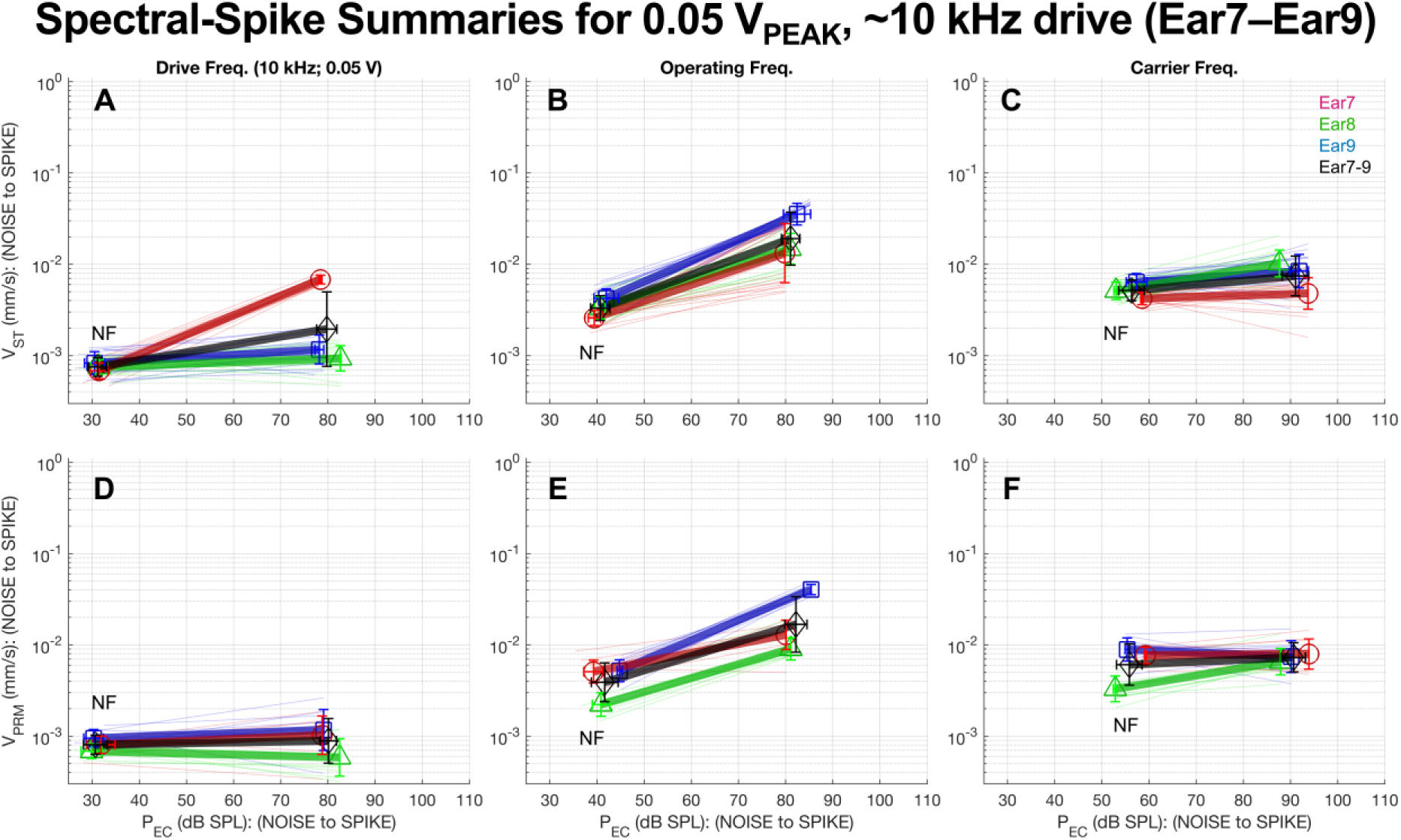
Summaries of the spectral spikes when driven by a single tone at 0.05 VPEAK and ∼10 kHz. See the caption to Fig. 7 for a detailed explanation.

Each pair of corresponding pressure and velocity amplitudes is represented as a line segment beginning with the pressure and velocity NF as the first pair of coordinates (P_EC_, V_ST_)|_NF_ or (P_EC_, V_PRM_)|_NF_ (for the top and bottom row of subpanels, respectively) and ending with the pressure and velocity spike amplitudes as the second pair of coordinates (P_EC_, V_ST_)|_SPIKE_ or (P_EC_, V_PRM_)|_SPIKE_ (also for the top and bottom row of subpanels, respectively). The horizontal projection of each line segment represents the height of the P_EC_ spike above the surrounding NF (in dB SPL), and the vertical projection represents the height of the V_ST_ spike (top row of subpanels) or V_PRM_ spike (bottom row of subpanels) above the surrounding noise (in mm/s).

If the slope of a given line segment is angled upward, then the P_EC_ spike corresponds to a velocity spike that rose above the surrounding spectral noise. On the other hand, if a given line segment is angled close to horizontally, then the P_EC_ spike does not correspond to a velocity that rose significantly above the surrounding spectral noise.

The three columns of subpanels in Figs 7–10 summarize the spikes at the drive frequency, operating frequency, and carrier frequency, respectively. The results from each specimen are shown in different colors (RGB), with red representing Ear7, green representing Ear8, and blue representing Ear9. Within each subpanel, individual spikes are represented as thin semitransparent line segments. The mean of each ensemble of (P, V) coordinates was also calculated, both for individual specimens and across specimens, and these are shown as colored symbols (Ear7=red circles, Ear8=green triangles, Ear9=blue squares, Ear7–9=black diamonds) joined by thick semitransparent lines of the same color. The velocity means were calculated on a log scale, and the pressure means were calculated on a linear scale using the dB SPL values. The vertical and horizontal error bars extending from the center of each symbol represent the standard deviation (STD), also calculated and plotted using the scaling appropriate for each axis.

#### Responses to a 1 kHz drive tone

Figure 7 presents the individual and statistically summarized results as driven by a 0.2 V_PEAK_, ∼1-kHz tone, for Ear7 (red), Ear8 (green), Ear9 (blue), and the combined **Ear7–9** ensemble (black). At the drive frequency (Fig. 7A), the (P_EC_, V_ST_)|_PEAK_ points for all 3 Ears fall in the upper right-hand corner of the subpanel (respective means for Ear7, Ear8, Ear9, and **Ear7–9**: 103.8, 100.4, 104.9, and **103.0** dB SPL; 0.371, 0.289, 0.136, and **0.242** mm/s), and the corresponding (P_EC_, V_ST_)|_NF_ points are all situated in the lower left-hand corner of the subpanel (respective means: 43.2, 43.6, 46.1, and **44.3** dB SPL; 0.00134, 0.00140, 0.00107, and **0.00126** mm/s). Both the P_EC_ and V_ST_ spike means rose significantly above the NF means at the drive frequency, by: 60.6 (Ear7), 56.8 (Ear8), 58.9 (Ear9), and **58.7** (Ear7–9) dB SPL; and 48.8 (Ear7), 46.3 (Ear8), 42.1 (Ear9), and **45.7** (Ear7–9) dB, respectively.

In the three ears tested, we conducted test-retest measurements by taking 10 separate (P_EC_, V_ST_) measurements, removing the transducer, returning it to its original location, and making 10 additional measurements (for a total of 20 retests). As indicated in Figure 7, the STD values for Ear7 and Ear8 were relatively small, while Ear9 had a larger STD. The larger V_ST_ variability in Ear9 was caused in some way by the repositioning of the transducer after the first batch of 10 measurements, as the V_ST_ spikes from the first and second batches of measurements form two distinct groups.

At the operating frequency (Fig. 7B), the (P_EC_, V_ST_)|_SPIKE_ points (respective means: 79.8, 81.0, 82.4, and **81.1** dB SPL; 0.0170, 0.0174, 0.0500, and **0.0247** mm/s) generally do not extend as high or as far to the right of the subpanel as the corresponding drive-frequency points, and the (P_EC_, V_ST_)|_NF_ points (respective means: 39.4, 40.8, 41.9, and **40.7** dB SPL; 0.00362, 0.00377, 0.00552, and **0.00423** mm/s) lie a little further to the left and somewhat higher than the respective drive-frequency points. As mentioned previously, this indicates that the velocity NF is higher at the operating frequency than at the drive frequency and that the spike amplitudes at the operating frequency are not as high as at the drive frequency. Nonetheless, both the P_EC_ and V_ST_ spike means at the operating frequency still do rise above the NF means, by: 40.4 (Ear7), 40.2 (Ear8), 40.5 (Ear9), and **40.4** (Ear7–9) dB SPL; and 13.4 (Ear7), 13.3 (Ear8), 19.1 (Ear9), and **15.3** (Ear7–9) dB, respectively. The V_ST_ spike means at the drive frequency lie above those at the operating frequency, by: 26.8 (Ear7), 24.4 (Ear8), 8.7 (Ear9), and **19.8** (Ear7–9) dB.

At the carrier frequency (Fig. 7C), the (P_EC_, V_ST_)|_SPIKE_ points (respective means: 93.7, 87.6, 91.8, and **91.0** dB SPL; 0.00771, 0.0111, 0.0105, and **0.00978** mm/s) lie a little further to the right, do not reach as high, and exhibit more variability in the vertical direction, as compared to the corresponding spike points at the operating frequency. The carrier-frequency (P_EC_, V_ST_)|_NF_ points (respective means: 58.5, 53.0, 57.4, and **56.2** dB SPL; 0.00597, 0.00569, 0.00861, and **0.00665** mm/s) are shifted considerably to the right and are somewhat higher on average than the respective points at the operating frequency. The P_EC_|_SPIKE_ means at the carrier frequency still retain a reasonably high SNR, of: 35.3 (Ear7), 34.6 (Ear8), 34.4 (Ear9), and **34.8** (Ear7–9) dB SPL. However, the V_ST_|_SPIKE_ means at the carrier frequency rise very little above the NF means, by: 2.2 (Ear7), 5.8 (Ear8), 1.7 (Ear9), and **3.3** (Ear7–9) dB; and the spike and NF vertical STD ranges overlap in all three cases. Therefore, these measured V_ST_ “spikes” at the carrier frequency are considered to be within the averaged spectral noise.

For the case where V_PRM_ was measured at the drive frequency (Fig. 7D), the (P_EC_, V_PRM_)|_SPIKE_ points (respective means: 104.2, 99.4, 106.1, and **103.2** dB SPL; 0.00206, 0.00109, 0.00283, and **0.00185** mm/s) lie at a similar horizontal position, but are much lower vertically than the respective V_ST_|_SPIKE_ points, whereas the (P_EC_, V_PRM_)|_NF_ points (respective means: 42.2, 42.5, 45.6, and **43.6** dB SPL; 0.00116, 0.00103, 0.00115, and **0.00112** mm/s) are placed similarly to the respective (P_EC_, V_ST_)|_NF_ points. Each V_ST_|_SPIKE_ average at the drive frequency is higher than the corresponding V_PRM_|_SPIKE_ average by 45.1 (Ear7), 48.5 (Ear8), 33.6 (Ear9), and **42.3** (Ear7–9) dB. The SNRs for P_EC_ in subpanel D: 62.0 (Ear7), 56.8 (Ear8), 60.5 (Ear9), and **59.8** (Ear7–9) dB SPL; are similar to the corresponding values from subpanel A, as expected. However, the V_PRM_|_SPIKE_ points rise only a little above the noise at the drive frequency, by: 4.9 (Ear7), 0.47 (Ear8), 7.8 (Ear9), and **4.4** (Ear7–9) dB.

At the operating frequency for the V_PRM_ measurements (Fig. 7E), the (P_EC_, V_PRM_)|_SPIKE_ points (respective means: 80.0, 81.1, 85.4, and **82.2** dB SPL; 0.0145, 0.00886, 0.0422, and **0.0176** mm/s) mostly occur at similar horizontal positions as compared to the respective (P_EC_, V_ST_)|_SPIKE_ points in subpanel B, except for the Ear9 P_EC_|_SPIKE_ readings, which are shifted to the right by 2–3 dB in subpanel E. The (P_EC_, V_PRM_)|_NF_ points (respective means: 40.6, 40.0, 44.7, and **41.8** dB SPL; 0.00467, 0.00288, 0.00319, and **0.00350** mm/s) are in the same vicinity as the respective (P_EC_, V_ST_)|_NF_ points from subpanel B, but with the Ear9 pressure shifted to the right (see the next paragraph for an explanation), the Ear7 velocity shifted slightly up, and the Ear8 and Ear9 velocities shifted slightly down. The P_EC_ and V_PRM_ spike means at the operating frequency rose above the NF means by: 39.4 (Ear7), 41.1 (Ear8), 40.6 (Ear9), and **40.4** (Ear7–9) dB SPL; and 9.9 (Ear7), 9.7 (Ear8), 22.4 (Ear9), and **14.0** (Ear7–9) dB, respectively. The mean V_PRM_ spike amplitudes at the operating frequency are significantly higher than the corresponding V_PRM_ spikes at the drive frequency, by: 17.0 (Ear7), 18.2 (Ear8), 23.5 (Ear9), and **19.5** (Ear7–9) dB. The V_PRM_ spike means at the operating frequency are also comparable to the V_ST_ spike means at the operating frequency, with V_ST_ higher than V_PRM_ by a mere: 1.4 (Ear7), 5.8 (Ear8), 1.5 (Ear9), and **2.9** (Ear7–9) dB.

The operating-frequency measurements suggest that, at the ∼100 kHz operating frequency, the device under test produces significant vibrations, with similar amplitudes, of both the stapes and the cochlear promontory. This occurred even though care was taken during the experiments to reduce the amount of direct BC transmission from the transducer to the temporal bone by using two foam ear tips to: 1) isolate the transducer from the coupling fixture, and 2) isolate the coupling fixture from the ear canal. If such steps had not been taken, the amount of bone vibration recorded at the operating frequency might have been higher.

The average SNR for the V_PRM_ measurements at the operating frequency was 14 dB, which is 9.6 dB higher than the V_PRM_ SNR for the drive frequency, even though the average spectral noise is higher near the operating frequency. This implies that the V_PRM_ spikes at the operating frequency are significant and should be considered equally as important as the V_ST_ spikes at the operating frequency. In other words, at the operating frequency the distinction between V_ST_ and V_PRM_ becomes less important, as both contribute significant vibrational inputs to the cochlea. It is unclear what amplitude should be considered safe when it comes to cochlear exposure. Knowing that the V_ST_ amplitudes are less than those caused by the drive signal is better than if it were equal or higher, but how the cochlea could be affected by differential inputs at this ultrasonic frequency is not known or understood. The higher V_PRM_ amplitude suggests that BC vibration exposure can be higher.

The rightward shift of many of the individual P_EC_ readings in Ear9 (i.e., 10 of the 20 readings in subpanel B and all 10 of the readings in subpanel E) corresponds to the removal and reinsertion of the ear tip into the ear canal after the first set of 10 (P_EC_, V_ST_) measurements were made. That reinsertion must have produced a tighter seal, as it corresponds to a boost in pressure for all of the measurements that were made afterward, including the only set of 10 (P_EC_, V_PRM_) measurements in subpanel E and the second set of 10 (P_EC_, V_ST_) measurements in subpanel B. Because of the order in which the various experimental variables were changed over the course of the Ear9 experiment (i.e., the drive voltage, the drive frequency, and whether V_ST_ or V_PRM_ is being measured using the 3D LDV), the Ear9 P_EC_ readings fall into one of two distinct groups depending on whether they were recorded before or after the ear tip reinsertion, which is especially evident in the full dataset for the operating frequency shown in Fig. 11.

**Figure 11:**
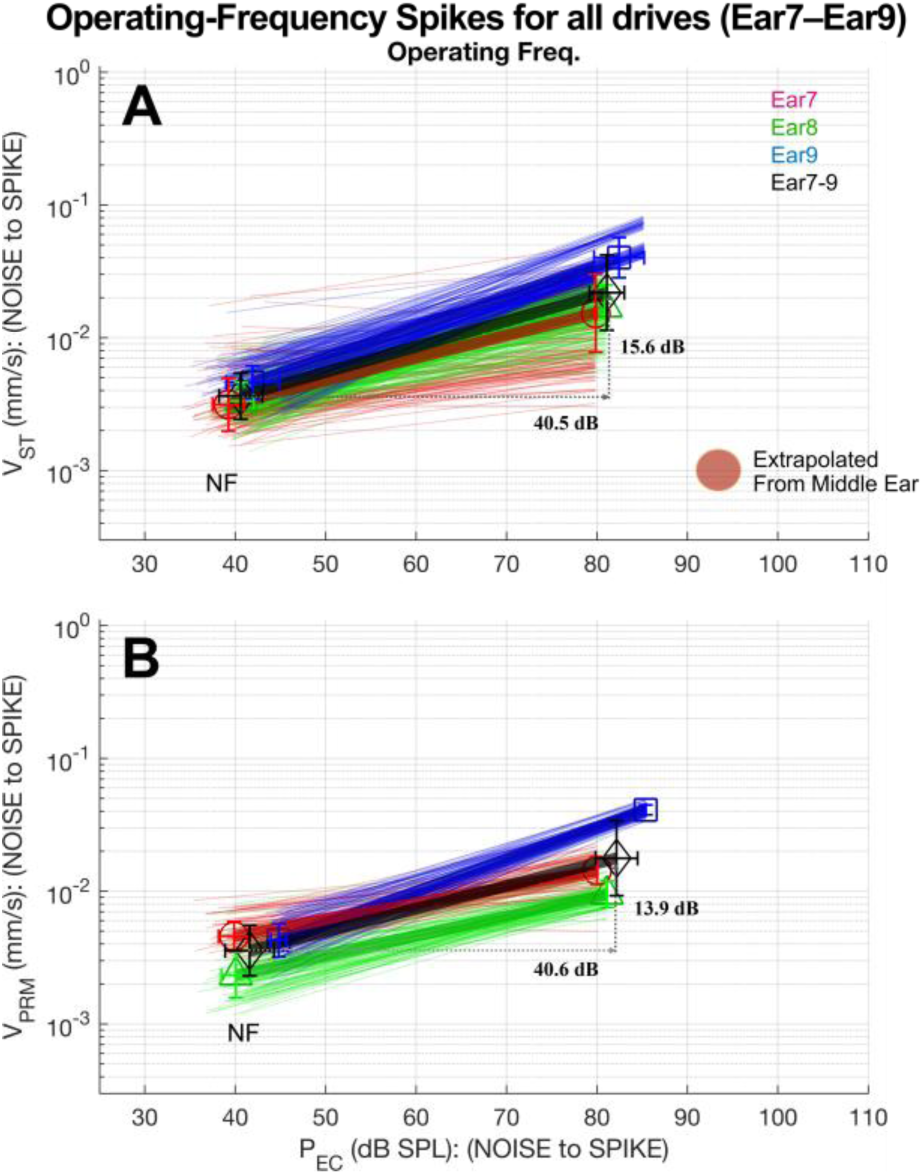
Summary of the operating-frequency spectral spikes across all stimulus-drive amplitudes (0.2 and 0.05 VPEAK) and frequencies (∼0.5, ∼1, ∼3, ∼10, and ∼20 kHz) for the Ear7 (red, circles), Ear8 (green, triangles), Ear9 (blue, squares), and Ear7–Ear9 (black, diamonds) ensembles. The VST summaries are shown in (A) and the VPRM summaries are shown in (B). Assuming the middle ear vibration decreases with frequency by about 20 dB/decade, the extrapolated stapes velocity at the operating frequency (∼100 kHz) at 1Pa (94 dB SPL) would be about 0.001 (mm/s), as shown in brown circle in (A).

At the carrier frequency (Fig. 7F), the (P_EC_, V_PRM_)|_SPIKE_ points (respective means: 93.9, 87.9, 90.1, and **90.6** dB SPL; 0.00803, 0.00823, 0.00683, and **0.00767** mm/s) lie in the same vicinity as the corresponding (P_EC_, V_ST_)|_SPIKE_ points (Fig. 7C), and the (P_EC_, V_PRM_)|_NF_ points (Fig. 7F) are also similarly placed (respective means: 59.3, 52.5, 55.6, and **55.8** dB SPL; 0.00766, 0.00449, 0.00602, and **0.00592** mm/s) compared to the corresponding (P_EC_, V_ST_)|_NF_ points (Fig. 7C). As for (P_EC_, V_ST_) in Fig. 7C, the P_EC_ spike means in Fig. 7F rise above the noise, by: 34.6 (Ear7), 35.4 (Ear8), 34.4 (Ear9), and **34.8** (Ear7–9) dB SPL; while the V_PRM_ spike means in Fig. 7F barely rise above the NF means, by: 0.41 (Ear7), 5.3 (Ear8), 1.1 (Ear9), and **2.3** (Ear7–9) dB. The V_PRM_ spikes all have overlapping STD error bars with those of the NF at the carrier frequency, such that the V_PRM_ “spikes” at the carrier frequency are also considered to be within the measurement noise.

The measurements at ∼1 kHz were repeated with a 0.05 V_PEAK_ stimulus voltage (Fig. 8). As anticipated, the (P_EC_, V_ST_)|_SPIKE_ means at the stimulus-drive frequency (Fig. 8A; respective means: 91.8, 88.2, 92.8, and **90.9** dB SPL; 0.0916, 0.0710, 0.0336, and **0.0602** mm/s) decreased by about a factor 4, or ∼12 dB, relative to the corresponding measurements at 0.2 V_PEAK_ (Fig. 8A), with respective P_EC_ differences of –12.1, –12.2, –12.1, and **-12.1** dB; and respective V_ST_ differences of –12.1, –12.2, –12.1, and **-12.1** dB. The reduced stimulus voltage left V_PRM_ at the drive frequency well within the NF (Fig. 8D). The responses at the operating frequency (Fig. 8B, E) and carrier frequency (Fig. 8C, F) were remarkably similar to those obtained with 0.2 V_PEAK_ stimulus tones (in the respective subpanels of Fig. 7). The average deviations between the 0.2 and 0.05 V_PEAK_ groups across the three specimens were less than 0.05 dB for P_EC_ and less than 1.1 dB for either velocity, at both the operating and carrier frequencies, which suggests that the amplitudes of the ultrasonic leakage at the operating and carrier frequencies, and the corresponding ME vibrations, are not at all dependent on the drive amplitude of the audio-band stimulus.

#### Responses to a 10 kHz drive tone

Figure 9 shows the results for a drive tone at ∼10 kHz with a 0.2 V_PEAK_ stimulus voltage. With some minor differences, the responses at the operating and carrier frequencies were nearly identical to those with a ∼1 kHz drive tone as shown in Fig. 7. The biggest difference was at the ∼1 vs. ∼10 kHz stimulus-drive frequency itself, where P_EC_ and V_ST_ with a ∼10 kHz drive tone decreased (Fig. 9A) by 13.3 (Ear7), 5.6 (Ear8), 14.6 (Ear9), and **11.2** (Ear7–9) dB SPL; and 22.8 (Ear7), 43.5 (Ear8), 32.0 (Ear9), and **32.7** dB; respectively. The associated P_EC_ and V_ST_ NF readings surrounding the ∼10 kHz spikes also decreased relative to ∼1 kHz, by: 11.5 (Ear7), 12.7 (Ear8), 15.6 (Ear9), and **13.3** (Ear7–9); and 6.0 (Ear7), 6.1 (Ear8), 2.5 (Ear9), and **4.9** (Ear7–9) dB; respectively. Notably, Ear7 had a significantly higher V_ST_ spike at ∼10 kHz than the other Ears (see also Fig. 5).

The measurements at ∼10 kHz were repeated with a 0.05 V_PEAK_ stimulus voltage. As shown in Figure 10, the responses at the operating frequency (Fig. 10B, E) and the carrier frequency (Fig. 10C, F) were remarkably similar to those obtained when 0.2 V_PEAK_ drive tones were used. As anticipated, V_ST_ at the stimulus-drive frequency decreased with the reduced drive voltage to levels close to the NF, except for Ear7, which remained above the NF in spite of its expected ∼12 dB decrease. V_PRM_ with a 0.05 V_PEAK_ drive was predominantly in the noise (Fig. 10D).

In summary, the pressure at the operating frequency remained remarkably constant across different stimulus-drive frequencies and voltages; and the corresponding V_ST_ and V_PRM_ spikes also remained remarkably independent of the stimulus-drive frequency or voltage, and both tended to remain above the NF. Similarly, the pressure at the carrier frequency also showed minimal variation across stimulus-drive frequency conditions, but the corresponding V_ST_ and V_PRM_ values were in the NF range at that frequency. The V_PRM_ spikes at the operating frequency were typically not much lower than the corresponding V_ST_ spikes, suggesting that both AC and BC factors may be contributing significantly to cochlear ultrasound exposure at the operating frequency.

These findings suggest that: 1) at the audio-band drive frequency, the primary route for sound transmission was through the AC pathway (as indicated by V_ST_) rather than through the BC pathway (as indicated by V_PRM_); 2) at the ∼100 kHz ultrasonic operating frequency, both the AC and BC ME-velocity indicators remain comparable and above the noise; and 3) at the carrier frequency, no significant AC or BC vibrations are discernible from the elevated NF environment near that frequency. Similar trends were also observed at the other tested drive frequencies of ∼0.5, ∼3, and ∼20 kHz (not shown).

Viewed as a whole, the measured pressure and velocity amplitudes at the operating frequency do not show any dependence on the voltage or frequency that happened to be assigned for the stimulus signal at any given time. For this reason, along with the fact that no significant velocity spikes at the carrier frequency were discernible from the NF, the entire set of measurements at the operating frequency (199–200 per specimen for V_ST_ and 100 per specimen for V_PRM_) were pooled together to analyze the overall ultrasonic vibrations measured at the operating frequency (described next).

#### Summary across ears at the operating frequency

Figure 11 provides a comprehensive summary at the operating frequency (for Transducer 1b) of the measured relationships between the P_EC_ and V_ST_ spikes (Fig. 11A) and between the P_EC_ and V_PRM_ spikes (Fig. 11B); across all stimulus-drive frequencies (∼0.5, ∼1, ∼3, ∼10, and ∼20 kHz) and voltages (0.2 and 0.05 V_PEAK_); for Ear7, Ear8, and Ear9 individually (red lines and circles, green lines and triangles, and blue lines and squares; respectively); and for the combined Ear7–9 ensemble (black lines and diamonds). A total of 200 measurements were acquired per specimen for the (P_EC_, V_ST_) pairs, except for Ear7, for which 199 measurements are shown because one was deemed faulty. The (P_EC_, V_PRM_) pairs for all three Ears consisted of 100 measurements each. As before, the mean and STD ranges are shown for each ear and for the combined Ear7–9 ensemble.

The overall P_EC_|_SPIKE_ mean±STD summaries from the (P_EC_, V_ST_)|_SPIKE_ ensemble of points at the operating frequency (Fig. 10A) were 79.9±0.1 (Ear7), 81.0±0.1 (Ear8), 82.4±2.8 (Ear9), and **81.1±1.9** (Ear7–9) [dB SPL]±[dB SPL]. As explained above, the larger variation of the Ear9 pressures was due to a change after removing and reinserting the ear tip into the ear canal of that specimen. The corresponding V_ST_|_SPIKE_ summaries from the same ensemble were 0.0153±5.6 (Ear7), 0.0170±3.4 (Ear8), 0.040±3.0 (Ear9), and **0.0218±5.7** (Ear7–9) [mm/s]±[dB].

The overall P_EC_|_NF_ summaries from the (P_EC_, V_ST_)|_NF_ ensemble of points at the operating frequency (Fig. 10A) were 39.2±1.8 (Ear7), 40.6±1.4 (Ear8), 41.9±2.9 (Ear9), and **40.6±2.4** (Ear7–9) [dB SPL]±[dB SPL]. The corresponding V_ST_|_NF_ summaries from the same ensemble were 0.00313±4.0 (Ear7), 0.00333±2.8 (Ear8), 0.00461±2.5 (Ear9), and **0.00364±3.5** (Ear7–9) [mm/s]±[dB].

The overall P_EC_|_SPIKE_ mean±STD summaries from the (P_EC_, V_PRM_)|_SPIKE_ ensemble of points at the operating frequency (Fig. 10B) were 80.0±0.1 (Ear7), 81.1±0.03 (Ear8), 85.4±0.03 (Ear9), and **82.2±2.3** (Ear7–9) [dB SPL]±[dB SPL]. The relative lack of variability of the Ear9 pressures for this ensemble is because all 10 of those measurements were taken *after* the ear tip was reinserted. The corresponding V_PRM_|_SPIKE_ summaries from the same ensemble were 0.0143±2.1 (Ear7), 0.00946±2.0 (Ear8), 0.0410±0.7 (Ear9), and **0.0177±5.6** (Ear7–9) [mm/s]±[dB].

Finally, the overall P_EC_|_NF_ summaries from the (P_EC_, V_PRM_)|_NF_ ensemble of points at the operating frequency (Fig. 11B) were 39.9±1.7 (Ear7), 40.1±1.6 (Ear8), 44.8±0.9 (Ear9), and **41.6±2.7** (Ear7–9) [dB SPL]±[dB SPL]. The corresponding V_PRM_|_NF_ summaries from the same ensemble were 0.00457±2.2 (Ear7), 0.00233±3.4 (Ear8), 0.00427±2.5 (Ear9), and **0.00357±3.8** (Ear7–9) [mm/s]±[dB].

The P_EC_ SNRs from the (P_EC_, V_ST_) ensemble at the operating frequency (Fig. 11A), determined between the spike and noise means, were 40.6 (Ear7), 40.3 (Ear8), 40.5 (Ear9), and **40.5** (Ear7– 9) dB SPL. The corresponding V_ST_ SNRs from the same ensemble were 13.8 (Ear7), 14.1 (Ear8), 18.8 (Ear9), and **15.6** (Ear7–9) dB.

The P_EC_ SNRs from the (P_EC_, V_PRM_) ensemble at the operating frequency (Fig. 11B) were 40.1 (Ear7), 41.0 (Ear8), 40.6 (Ear9), and **40.6** (Ear7–9) dB SPL. The corresponding V_PRM_ SNRs from the same ensemble were 9.9 (Ear9), 12.2 (Ear8), 19.6 (Ear9), and **13.9** (Ear7–9) dB.

The V_ST_|_SPIKE_ ensembles at the operating frequency lie above the respective V_PRM_|_SPIKE_ ensembles by: 0.5 (Ear7), 5.1 (Ear8), –0.2 (Ear9), and **1.8** (Ear7–9) dB. Both Ear7 and Ear9 exhibited remarkably similar average V_ST_|_SPIKE_ and V_PRM_|_SPIKE_ responses, but for Ear8 the V_ST_|_SPIKE_ responses were significantly higher on average.

## Discussion

In this study, we characterized the performance of a novel type of MEMS-based audio transducer that operates at ultrasonic frequencies. The Transducer 1b design generated AM-modulated air-pressure pulses at an ultrasonic carrier frequency of around 200 kHz through rapid-fire mechanical motions. Additionally, it produced AM-modulated ultrasonic motions at an operating frequency of about 100 kHz, effectively demodulating the audio signal into the baseband audio pressure while also emitting residual ultrasonic tones at the carrier and operating frequencies. At both the carrier and operating frequencies, we anticipated high sound pressure levels. Our objective was to understand how those ultrasonic pressure waves are transmitted in humans through the middle ear, or mechanically vibrated through BC pathway to the inner ear. To achieve this, we used human cadaveric temporal bones and measured pressure in the ear canal (P_EC_) and stapes velocity (V_ST_) as a pair, or P_EC_ and promontory velocity (V_PRM_) as a pair, for frequencies up to 240 kHz, which encompassed both the operating and carrier ultrasonic frequencies. These measurements at ultrasonic frequencies in human cadaveric temporal bones are novel and as far as we are aware, not previously reported.

Across various stimulus-drive frequencies (approximately 0.5, 1, 3, 10, and 20 kHz) and voltages (0.2 and 0.05 V_PEAK_), and for the combined Ear7–9 ensemble, we observed relatively consistent P_EC_ measurements at the ultrasonic frequencies, with minimal variation across ears. When the input signal contained tones within the audio band, the corresponding P_EC_ output responses were linear, with harmonic distortions usually less than 1%.

At the operating frequency (∼100 kHz), the overall P_EC_ was **82.2±2.3** dB SPL, with an SNR of about 40.6 dB, when V_PRM_ was measured. When V_ST_ was measured, P_EC_ happened to be about 1 dB lower on average, and the SNR was nearly identical. At the carrier frequency (∼200 kHz), the overall P_EC_ increased slightly to **90.6** dB SPL, accompanied by an SNR of **34.8** dB for a 1 kHz input signal. These measurements of pure-tone emissions will be compared to previously reported safety guidelines (based on 1/3-octave bands) in the literature (see below).

At the operating frequency, the overall V_ST_ was **0.0218±5.7** [mm/s]±[dB]. It remained independent of the stimulus-drive frequency and voltage, and did not vary much across specimens. V_ST_ tended to stay above the NF, with an overall SNR of **15.6** dB. Similarly, the overall V_PRM_ at the operating frequency was **0.0177±5.6** [mm/s]±[dB], with an SNR of 13.9 dB. V_ST_ was slightly above V_PRM_, by about 1.8 dB. The fact that the V_PRM_ spikes at the operating frequency were typically not significantly lower than the corresponding V_ST_ spikes suggests that both AC and BC factors may be contributing significantly to cochlear ultrasound exposure at the operating frequency. On the other hand, at the carrier frequency, the corresponding V_ST_ and V_PRM_ values were within the NF range.

### Air-conduction and bone-conduction transmission

These findings indicate that at the audio-band stimulus-drive frequency, sound transmission primarily occurred through the AC pathway (as indicated by higher V_ST_) rather than through the BC pathway (as indicated by reduced V_PRM_). However, at the ∼100 kHz ultrasonic operating frequency, both the AC and BC ME-velocity indicators remained comparable and above the noise level. Interestingly, at the ∼200 kHz carrier frequency, no significant AC or BC vibrations were discernible from the relatively high NF environment near that frequency.

At audio-band frequencies, V_ST_ motion typically occurs because the acoustic output of a transducer passes into the ear canal and is transmitted through the eardrum and middle-ear bones to vibrate the stapes. The V_PRM_ results would also originate from the transducer at audio-band frequencies, but through mechanical coupling to the temporal bone through one or more mechanisms rather than directly through acoustic coupling. The transducer body did vibrate significantly at audio-band frequencies, but those vibrations were significantly attenuated by the foam ear tips of the transducer and coupler, and the corresponding V_PRM_ measurements at the stimulus-drive frequency were significantly lower than the V_ST_ measurements.

However, the fact that the V_ST_ response was comparable to the V_PRM_ response at the ∼100 kHz operating frequency suggests that mechanical coupling to the temporal bone of some kind was occurring at that frequency, and this needs further consideration. Because V_ST_ is primarily acoustically driven through the middle ear at audio-band frequencies, one might be tempted to think that the high output of V_ST_ at the ultrasonic operating frequency was also due to the AC pathway. However, in this case the MEMS transducer generated both acoustic and vibrational outputs simultaneously at ultrasonic frequencies, so it is possible that some or most of the V_ST_ response at ∼100 kHz was not due to the AC pathway but rather due to one or more BC pathways.

Among the five different mechanisms identified for BC transmission to the cochlea [13], cochlear fluid inertia and cochlear fluid compression are two of the mechanisms that could elicit stapes motion at ultrasonic frequencies. This would be due to vibration of the transducer coupled through the coupler ear tip in contact with the ear canal and propagated through the temporal bone. These BC mechanisms would originate from the cochlear fluid and vibrate the stapes from the inside. We don’t know at this time which of these mechanisms might be involved, and further experiments are needed. These experiments might include V_ST_ and V_PRM_ measurements before and after cutting the ossicles, or before and after draining the cochlea. It is also possible that some amount of the airborne ultrasound from the ear canal may have passed through the eardrum, through the MEC airspace, and through the round-window membrane to stimulate the cochlear fluid [14].

### Previously reported safety guidelines on ultrasound in humans

A primary objective of this study was to assess the safety risk of human exposure to the ultrasonic emissions of a new type of sound-from-ultrasound MEMS-based transducer. While we conducted measurements of the output pressure generated by the transducer and the vibrations it induced in the middle ear and inner ear, we cannot directly address the safety question both due to our inability to measure these levels on living subjects, and due to the limited available information about the physiological consequences of long-term exposure to these ultrasonic levels and frequencies. Nevertheless, we can compare our measurements to the available guidelines from published safety standards to begin to address the safety concerns. These guidelines are summarized below.

#### Environmental health criteria for airborne ultrasound exposure

In 1984, the International Radiation Protection Association (IRPA) published its “interim guidelines” for airborne ultrasonic exposure [7] based on its earlier 1982 document titled “Environmental Health Criteria for Ultrasound” [24]. The 1984 guidelines specifically apply to airborne ultrasound and do not apply when the transducer comes into direct contact with the body. Additionally, IRPA distinguished between workplace exposure, assuming an 8-hour workday, and continuous exposure for the general population. For 8 hours of occupational exposure, the published limit is 110 dB SPL for the 1/3-octave band surrounding 100 kHz. For continuous public exposure, the published limit is 100 dB SPL for the 1/3-octave band surrounding 100 kHz. No guidance was provided for 1/3-octave bands above the 100 kHz center frequency, and no guidance was provided for pure-tone exposure such as are produced by the device tested in this study.

As reviewed in Leighton [6], the 1984 IRPA guidelines still form the basis for many of the official guidelines for airborne ultrasound throughout the world, in spite of their very limited and questionable scientific foundations. As recently as 2024, the International Commission on Non-Ionizing Radiation Protection (ICNIRP) published a statement acknowledging the need for more careful research in order to review the validity of and potentially improve upon the 1984 guidelines [8].

#### Potential physiological effects of airborne ultrasound exposure

As Leighton (2016) reviewed, experiments with ultrasound in air have resulted in the death of insects and mice within 10 to 180 seconds from a 169 dB SPL exposure at 20 kHz, and humans have experienced “dizziness and mild-to-very-unpleasant heating at crevices (fingers, skin clefts, nasal passages)” for airborne exposures of 140 dB SPL or higher [6].

Duck and Leighton observed that, for frequencies below 500 kHz, the primary concern associated with ultrasound in fluid or tissue is cavitation, whereas heating effects become the primary concern above 500 kHz [25]. Still, for the sorts of airborne ultrasound levels that people are likely to be exposed to, Leighton asserted that cavitation should not be an issue [6]. However, Leighton did propose an untested hypothetical mechanism by which airborne ultrasound could potentially induce unpleasant effects in some individuals, such as “migraine, nausea, tinnitus, headaches, fatigue, dizziness, feelings of ‘pressure’”. Much more research would be needed to determine whether this proposed mechanism holds any validity whatsoever, much less whether any of the mentioned side effects could be induced by the specific pattern of ultrasound emissions produced by the device of the present study. Nonetheless, given the many unknowns, caution is warranted.

#### Ultrasound bone conduction hearing in human subjects

Humans may be able to sense ultrasonic frequencies up to 100 kHz through the BC pathway [10–12, 26]. Békésy was the first to demonstrate that, at least in the audio-band frequency region, that once sound reached the cochlea through either AC or BC pathways, they both created the classic traveling wave [27–29]. At ultrasonic frequencies, the situation is likely similar but with some important differences.

For AC stimulation the ME attenuates sounds at ultrasonic frequencies, but seemingly not as much we might have expected. Our results show that at a pressure of 81 dB SPL, the transmission to the stapes at 100 kHz was approximately 0.022 mm/s (Fig. 11A). This can be extrapolated to about 0.1 (mm/s)/Pa. For comparison, the average response at 1 kHz was approximately 0.12 (mm/s)/Pa and gradually decreases by about 20 dB/decade. Consequently, at 10 kHz, the response was around 0.012 (mm/s)/Pa (Fig. 5A). If we assume this trend persists, V_ST_ at 100 kHz would be expected to be approximately 1.2 (µm/s)/Pa (Fig. 11A: Brown circle). However, our measured V_ST_ of about 0.1 (mm/s)/Pa was significantly higher than what one might expect from extrapolation and in fact not much lower than at mid frequencies of 1 kHz. This states that it may not be the middle ear that causes stapes vibrations at ultrasound hearing with AC stimulation, but rather the cochlea itself through BC. Thus, it is possible that the stapes moves not due to the middle ear AC pathway, but rather through cochlear fluid motion that drives the stapes in reverse. Future analysis of the stapes vibration phase could provide insight into this mechanism [20].

Regardless of the mechanism, what are the mechanisms that allows perception of ultrasonic sounds in humans? One possibility is the “cochlear place theory for ultrasonic frequencies” which was derived from pitch matching [30] and loudness matching experiments [26] conducted by Corso and Levine. In the pitch matching experiments, they observed that pitch matching at ultrasonic frequencies was localized to the pitch of the highest audio frequency. When compared to a 57 kHz reference tone, subjects perceived pitches closer to 16 kHz tones as having higher pitch. Notably, they also judged a 57 kHz tone to be higher in pitch than a 64 kHz tone [30]. In a separate study, Corso and Levine conducted loudness matching experiments for AC and BC tones. The AC loudness matching revealed a significant increase in levels for frequencies above 14 kHz, with the response terminating near 17 kHz, close to the upper limit of hearing. Since our middle-ear transmission measurements do not show such a sharp decrease in response above 10 kHz (Fig. 3), it suggests that the cochlea limits AC at high frequencies rather than the middle ear. However, this is not without controversy, because the opposite was argued based on measurements of high-frequency psychophysical tunning curves [31]. The BC loudness-matching experiments indicated that the level increased from approximately 12 kHz to 80 kHz and slightly decreased at 94 kHz. These results corroborate previous studies that the upper limit of BC extends out to 100 kHz [10–12]. Collectively, these studies suggest that the auditory system can process information about loudness discrimination for BC tones in the ultrasonic range, spanning from 20 kHz to at least 85 kHz, but not pitch-discrimination information.

These studies suggest that it is the basal end of the cochlea that responds to ultrasonic frequencies. However, there is no known tonotopic location for frequencies higher than the auditory range. A tone in the auditory range sets up a traveling wave whose wavelength decreases from the base to the apex. In the more basal, long-wavelength region, the amplitude of the traveling-wave envelope increases in the stiffness-dominated region. It reaches a peak at its characteristic place with a medium wavelength. As the wavelength gets even shorter, the amplitude decreases rapidly due to fluid viscosity. When an ultrasonic tone causes the basal end to respond, it lacks time to develop the classic traveling wave. Instead, it likely responds like a stiffness-mass-damping resonant structure, where the resonant frequency is higher than the audiometric frequency and thus off the basal end of the cochlea. Consequently, the cochlear partition responds to the high side of a resonant tuning, likely the mass-dominated region. In this manner, as the ultrasonic frequency increases, it causes changes in the amplitude of the envelope of the basal end and not the pitch. Changes in the amplitude of a fixed ultrasonic frequency can cause a change in the amplitude of the basal end of the cochlear partition. The observation that, at ultrasonic frequencies, amplitude changes can cause a change in hearing perception by the brainstem indicates that the primary auditory nerve fibers, originating from inner hair cells, are still responding to the envelope of the ultrasonic tone and have not yet saturated. This is likely the reason there is virtually no pitch discrimination with ultrasound stimulation [19].

### Animal measurements with ultrasound at high levels

Animal studies have shown that ultrasound can stimulate neural activity in the ascending auditory pathway. This stimulation is observed in the inferior colliculus (IC) and primary auditory cortex (A1), with activation latencies similar to those triggered by acoustic stimuli. This effect is consistent across a wide range of ultrasound stimulation parameters. The activation is mediated via a cochlear fluid pathway, where vibrations of the cerebrospinal fluid (CSF) cause vibrations within the cochlea through the cochlear aqueduct (this is one of the five BC mechanisms mentioned earlier), ultimately activating the ascending auditory pathway [32].

In a recent guinea pig study, ultrasound was used to evaluate stimulation parameters that safely and effectively engaged the auditory system during continuous ultrasound stimulation through IC recordings. Non-invasive physiological measures of auditory brainstem response (ABR) and distortion-product otoacoustic emissions (DPOAEs) were measured [33]. In chronic ultrasound experiments, animals were subjected to 10 minutes of continuous transcranial ultrasound delivered through a direct contact transducer (carrier frequency: 500 kHz, unmodulated, pulse duration: 5 ms, onset/offset ramps: 5 ms, duty cycle: 50%, and pressure range: 50-100 kPa).

In these chronic preliminary experiments, 10 minutes of transcranial ultrasound exposure at 50 kPa (188 dB SPL re 20 µPa) did not produce any measurable ABR (1–32 kHz) or DPOAE threshold shifts over 14 days. However, a threshold increase was observed in the single animal tested at 100 kPa (194 dB SPL) over the same 14-day period. These preliminary animal experiments suggest that no noise damage was observed in live animals due to 10 minutes of continuous exposure to 500 kHz ultrasound at 50 kPa.

It is unclear how these findings translate to potentially much longer exposure times to 100 kHz ultrasound in humans. The SPLs generated by the device we tested were lower than 91 dB SPL, which is more than five orders of magnitude less than the 194 dB SPL that was shown to damage cochlear structures in the acute animal experiments.

### Summary of key findings

With Transducer 1b, the maximum ear-canal pressure, measured within approximately 20 mm of the eardrum, was usually less than 82 dB SPL at the operating frequency of about 100 kHz, and 91 dB SPL at the 200 kHz carrier frequency. The level at ∼100 kHz is significantly lower than the current exposure guidelines, which set occupational limits of 110 dB SPL and public exposure limits of 100 dB SPL at 100 kHz, assuming an 8-hour workday and 24-hour continuous exposure, respectively [7]. One limitation is that the guidelines assume free-field exposure, and our measurements were taken at the ear canal. Due to standing waves in tubes, levels at the ear canal can vary depending on measurement location in the canal, but near the tympanic membrane they can be expected to even higher by as much as 10 dB. This would raise the effective guidelines to 120 dB SPL for the 8-hour occupational limit and 110 dB SPL for the continuous public-exposure limit for pressures measured in the ear canal. Especially with this greater margin of error, the 82 dB SPL levels at ∼100 kHz from Transducer 1b can be said to fit well within these existing exposure guidelines.

Although the exposure guidelines do not cover the 200 kHz region occupied by the carrier frequency, the fact that the measured vibrations in the middle ear were within the spectral noise at that frequency suggests that the effective vibrational exposure to the inner ear at the carrier frequency may be lower than at the operating frequency in spite of its somewhat higher pressure (91 vs. 82 dB SPL).

While the free-field guidelines are important, the literature has not yet addressed how much ultrasonic vibration is transmitted to the cochlea through the middle-ear or bone-conduction route(s).

Our results show that at a pressure of 81 dB SPL, the transmission to the stapes at ∼100 kHz was approximately 0.022 mm/s (Fig. 11A). This can be represented as about 0.1 (mm/s)/Pa. As we described earlier, for comparison, this was significantly higher than what one would expect from extrapolation based on the middle-ear transfer function alone (Fig. 11A: Brown circle, 1.2 (µm/s)/Pa). This states that it may not be the middle ear that limits ultrasonic hearing, but rather the cochlea itself.

The V_PRM_/P_EC_ ratio at the ∼100 kHz operating frequency was measured to be approximately 0.07 (mm/s)/Pa, which is less than the 0.1 (mm/s)/Pa value measured for V_ST_/P_EC_. These measurements suggest that Transducer 1b is providing significant input to the cochlea at the operating frequency. Due to the tonotopic organization of sound first described by Békésy [34], ultrasonic sounds will not stimulate most of the cochlear locations except possibly in the very basal region of the cochlea closest to the oval and round windows [11]. Above about 10–15 kHz, the threshold of hearing in humans increases significantly [35]. Thus, it follows that the level of ultrasonic sounds would need to be quite high in order to stimulate the basal end of the cochlea. This was borne out in the animal measurements which suggest, for a 500 kHz stimulus, that the input levels would need to be several orders of magnitude higher to cause damage [33]. At the carrier frequency, V_ST_ and V_PRM_ were in the noise and thus immeasurable.

### Conclusions

By conducting human cadaveric temporal-bone experiments (N=3) with a new type of MEMS-based sound-from-ultrasound transducer (Transducer 1b) as the driver coupled to the ear canal, we determined that this particular device’s characteristic airborne ultrasonic emissions at ∼100 kHz produce an average ear-canal pressure of 81 dB SPL, which is well below the 100 dB SPL free-field limit for continuous public exposure to airborne ultrasound as stated in existing guidelines [7].

However, the corresponding measured vibrations of the middle ear in response to this ∼100 kHz emission revealed that a significant amount of vibration was able to consistently reach the cochlea at that frequency, apparently due to both air-conducted and bone-conducted transmission mechanisms, even though the transducer was mechanically isolated from the temporal bone using two sets of foam ear tips.

This finding suggests that design adjustments should be employed to reduce the ultrasonic vibration of the device housing as much as possible in order to prevent potentially much higher vibration levels from reaching the cochlea during practical usage scenarios in which the device housing is placed in direct contact with the outer ear.

For the other major ultrasonic emission of the device, at ∼200 kHz, the average ear-canal pressure was higher, at 91 dB SPL. Although the safety guideline cited above does not specifically cover the 200 kHz region, the corresponding middle-ear vibrations at this frequency were lower than those at ∼100 kHz, and were mostly indistinguishable from the surrounding spectral noise. Therefore, the ∼200 kHz emission appears to pose less of an active concern than the ∼100 kHz emission.

Devices differ in their design specifics. Ultrasonic exposure can vary in frequency and intensity and duration and means of coupling into the ear. Little is known about the actual long-term effects of such exposure, so it is wise to proceed with caution. Existing guidelines have limited scientific basis. However, if they are to be trusted, this device we tested appears to fit within its criteria for safety.

## Acknowledgements

We thank Dr. Anbuselvan Dharmarajan for help with the temporal measurements. This work was supported by a research contract from xMEMS Labs to the Puria Otobiomechanics Lab.

## Appendix: Theory of Operation

### System Architecture and Modulation Strategy

An ultrasonic pulsed-air speaker system akin to the xMEMS Cypress system is illustrated schematically in Appendix-Fig-1. The architecture combines custom high-voltage drive electronics with a capacitive MEMS ultrasonic transducer designed for audio-band sound reproduction. The operational principle involves the production of specially designed carrier and demodulation signals from the input audio waveform and a power source, such that the signals control two pairs of flaps in the MEMS device, labeled SM and SV, in a prescribed way at ultrasonic rates. The sequence of choreographed movements of the flaps, as determined by the carrier and demodulation signals, are such that the device emits a rapid-fire sequence of positive or negative pressure pulses to the external environment that, taken together, form a waveform envelope that effectively reproduces the input audio signal.

The device forces demodulation in a prescribed way, not as an automatic consequence of non-linear signal exposure. The SM and SV flaps shape and control the pressure released in each burst, thereby engineering the demodulation process. Both modulation and demodulation are prescribed by active mechanical components and signals. Neither is a passive harmonic or subharmonic of the other, as they would be actively controlled at specified rates and their 1:2 frequency relationship determined by the operation design. Harmonics could occur due to the mechanical characteristics of the piezo-coated silicon flaps, but they would be expected to occur outside of the band of consideration.

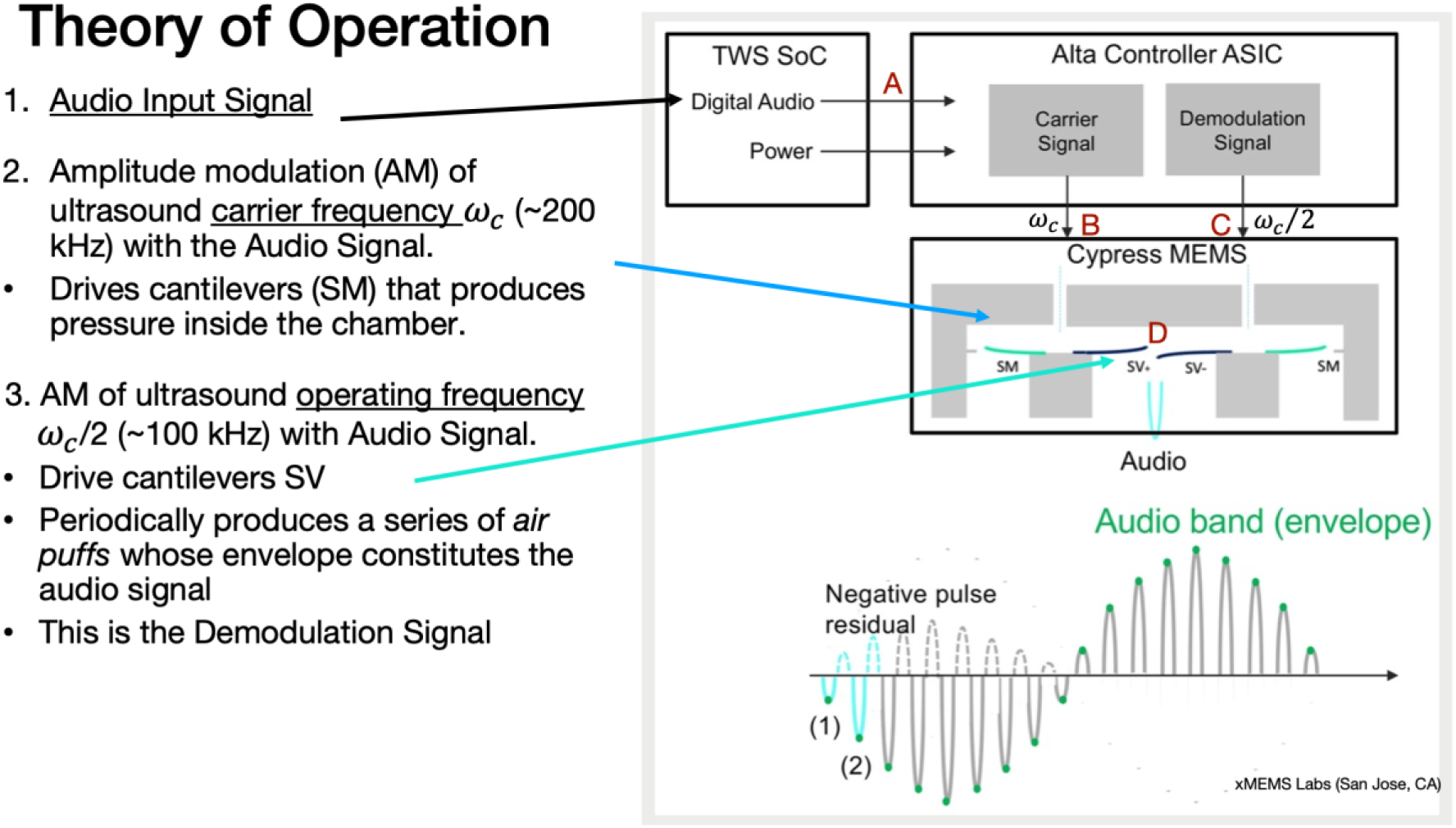
Appendix-Fig-1:

The incoming baseband analog audio signal (Appendix-Fig-1A) is amplitude-modulated by applying it to a switching driver circuit operating at a fixed (switching) carrier frequency of *f*_c_ (∼200 kHz), resulting in signal S_M_ (Appendix-Fig-1B) to control the SM flaps. A demodulation signal S_V_ with twice the period of *f*_c_, or *f*_c_/2 (the ∼100 kHz operating frequency) is also produced, to control the SV flaps (Appendix-Fig-1C). The selection of the carrier and operating frequencies is critical: they must be high enough to minimize audible artifacts, while remaining within the transducer’s mechanical frequency-response bandwidth. This places constraints on the mechanical design of the device. Simultaneously with the modulation signal, the drive circuitry synthesizes a phase-coherent demodulation reference signal (Appendix-Fig-1C) that is applied across the MEMS transducer electrodes. This ensures that the electrostatic forces driving the flaps remain precisely synchronized with the modulation envelope. The resulting time-varying electric field drives rapid, ultrasonic-rate motion of the capacitive flaps (Appendix-Fig-1D).

On a cycle-by-cycle basis, the pressurization cavity within the transducer gets pressurized and depressurized relative to the surrounding air using the peripheral SM flaps during the phase when the central SV flaps are closed. If a positive pressure is produced in the chamber by an inward flexing of both SM flaps while the SV flaps are closed, then briefly opening the SV flaps would release a puff of air with positive pressure. After opening the SV flaps, the pressure would have been temporarily equalized between the chamber and the outside ambient air. If the SM flaps were to retain their flexed position as the SV flaps close, then they could create a negative pressure in the chamber by unflexing to their resting position while the SV flaps were closed.

Then, the next time the SV flaps were opened, a “puff” of negative pressure would be released from the chamber. If the SM flaps were to return to their flexed position while the SV flaps remained open (after the negative puff had been released), then after the SV flaps closed again the SM flaps would be positioned to produce another negative pressure in the chamber by unflexing. Conversely, if the SM flaps were to unflex to their starting position while the SV flaps were open (after the puff had been released), then once the SV flaps were closed the SM flaps would be in a position to either produce a positive pressure in the chamber by flexing inward, or to produce no net pressure by simply not moving by the time the SV flaps opened next (Appendix-Fig-1D).

The requirement that the SM flaps must 1) produce either a compression or decompression in the chamber while the SV flaps are closed, and 2) reposition themselves appropriately while the SV flaps are open so they are ready to compress/decompress again the next time the SV flaps close, explains why the SM flaps at the carrier frequency must operate at twice the frequency of the SV flaps in order to generate the audio-band envelope (bottom of Appendix-Fig-1).

### Advantages Over Conventional Electrodynamic Transducers

The ultrasonic approach described above offers distinct advantages over conventional electrodynamic loudspeakers for compact audio applications. First, because the MEMS flaps oscillate at ultrasonic frequencies rather than at audio frequencies, the displacement requirements are dramatically reduced. From Euler-Bernoulli beam theory, the displacement of a vibrating membrane is approximately inversely proportional to the square of the driving frequency: 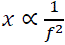 Thus, operating at ∼200 kHz rather than at audio frequencies (1–20 kHz) reduces the required displacement by 3–5 orders of magnitude. This enables compact transducer designs with small diaphragms and reduced moving mass, yielding significant reductions in weight and volume over conventional loudspeakers that favor integration into portable consumer devices such as earbuds, VR headsets, and smart watches. Second, the frequency bandwidth of the MEMS transducer can be engineered to cover a specific ultrasonic range, allowing efficient operation at a fixed operating point rather than requiring multiple resonant frequencies to span the entire audio bandwidth.

## Footnotes

1 The 3D LDV was calibrated to NIST standards by PolyTec in 2017. The PXI system was calibrated by NI in 2021. The optical microphone was calibrated in 2025.

