## Supplemental Information for "Sound from Ultrasound: Characterization of a MEMS-based personal-audio device in human temporal bones"

**Supporting Information**

09-09-2026

Mass General Brigham
Mass Eye and Ear

### Characterization of Extraneous Peaks (Spikes) in Ultrasound Transducer Measurements by Systematically Connecting/Disconnecting and Powering On/Off Each Device in the Signal Chains at the Otobiomech Lab at EPL, Chamber 3

### Table of Contents


Measurement Schematic

Bench-Test Setup

1. Full loopbacks through the USB-6346
2. Loopbacks through the Krohn-Hite 3384 LP filter
3. Everything is connected but powered off
4. Power off everything except the 3384 LP filter
5. Power on JUST the 3384 and xMEMS microphone amp
6. Power on JUST the 3384 and 3D LDV
7. Power on everything EXCEPT the xMEMS speaker driver
8. Power on EVERYTHING
9. Power on everything EXCEPT the 3D carriage controller

Identifying Peaks

Conclusions

### Measurement Schematic

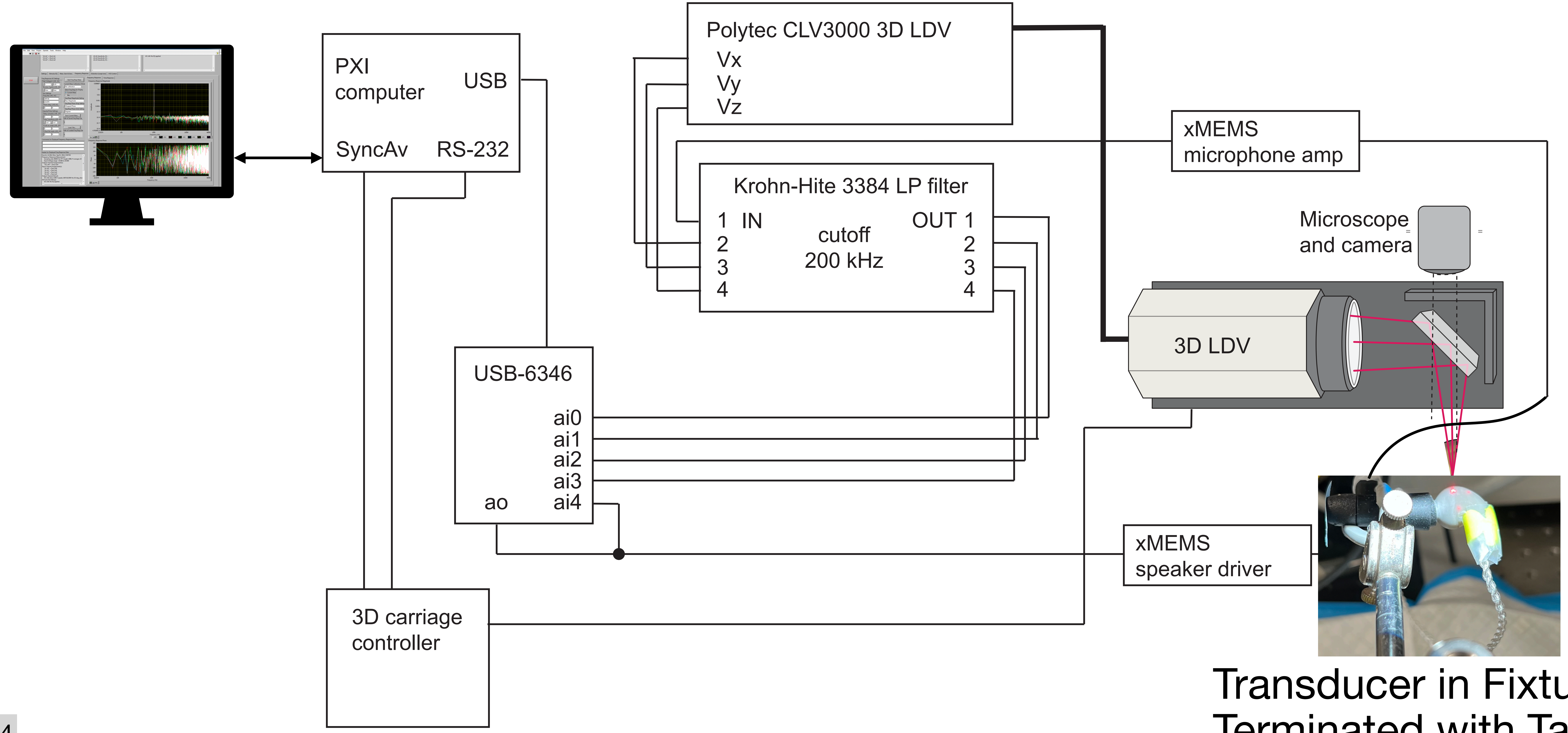


Transducer in Fixture,
Terminated with Tape

### Bench-Test Setup

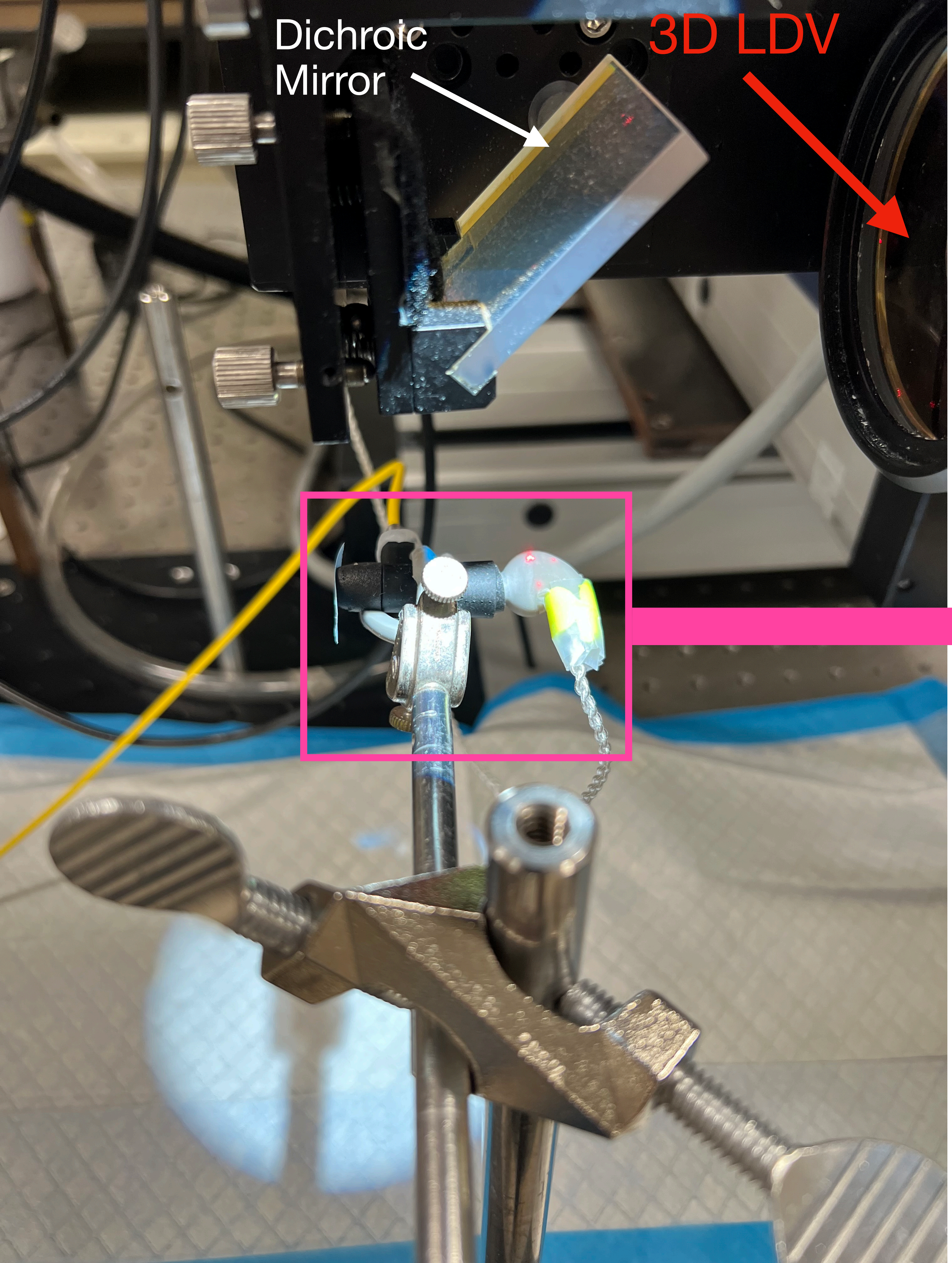


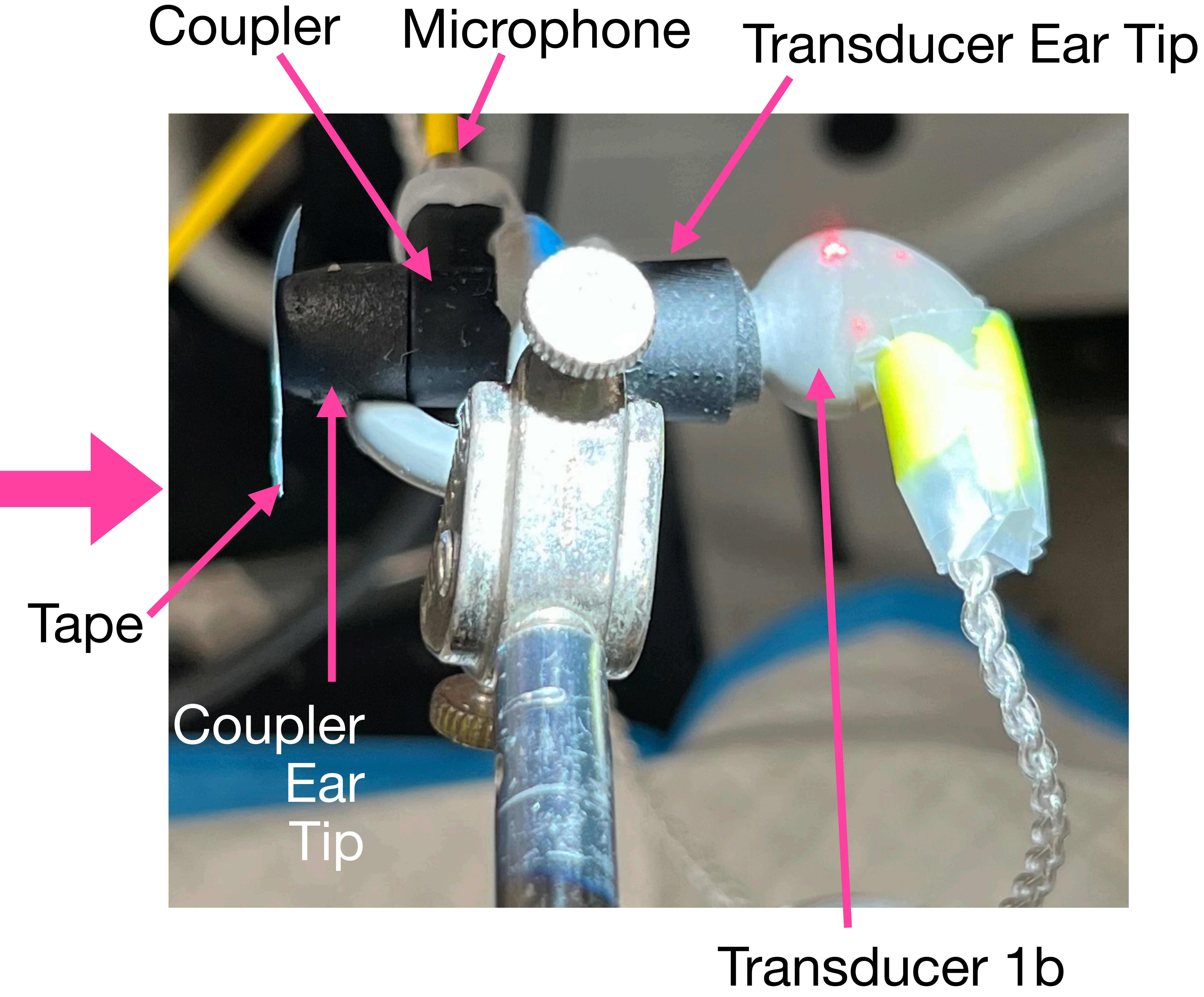

### 1. Full loopbacks through the USB-6346

- **Everything else is disconnected and powered off**
  - ao is connected directly (via BNC t-junctions and short cables) to ai0, ai1, ai2, ai3, and ai4 of the USB-6346
  - The 3384 LP filter is DISCONNECTED and powered OFF
  - The xMEMS microphone amp is DISCONNECTED and powered OFF
  - The 3D LDV is DISCONNECTED and powered OFF (the LDV should have been aimed at the transducer before powering it off)
  - The xMEMS speaker driver is DISCONNECTED and powered OFF
- **Purpose:**
  - This case is to test the consistency of all input channels of the USB acquisition device and to establish a baseline FFT response for a given output signal with no possible contributions from other electronic or measurement components (they are all disconnected and powered off).

### 1. Full loopbacks through the USB-6346

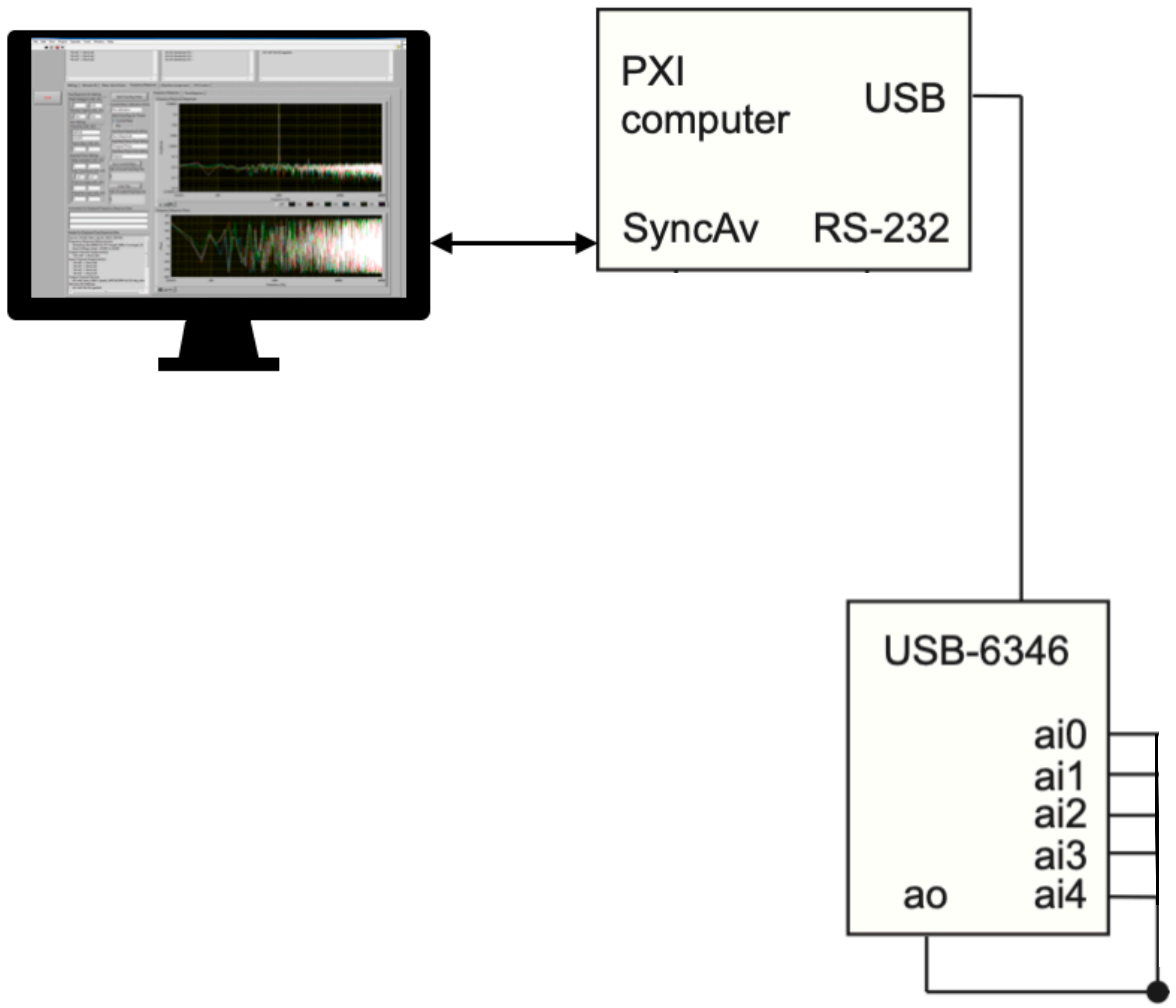

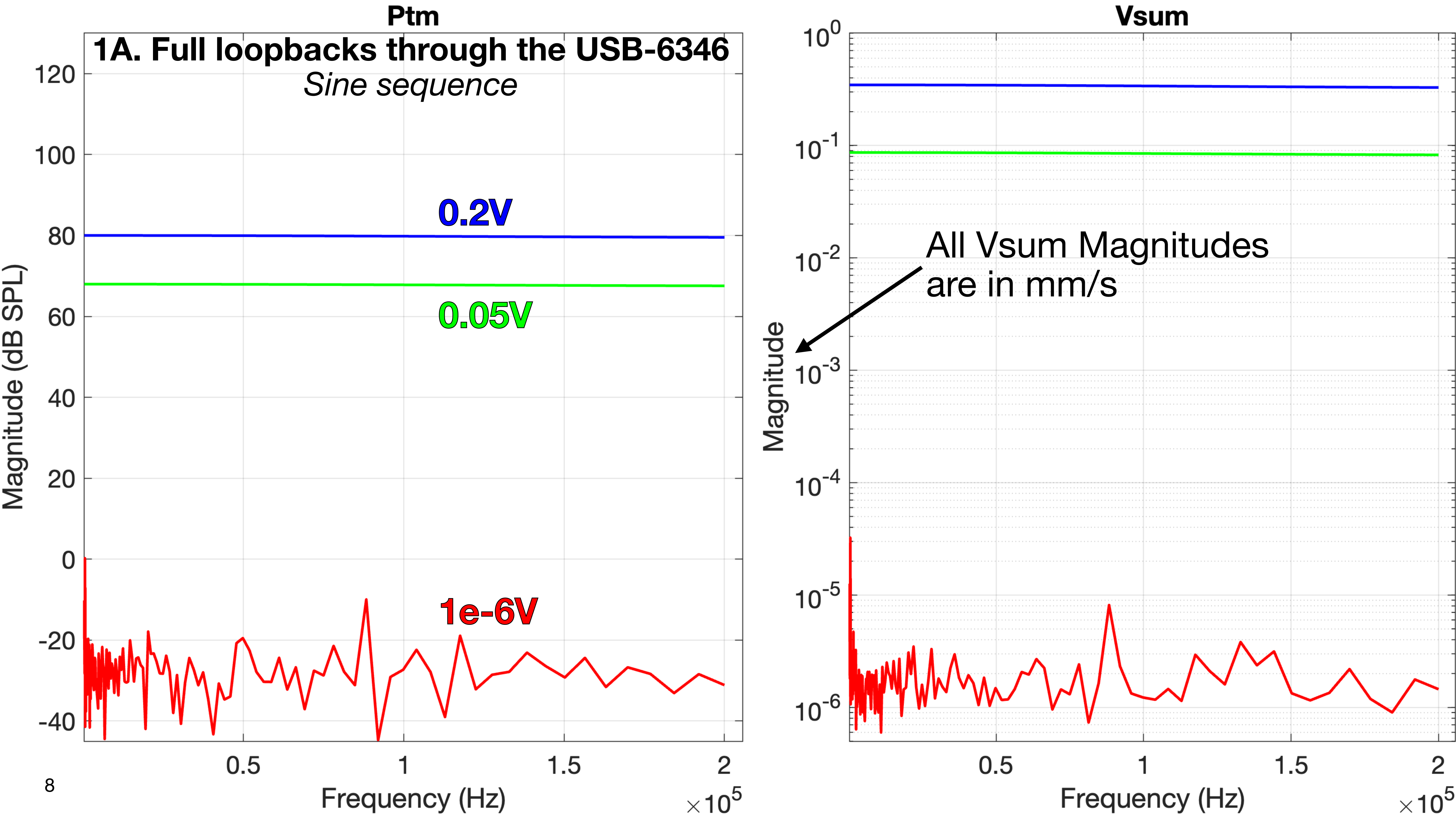

Ptm
1A. Full loopbacks through the USB-6346
Sine sequence
0.2V
0.05V
1e-6V
Magnitude (dB SPL)
Frequency (Hz)
×10^5
Vsum
All Vsum Magnitudes are in mm/s
Magnitude
Frequency (Hz)
×10^5

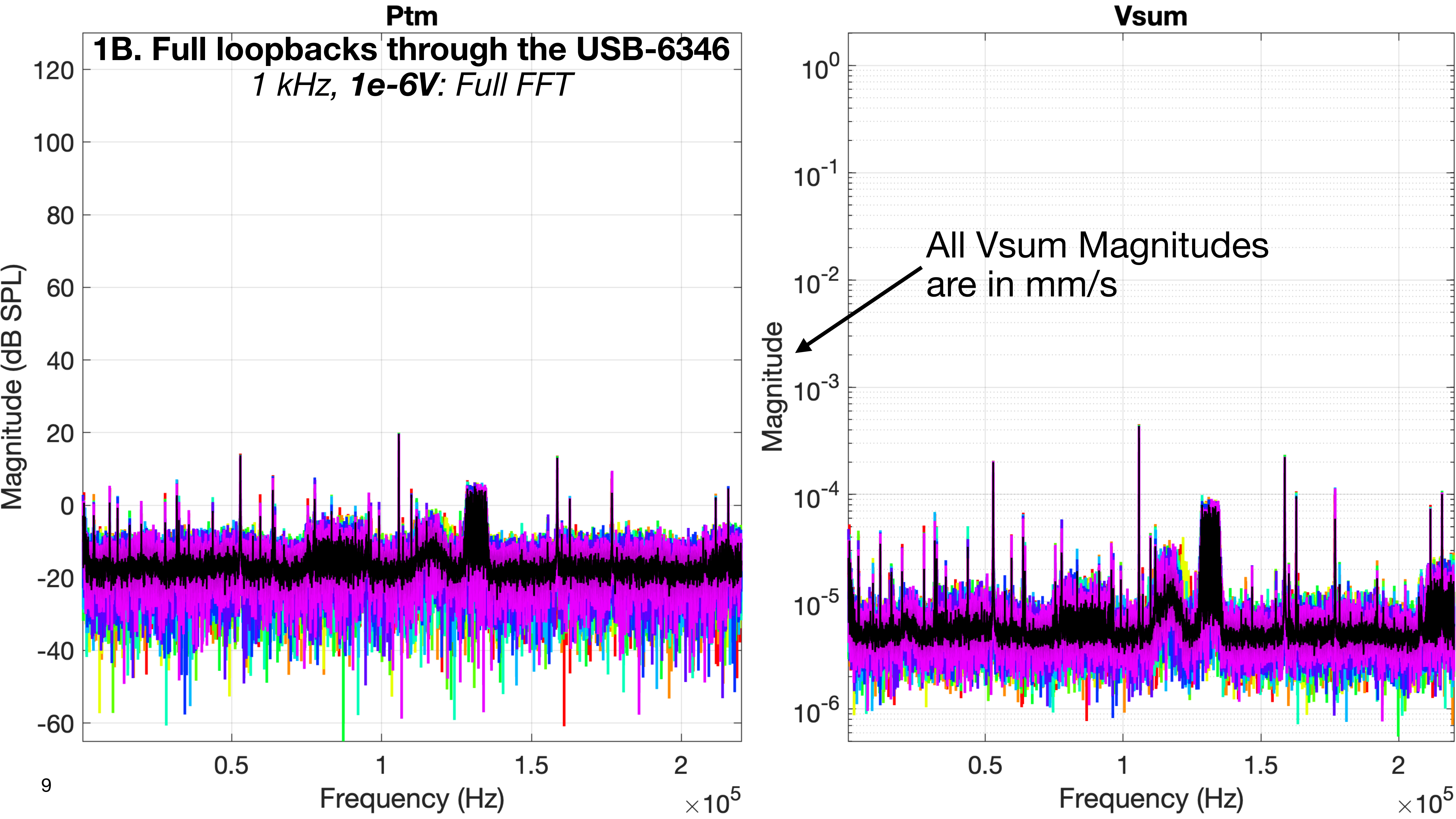
Ptm
1B. Full loopbacks through the USB-6346
1 kHz, 1e-6V: Full FFT
Magnitude (dB SPL)
Frequency (Hz)
×10^5
Vsum
All Vsum Magnitudes
are in mm/s
Magnitude
Frequency (Hz)
×10^5

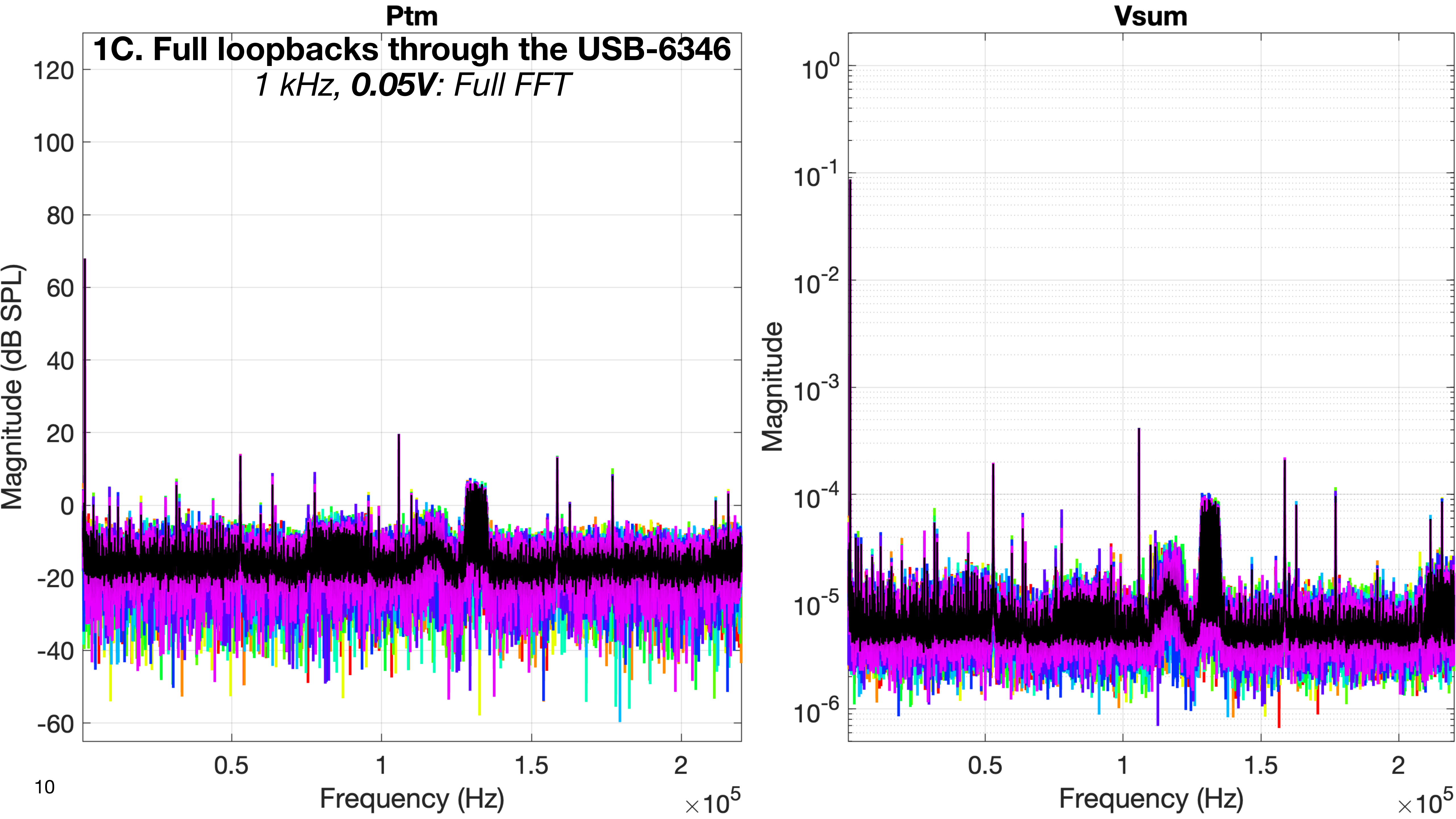
Ptm
1C. Full loopbacks through the USB-6346
1 kHz, 0.05V: Full FFT
Magnitude (dB SPL)
Frequency (Hz)
×10^5
Vsum
Magnitude
Frequency (Hz)
×10^5

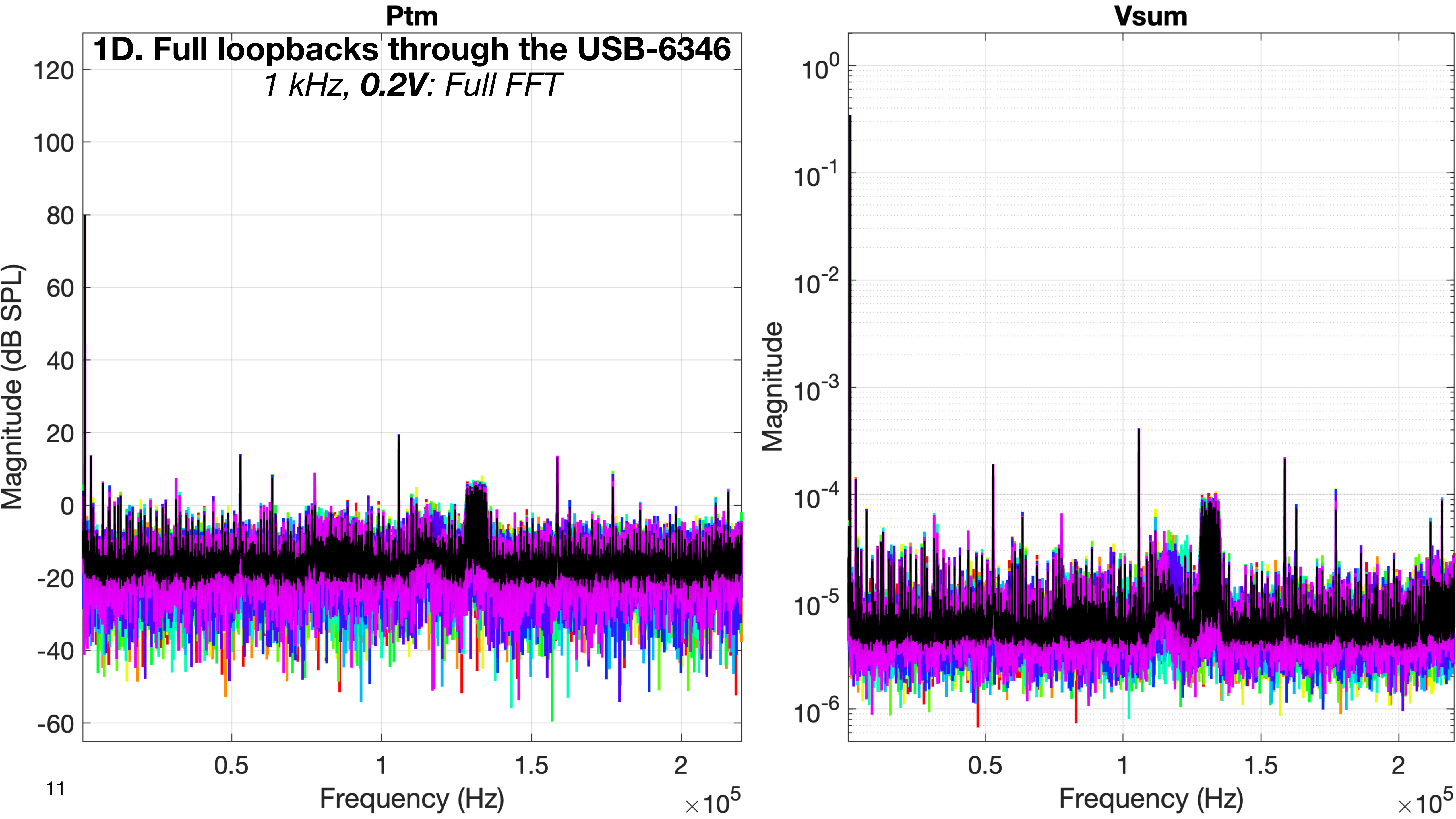
Ptm
1D. Full loopbacks through the USB-6346
1 kHz, 0.2V: Full FFT
Magnitude (dB SPL)
120
100
80
60
40
20
0
-20
-40
-60
0.5
1
1.5
2
Frequency (Hz)
×10^5
Vsum
Magnitude
10^0
10^-1
10^-2
10^-3
10^-4
10^-5
10^-6
0.5
1
1.5
2
Frequency (Hz)
×10^5

### 2. Loopbacks through the Krohn-Hite 3384 LP filter

- **Everything else is disconnected and powered off**
  - ao is looped back to ai4 as usual, and ao is replicated on inputs 1, 2, 3, and 4 of the Krohn-Hite 3384 LP filter.
  - The 4 outputs of the 3384 LP filter are connected respectively to the ai0, ai1, ai2, and ai3 inputs of the USB-6346.
  - The 3384 LP filter is CONNECTED and powered ON, with its normal settings (200 kHz cutoff, etc.)
  - The xMEMS microphone amp is DISCONNECTED and powered OFF (the microphone is in the fixture)
  - The 3D LDV is DISCONNECTED and powered OFF (the LDV should have been aimed at the transducer before powering it off)
  - The xMEMS speaker driver is DISCONNECTED and powered OFF
- **Purpose:**
  - This is to test the behavior of the lowpass filter on its own with no other contributions (they are all out of the signal path and powered off).

### 2. Loopbacks through the Krohn-Hite 3384 LP filter

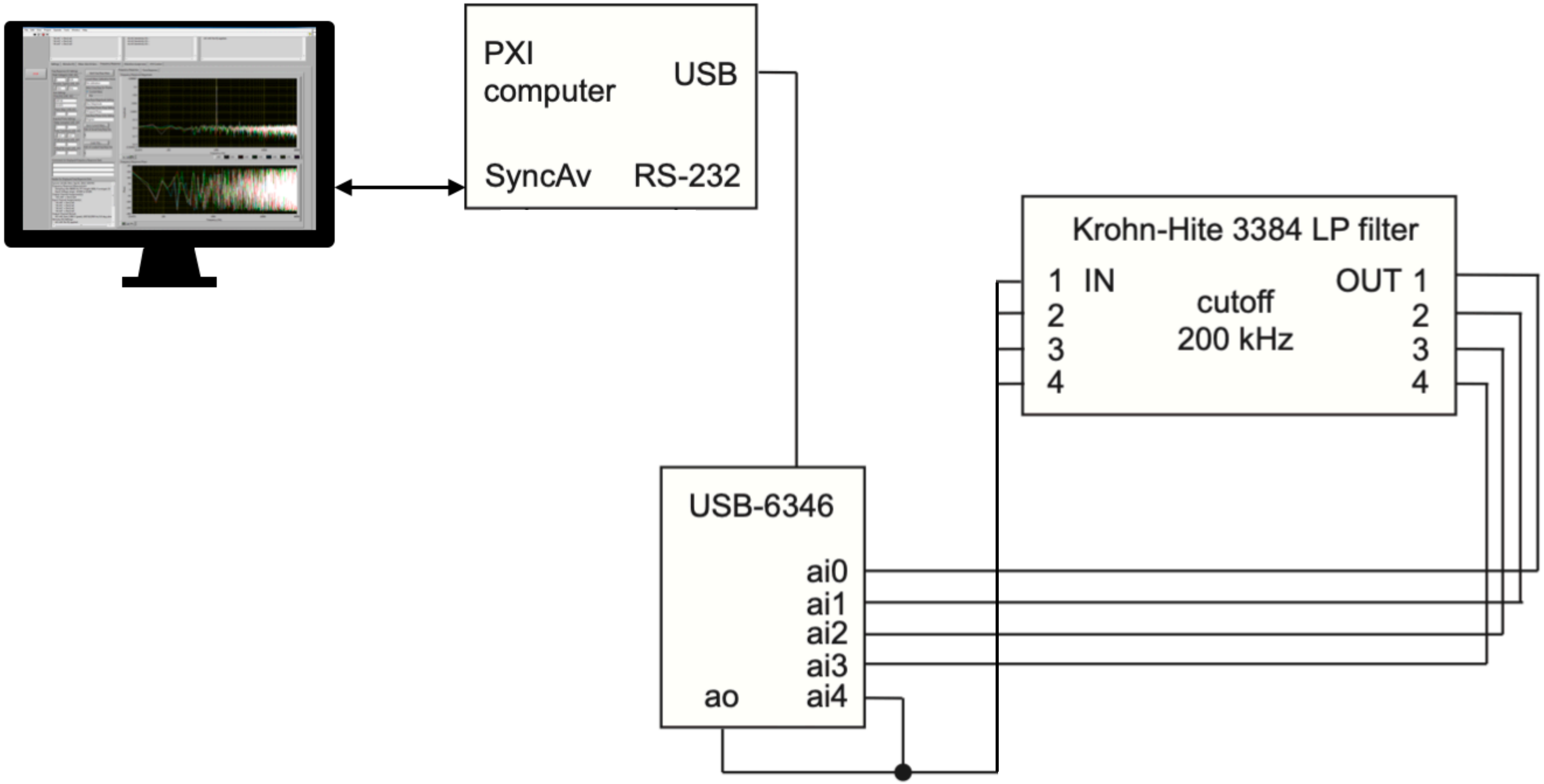

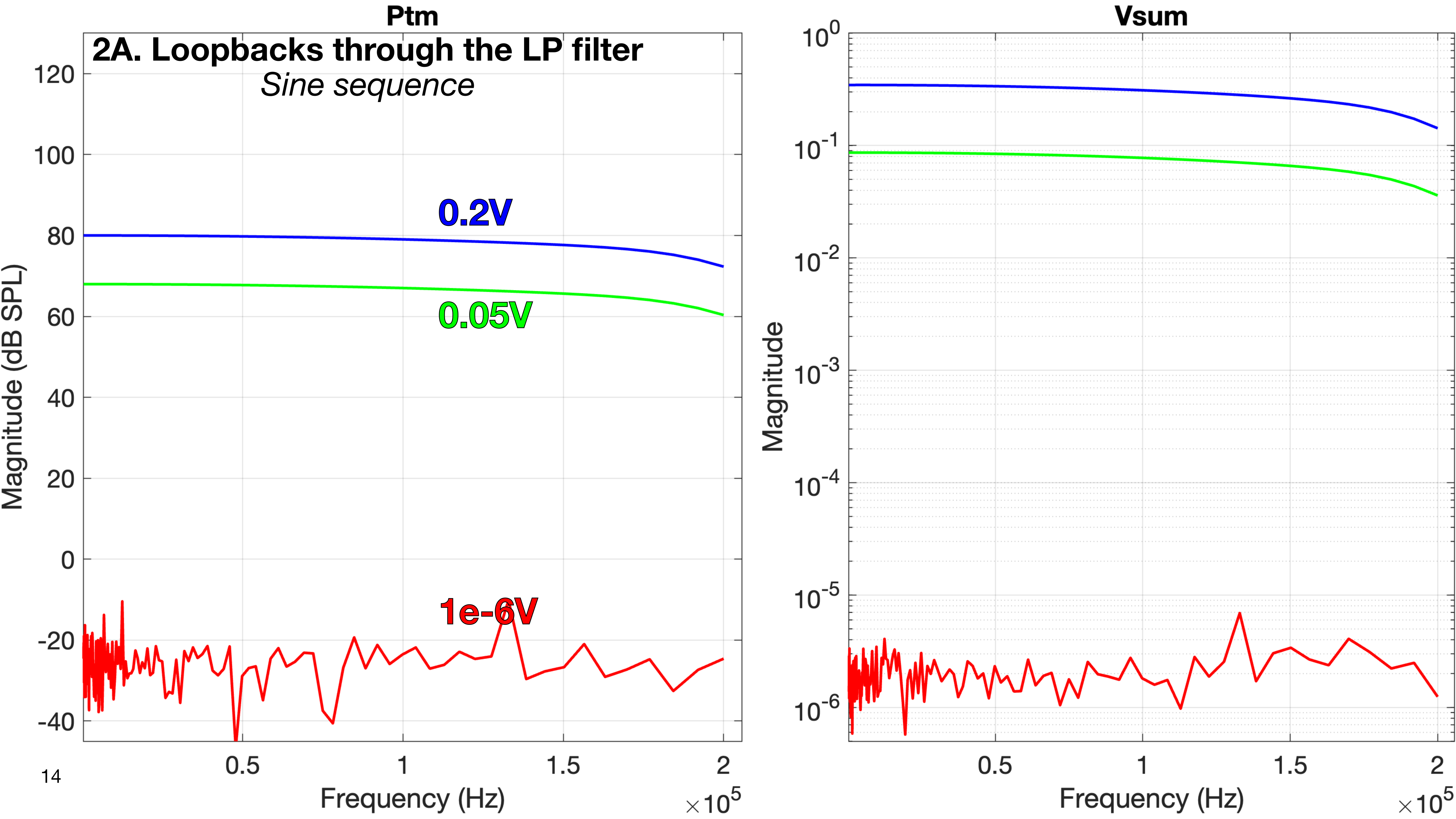

Ptm
2A. Loopbacks through the LP filter
Sine sequence
0.2V
0.05V
1e-6V
Magnitude (dB SPL)
Frequency (Hz)
Vsum
Magnitude
Frequency (Hz)

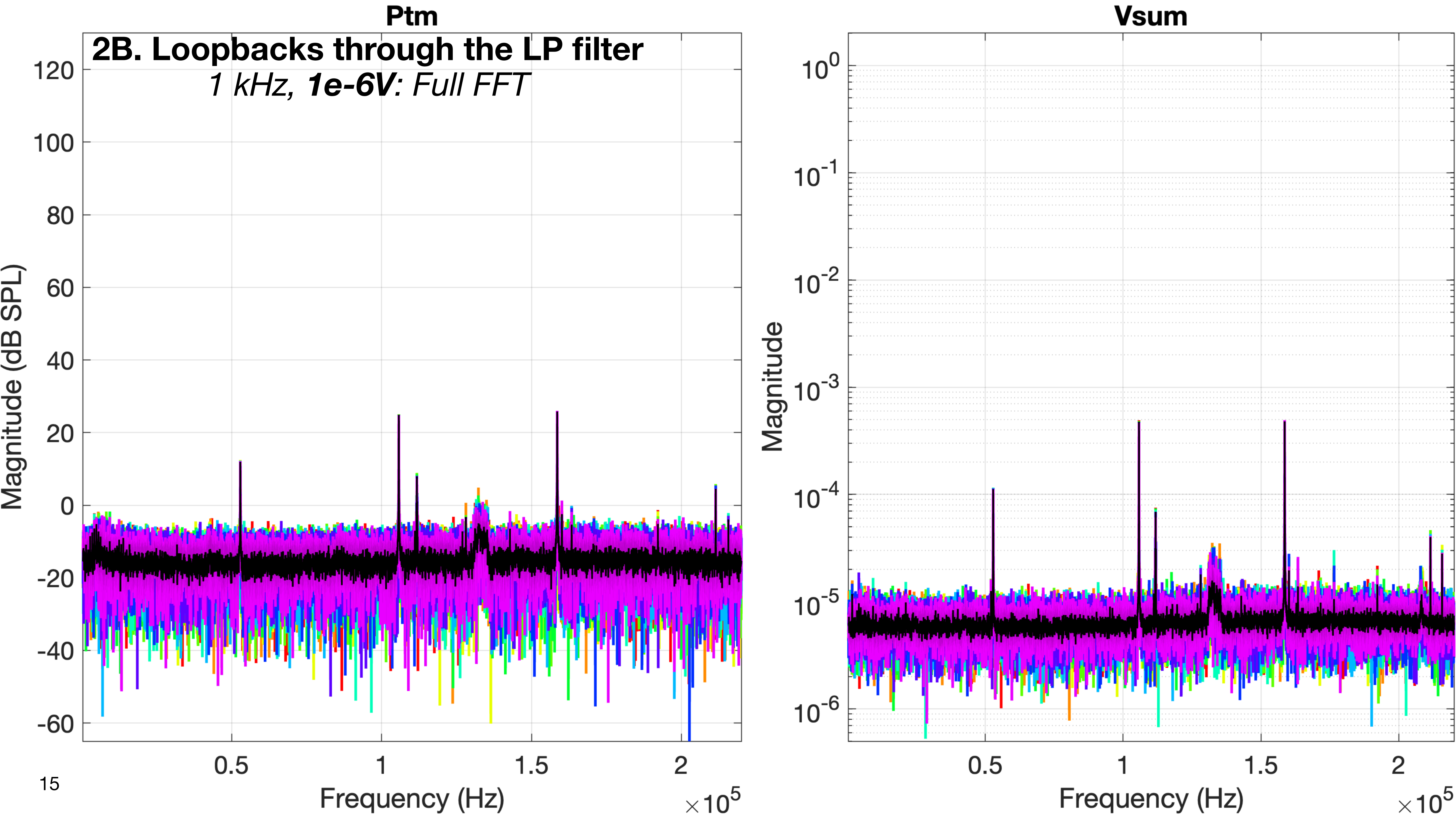
Ptm
2B. Loopbacks through the LP filter
1 kHz, 1e-6V: Full FFT
Magnitude (dB SPL)
120
100
80
60
40
20
0
-20
-40
-60
0.5
1
1.5
2
Frequency (Hz)
×10^5
Vsum
Magnitude
10^0
10^-1
10^-2
10^-3
10^-4
10^-5
10^-6
0.5
1
1.5
2
Frequency (Hz)
×10^5

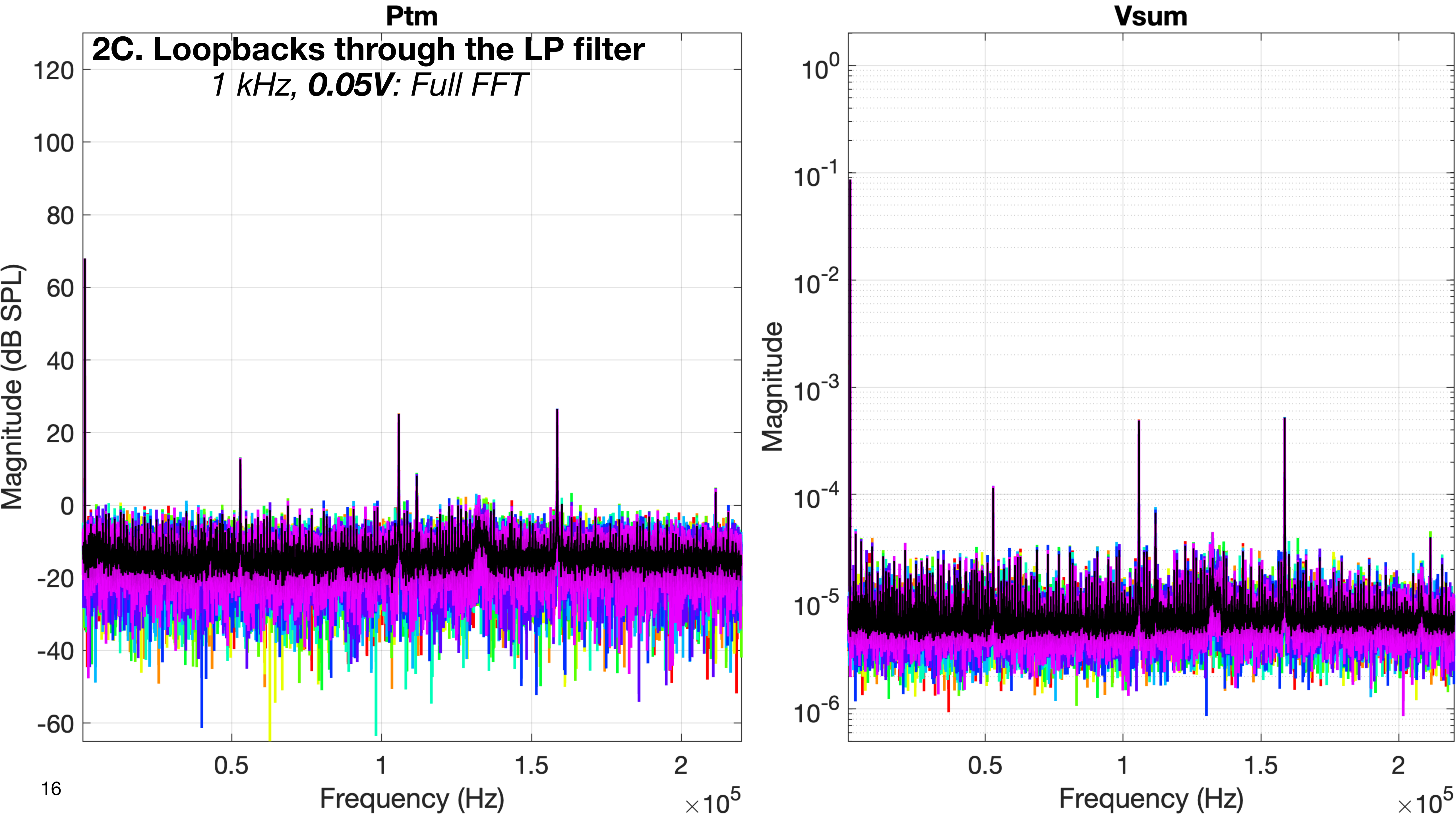

Ptm
2C. Loopbacks through the LP filter
1 kHz, 0.05V: Full FFT
Magnitude (dB SPL)
120
100
80
60
40
20
0
-20
-40
-60
0.5
1
1.5
2
Frequency (Hz)
×10^5
Vsum
Magnitude
10^0
10^-1
10^-2
10^-3
10^-4
10^-5
10^-6
0.5
1
1.5
2
Frequency (Hz)
×10^5

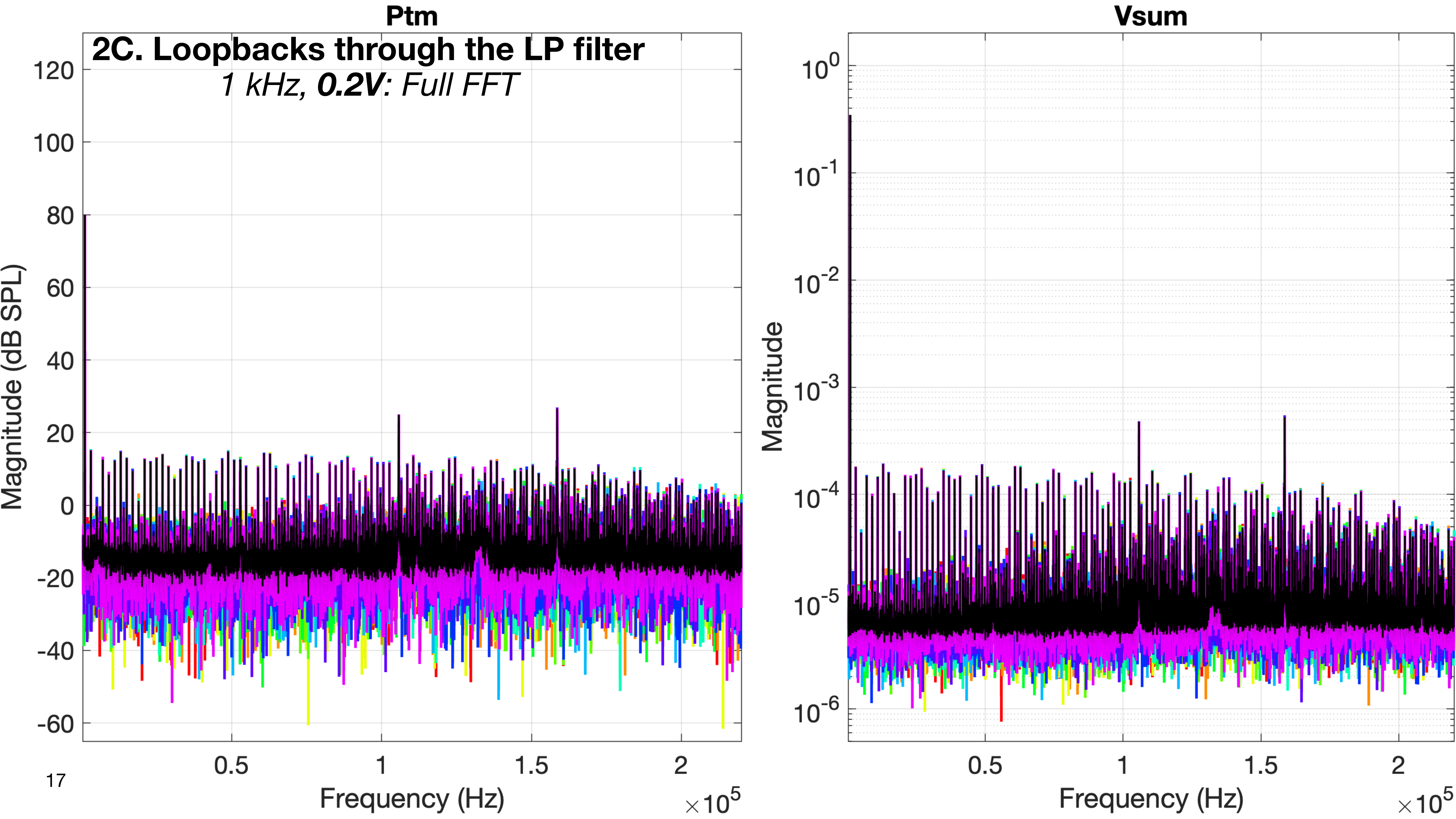
Ptm
2C. Loopbacks through the LP filter
1 kHz, 0.2V: Full FFT
Magnitude (dB SPL)
120
100
80
60
40
20
0
-20
-40
-60
0.5
1
1.5
2
Frequency (Hz)
×10^5
Vsum
Magnitude
10^0
10^-1
10^-2
10^-3
10^-4
10^-5
10^-6
0.5
1
1.5
2
Frequency (Hz)
×10^5

### 3. Everything is connected but powered off

- ao is looped back to ai4 as usual, and additionally connected to the xMEMS speaker driver, as for normal measurements
- input 1 of the 3384 LP filter is connected to the xMEMS microphone amp, as for normal measurements
- inputs 2 through 4 of the 3384 LP filter are connected respectively to Vx, Vy, and Vz of the 3D LDV, as for normal measurements
- The 3384 LP filter is CONNECTED and powered OFF
- The xMEMS microphone amp is CONNECTED and powered OFF (the microphone is in the fixture)
- The 3D LDV is CONNECTED and powered OFF (the LDV should have been aimed at the transducer before powering it off)
- The xMEMS speaker driver is CONNECTED and powered OFF

- **Purpose:**
  - This to test any signal leakage through the normal wiring when all components are connected but powered off.

### 3. Everything is connected but powered off

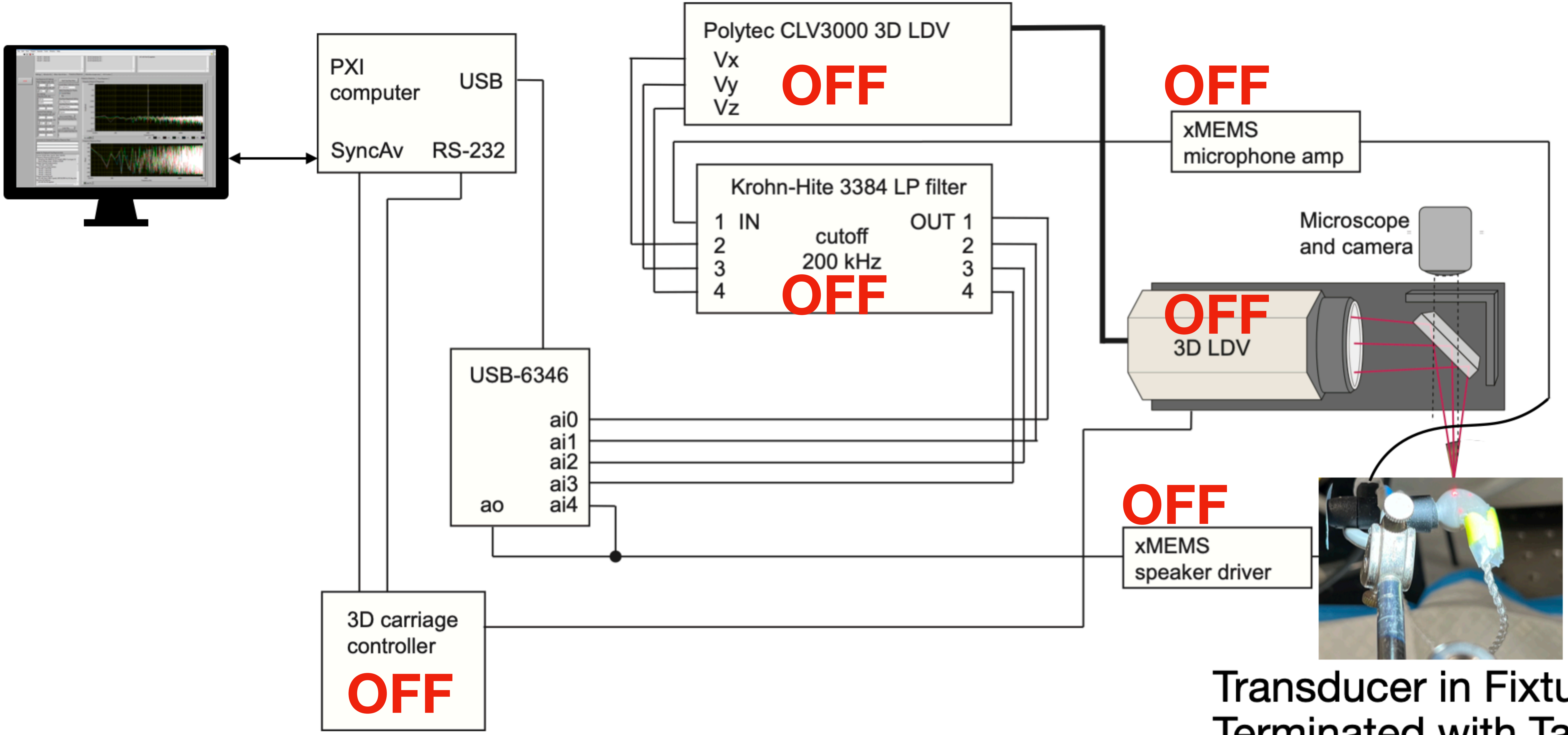


Transducer in Fixture,
Terminated with Tape

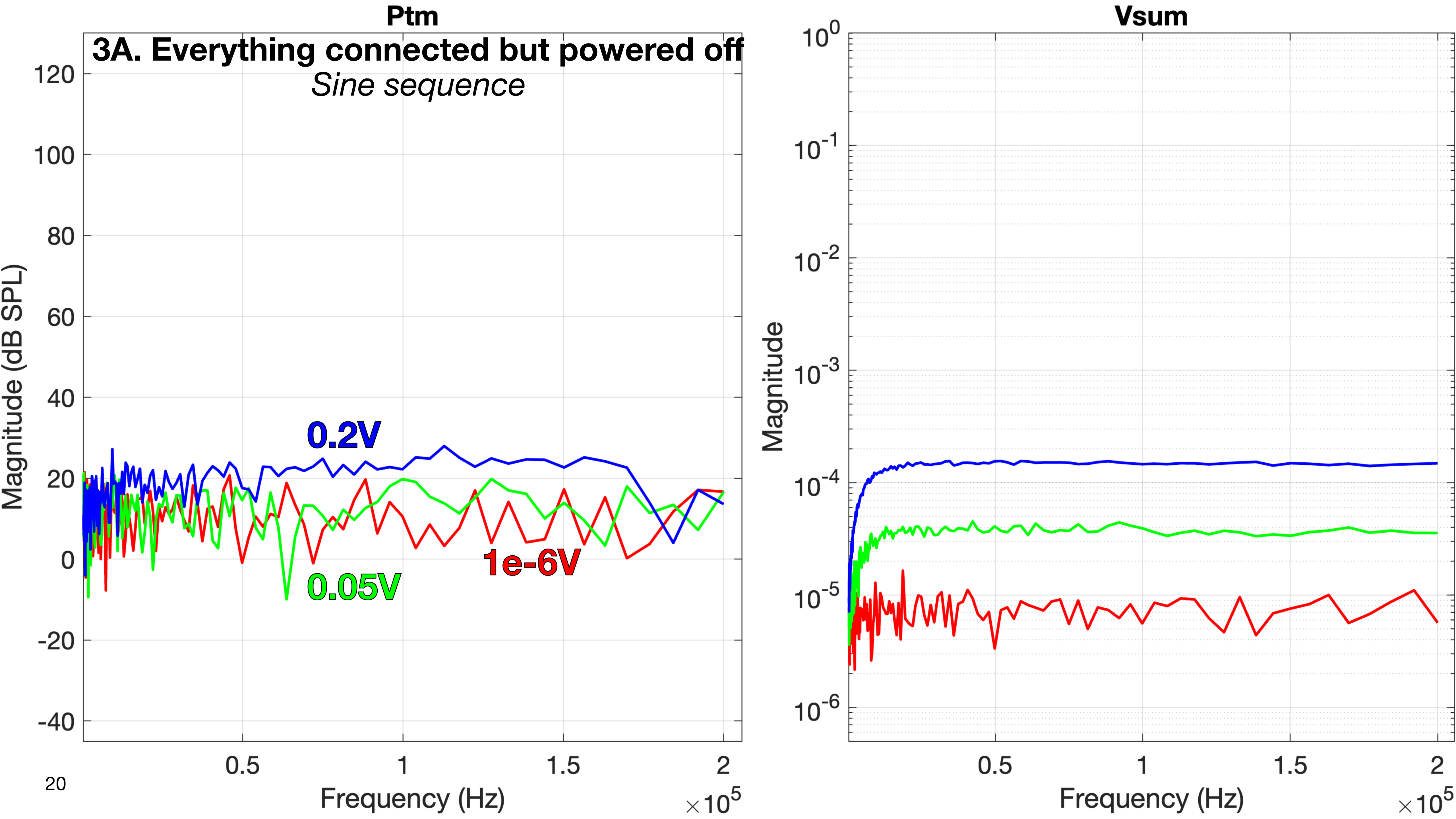

Ptm
3A. Everything connected but powered off
Sine sequence
Magnitude (dB SPL)
120
100
80
60
40
20
0
-20
-40
0.2V
0.05V
1e-6V
0.5
1
1.5
2
Frequency (Hz)
×10^5
Vsum
Magnitude
10^0
10^-1
10^-2
10^-3
10^-4
10^-5
10^-6
0.5
1
1.5
2
Frequency (Hz)
×10^5

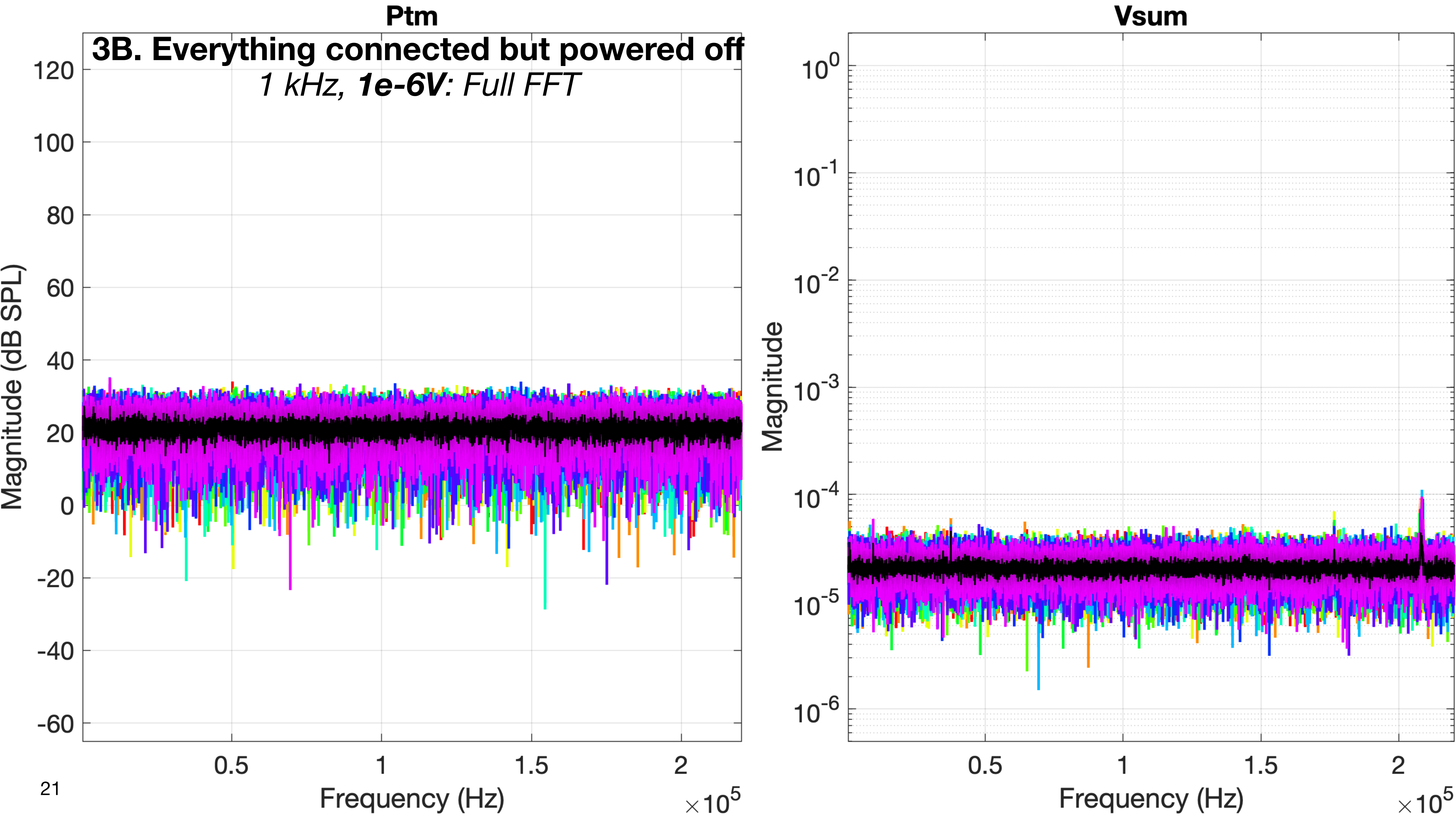
Ptm
3B. Everything connected but powered off
1 kHz, 1e-6V: Full FFT
Magnitude (dB SPL)
120
100
80
60
40
20
0
-20
-40
-60
0.5
1
1.5
2
Frequency (Hz)
×10^5
Vsum
Magnitude
10^0
10^-1
10^-2
10^-3
10^-4
10^-5
10^-6
0.5
1
1.5
2
Frequency (Hz)
×10^5

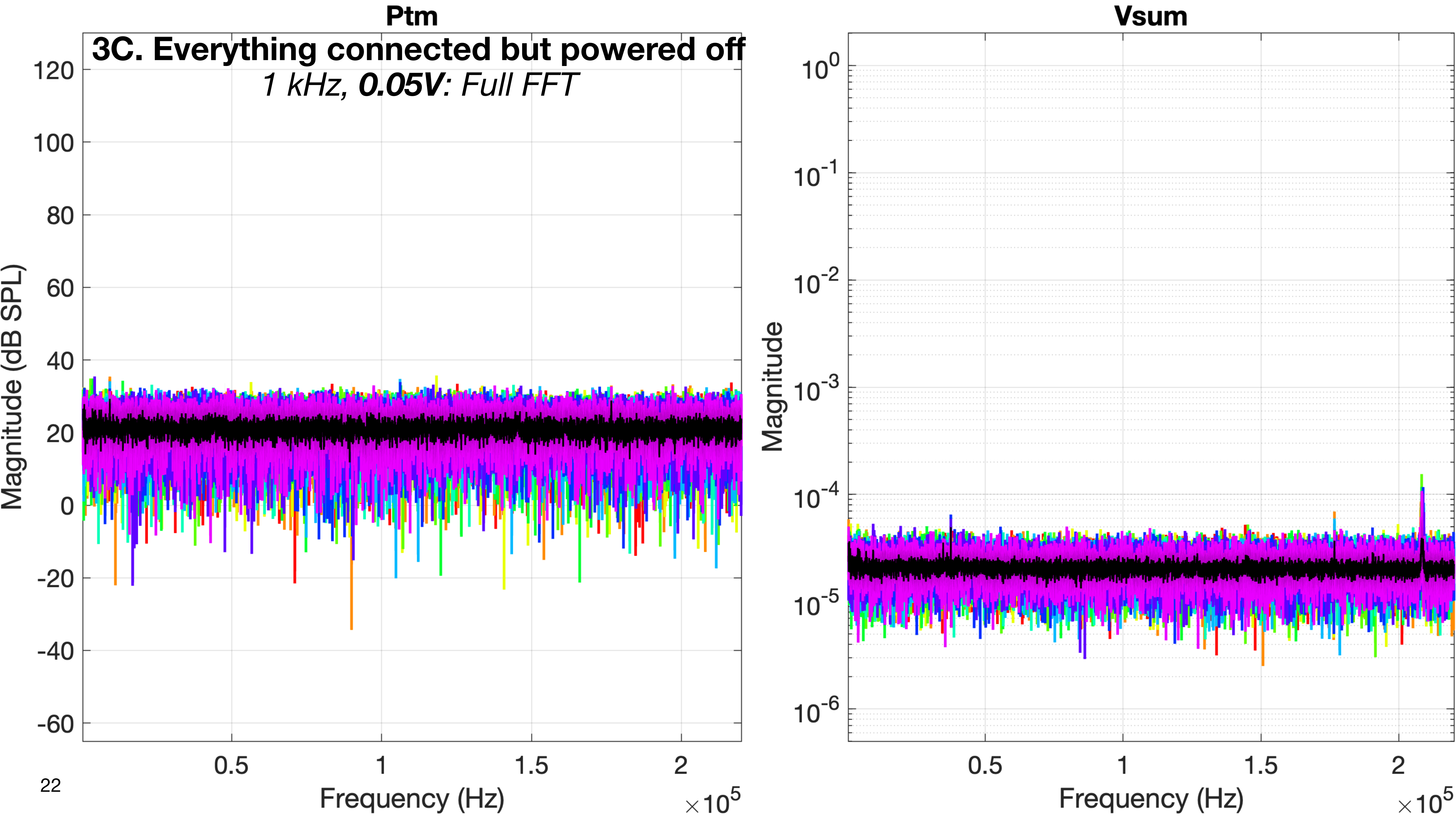
Ptm
3C. Everything connected but powered off
1 kHz, 0.05V: Full FFT
Magnitude (dB SPL)
120
100
80
60
40
20
0
-20
-40
-60
0.5
1
1.5
2
Frequency (Hz)
×10^5
Vsum
Magnitude
10^0
10^-1
10^-2
10^-3
10^-4
10^-5
10^-6
0.5
1
1.5
2
Frequency (Hz)
×10^5

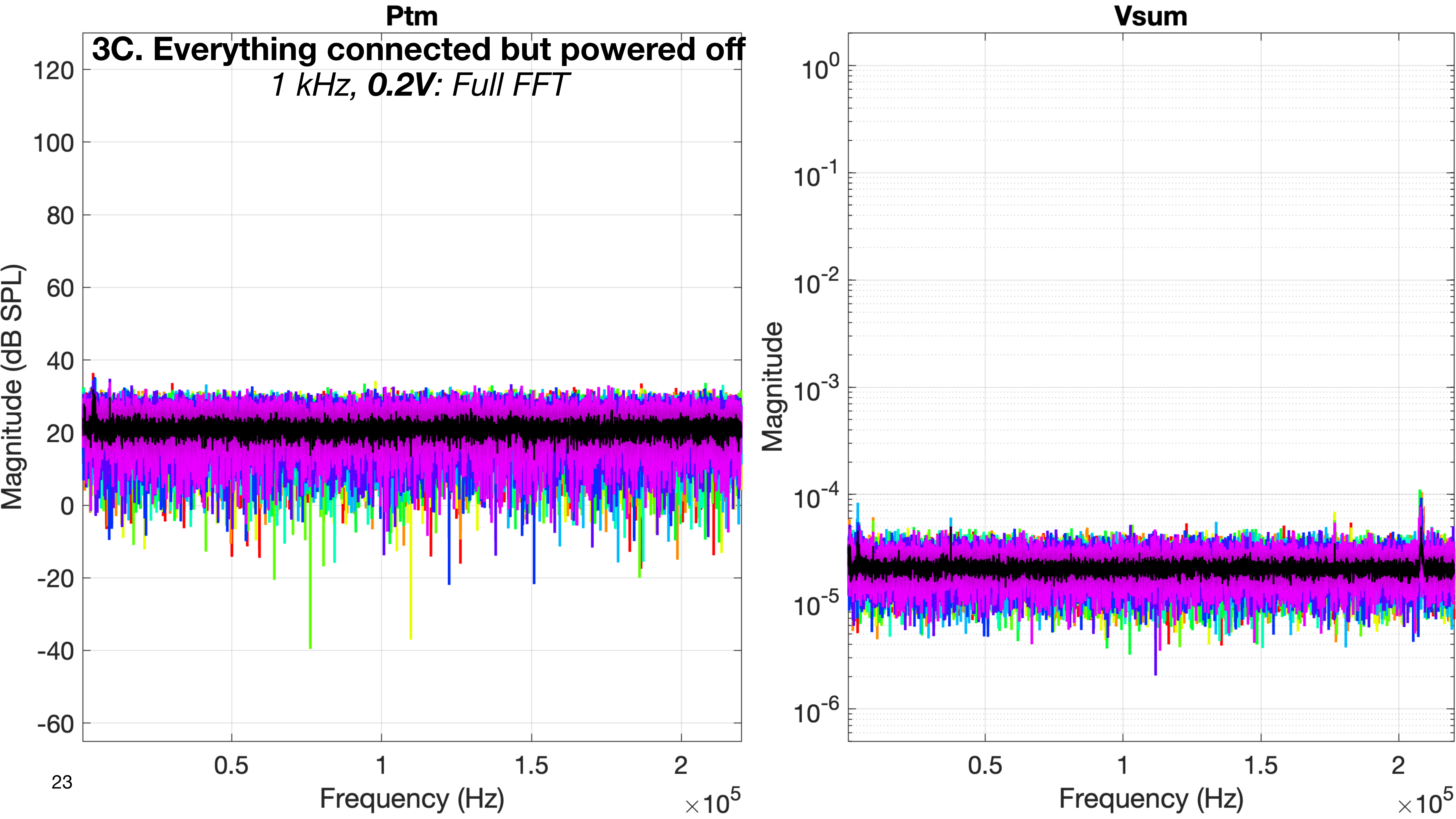
Ptm
3C. Everything connected but powered off
1 kHz, 0.2V: Full FFT
Magnitude (dB SPL)
120
100
80
60
40
20
0
-20
-40
-60
0.5
1
1.5
2
Frequency (Hz)
×10^5
Vsum
Magnitude
10^0
10^-1
10^-2
10^-3
10^-4
10^-5
10^-6
0.5
1
1.5
2
Frequency (Hz)
×10^5

### 4. Power off everything except the 3384 LP filter

- ao is looped back to ai4 as usual, and additionally connected to the xMEMS speaker driver, as for normal measurements
- input 1 of the 3384 LP filter is connected to the xMEMS microphone amp, as for normal measurements
- inputs 2 through 4 of the 3384 LP filter are connected respectively to Vx, Vy, and Vz of the 3D LDV, as for normal measurements
- The 3384 LP filter is CONNECTED and powered ON
- The xMEMS microphone amp is CONNECTED and powered OFF (the microphone is in the fixture)
- The 3D LDV is CONNECTED and powered OFF (the LDV should have been aimed at the transducer before powering it off)
- The xMEMS speaker driver is CONNECTED and powered OFF

- **Purpose:**
  - This is to establish a baseline noise reading of the 3384 LP filter when all other drive and measurement components are connected but not powered on.

### 4. Power off everything except the 3384 LP filter

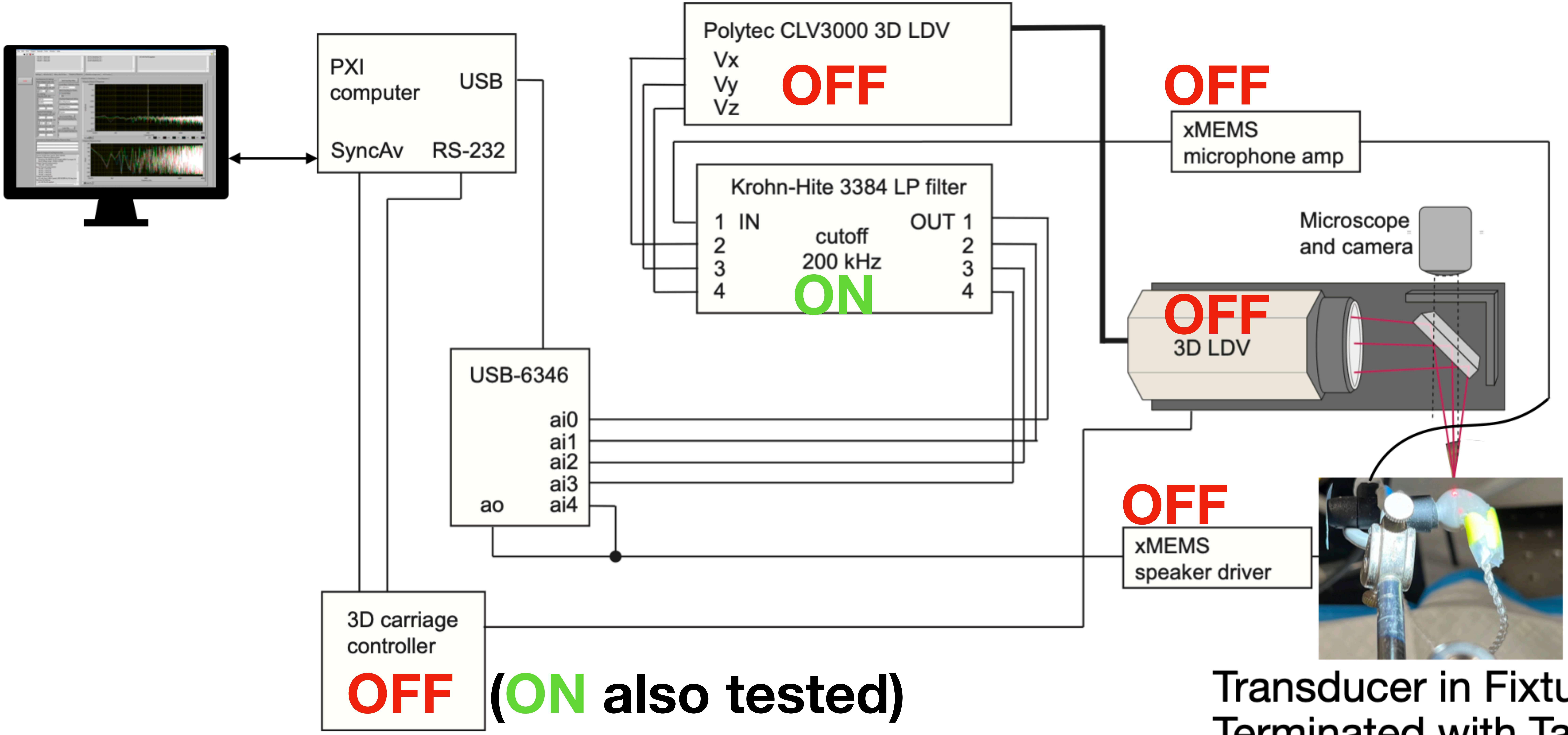


(ON also tested)

Transducer in Fixture, Terminated with Tape

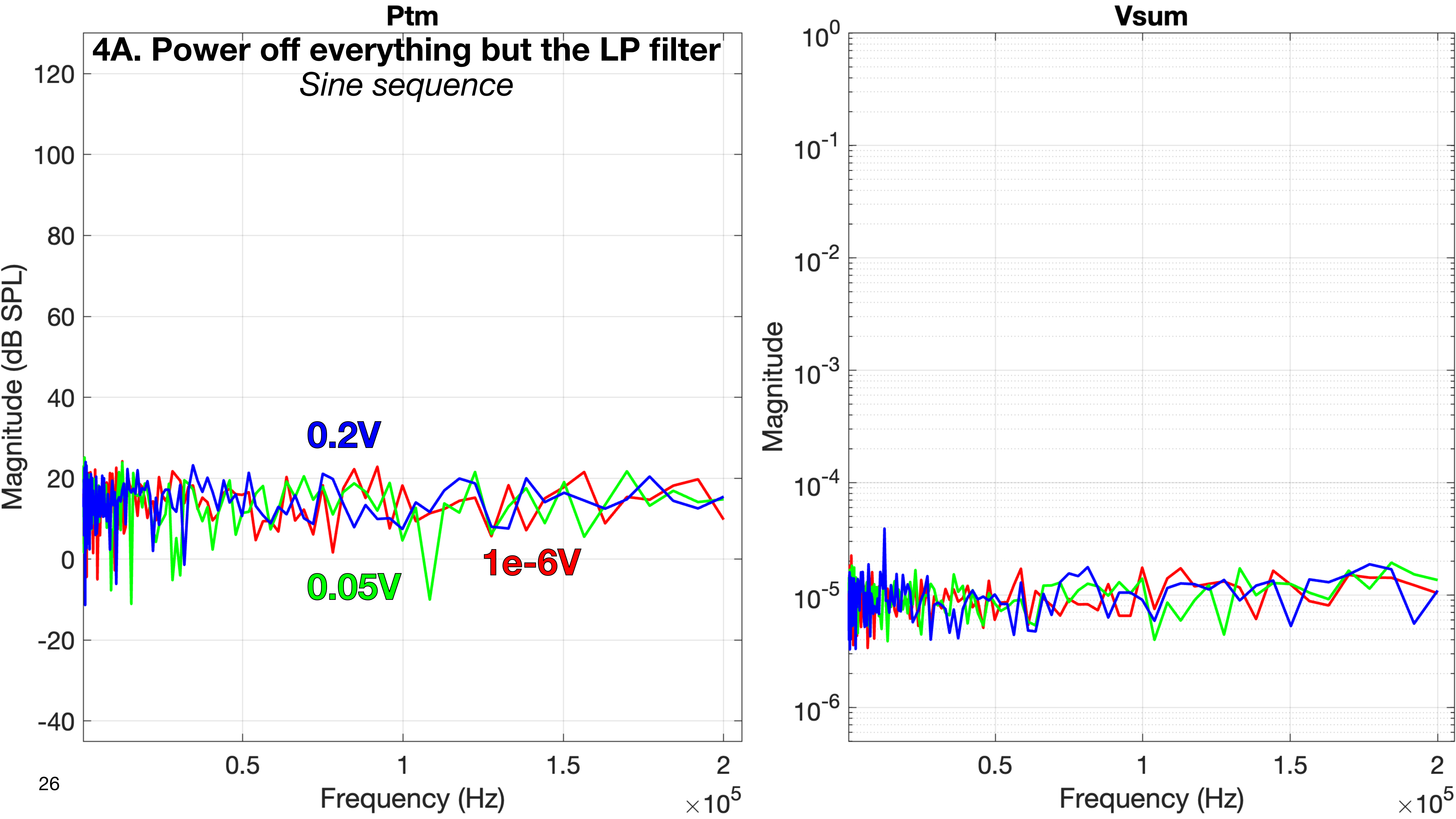

Ptm
4A. Power off everything but the LP filter
Sine sequence
Magnitude (dB SPL)
120
100
80
60
40
20
0
-20
-40
0.2V
0.05V
1e-6V
0.5
1
1.5
2
Frequency (Hz)
×10^5
Vsum
Magnitude
10^0
10^-1
10^-2
10^-3
10^-4
10^-5
10^-6
0.5
1
1.5
2
Frequency (Hz)
×10^5

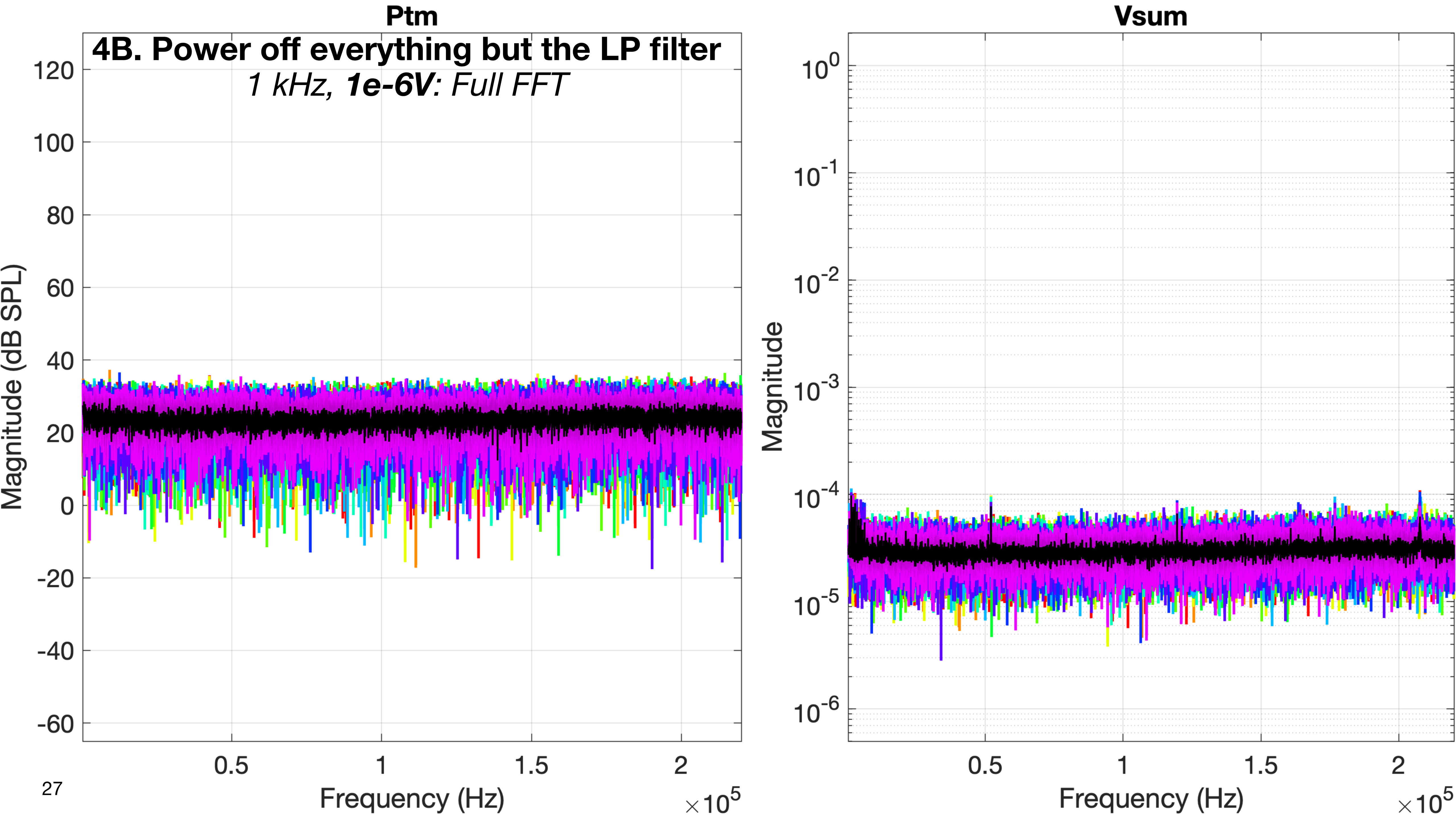
Ptm
4B. Power off everything but the LP filter
1 kHz, 1e-6V: Full FFT
Magnitude (dB SPL)
120
100
80
60
40
20
0
-20
-40
-60
0.5
1
1.5
2
Frequency (Hz)
×10^5
Vsum
Magnitude
10^0
10^-1
10^-2
10^-3
10^-4
10^-5
10^-6
0.5
1
1.5
2
Frequency (Hz)
×10^5

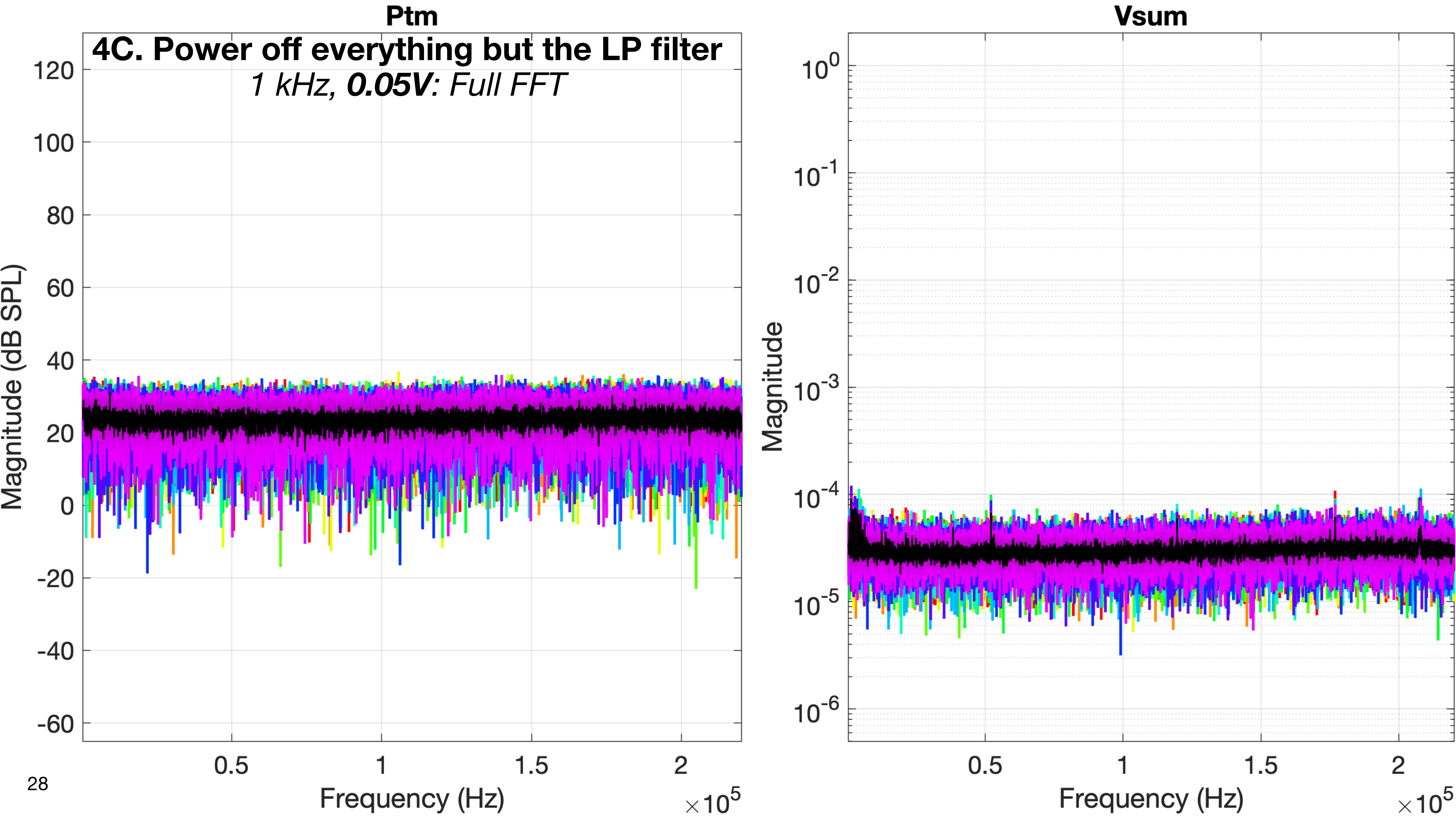

Ptm
4C. Power off everything but the LP filter
1 kHz, 0.05V: Full FFT
Magnitude (dB SPL)
Frequency (Hz)
Vsum
Magnitude
Frequency (Hz)

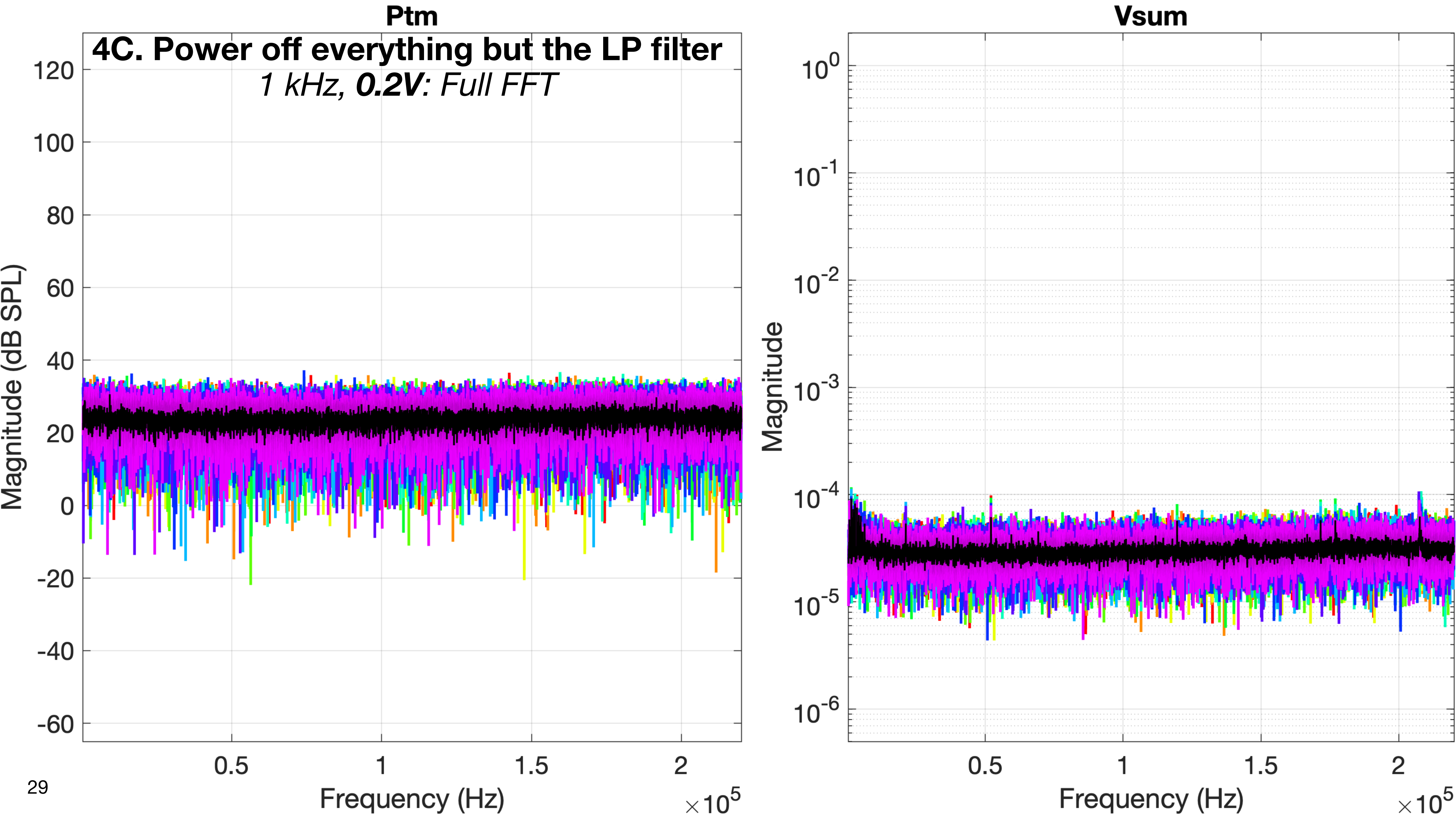
Ptm
4C. Power off everything but the LP filter
1 kHz, 0.2V: Full FFT
Magnitude (dB SPL)
120
100
80
60
40
20
0
-20
-40
-60
0.5
1
1.5
2
Frequency (Hz)
×10^5
Vsum
Magnitude
10^0
10^-1
10^-2
10^-3
10^-4
10^-5
10^-6
0.5
1
1.5
2
Frequency (Hz)
×10^5

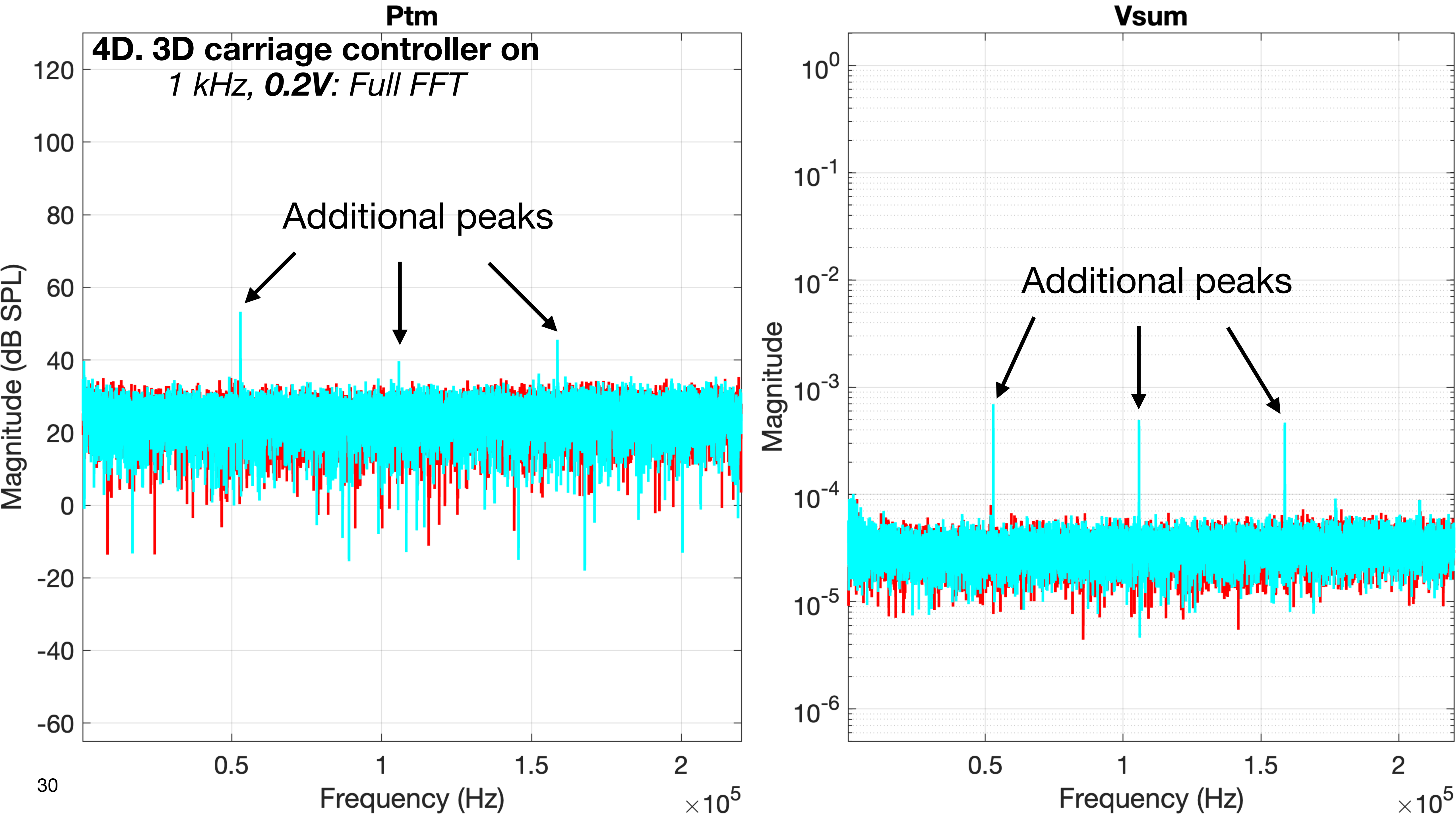
Ptm
4D. 3D carriage controller on
1 kHz, 0.2V: Full FFT
Additional peaks
Magnitude (dB SPL)
Frequency (Hz)
×10^5
Vsum
Additional peaks
Magnitude
Frequency (Hz)
×10^5

### 5. Power on JUST the 3384 and xMEMS microphone amp

- Wiring according to the normal schematic for measurements
- The 3384 LP filter is CONNECTED and powered ON
- The xMEMS microphone amp is CONNECTED and powered ON (the microphone is in the fixture)
- The 3D LDV is CONNECTED and powered OFF (the LDV should have been aimed at the transducer before powering it off)
- The xMEMS speaker driver is CONNECTED and powered OFF

- **Purpose:**
  - This is to establish the contribution of just the xMEMS microphone amp along its normal signal path when measuring a live signal of acoustic noise in the fixture with everything else off.

### 5. Power on JUST the 3384 and xMEMS microphone amp

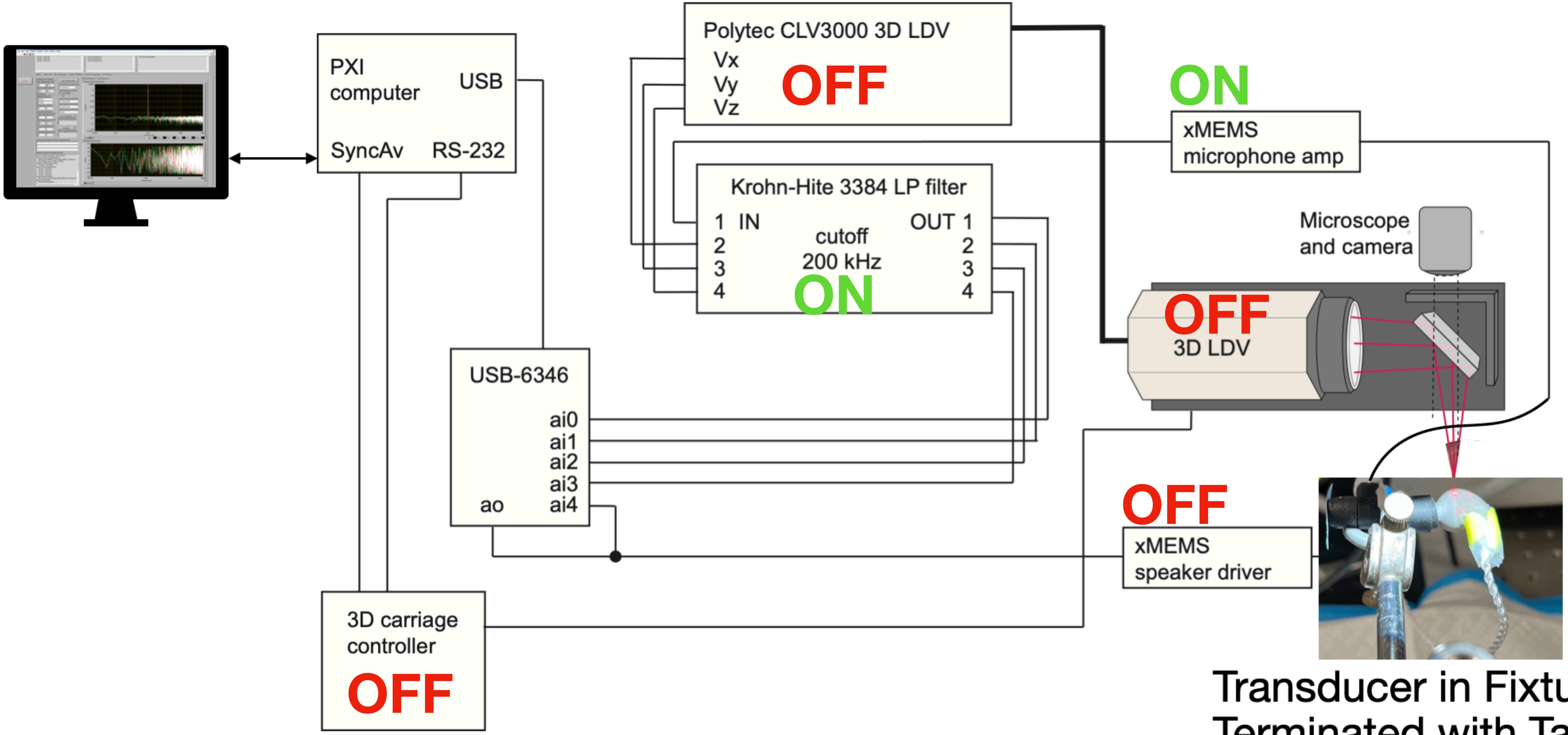


Transducer in Fixture,
Terminated with Tape

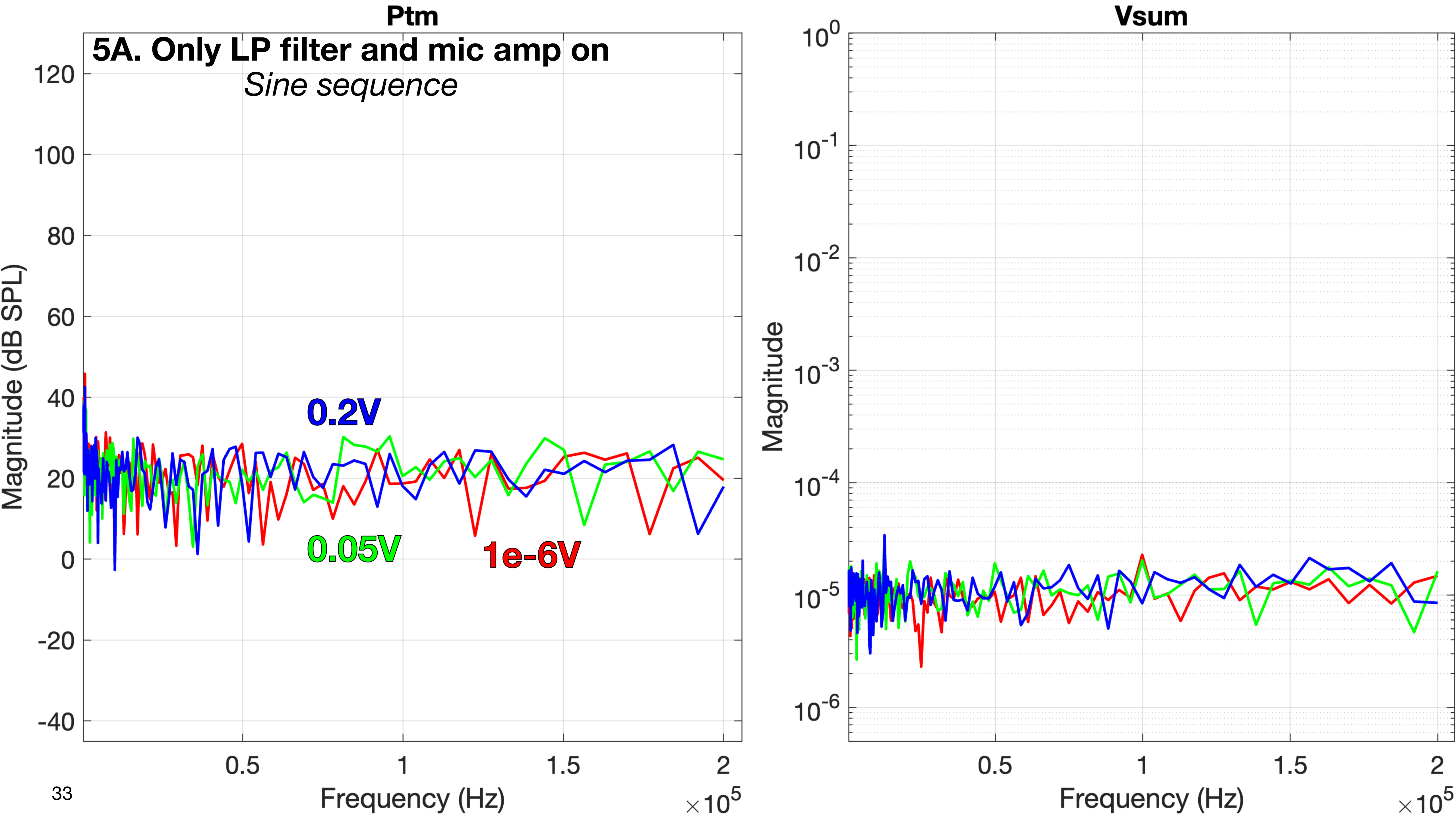

Ptm
5A. Only LP filter and mic amp on
Sine sequence
Magnitude (dB SPL)
120
100
80
60
40
20
0
-20
-40
0.2V
0.05V
1e-6V
0.5
1
1.5
2
Frequency (Hz)
×10^5
Vsum
Magnitude
10^0
10^-1
10^-2
10^-3
10^-4
10^-5
10^-6
0.5
1
1.5
2
Frequency (Hz)
×10^5

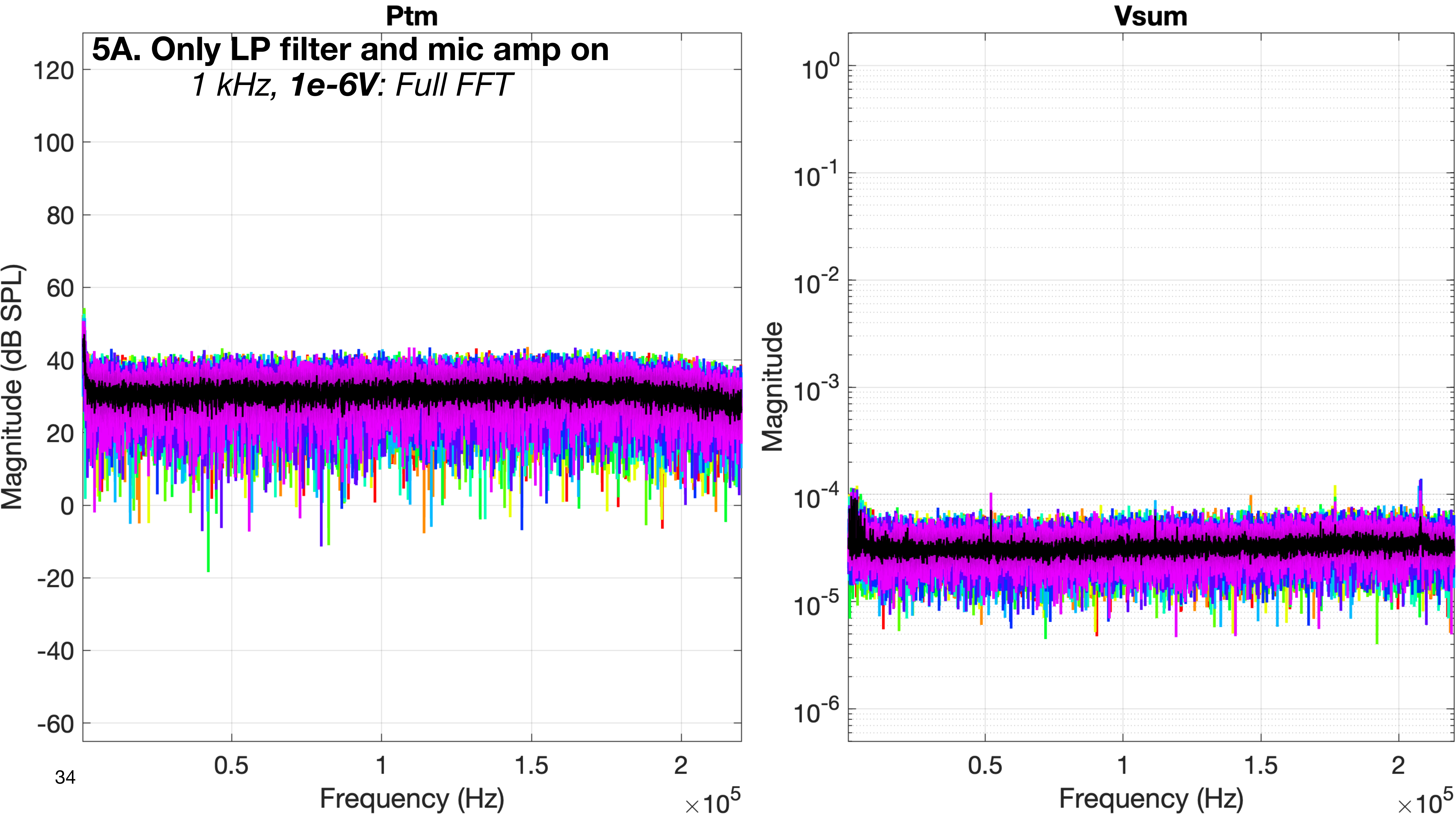

Ptm
5A. Only LP filter and mic amp on
1 kHz, 1e-6V: Full FFT
Magnitude (dB SPL)
Frequency (Hz)
×10^5
Vsum
Magnitude
Frequency (Hz)
×10^5

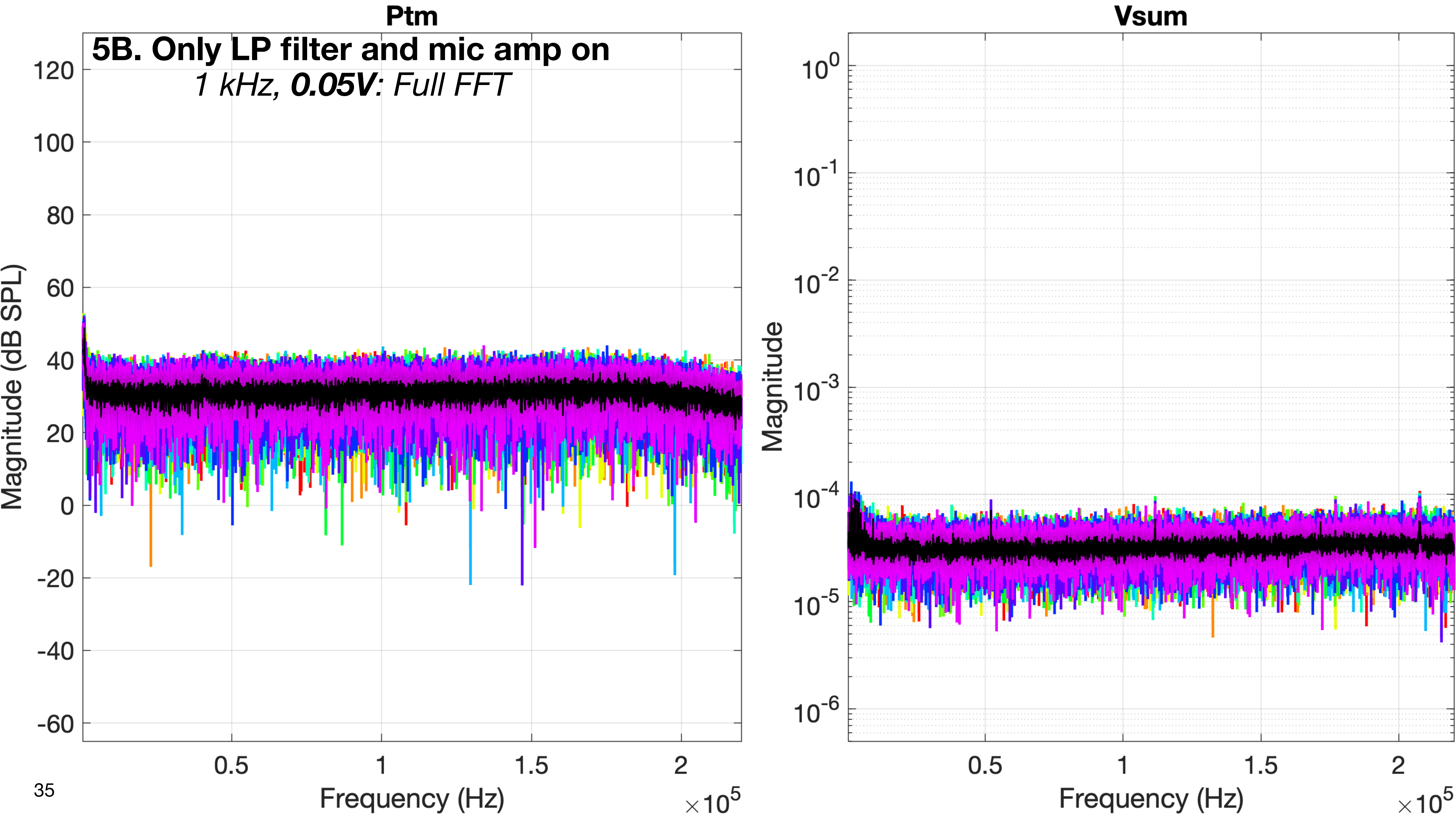
Ptm
5B. Only LP filter and mic amp on
1 kHz, 0.05V: Full FFT
Magnitude (dB SPL)
120
100
80
60
40
20
0
-20
-40
-60
0.5
1
1.5
2
Frequency (Hz)
×10^5
Vsum
Magnitude
10^0
10^-1
10^-2
10^-3
10^-4
10^-5
10^-6
0.5
1
1.5
2
Frequency (Hz)
×10^5

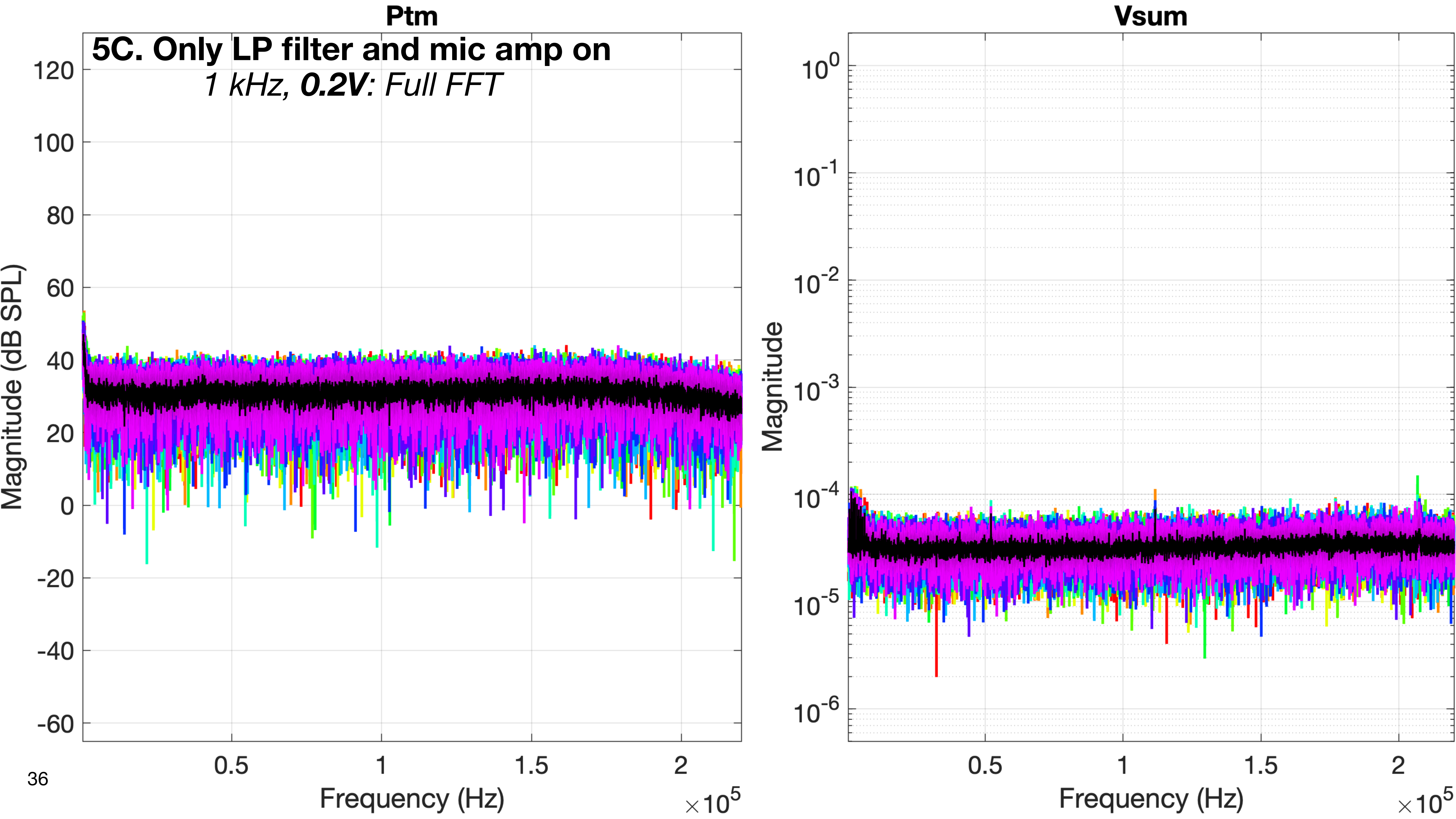

Ptm
5C. Only LP filter and mic amp on
1 kHz, 0.2V: Full FFT
Magnitude (dB SPL)
120
100
80
60
40
20
0
-20
-40
-60
0.5
1
1.5
2
Frequency (Hz)
×10^5
Vsum
Magnitude
10^0
10^-1
10^-2
10^-3
10^-4
10^-5
10^-6
0.5
1
1.5
2
Frequency (Hz)
×10^5

### 6. Power on JUST the 3384 and 3D LDV

- Wiring according to the normal schematic for measurements
- The 3384 LP filter is CONNECTED and powered ON
- The xMEMS microphone amp is CONNECTED and powered OFF (the microphone is in the fixture)
- The 3D LDV is CONNECTED and powered ON
- The xMEMS speaker driver is CONNECTED and powered OFF

- **Purpose:**
  - This is to establish the contribution of just the 3D LDV along its normal signal path when measuring live signals of vibrational noise on the undriven transducer and second eartip, with everything else off.

### 6. Power on JUST the 3384 and 3D LDV

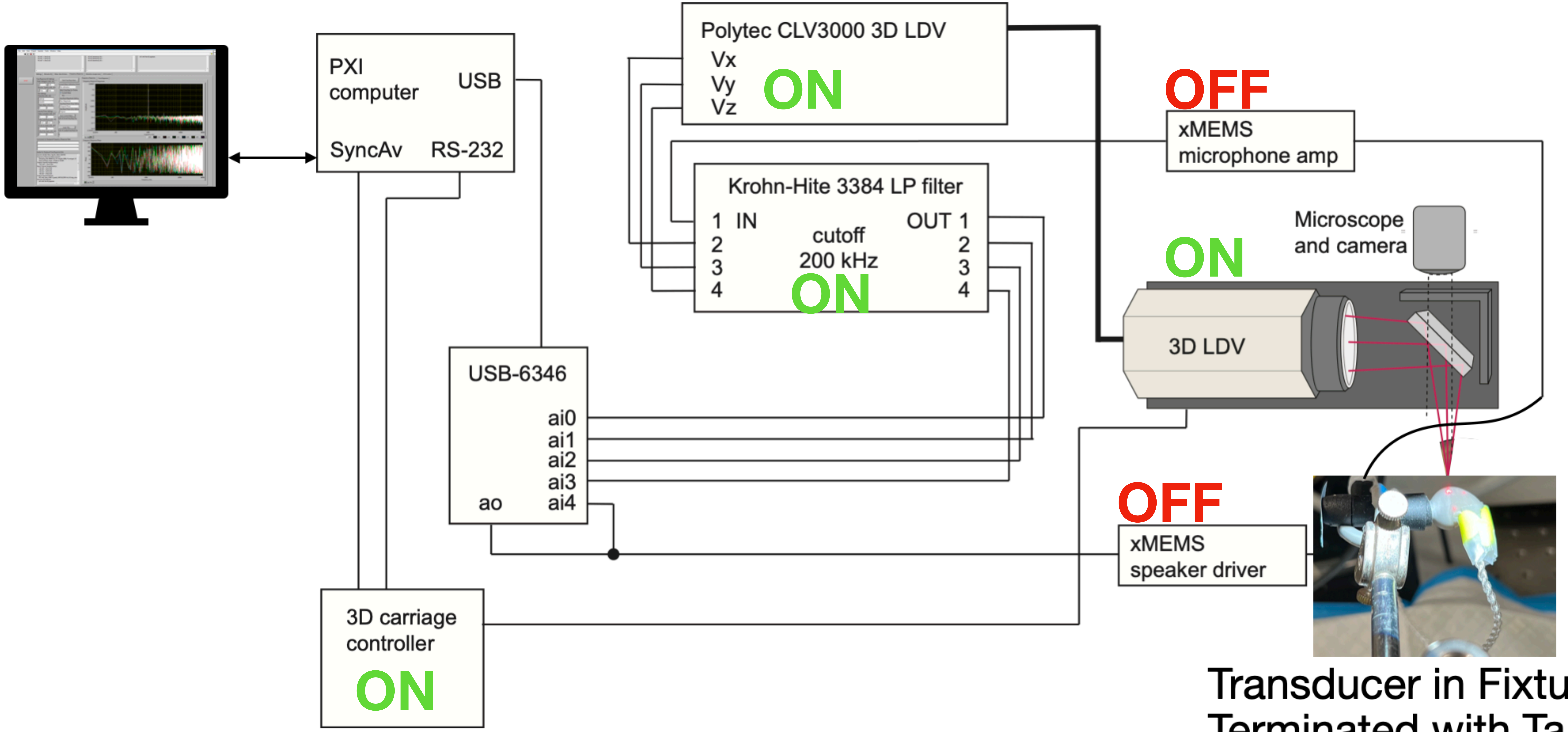


Transducer in Fixture,
Terminated with Tape

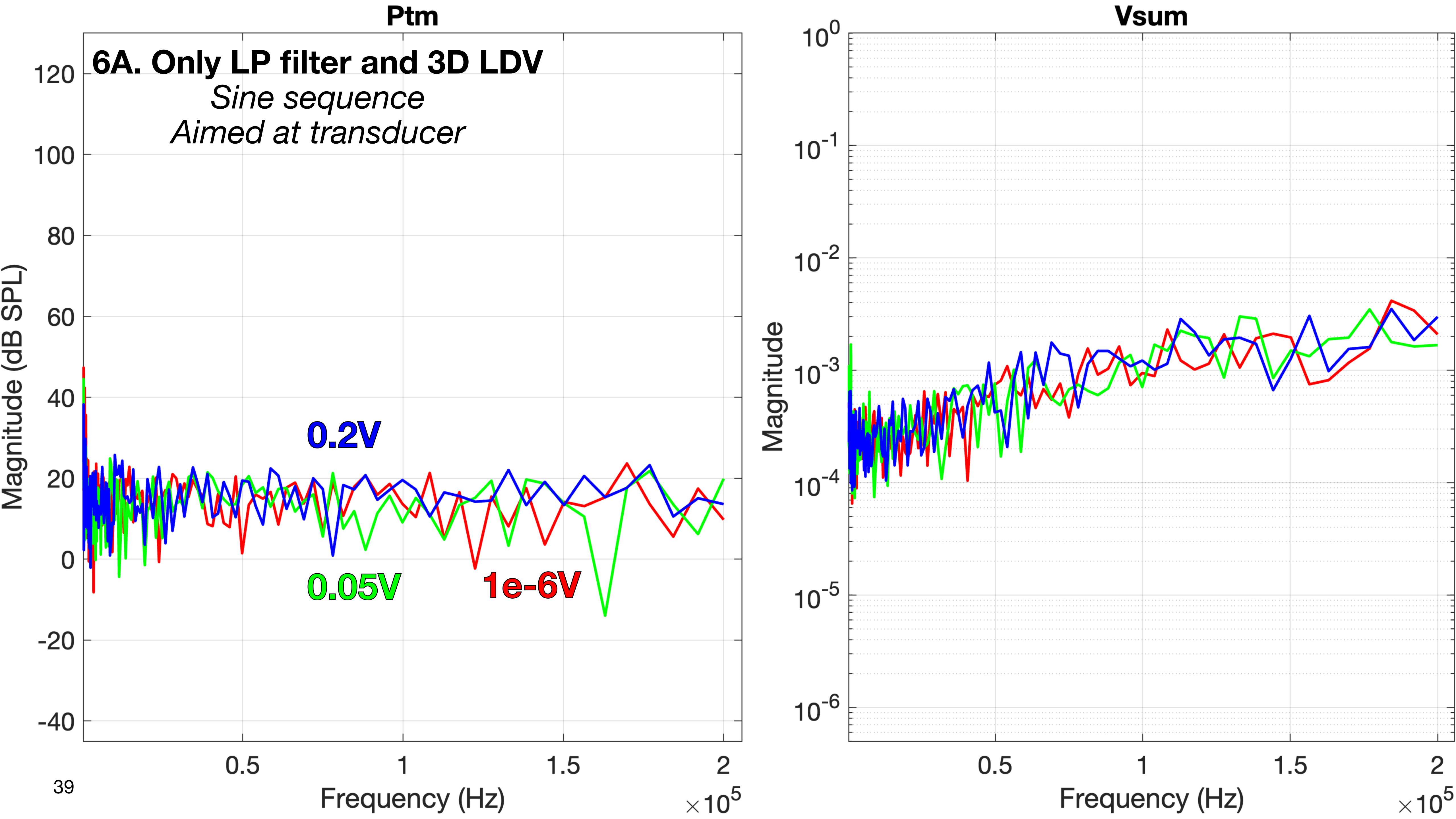

Ptm
6A. Only LP filter and 3D LDV
Sine sequence
Aimed at transducer
0.2V
0.05V
1e-6V
Magnitude (dB SPL)
120
100
80
60
40
20
0
-20
-40
0.5
1
1.5
2
Frequency (Hz)
×10^5
Vsum
10^0
10^-1
10^-2
10^-3
10^-4
10^-5
10^-6
Magnitude
0.5
1
1.5
2
Frequency (Hz)
×10^5

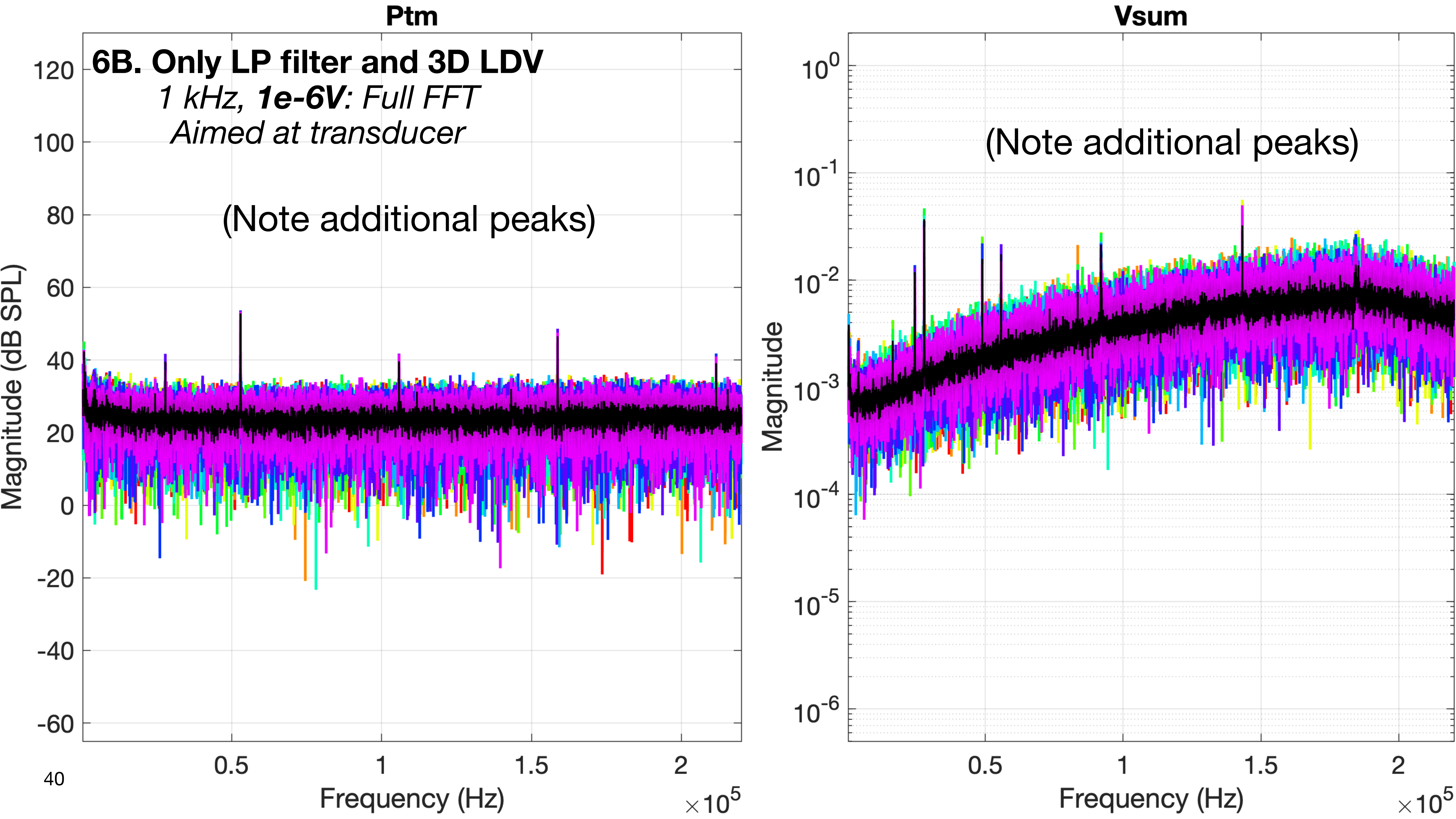
Ptm
6B. Only LP filter and 3D LDV
1 kHz, 1e-6V: Full FFT
Aimed at transducer
(Note additional peaks)
Magnitude (dB SPL)
120
100
80
60
40
20
0
-20
-40
-60
0.5
1
1.5
2
Frequency (Hz)
×10^5
Vsum
(Note additional peaks)
Magnitude
10^0
10^-1
10^-2
10^-3
10^-4
10^-5
10^-6
0.5
1
1.5
2
Frequency (Hz)
×10^5

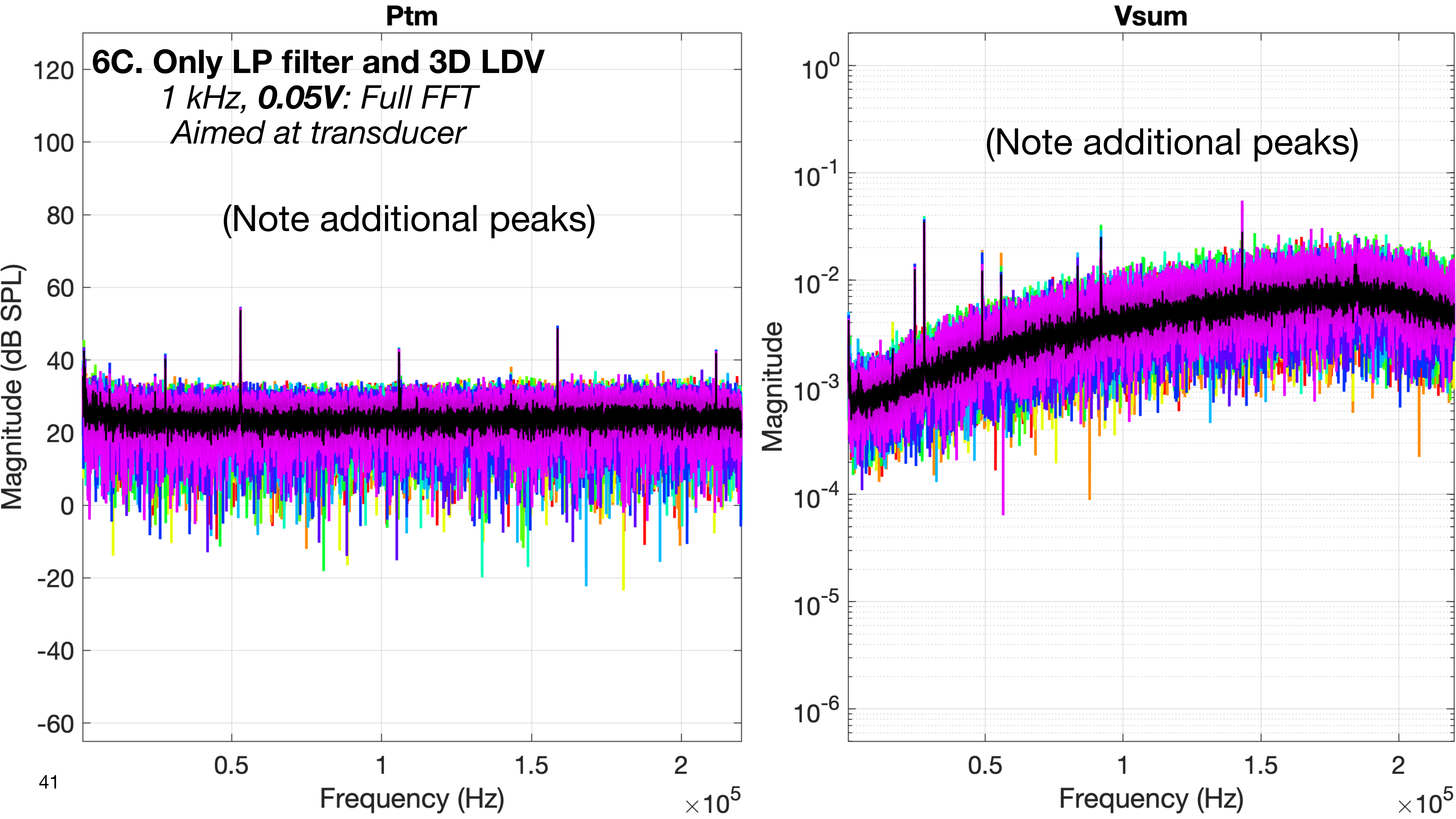

Ptm
6C. Only LP filter and 3D LDV
1 kHz, 0.05V: Full FFT
Aimed at transducer
(Note additional peaks)
Magnitude (dB SPL)
120
100
80
60
40
20
0
-20
-40
-60
0.5
1
1.5
2
Frequency (Hz)
×10^5
Vsum
(Note additional peaks)
Magnitude
10^0
10^-1
10^-2
10^-3
10^-4
10^-5
10^-6
0.5
1
1.5
2
Frequency (Hz)
×10^5

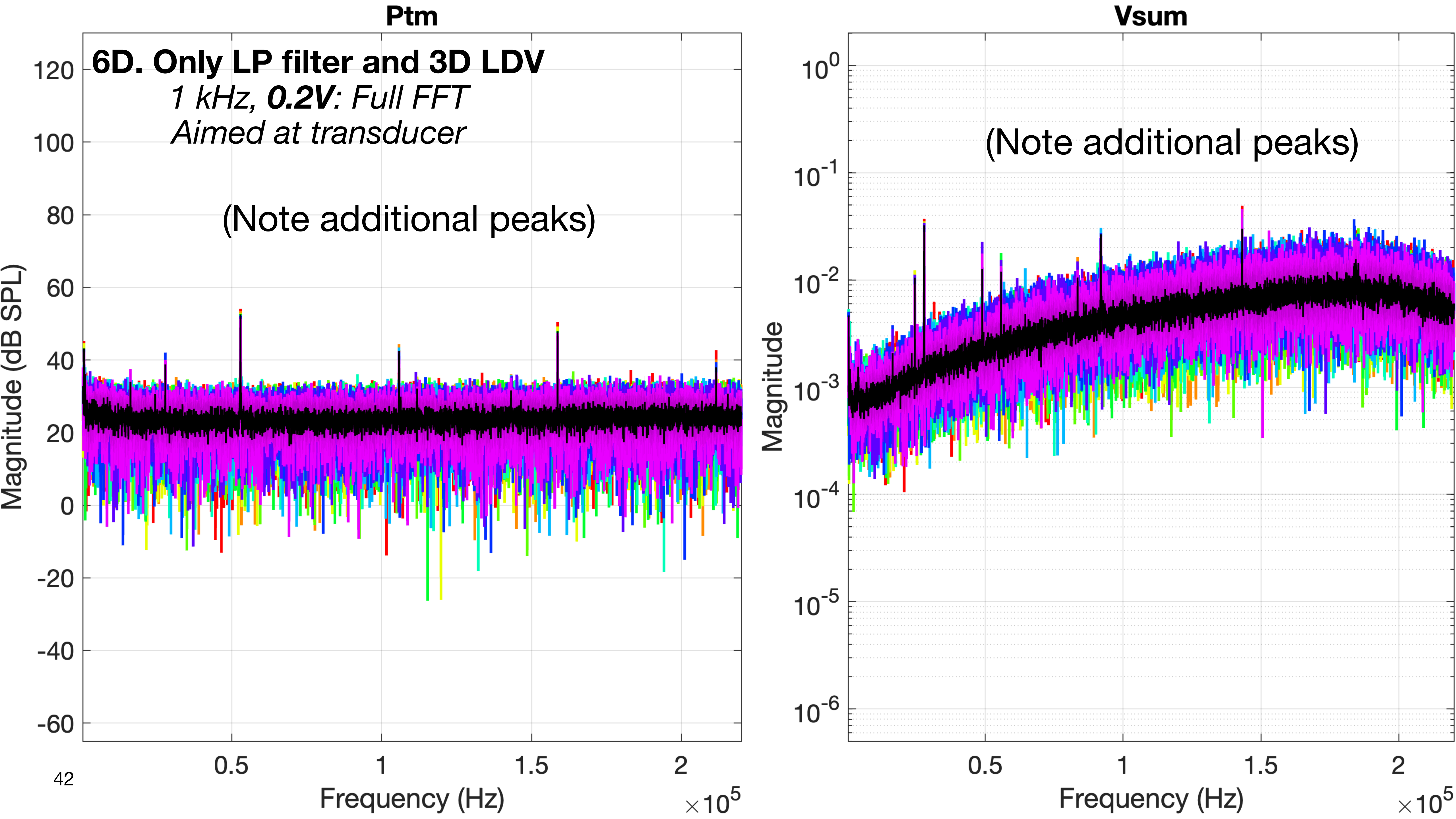
Ptm
6D. Only LP filter and 3D LDV
1 kHz, 0.2V: Full FFT
Aimed at transducer
(Note additional peaks)
Magnitude (dB SPL)
Frequency (Hz)
Vsum
(Note additional peaks)
Magnitude
Frequency (Hz)

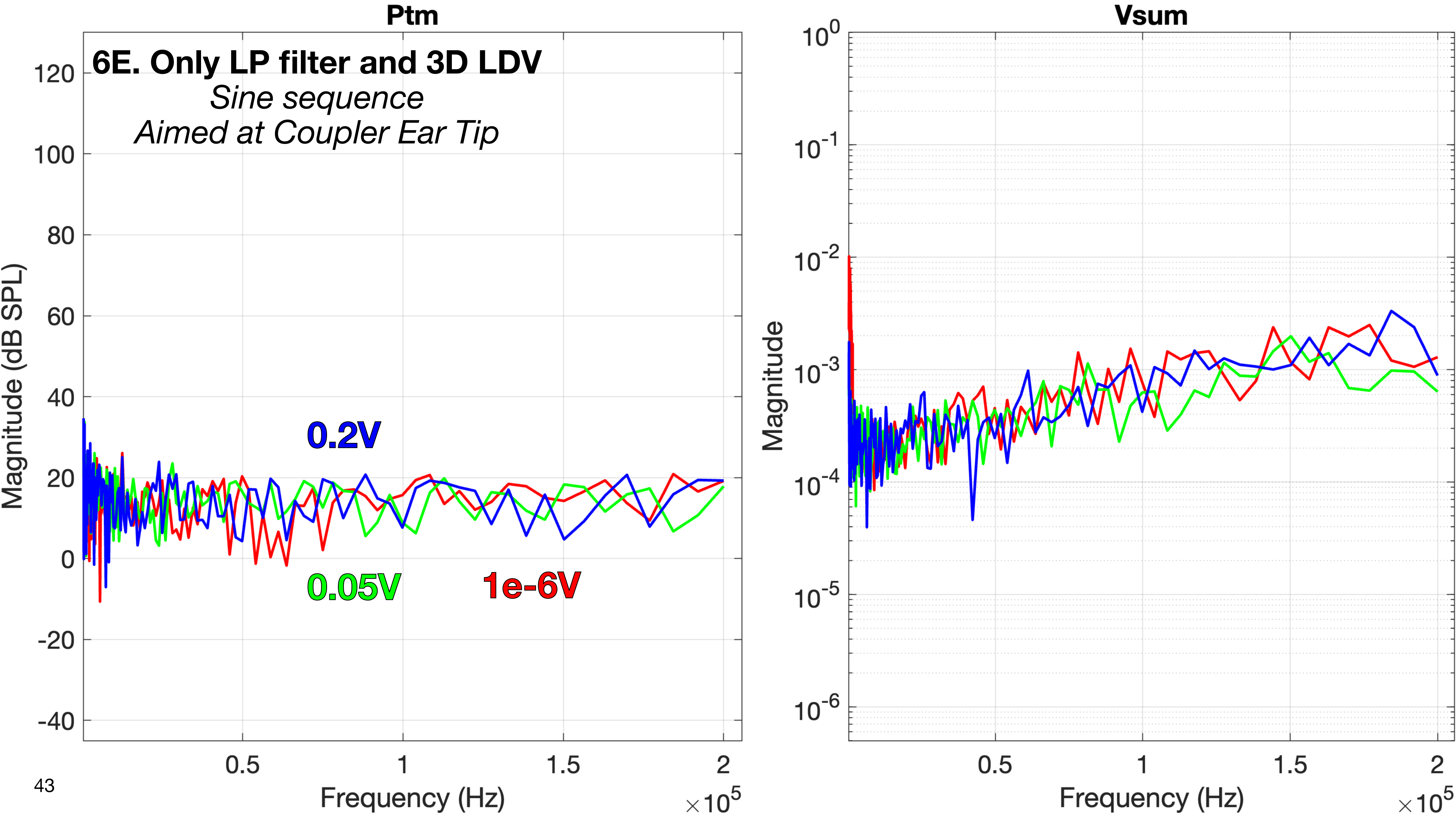
Ptm
6E. Only LP filter and 3D LDV
Sine sequence
Aimed at Coupler Ear Tip
0.2V
0.05V
1e-6V
Magnitude (dB SPL)
120
100
80
60
40
20
0
-20
-40
0.5
1
1.5
2
Frequency (Hz)
$\times 10^5$
Vsum
Magnitude
$10^0$
$10^{-1}$
$10^{-2}$
$10^{-3}$
$10^{-4}$
$10^{-5}$
$10^{-6}$
0.5
1
1.5
2
Frequency (Hz)
$\times 10^5$

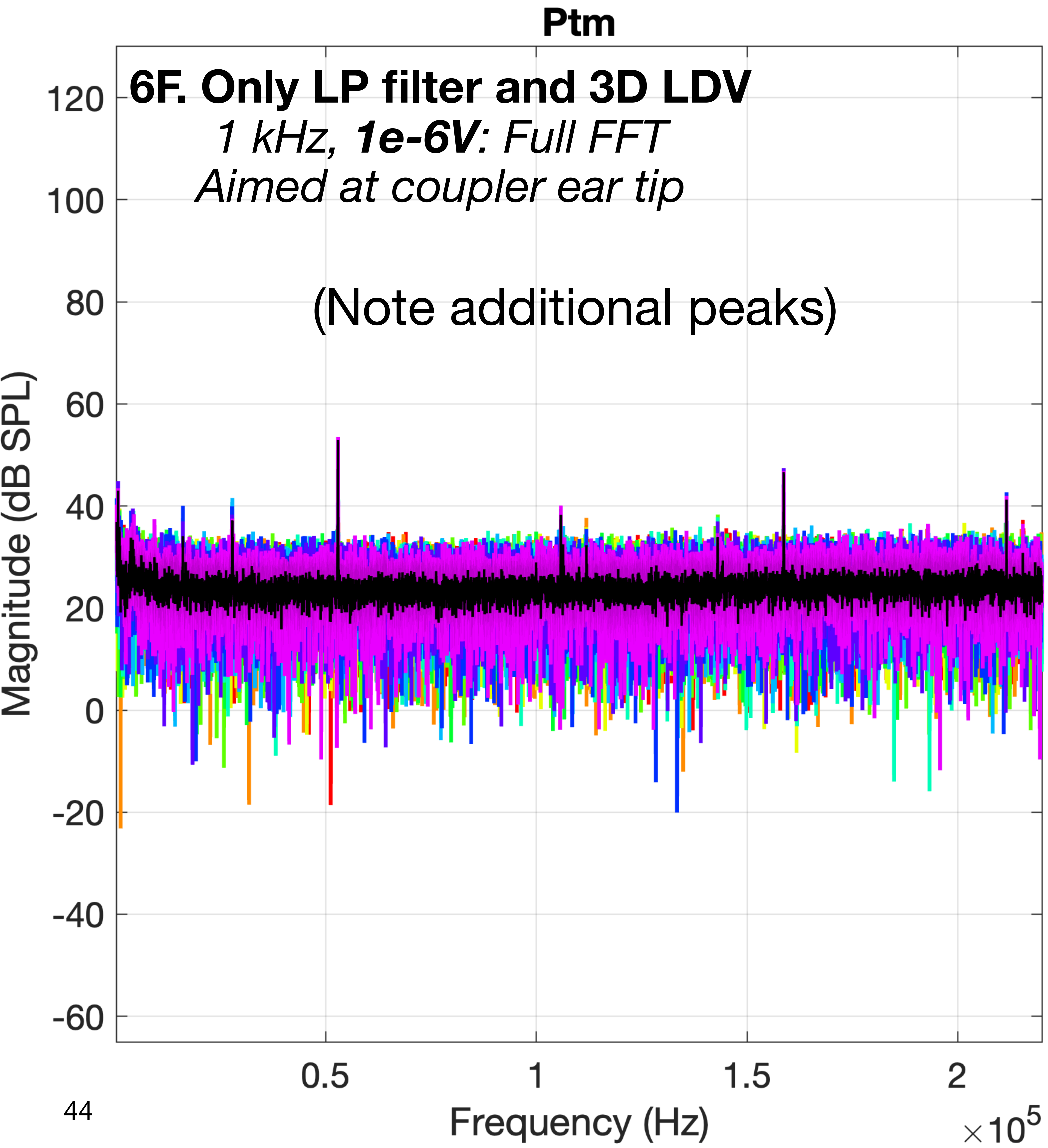
Ptm
6F. Only LP filter and 3D LDV
1 kHz, 1e-6V: Full FFT
Aimed at coupler ear tip
(Note additional peaks)
Magnitude (dB SPL)
120
100
80
60
40
20
0
-20
-40
-60
0.5
1
1.5
2
Frequency (Hz)
×10^5

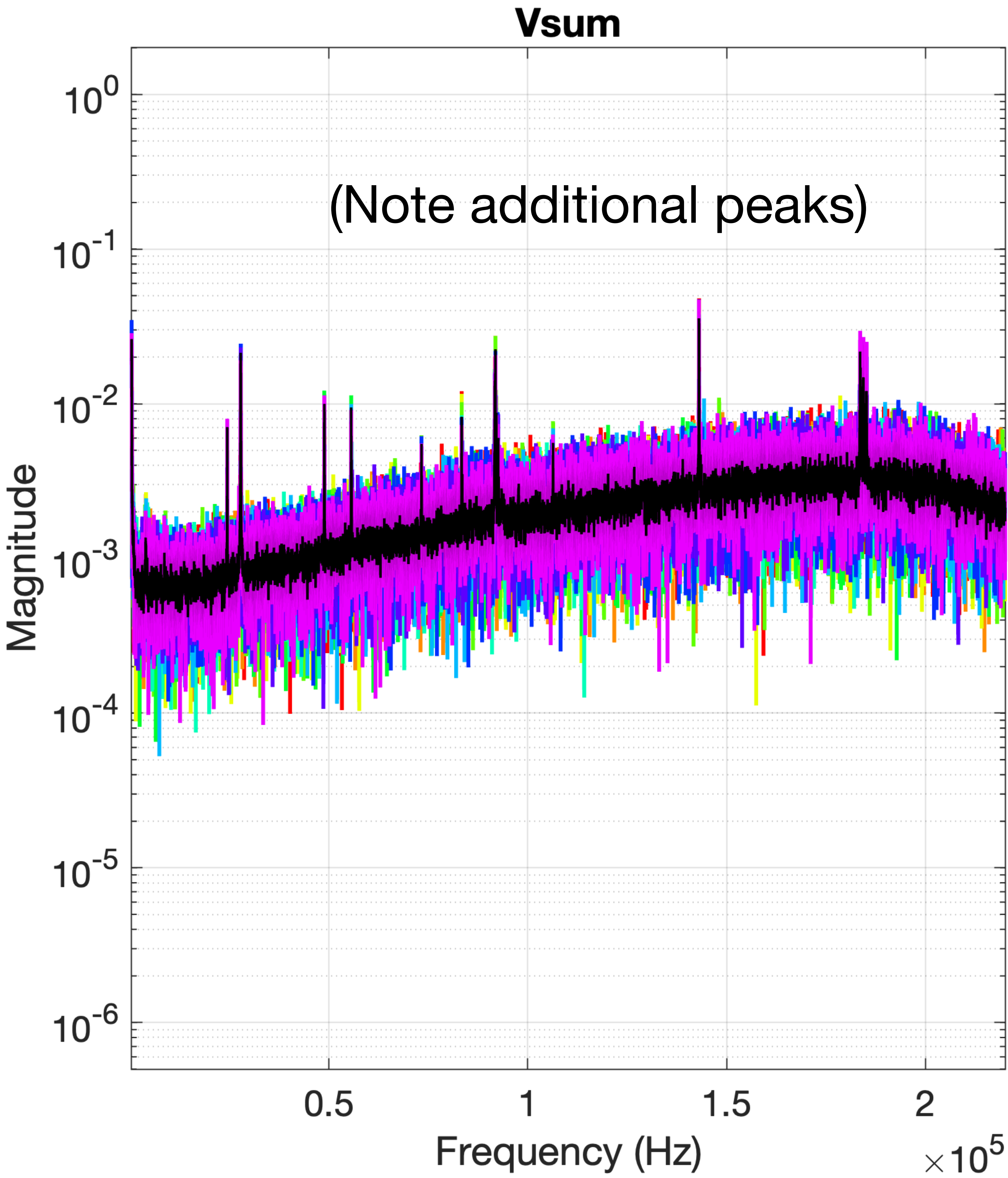
Vsum
(Note additional peaks)
Magnitude
10^0
10^-1
10^-2
10^-3
10^-4
10^-5
10^-6
0.5
1
1.5
2
Frequency (Hz)
×10^5

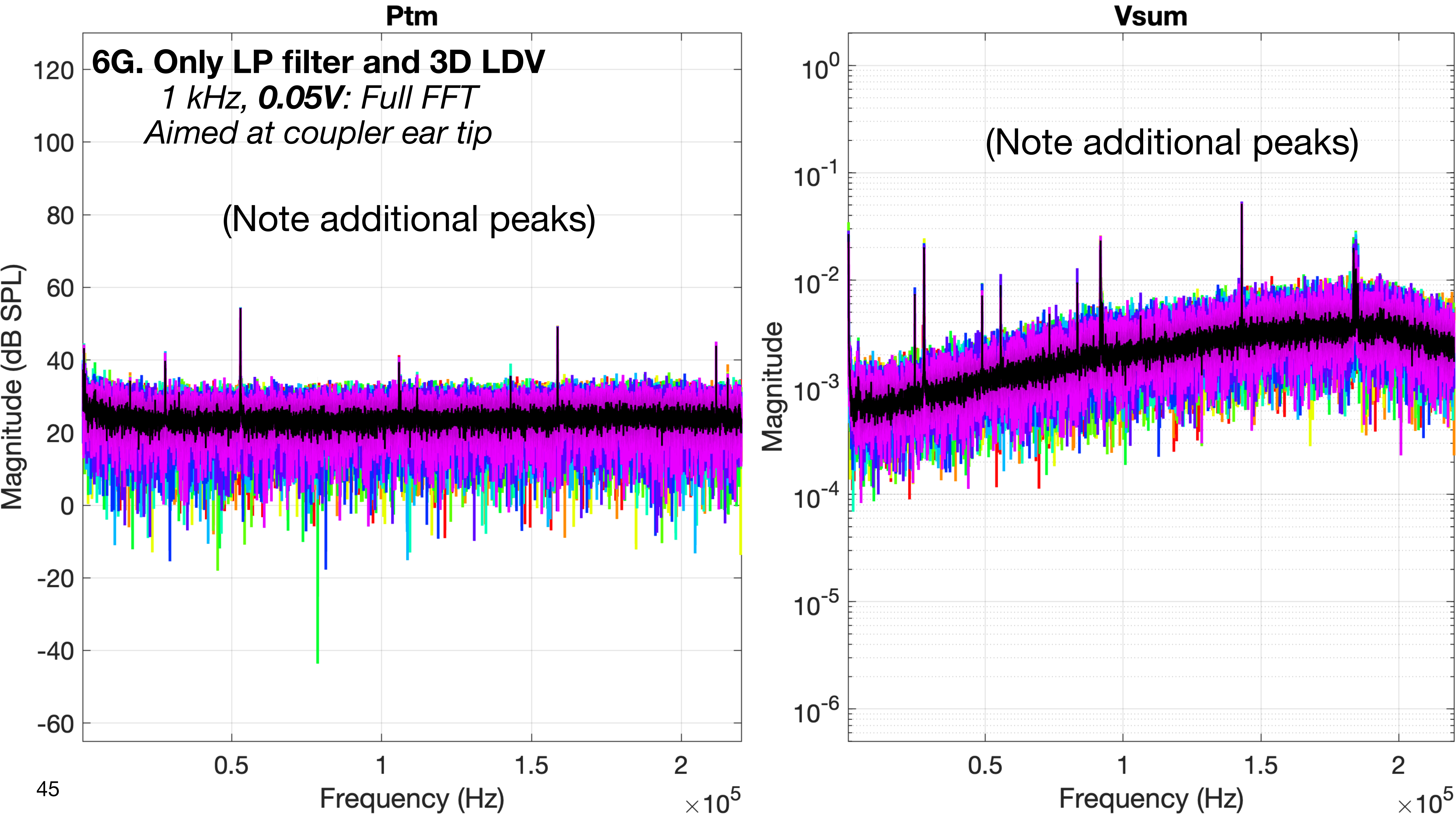
Ptm
6G. Only LP filter and 3D LDV
1 kHz, 0.05V: Full FFT
Aimed at coupler ear tip
(Note additional peaks)
Magnitude (dB SPL)
120
100
80
60
40
20
0
-20
-40
-60
0.5
1
1.5
2
Frequency (Hz)
$\times 10^5$
Vsum
(Note additional peaks)
Magnitude
$10^0$
$10^{-1}$
$10^{-2}$
$10^{-3}$
$10^{-4}$
$10^{-5}$
$10^{-6}$
0.5
1
1.5
2
Frequency (Hz)
$\times 10^5$

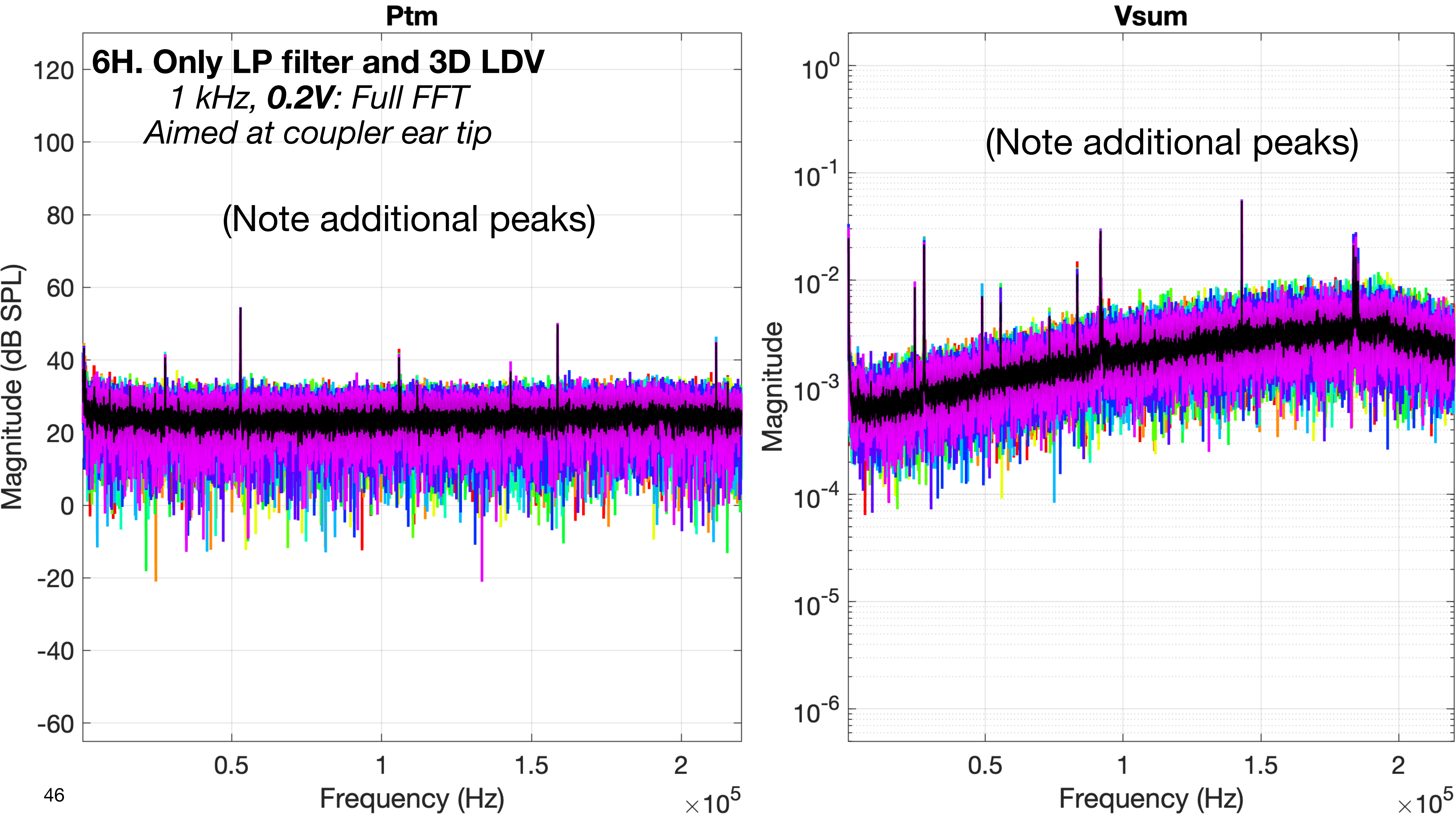

Ptm
6H. Only LP filter and 3D LDV
1 kHz, 0.2V: Full FFT
Aimed at coupler ear tip
(Note additional peaks)
Magnitude (dB SPL)
120
100
80
60
40
20
0
-20
-40
-60
0.5
1
1.5
2
Frequency (Hz)
×10^5
Vsum
(Note additional peaks)
Magnitude
10^0
10^-1
10^-2
10^-3
10^-4
10^-5
10^-6
0.5
1
1.5
2
Frequency (Hz)
×10^5

### 7. Power on everything EXCEPT the xMEMS speaker driver

- Wiring according to the normal schematic for measurements
- The 3384 LP filter is CONNECTED and powered ON
- The xMEMS microphone amp is CONNECTED and powered ON (the microphone is in the fixture)
- The 3D LDV is CONNECTED and powered ON
- The xMEMS speaker driver is CONNECTED and powered OFF

- **Purpose:**
  - This is to establish the contributions of the xMEMS microphone and 3D LDV along their normal signal paths when measuring live signals of acoustic and vibrational noise on the undriven transducer and second eartip, with everything else off.

### 7. Power on everything EXCEPT the xMEMS speaker driver

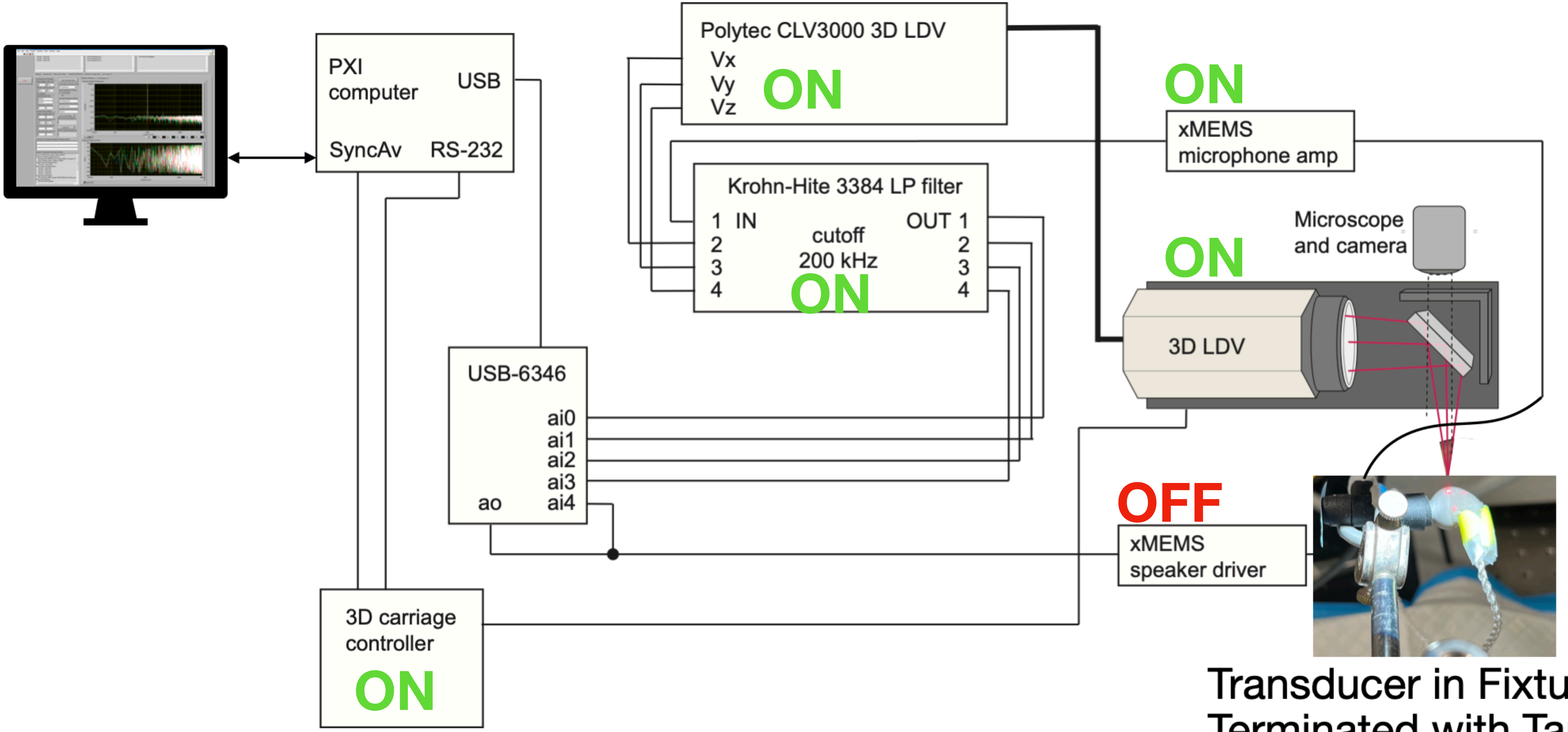


Transducer in Fixture, Terminated with Tape

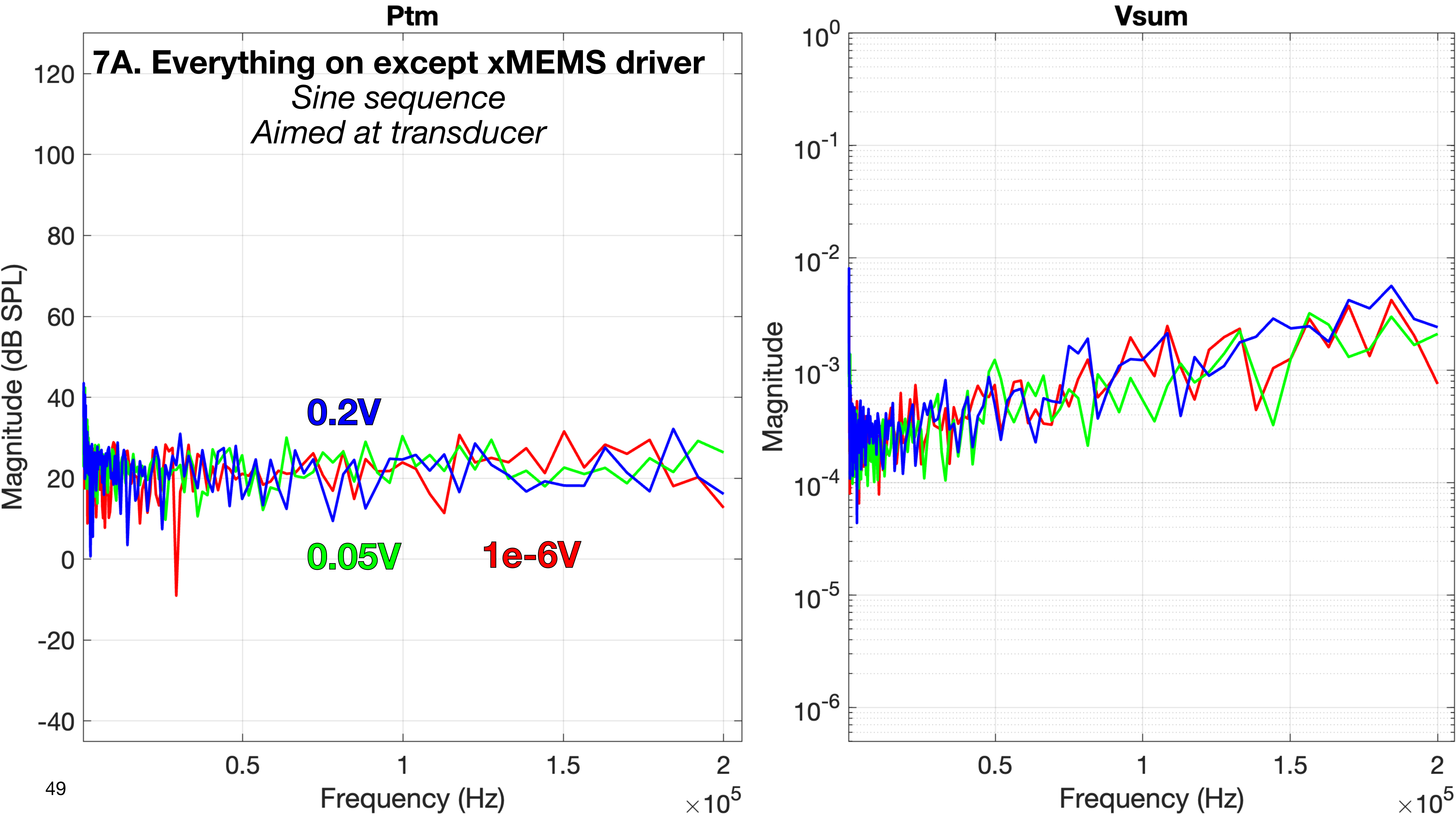

Ptm
7A. Everything on except xMEMS driver
Sine sequence
Aimed at transducer
Magnitude (dB SPL)
120
100
80
60
40
20
0
-20
-40
0.2V
0.05V
1e-6V
0.5
1
1.5
2
Frequency (Hz)
×10^5
Vsum
Magnitude
10^0
10^-1
10^-2
10^-3
10^-4
10^-5
10^-6
0.5
1
1.5
2
Frequency (Hz)
×10^5

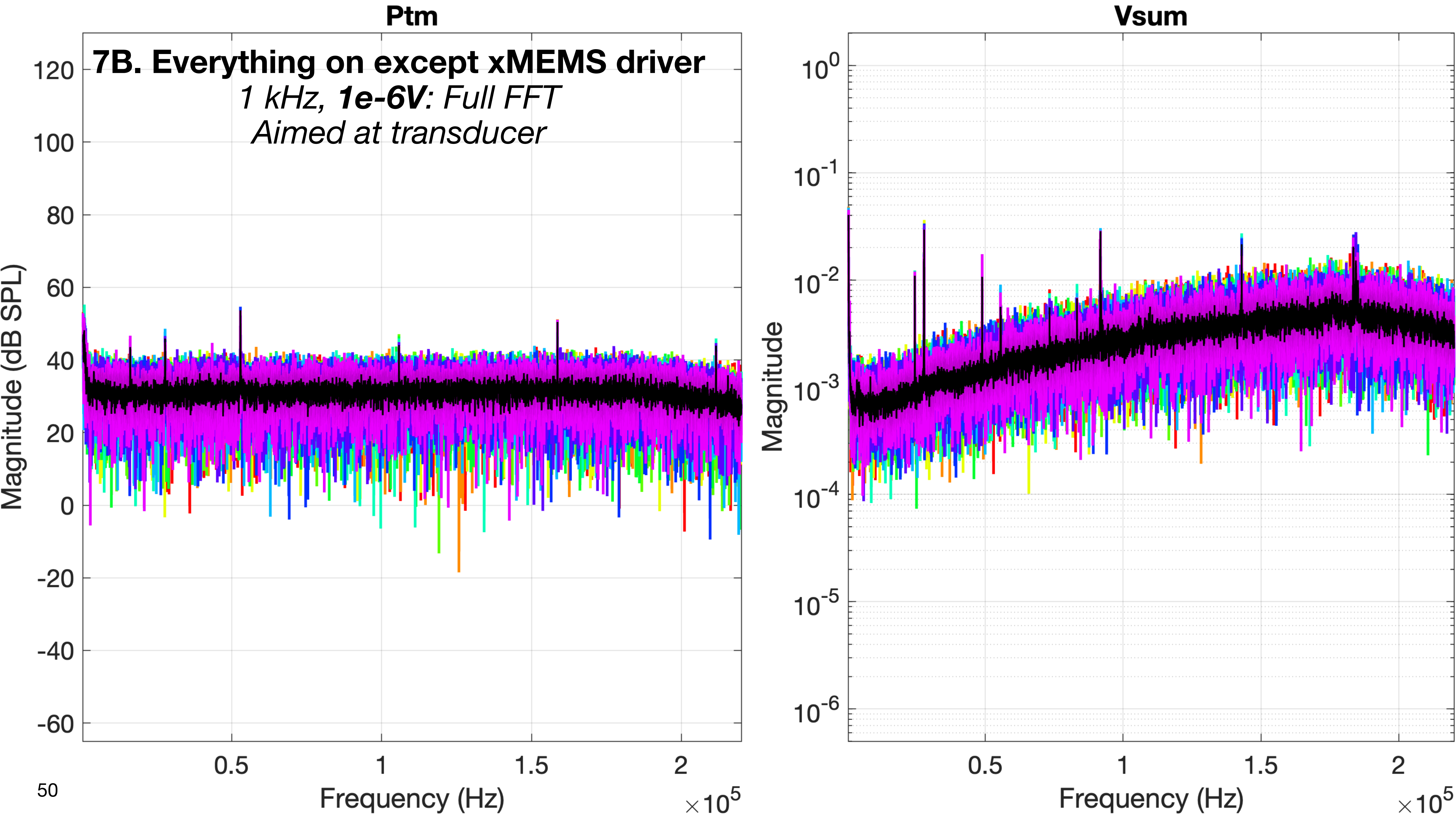
Ptm
Vsum
7B. Everything on except xMEMS driver
1 kHz, 1e-6V: Full FFT
Aimed at transducer
Magnitude (dB SPL)
Magnitude
Frequency (Hz)
Frequency (Hz)
×10^5
×10^5

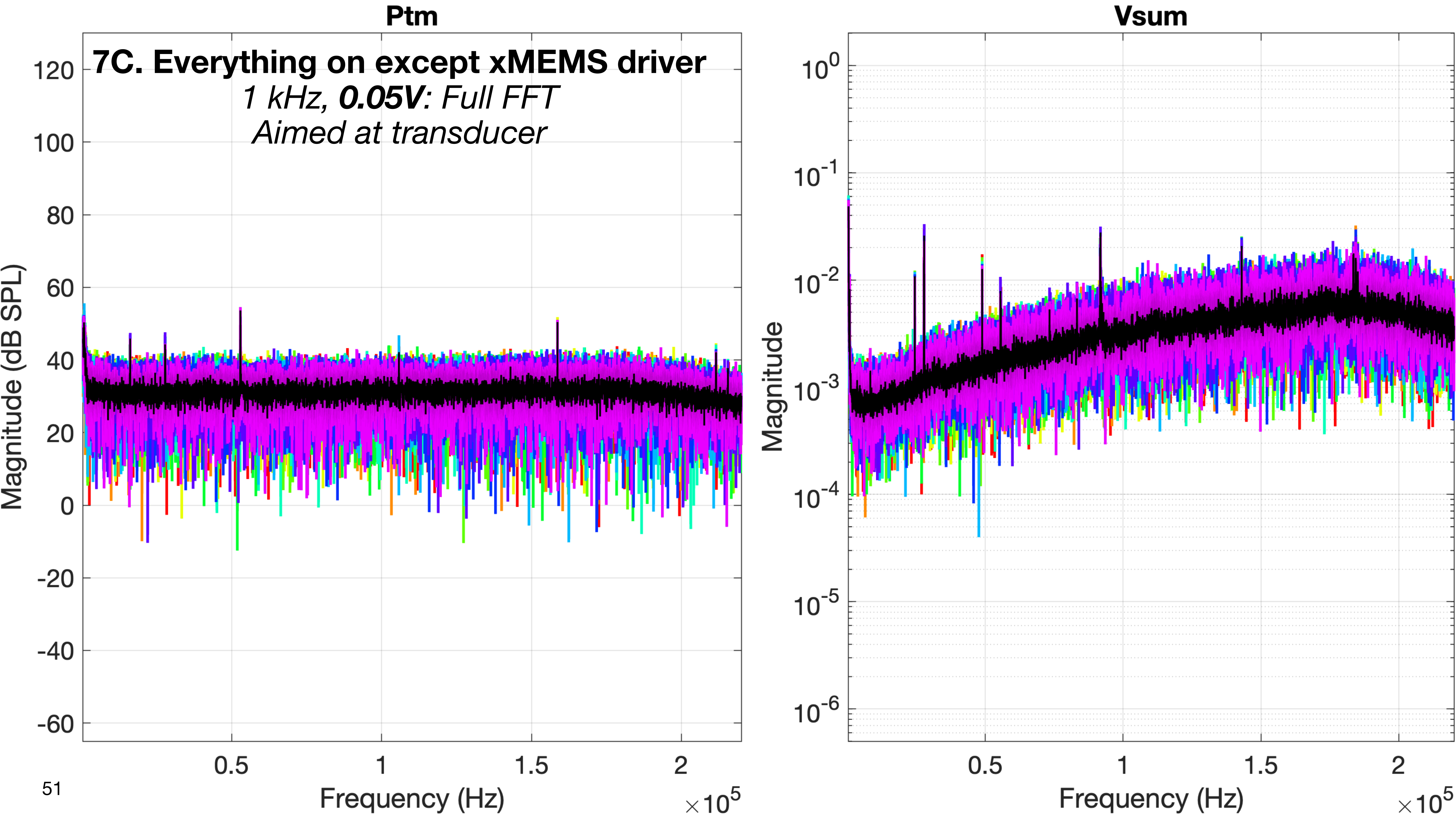
Ptm
7C. Everything on except xMEMS driver
1 kHz, 0.05V: Full FFT
Aimed at transducer
Magnitude (dB SPL)
120
100
80
60
40
20
0
-20
-40
-60
0.5
1
1.5
2
Frequency (Hz)
×10^5
Vsum
Magnitude
10^0
10^-1
10^-2
10^-3
10^-4
10^-5
10^-6
0.5
1
1.5
2
Frequency (Hz)
×10^5

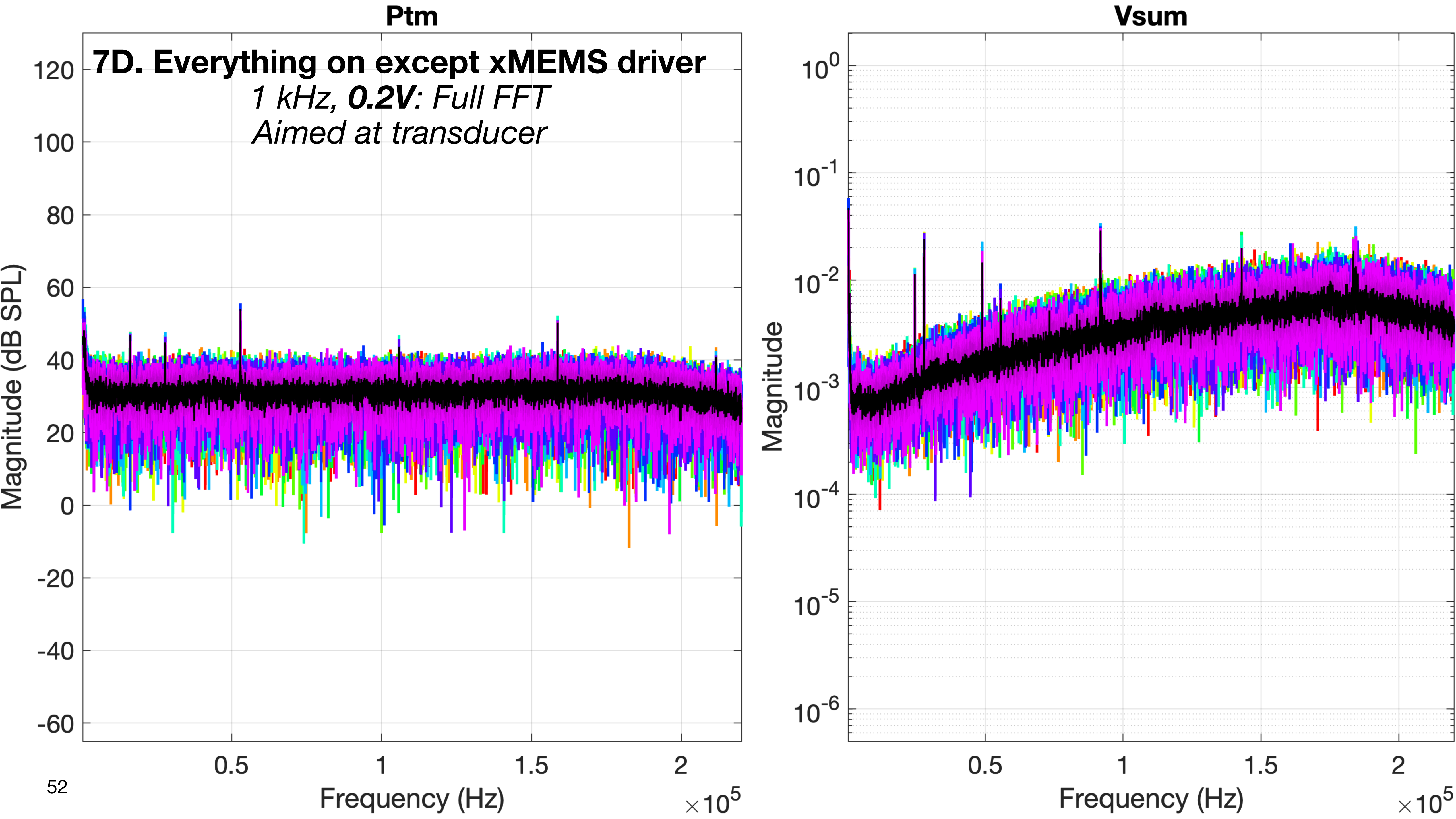
Ptm
7D. Everything on except xMEMS driver
1 kHz, 0.2V: Full FFT
Aimed at transducer
Magnitude (dB SPL)
120
100
80
60
40
20
0
-20
-40
-60
0.5
1
1.5
2
Frequency (Hz)
×10^5
Vsum
Magnitude
10^0
10^-1
10^-2
10^-3
10^-4
10^-5
10^-6
0.5
1
1.5
2
Frequency (Hz)
×10^5

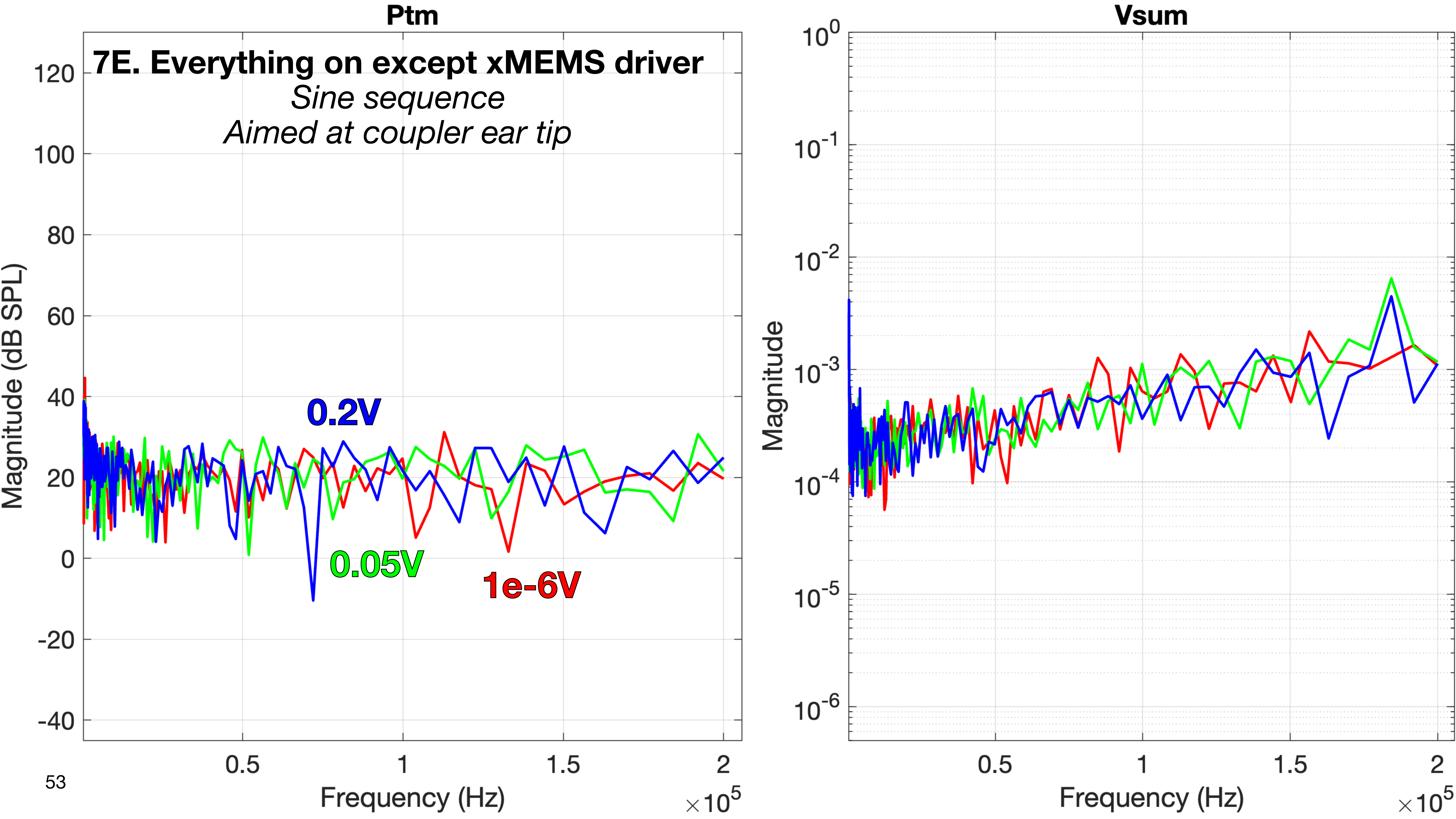

Ptm
7E. Everything on except xMEMS driver
Sine sequence
Aimed at coupler ear tip
Magnitude (dB SPL)
120
100
80
60
40
20
0
-20
-40
0.2V
0.05V
1e-6V
0.5
1
1.5
2
Frequency (Hz)
$\times 10^5$
Vsum
Magnitude
$10^0$
$10^{-1}$
$10^{-2}$
$10^{-3}$
$10^{-4}$
$10^{-5}$
$10^{-6}$
0.5
1
1.5
2
Frequency (Hz)
$\times 10^5$

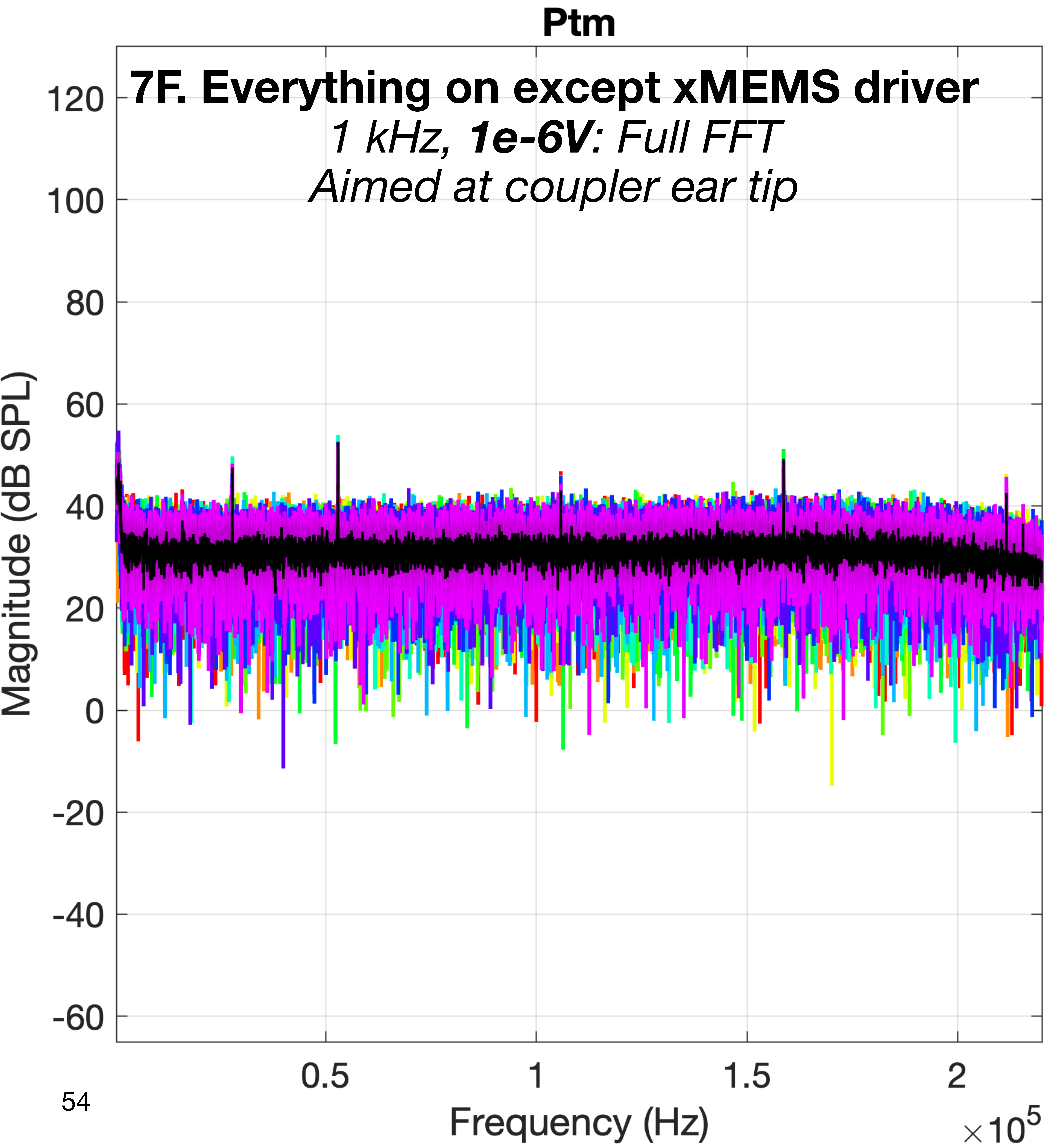
Ptm
7F. Everything on except xMEMS driver
1 kHz, 1e-6V: Full FFT
Aimed at coupler ear tip
Magnitude (dB SPL)
120
100
80
60
40
20
0
-20
-40
-60
0.5
1
1.5
2
Frequency (Hz)
×10^5

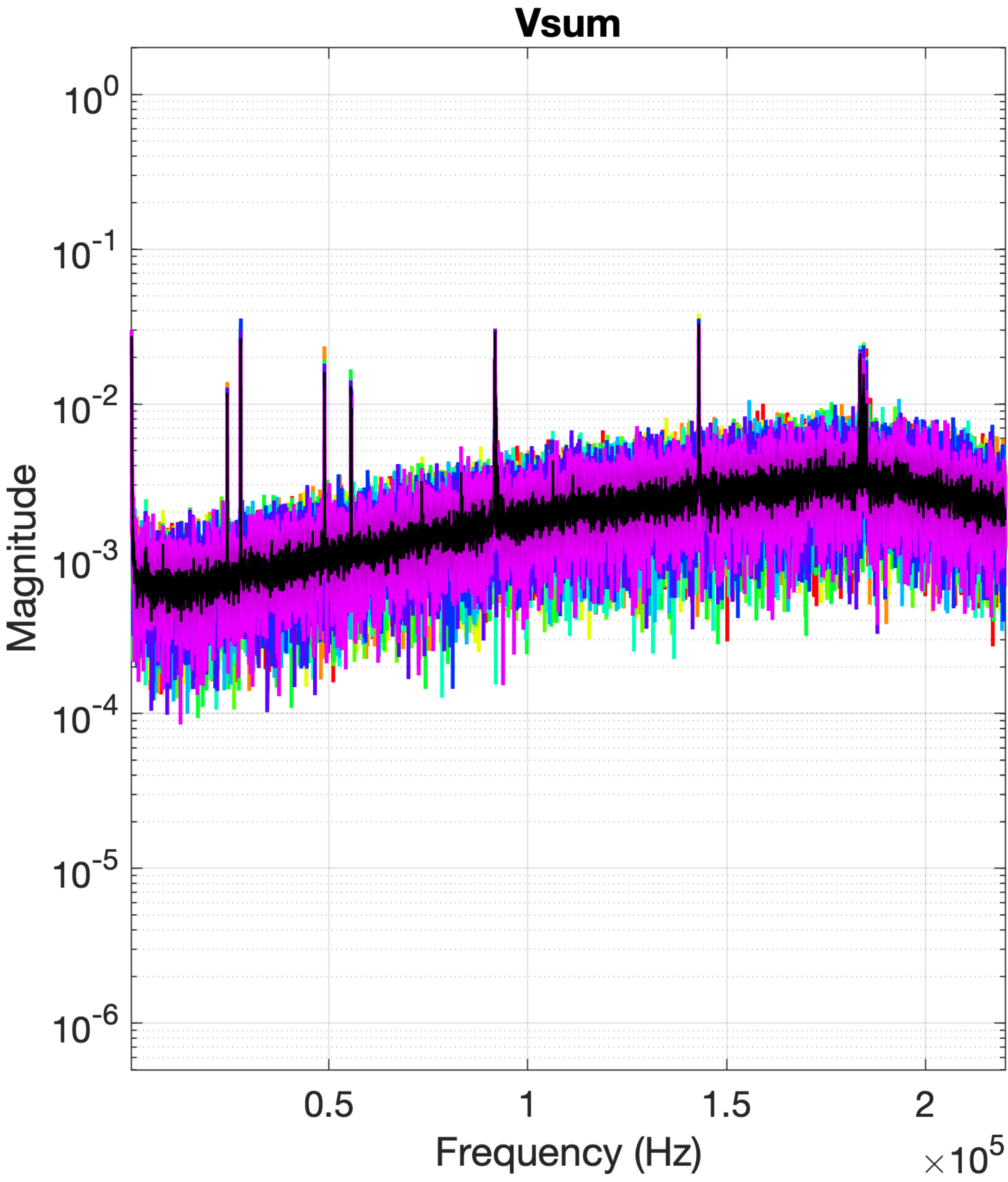
Vsum
Magnitude
10^0
10^-1
10^-2
10^-3
10^-4
10^-5
10^-6
0.5
1
1.5
2
Frequency (Hz)
×10^5

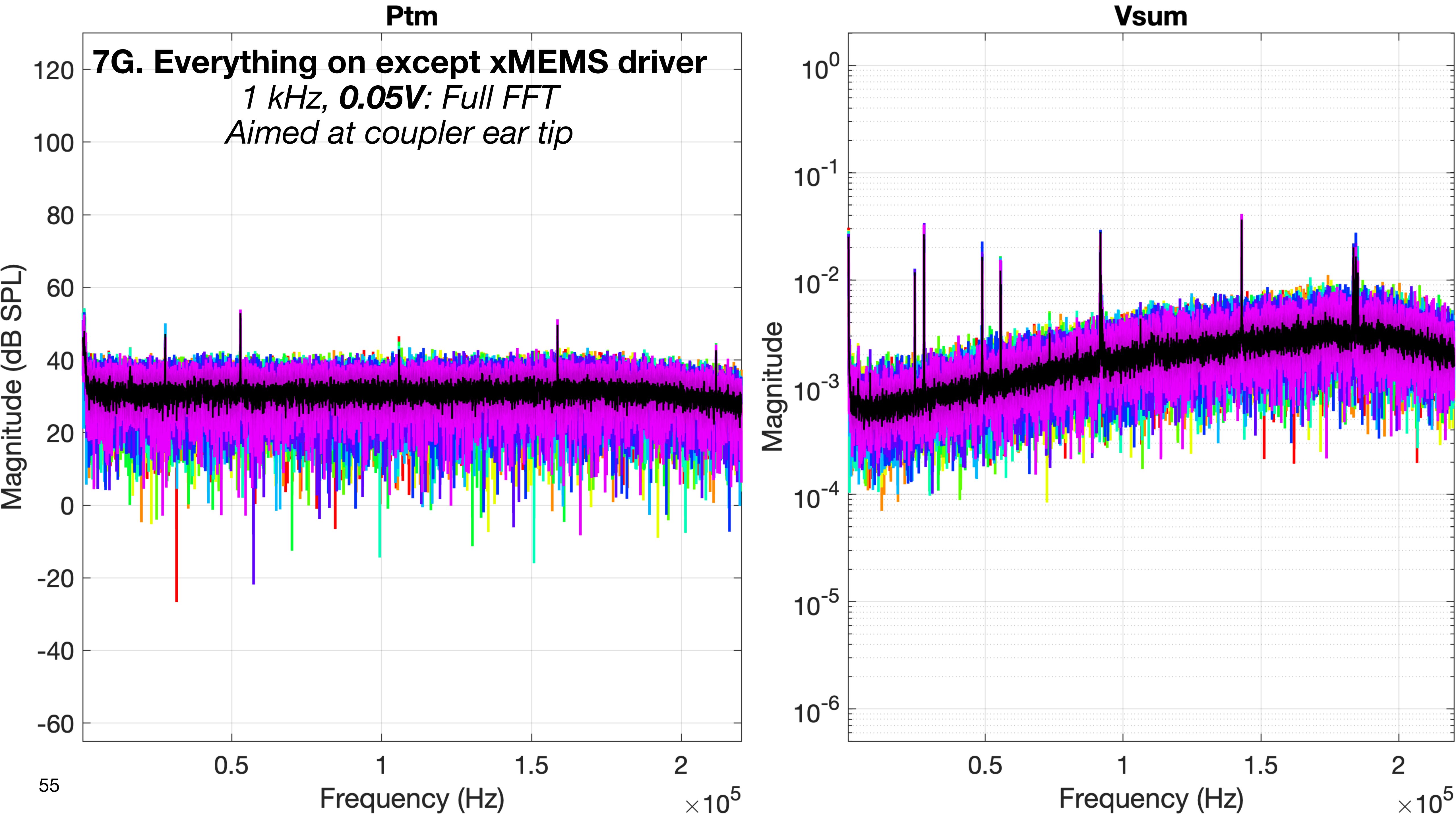
Ptm
7G. Everything on except xMEMS driver
1 kHz, 0.05V: Full FFT
Aimed at coupler ear tip
Magnitude (dB SPL)
120
100
80
60
40
20
0
-20
-40
-60
0.5
1
1.5
2
Frequency (Hz)
×10^5
Vsum
Magnitude
10^0
10^-1
10^-2
10^-3
10^-4
10^-5
10^-6
0.5
1
1.5
2
Frequency (Hz)
×10^5

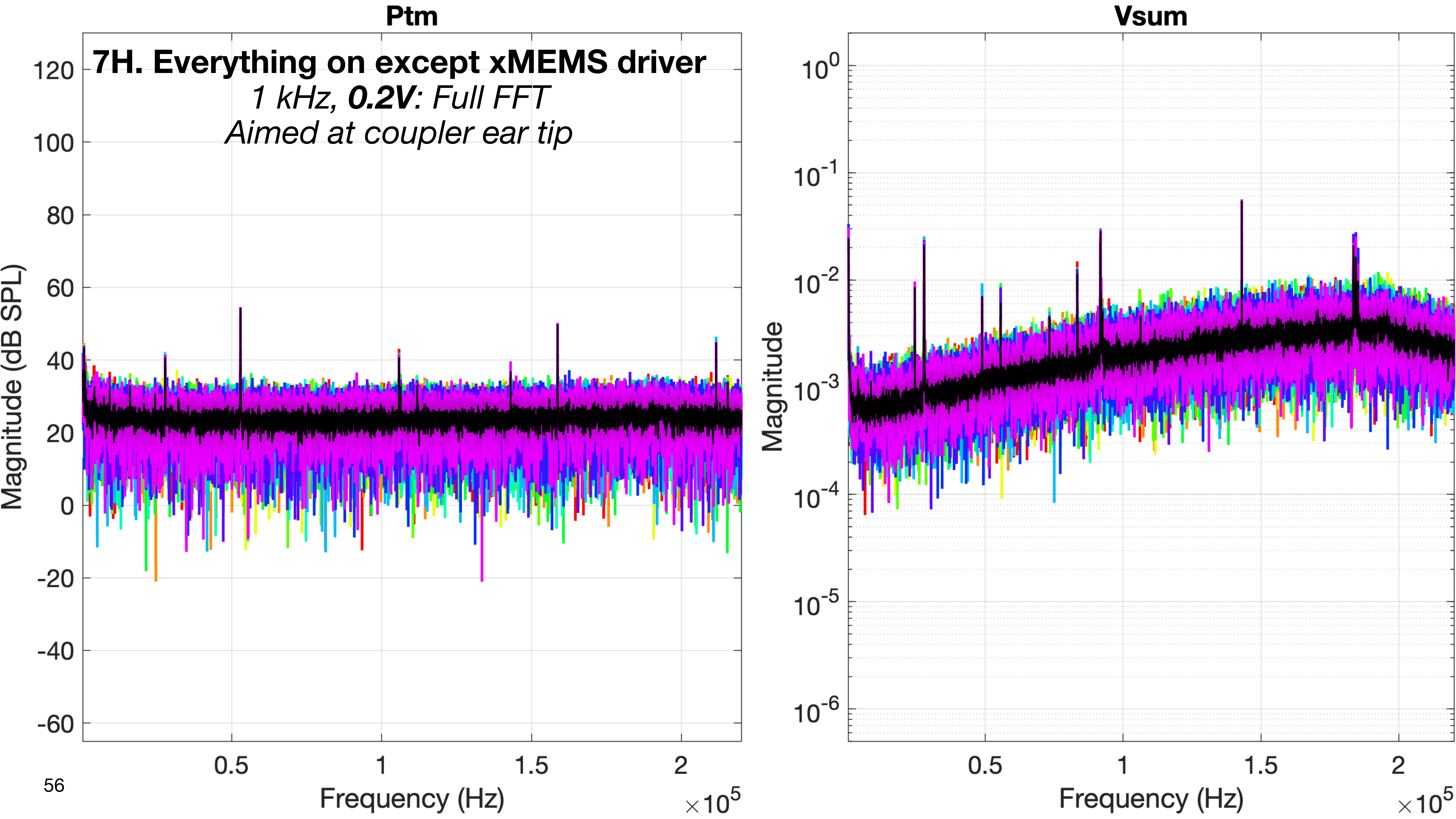

Ptm
7H. Everything on except xMEMS driver
1 kHz, 0.2V: Full FFT
Aimed at coupler ear tip
120
100
80
60
40
20
0
-20
-40
-60
Magnitude (dB SPL)
0.5
1
1.5
2
Frequency (Hz)
×10^5
Vsum
10^0
10^-1
10^-2
10^-3
10^-4
10^-5
10^-6
Magnitude
0.5
1
1.5
2
Frequency (Hz)
×10^5

### 8. Power on EVERYTHING

- Wiring according to the normal schematic for measurements
- The 3384 LP filter is CONNECTED and powered ON
- The xMEMS microphone amp is CONNECTED and powered ON (the microphone is in the fixture)
- The 3D LDV is CONNECTED and powered ON, with the LDV aimed at the transducer
- The xMEMS speaker driver is CONNECTED and powered ON

- **Purpose:**
  - This is to establish the baselines for the test fixture (with tape instead of a temporal bone) when all components of the measurement setup are connected and powered on normally.

### 8. Power on EVERYTHING

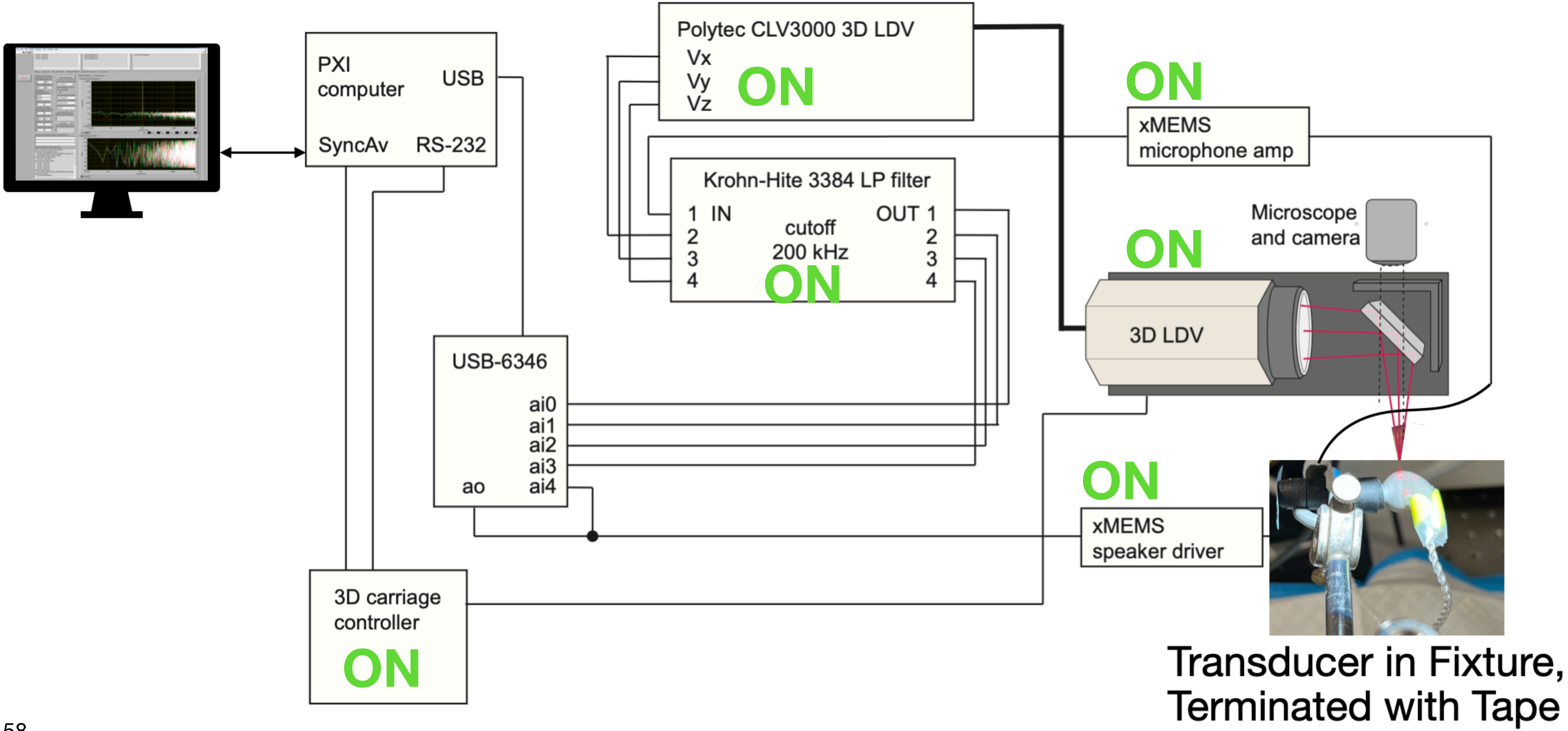

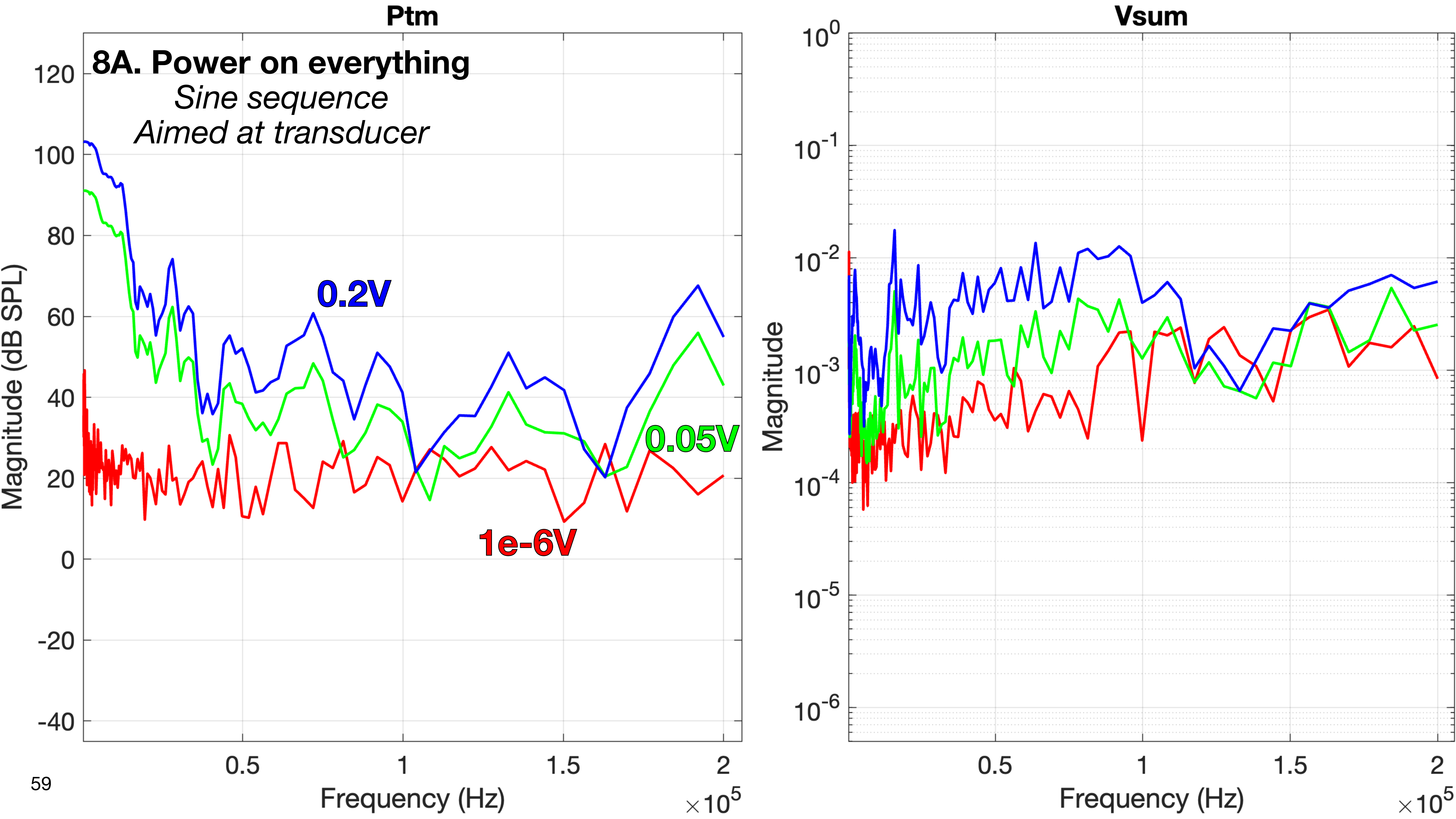

Ptm
8A. Power on everything
Sine sequence
Aimed at transducer
0.2V
0.05V
1e-6V
Magnitude (dB SPL)
Frequency (Hz)
$\times 10^5$
Vsum
Magnitude
Frequency (Hz)
$\times 10^5$

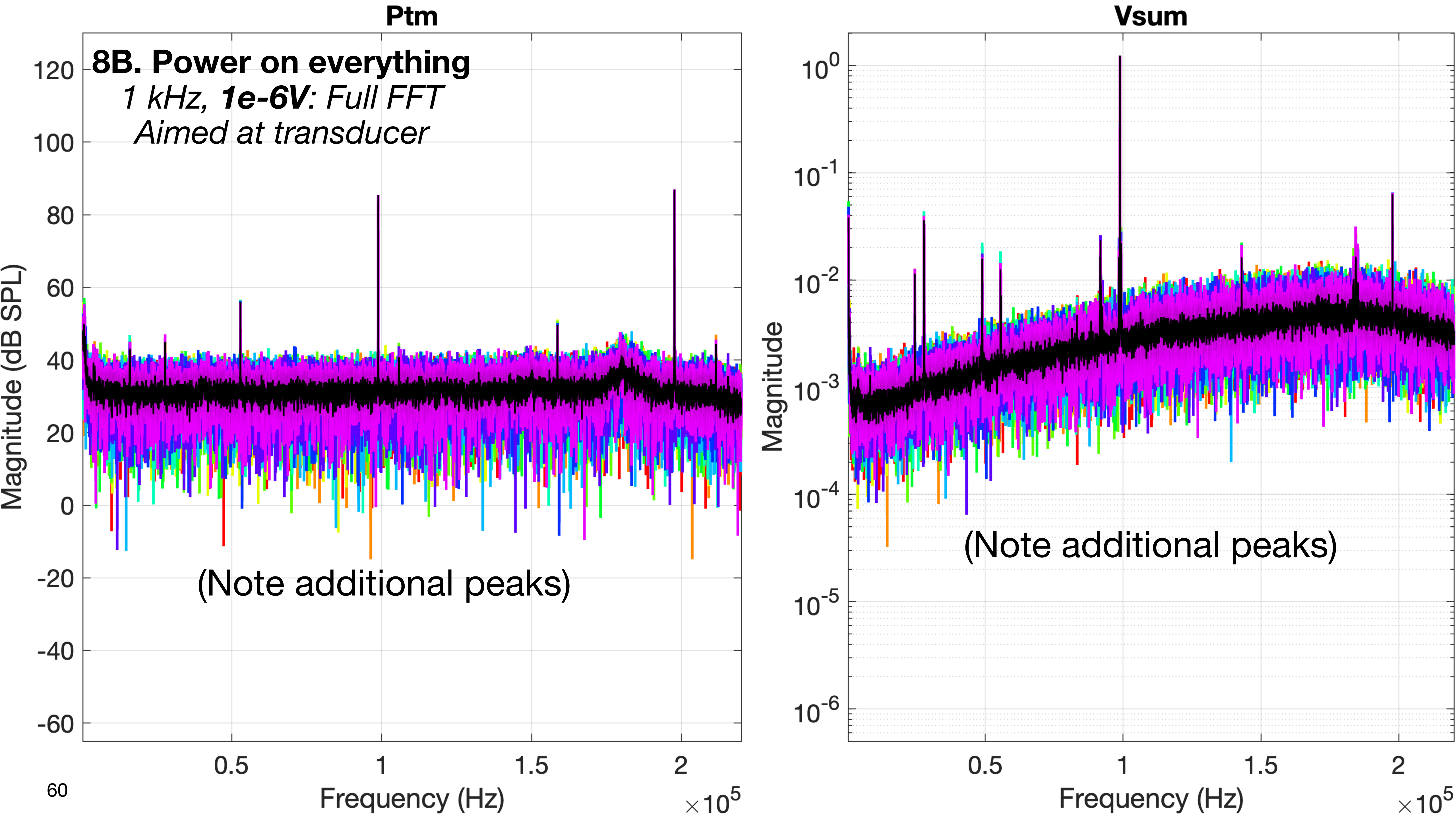
Ptm
8B. Power on everything
1 kHz, 1e-6V: Full FFT
Aimed at transducer
(Note additional peaks)
Magnitude (dB SPL)
120
100
80
60
40
20
0
-20
-40
-60
0.5
1
1.5
2
Frequency (Hz)
×10^5
Vsum
Magnitude
10^0
10^-1
10^-2
10^-3
10^-4
10^-5
10^-6
(Note additional peaks)
0.5
1
1.5
2
Frequency (Hz)
×10^5

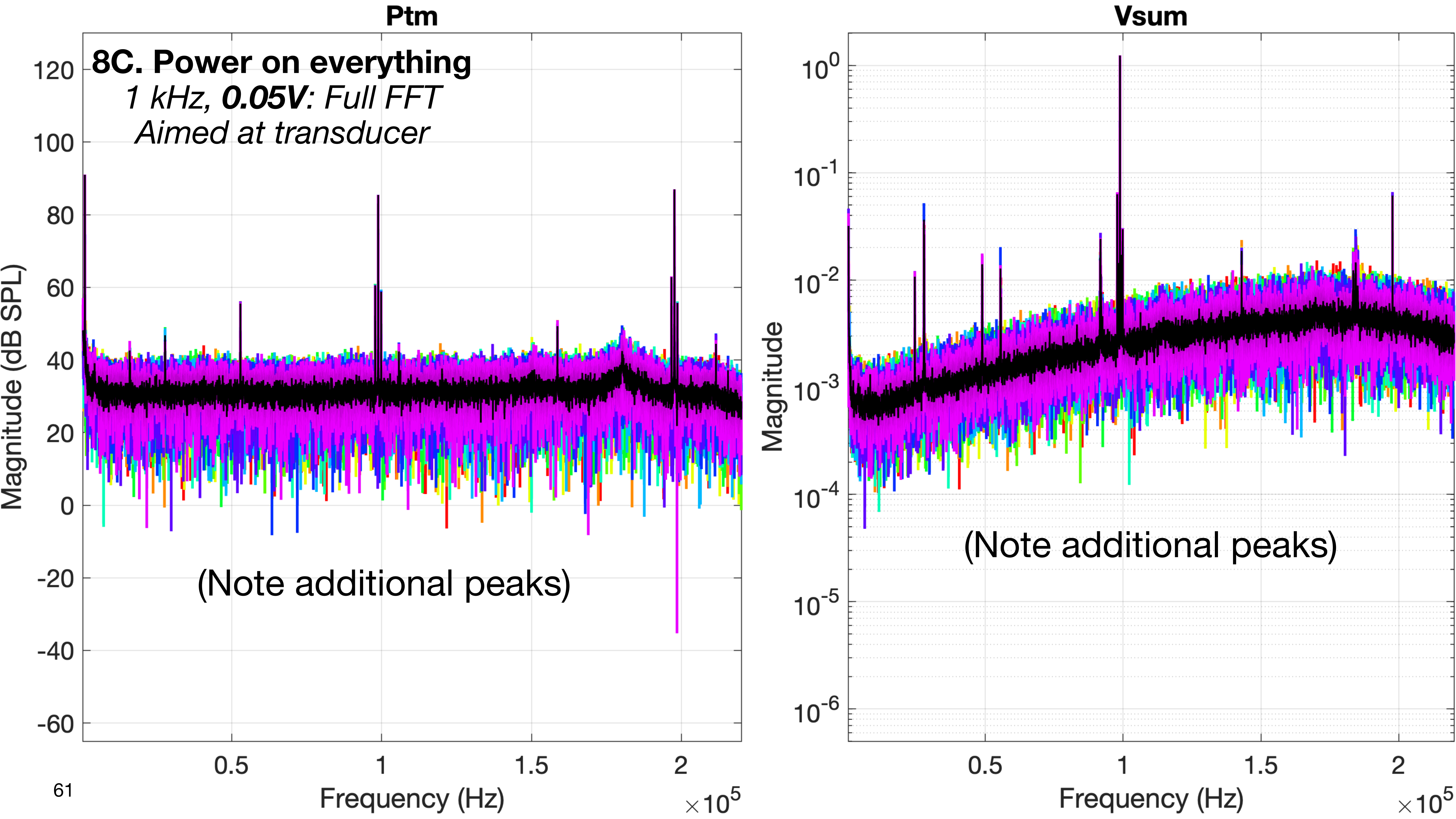
Ptm
8C. Power on everything
1 kHz, 0.05V: Full FFT
Aimed at transducer
(Note additional peaks)
Magnitude (dB SPL)
120
100
80
60
40
20
0
-20
-40
-60
0.5
1
1.5
2
Frequency (Hz)
$\times 10^5$
Vsum
Magnitude
$10^0$
$10^{-1}$
$10^{-2}$
$10^{-3}$
$10^{-4}$
$10^{-5}$
$10^{-6}$
(Note additional peaks)
0.5
1
1.5
2
Frequency (Hz)
$\times 10^5$

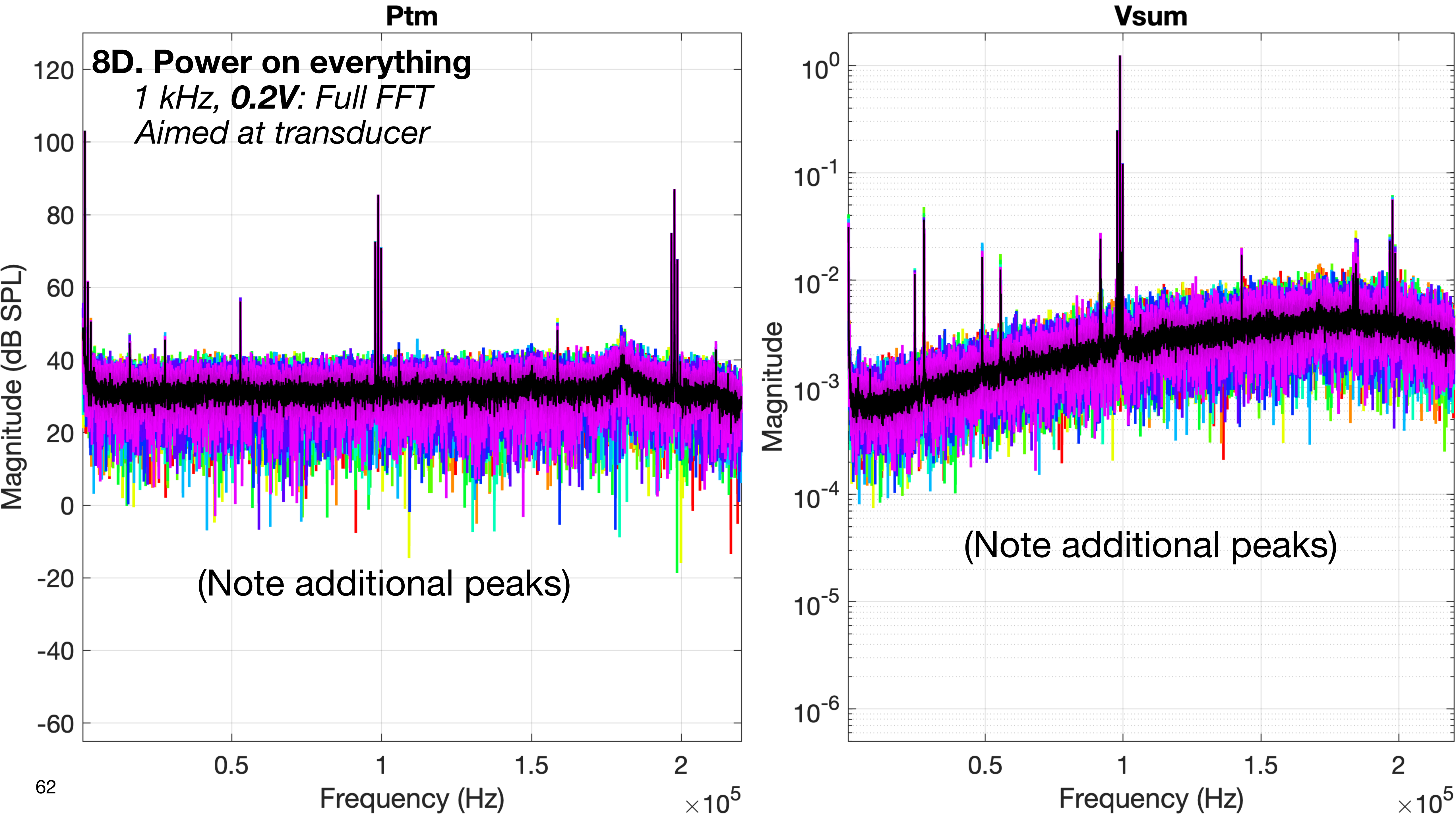
Ptm
8D. Power on everything
1 kHz, 0.2V: Full FFT
Aimed at transducer
Magnitude (dB SPL)
120
100
80
60
40
20
0
-20
-40
-60
(Note additional peaks)
0.5
1
1.5
2
Frequency (Hz)
×10^5
Vsum
Magnitude
10^0
10^-1
10^-2
10^-3
10^-4
10^-5
10^-6
(Note additional peaks)
0.5
1
1.5
2
Frequency (Hz)
×10^5

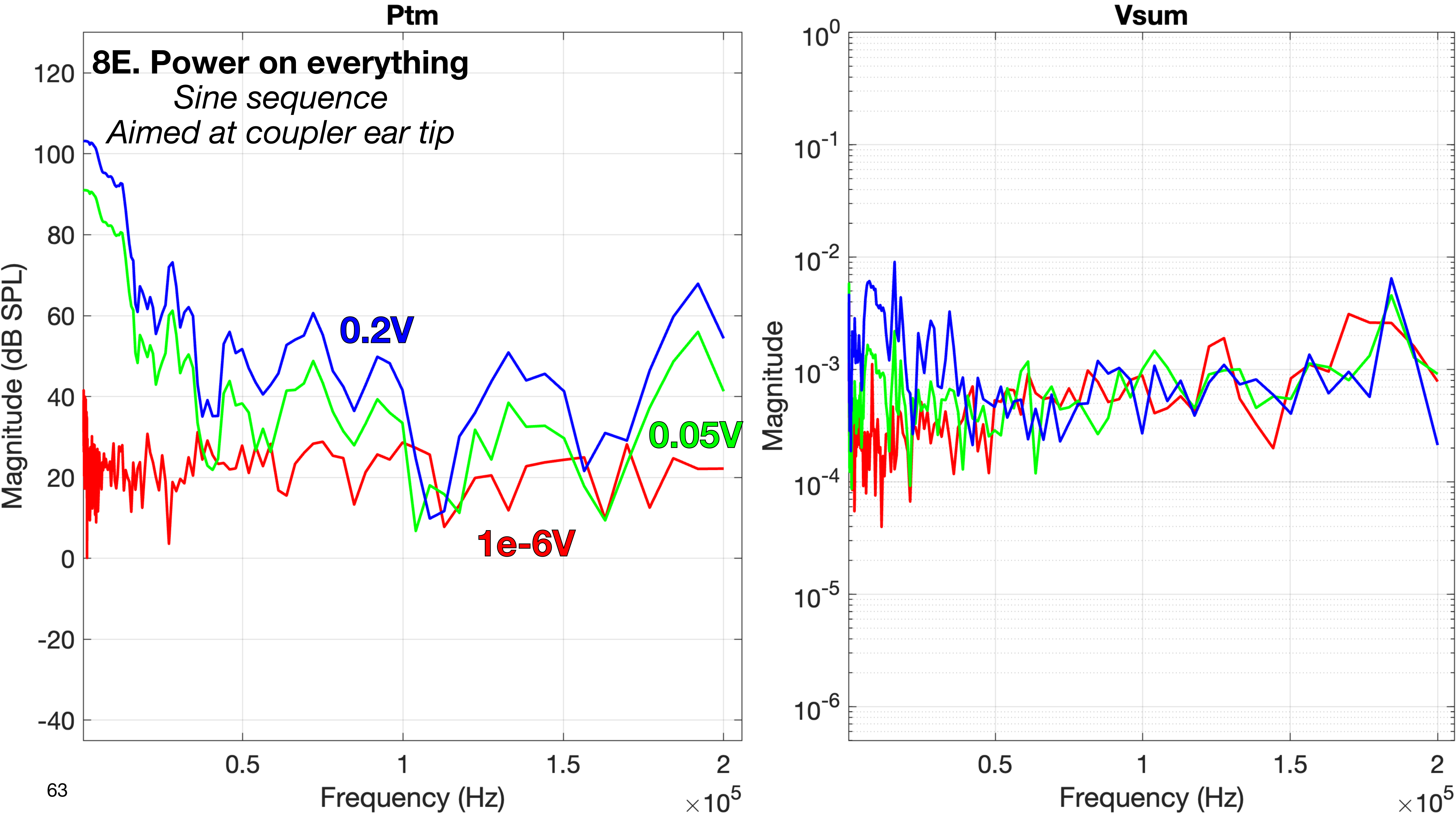

Ptm
8E. Power on everything
Sine sequence
Aimed at coupler ear tip
0.2V
0.05V
1e-6V
Magnitude (dB SPL)
120
100
80
60
40
20
0
-20
-40
0.5
1
1.5
2
Frequency (Hz)
$\times 10^5$
Vsum
Magnitude
$10^0$
$10^{-1}$
$10^{-2}$
$10^{-3}$
$10^{-4}$
$10^{-5}$
$10^{-6}$
0.5
1
1.5
2
Frequency (Hz)
$\times 10^5$

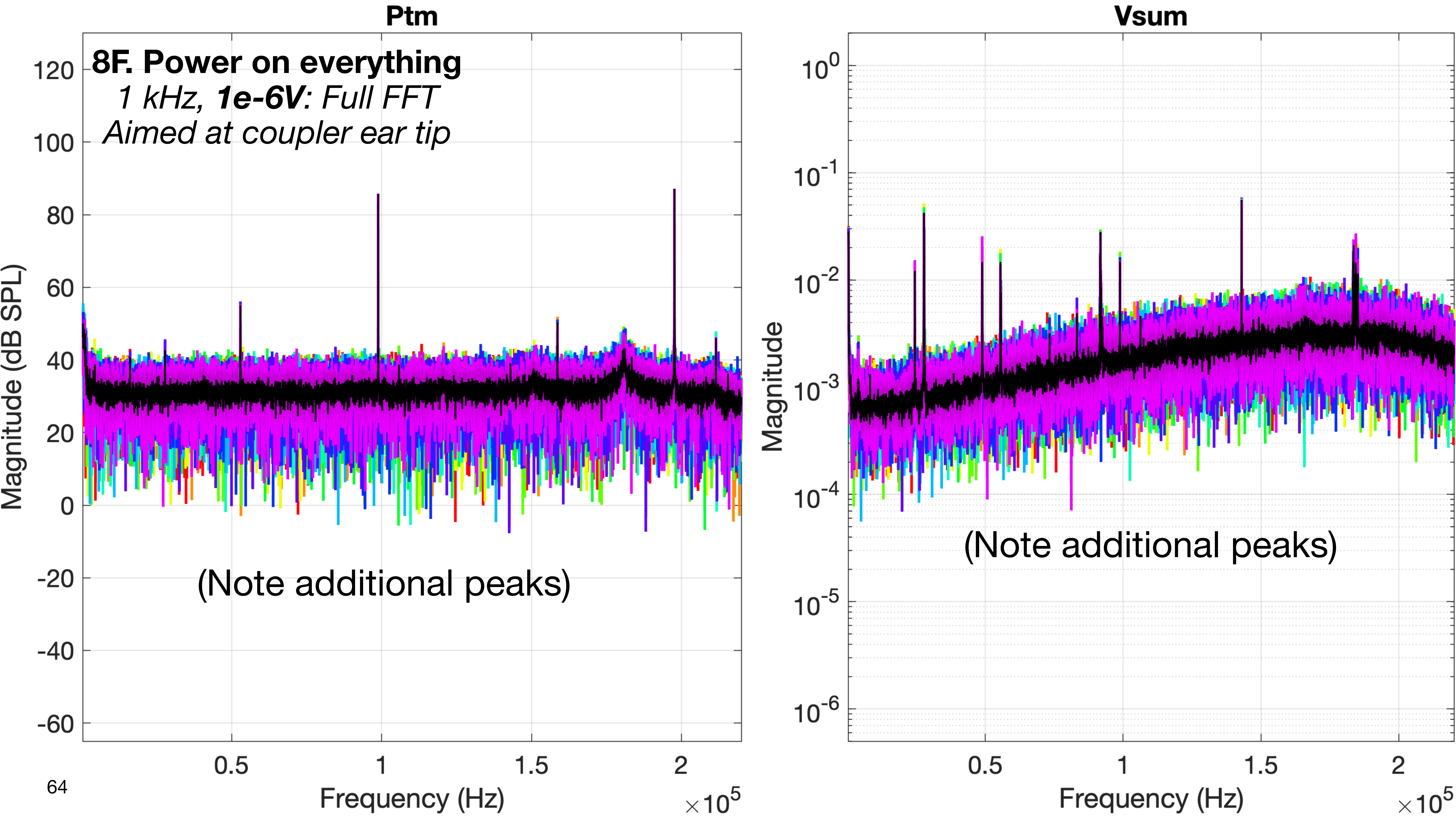
Ptm
8F. Power on everything
1 kHz, 1e-6V: Full FFT
Aimed at coupler ear tip
Magnitude (dB SPL)
120
100
80
60
40
20
0
-20
-40
-60
(Note additional peaks)
0.5
1
1.5
2
Frequency (Hz)
×10^5
Vsum
Magnitude
10^0
10^-1
10^-2
10^-3
10^-4
10^-5
10^-6
(Note additional peaks)
0.5
1
1.5
2
Frequency (Hz)
×10^5

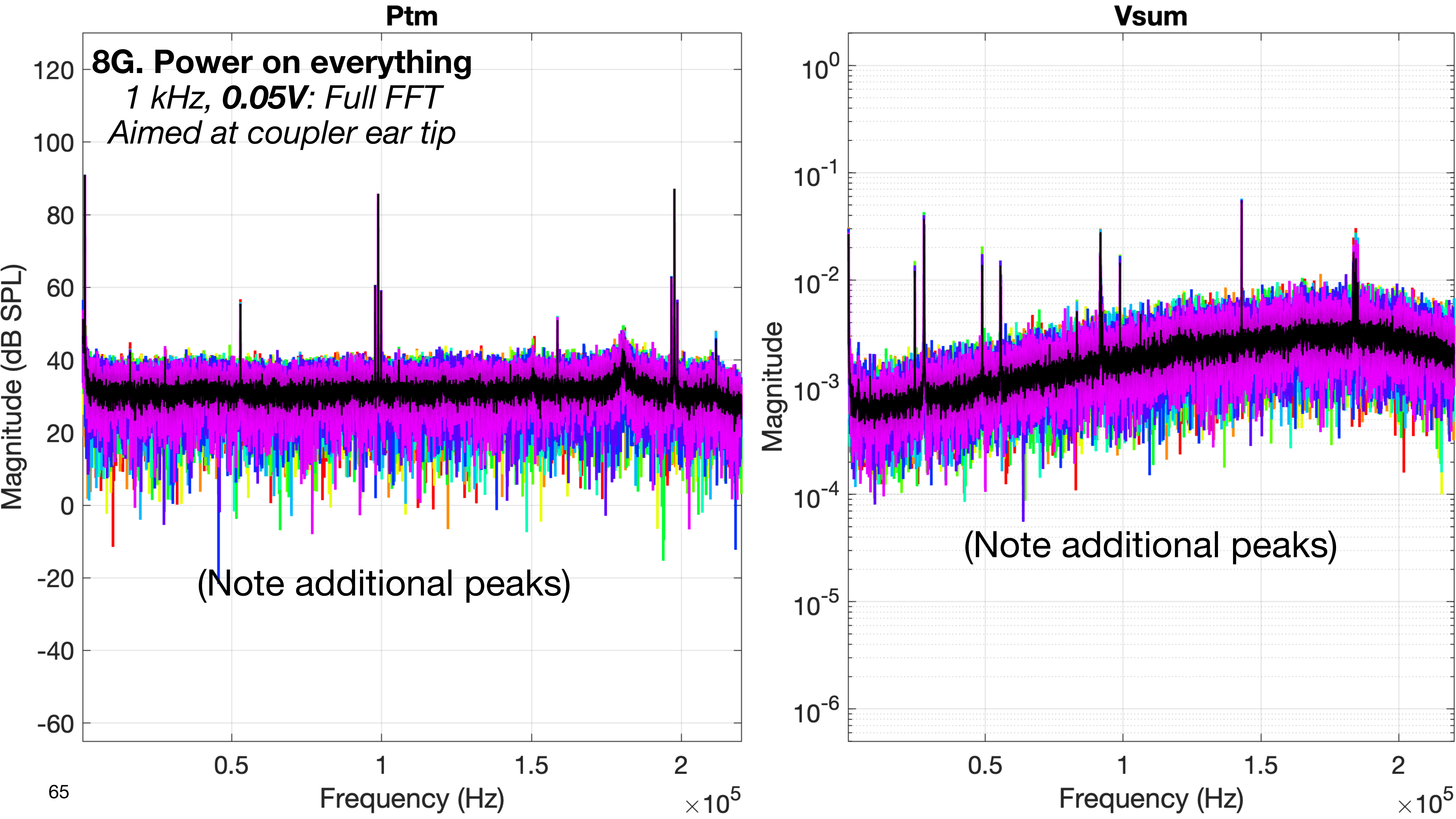

Ptm
8G. Power on everything
1 kHz, 0.05V: Full FFT
Aimed at coupler ear tip
(Note additional peaks)
Magnitude (dB SPL)
Frequency (Hz)
×10^5
Vsum
Magnitude
(Note additional peaks)
Frequency (Hz)
×10^5

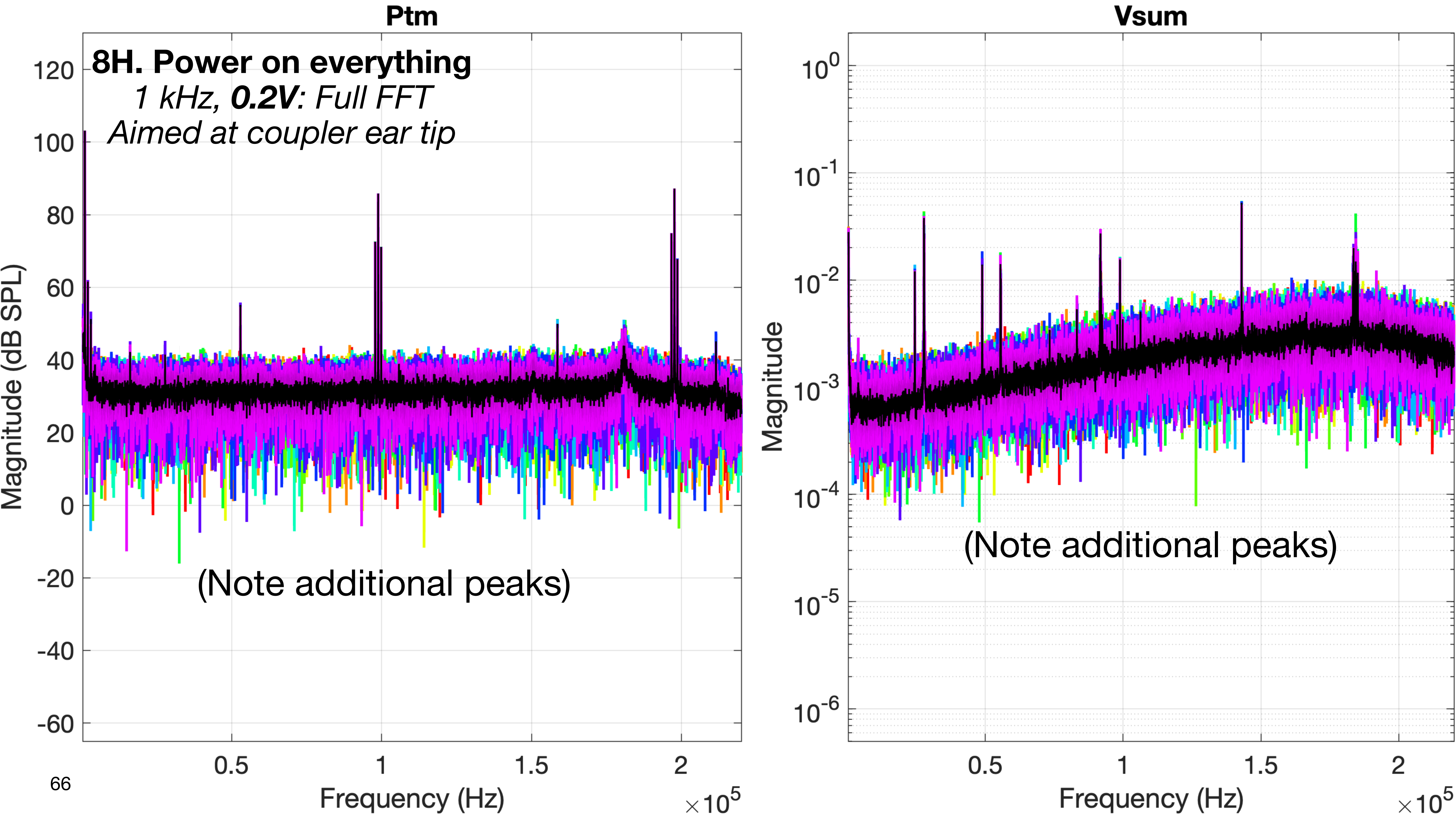
Ptm
8H. Power on everything
1 kHz, 0.2V: Full FFT
Aimed at coupler ear tip
(Note additional peaks)
Magnitude (dB SPL)
Frequency (Hz)
×10^5
120
100
80
60
40
20
0
-20
-40
-60
0.5
1
1.5
2
Vsum
Magnitude
10^0
10^-1
10^-2
10^-3
10^-4
10^-5
10^-6
(Note additional peaks)
Frequency (Hz)
×10^5

### 9. Power on everything EXCEPT the 3D carriage controller

- Wiring according to the normal schematic for measurements
- The 3384 LP filter is CONNECTED and powered ON
- The xMEMS microphone amp is CONNECTED and powered ON (the microphone is in the fixture)
- The 3D LDV is CONNECTED and powered ON, with the LDV aimed at the transducer
- The 3D carriage controller is powered OFF
- The xMEMS speaker driver is CONNECTED and powered ON

- **Purpose:**
  - This is to establish the baselines for the test fixture (with tape instead of a temporal bone) when all components of the measurement setup are connected and powered on normally.

### 9. Power on everything EXCEPT the 3D carriage controller

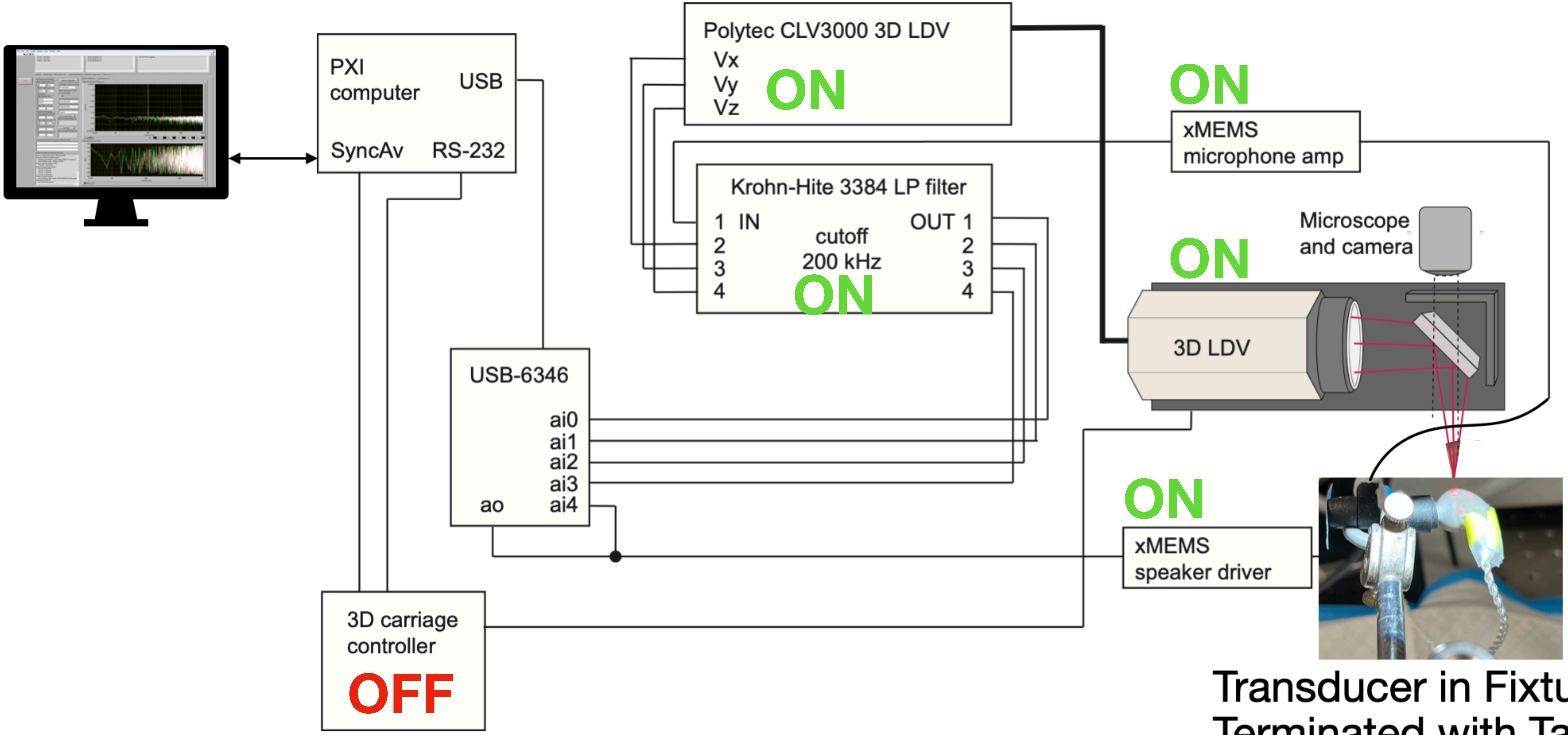


Transducer in Fixture, Terminated with Tape

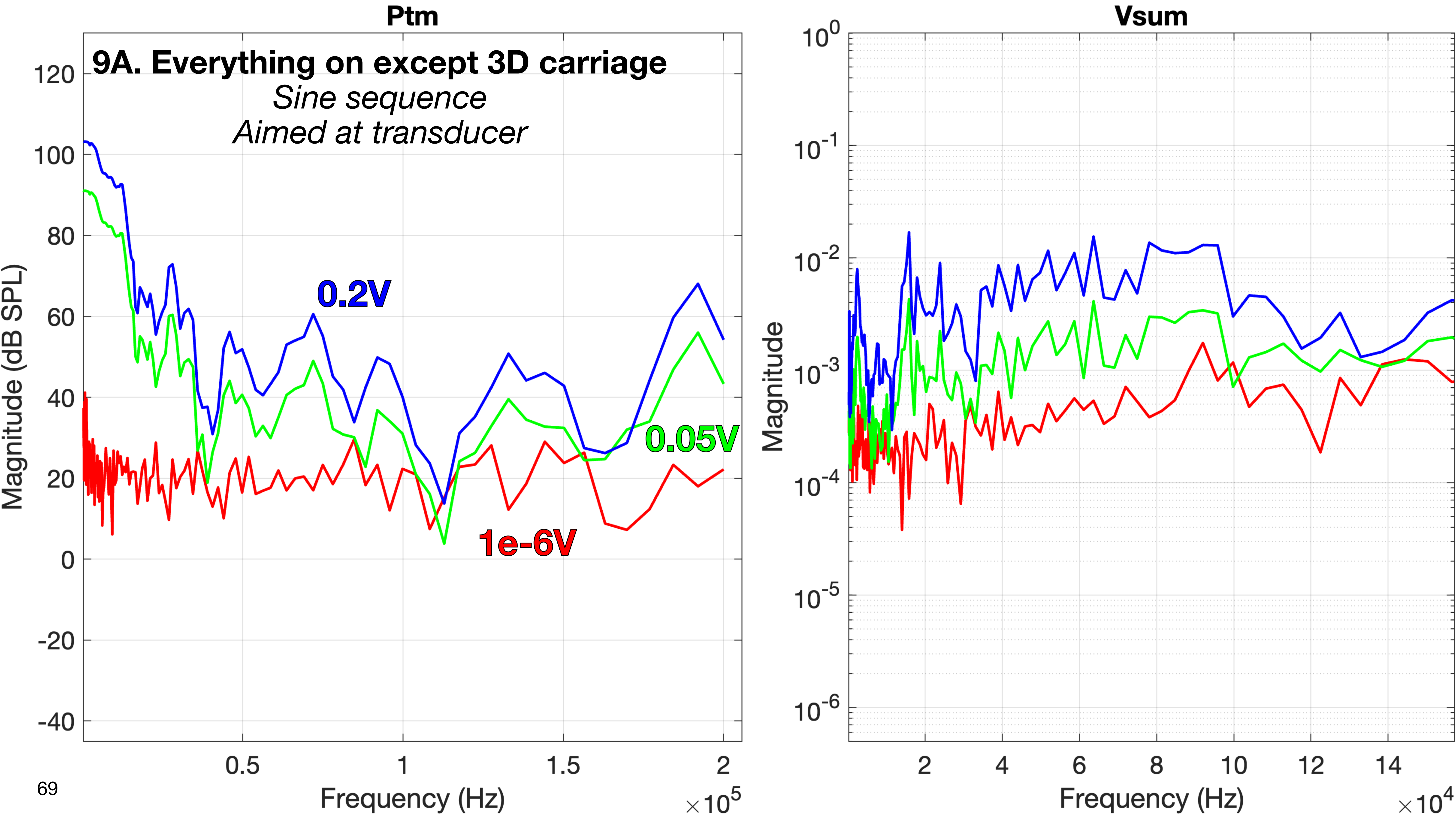

Ptm
9A. Everything on except 3D carriage
Sine sequence
Aimed at transducer
0.2V
0.05V
1e-6V
Magnitude (dB SPL)
120
100
80
60
40
20
0
-20
-40
0.5
1
1.5
2
Frequency (Hz)
×10^5
Vsum
Magnitude
10^0
10^-1
10^-2
10^-3
10^-4
10^-5
10^-6
2
4
6
8
10
12
14
Frequency (Hz)
×10^4

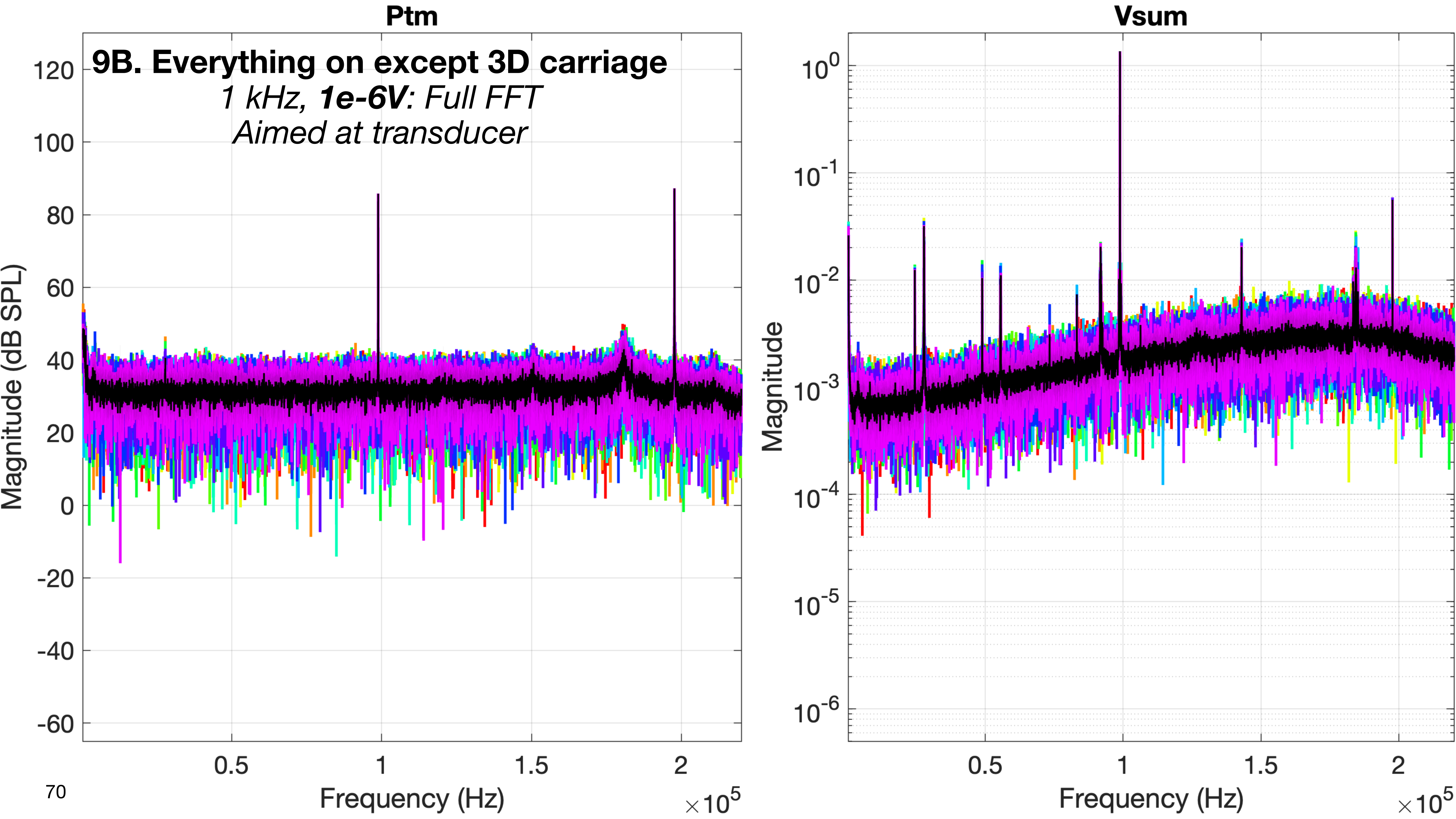
Ptm
9B. Everything on except 3D carriage
1 kHz, 1e-6V: Full FFT
Aimed at transducer
Magnitude (dB SPL)
120
100
80
60
40
20
0
-20
-40
-60
0.5
1
1.5
2
Frequency (Hz)
×10^5
Vsum
Magnitude
10^0
10^-1
10^-2
10^-3
10^-4
10^-5
10^-6
0.5
1
1.5
2
Frequency (Hz)
×10^5

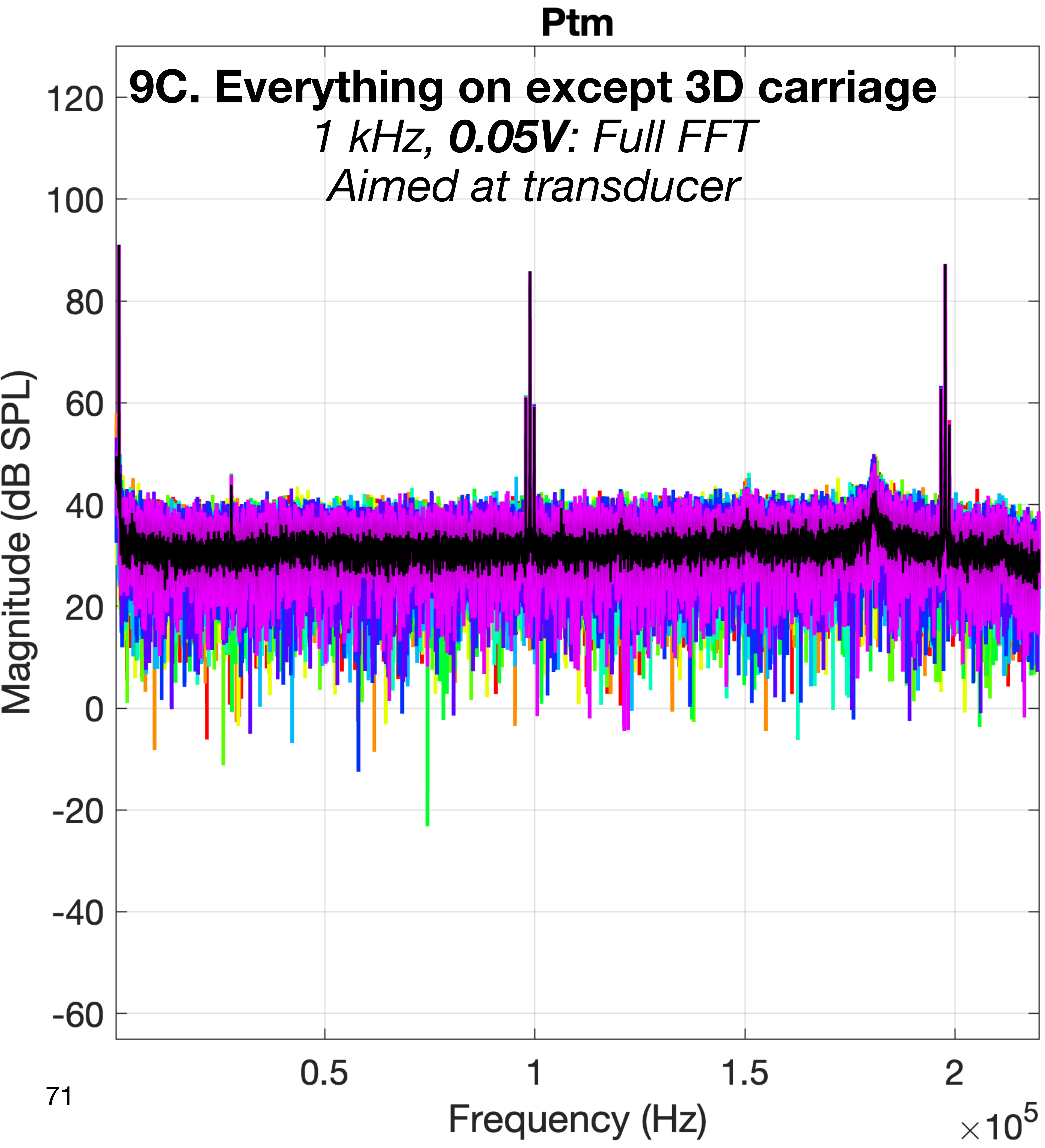
Ptm
9C. Everything on except 3D carriage
1 kHz, 0.05V: Full FFT
Aimed at transducer
Magnitude (dB SPL)
120
100
80
60
40
20
0
-20
-40
-60
0.5
1
1.5
2
Frequency (Hz)
×10^5

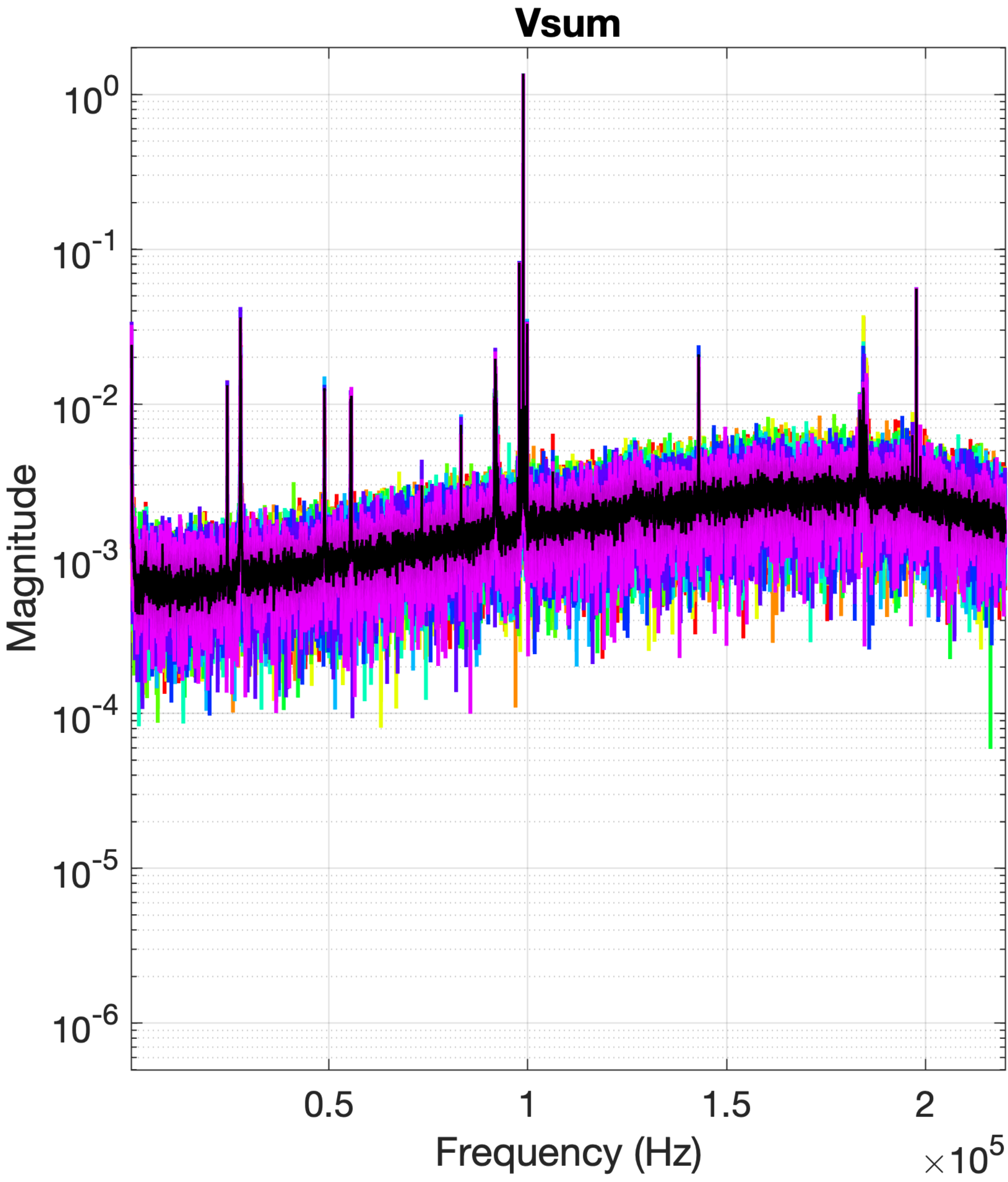
Vsum
Magnitude
10^0
10^-1
10^-2
10^-3
10^-4
10^-5
10^-6
0.5
1
1.5
2
Frequency (Hz)
×10^5

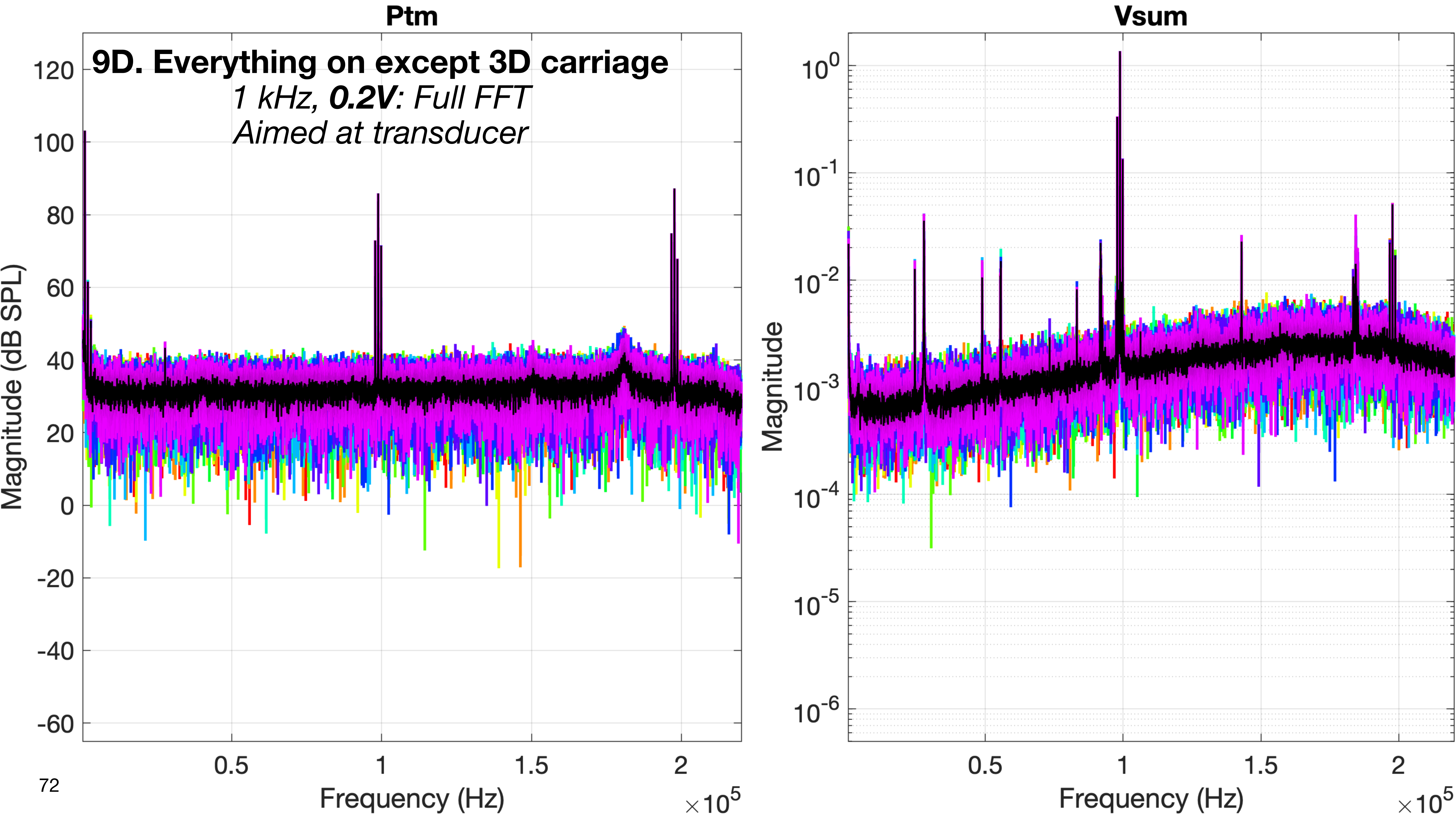
Ptm
9D. Everything on except 3D carriage
1 kHz, 0.2V: Full FFT
Aimed at transducer
Magnitude (dB SPL)
120
100
80
60
40
20
0
-20
-40
-60
0.5
1
1.5
2
Frequency (Hz)
×10^5
Vsum
Magnitude
10^0
10^-1
10^-2
10^-3
10^-4
10^-5
10^-6
0.5
1
1.5
2
Frequency (Hz)
×10^5

### Identifying Peaks (Spikes)

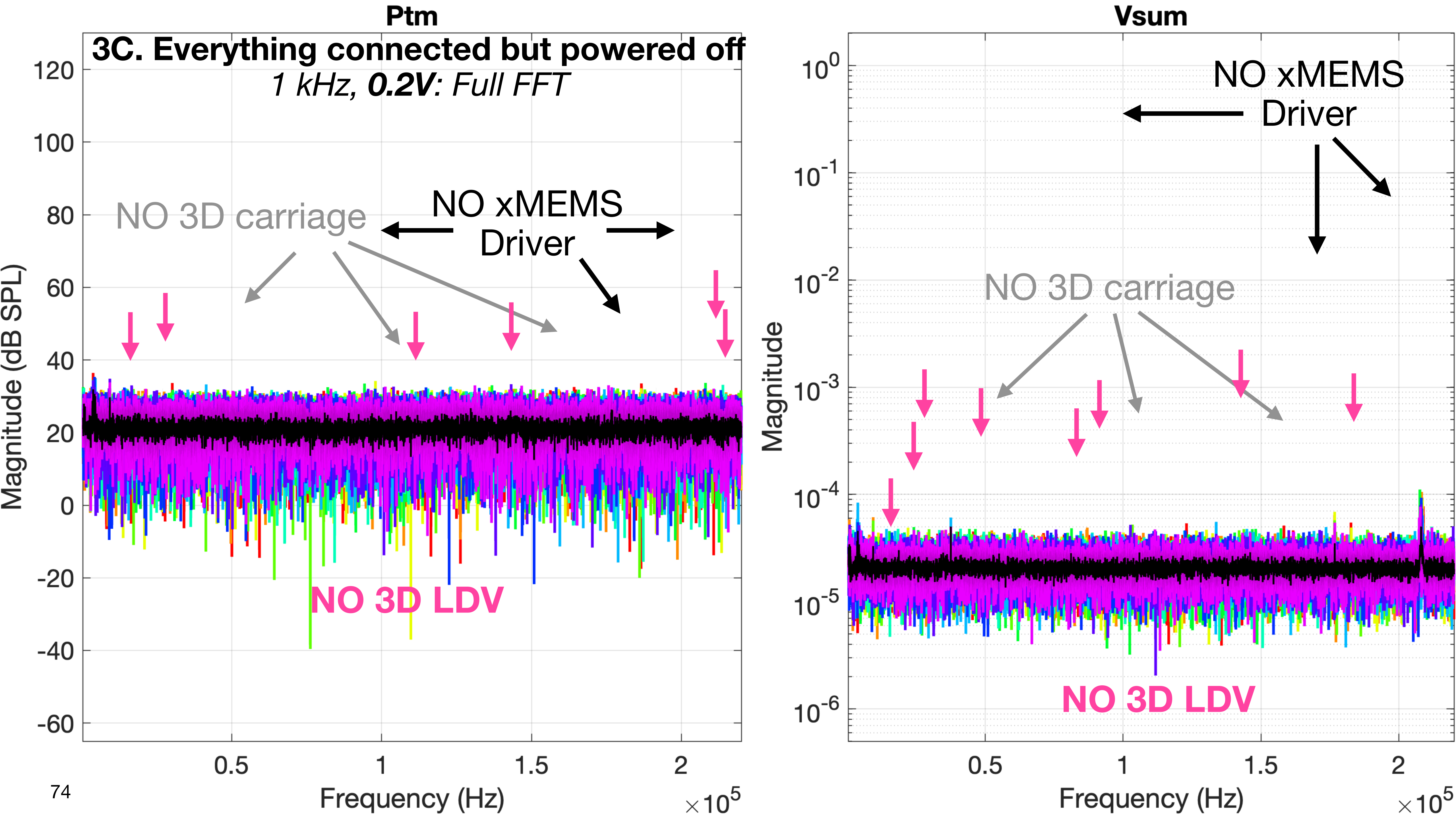
Ptm
3C. Everything connected but powered off
1 kHz, 0.2V: Full FFT
NO 3D carriage
NO xMEMS
Driver
NO 3D LDV
Magnitude (dB SPL)
120
100
80
60
40
20
0
-20
-40
-60
0.5
1
1.5
2
Frequency (Hz)
×10^5
Vsum
NO xMEMS
Driver
NO 3D carriage
NO 3D LDV
Magnitude
10^0
10^-1
10^-2
10^-3
10^-4
10^-5
10^-6
0.5
1
1.5
2
Frequency (Hz)
×10^5

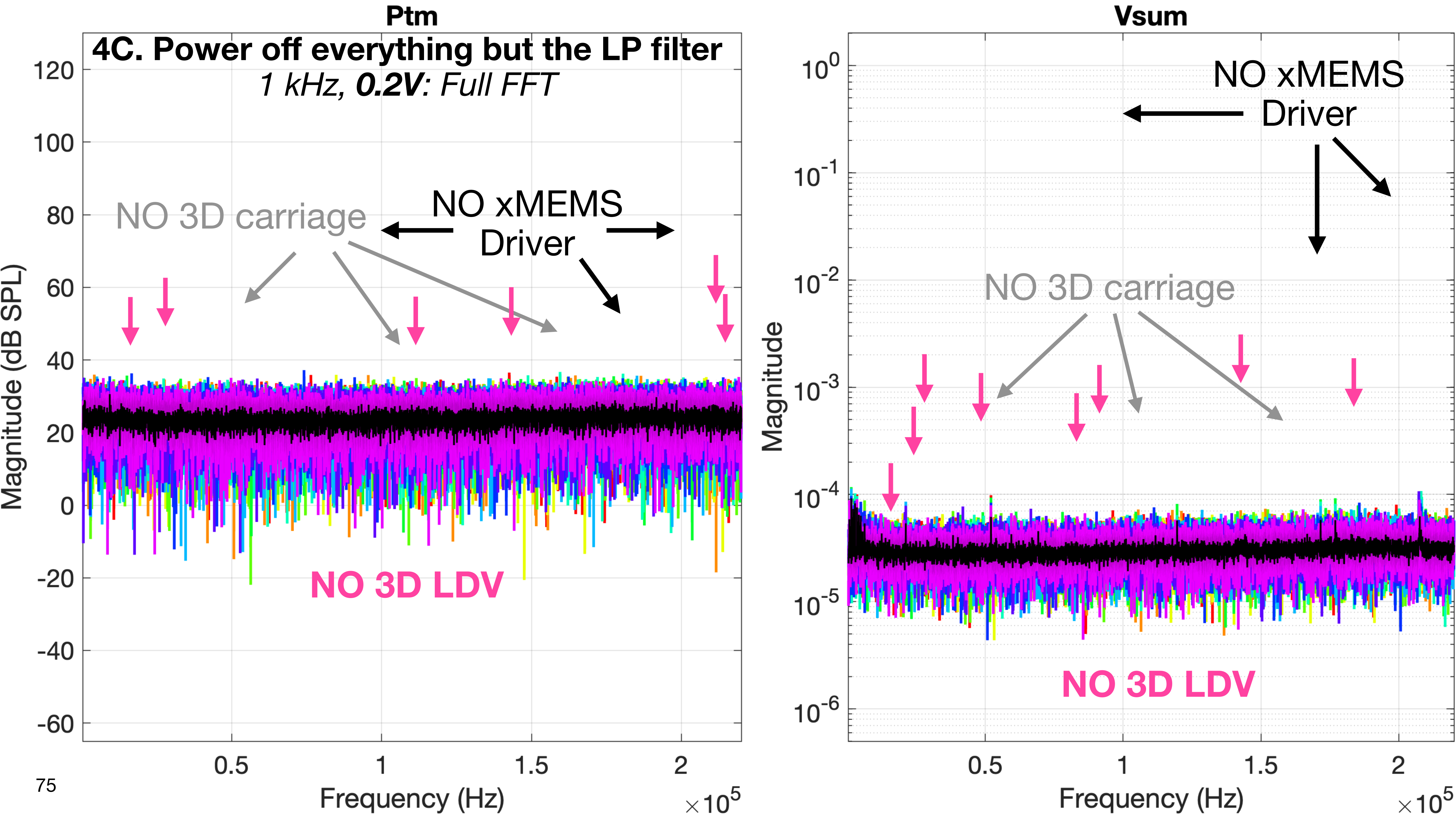
Ptm
4C. Power off everything but the LP filter
1 kHz, 0.2V: Full FFT
NO 3D carriage
NO xMEMS Driver
NO 3D LDV
Magnitude (dB SPL)
Frequency (Hz)
×10^5
Vsum
NO xMEMS Driver
NO 3D carriage
NO 3D LDV
Magnitude
Frequency (Hz)
×10^5

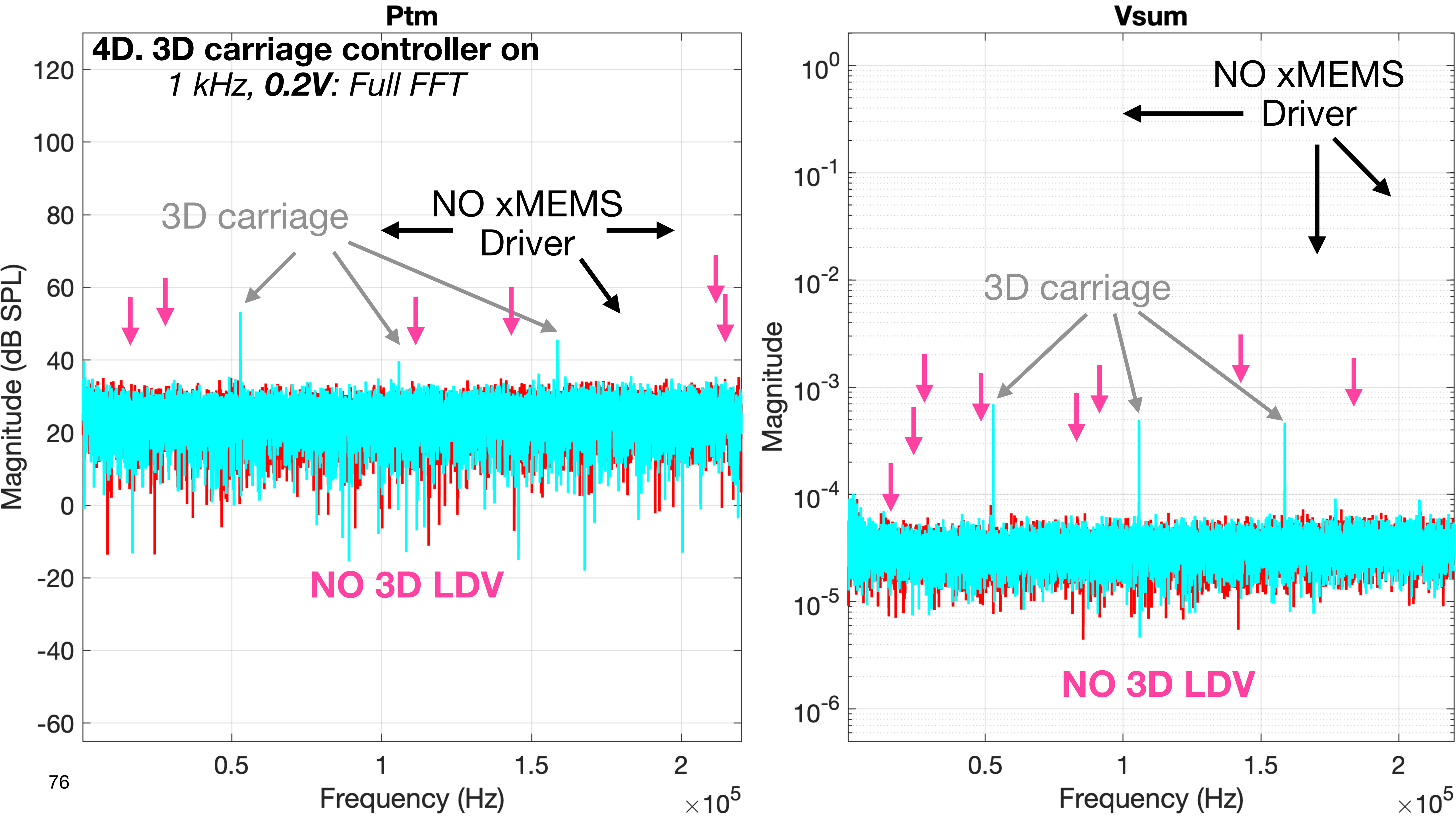

Ptm
4D. 3D carriage controller on
*1 kHz, **0.2V**: Full FFT*
3D carriage
NO xMEMS
Driver
NO 3D LDV
Magnitude (dB SPL)
Frequency (Hz)
×10^5
Vsum
NO xMEMS
Driver
3D carriage
NO 3D LDV
Magnitude
Frequency (Hz)
×10^5

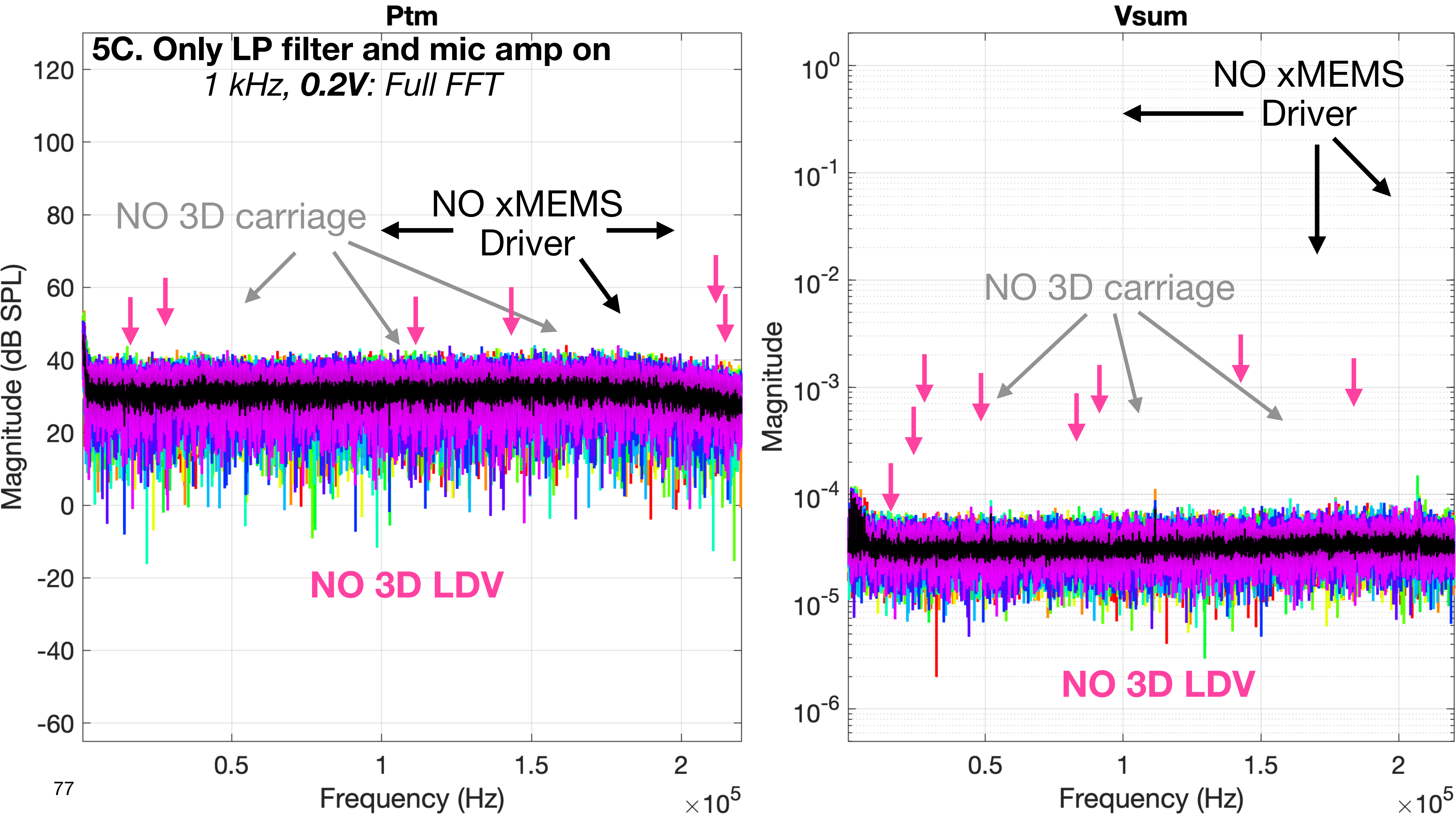
Ptm
5C. Only LP filter and mic amp on
1 kHz, 0.2V: Full FFT
NO 3D carriage
NO xMEMS Driver
NO 3D LDV
Magnitude (dB SPL)
Frequency (Hz)
Vsum
NO xMEMS Driver
NO 3D carriage
NO 3D LDV
Magnitude
Frequency (Hz)

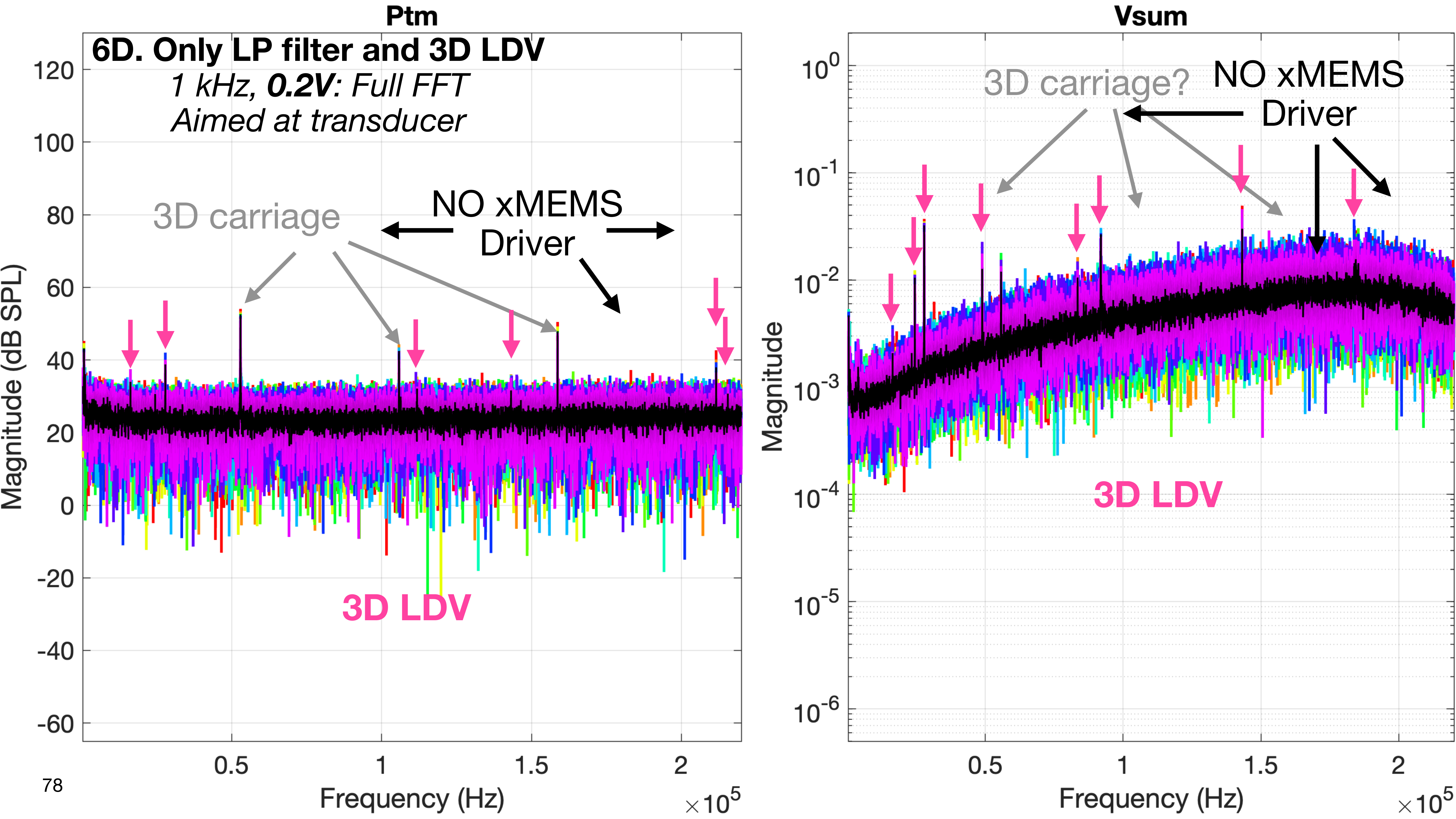
Ptm
6D. Only LP filter and 3D LDV
1 kHz, 0.2V: Full FFT
Aimed at transducer
3D carriage
NO xMEMS
Driver
3D LDV
Magnitude (dB SPL)
Frequency (Hz)
×10^5
Vsum
3D carriage?
NO xMEMS
Driver
3D LDV
Magnitude
Frequency (Hz)
×10^5

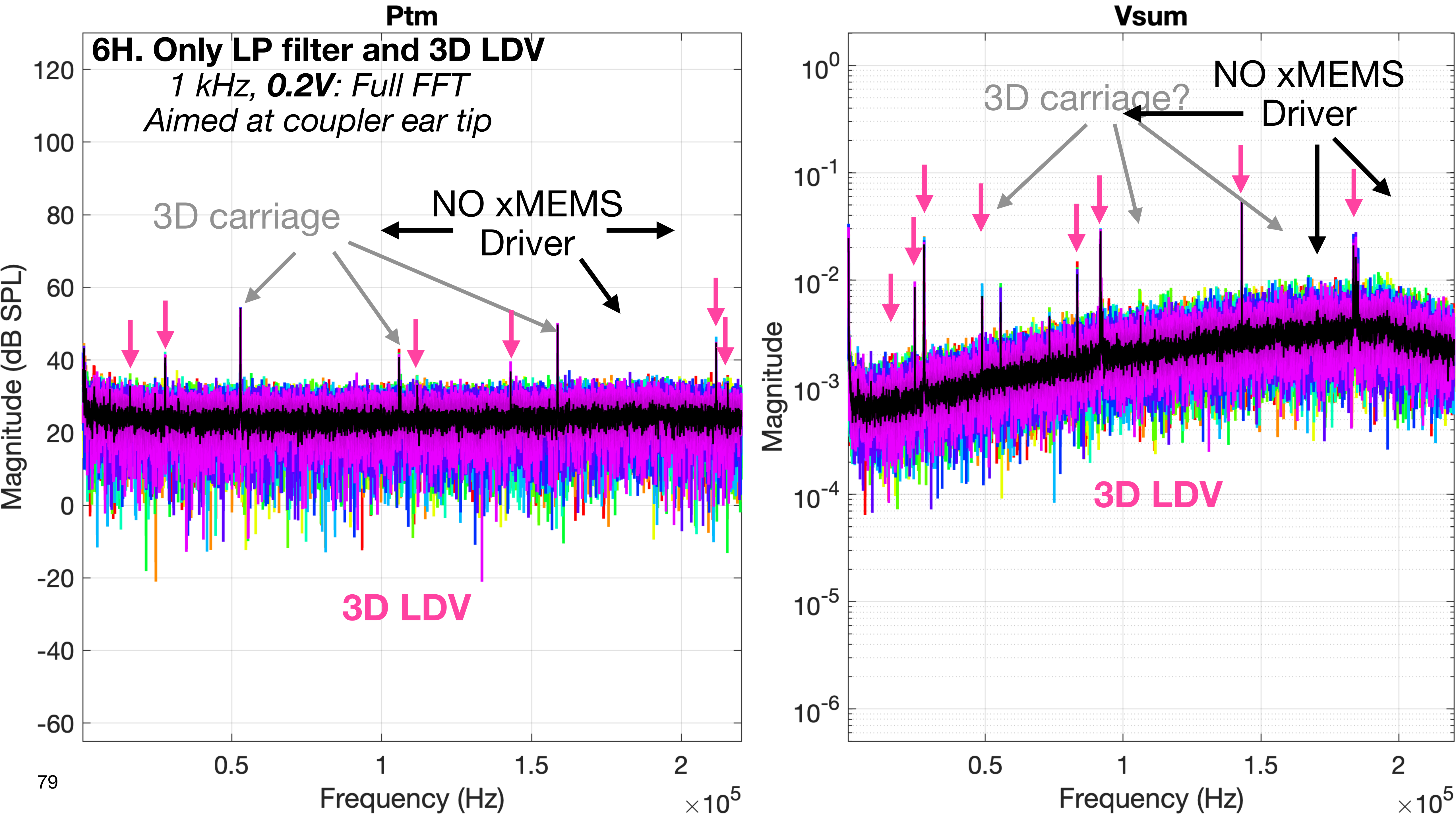
Ptm
6H. Only LP filter and 3D LDV
1 kHz, 0.2V: Full FFT
Aimed at coupler ear tip
3D carriage
NO xMEMS
Driver
3D LDV
Magnitude (dB SPL)
Frequency (Hz)
×10^5
Vsum
3D carriage?
NO xMEMS
Driver
3D LDV
Magnitude
Frequency (Hz)
×10^5

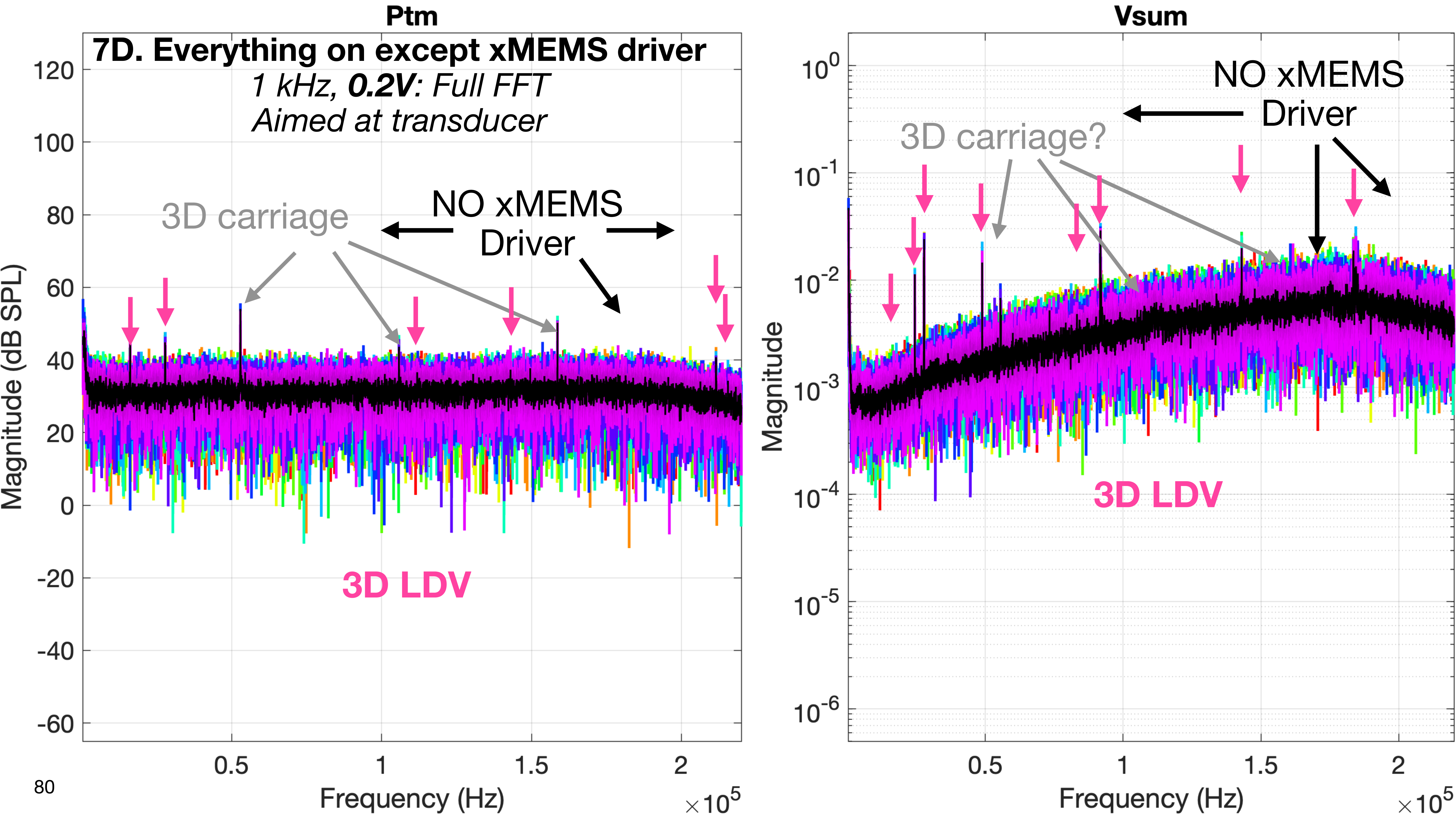

Ptm
7D. Everything on except xMEMS driver
1 kHz, 0.2V: Full FFT
Aimed at transducer
3D carriage
NO xMEMS
Driver
3D LDV
Magnitude (dB SPL)
Frequency (Hz)
×10^5
Vsum
NO xMEMS
Driver
3D carriage?
3D LDV
Magnitude
Frequency (Hz)
×10^5

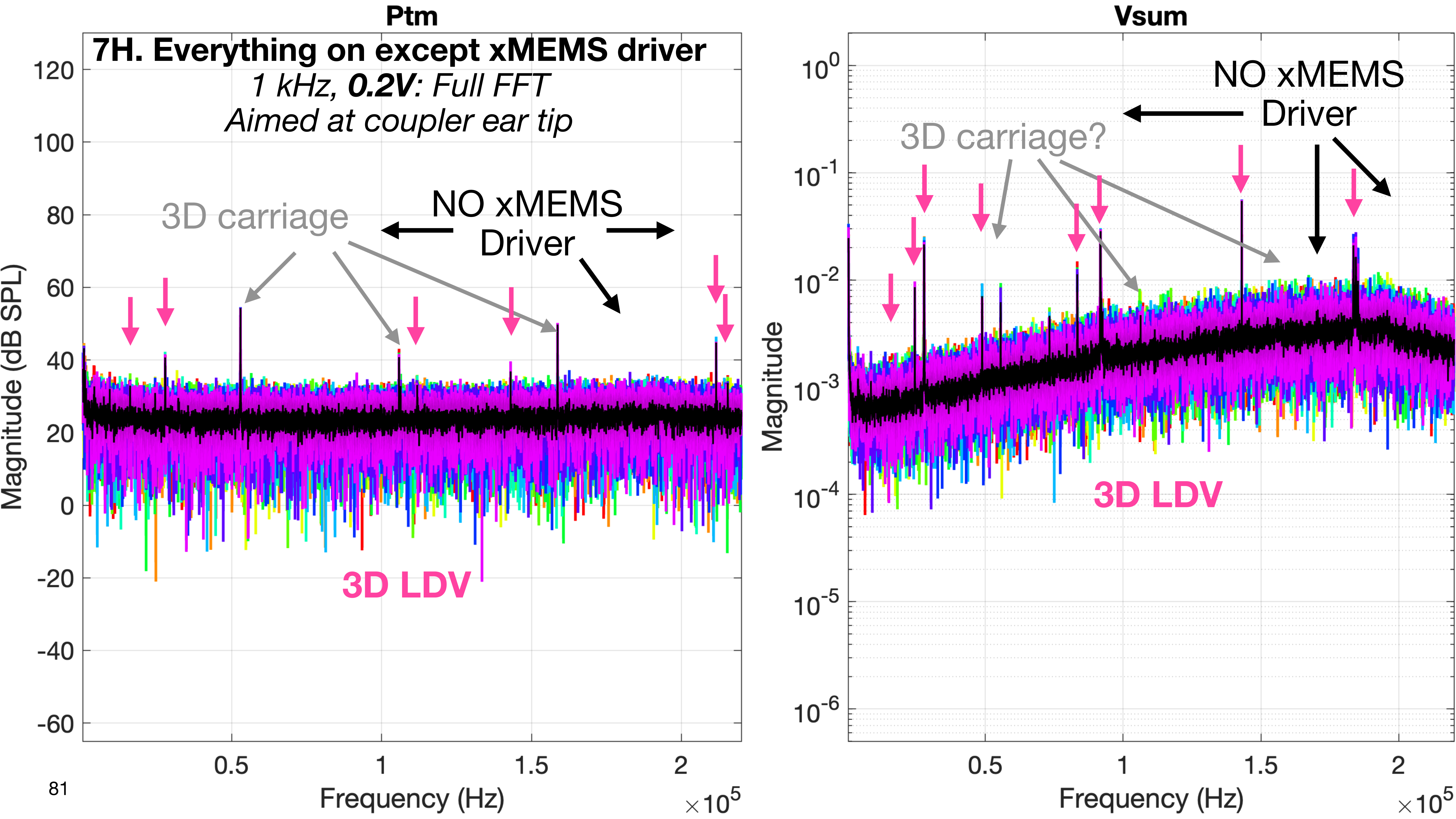

Ptm
7H. Everything on except xMEMS driver
1 kHz, 0.2V: Full FFT
Aimed at coupler ear tip
3D carriage
NO xMEMS
Driver
3D LDV
Magnitude (dB SPL)
Frequency (Hz)
×10^5
Vsum
NO xMEMS
Driver
3D carriage?
3D LDV
Magnitude
Frequency (Hz)
×10^5

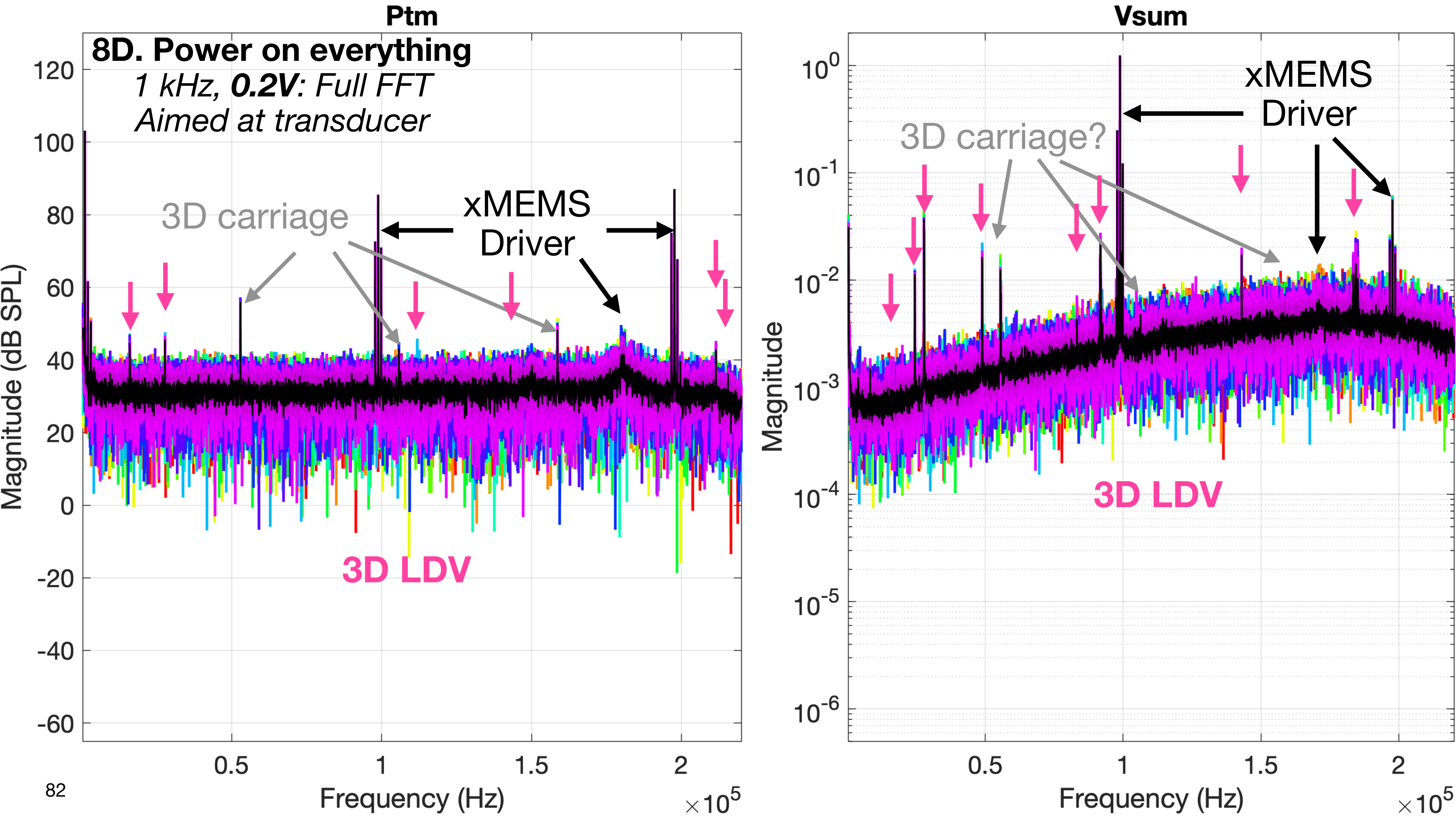
Ptm
8D. Power on everything
1 kHz, 0.2V: Full FFT
Aimed at transducer
3D carriage
xMEMS
Driver
3D LDV
Magnitude (dB SPL)
Frequency (Hz)
×10^5
Vsum
xMEMS
Driver
3D carriage?
3D LDV
Magnitude
Frequency (Hz)
×10^5

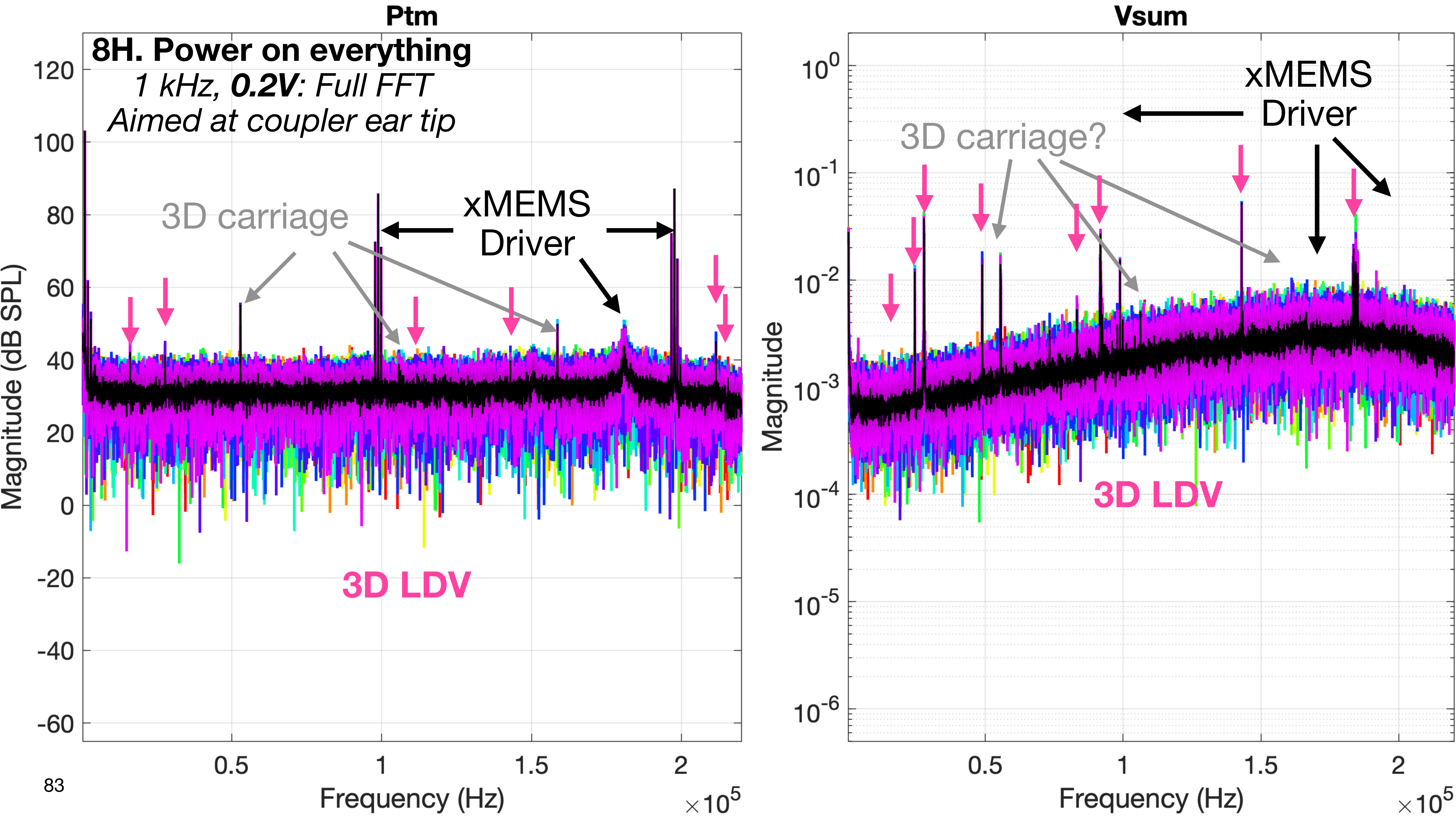

Ptm
8H. Power on everything
*1 kHz, **0.2V**: Full FFT*
*Aimed at coupler ear tip*
3D carriage
xMEMS
Driver
3D LDV
Magnitude (dB SPL)
Frequency (Hz)
×10^5
Vsum
xMEMS
Driver
3D carriage?
3D LDV
Magnitude
Frequency (Hz)
×10^5

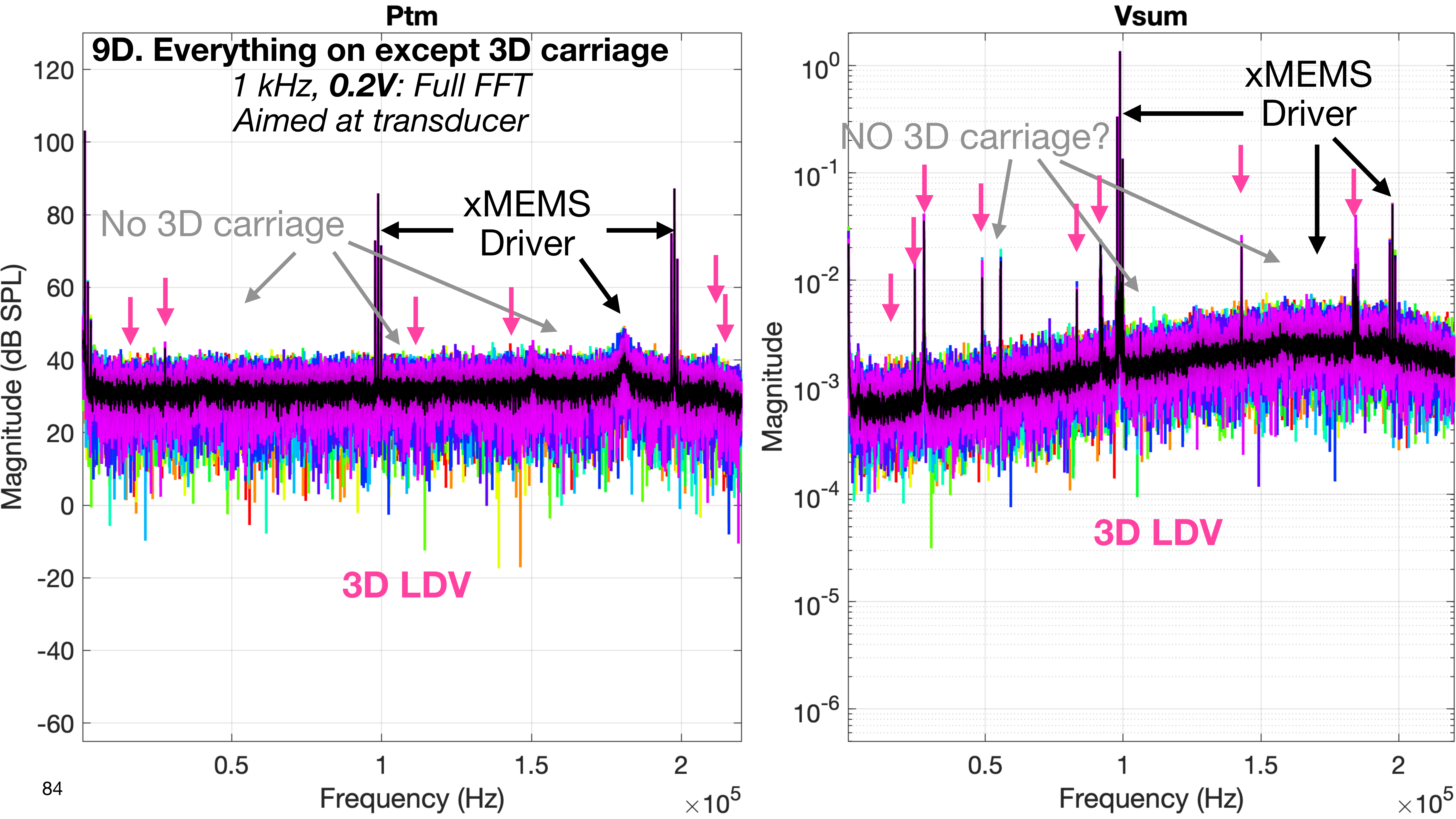
Ptm
9D. Everything on except 3D carriage
1 kHz, 0.2V: Full FFT
Aimed at transducer
No 3D carriage
xMEMS
Driver
3D LDV
Magnitude (dB SPL)
Frequency (Hz)
×10^5
Vsum
NO 3D carriage?
xMEMS
Driver
3D LDV
Magnitude
Frequency (Hz)
×10^5

### Measurements in temporal bone ears (TB7,TB8,TB9)

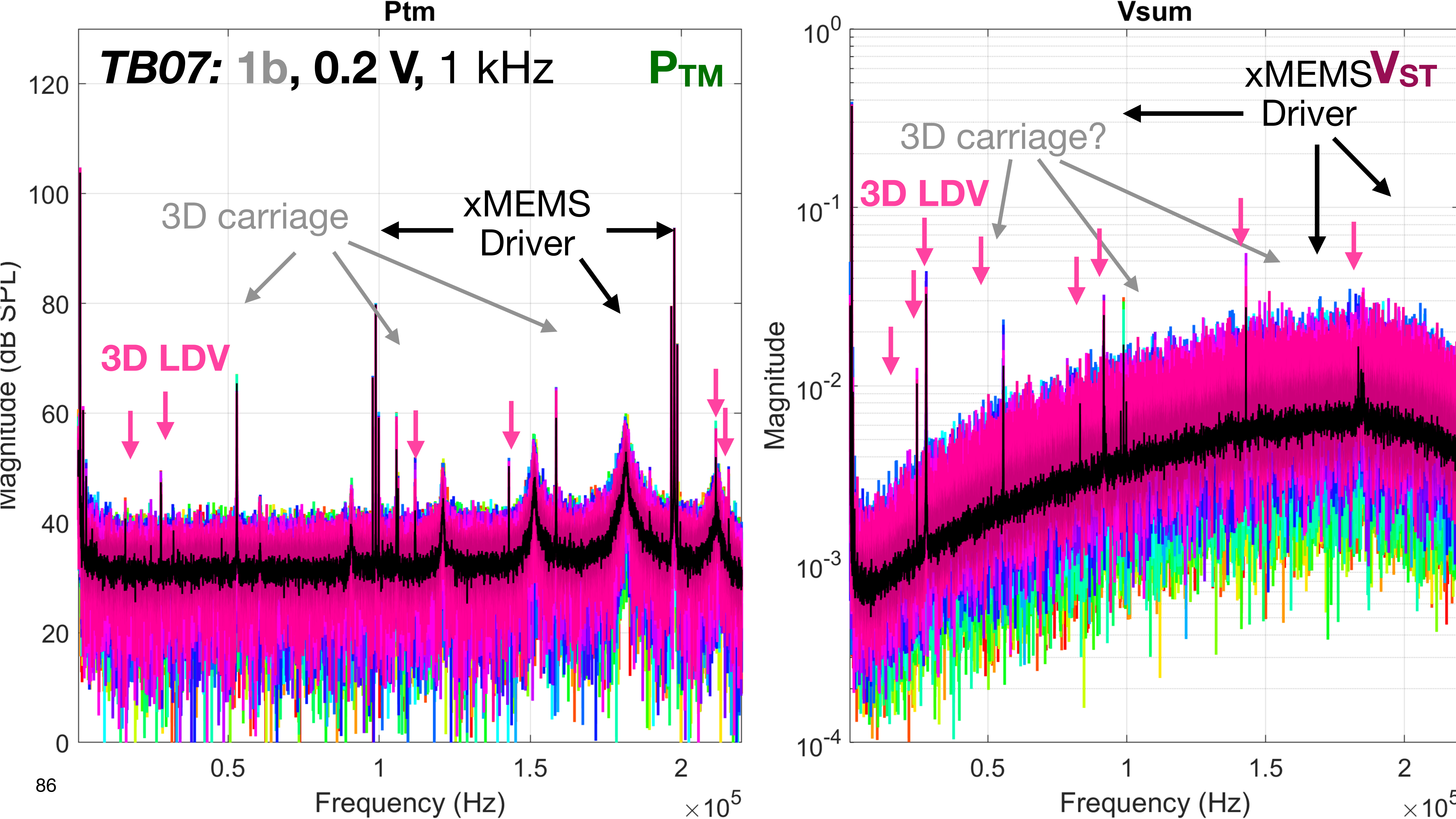
Ptm
TB07: 1b, 0.2 V, 1 kHz
PTM
3D carriage
xMEMS
Driver
3D LDV
Magnitude (dB SPL)
Frequency (Hz)
×10^5
Vsum
xMEMS VST
Driver
3D carriage?
3D LDV
Magnitude
Frequency (Hz)
×10^5

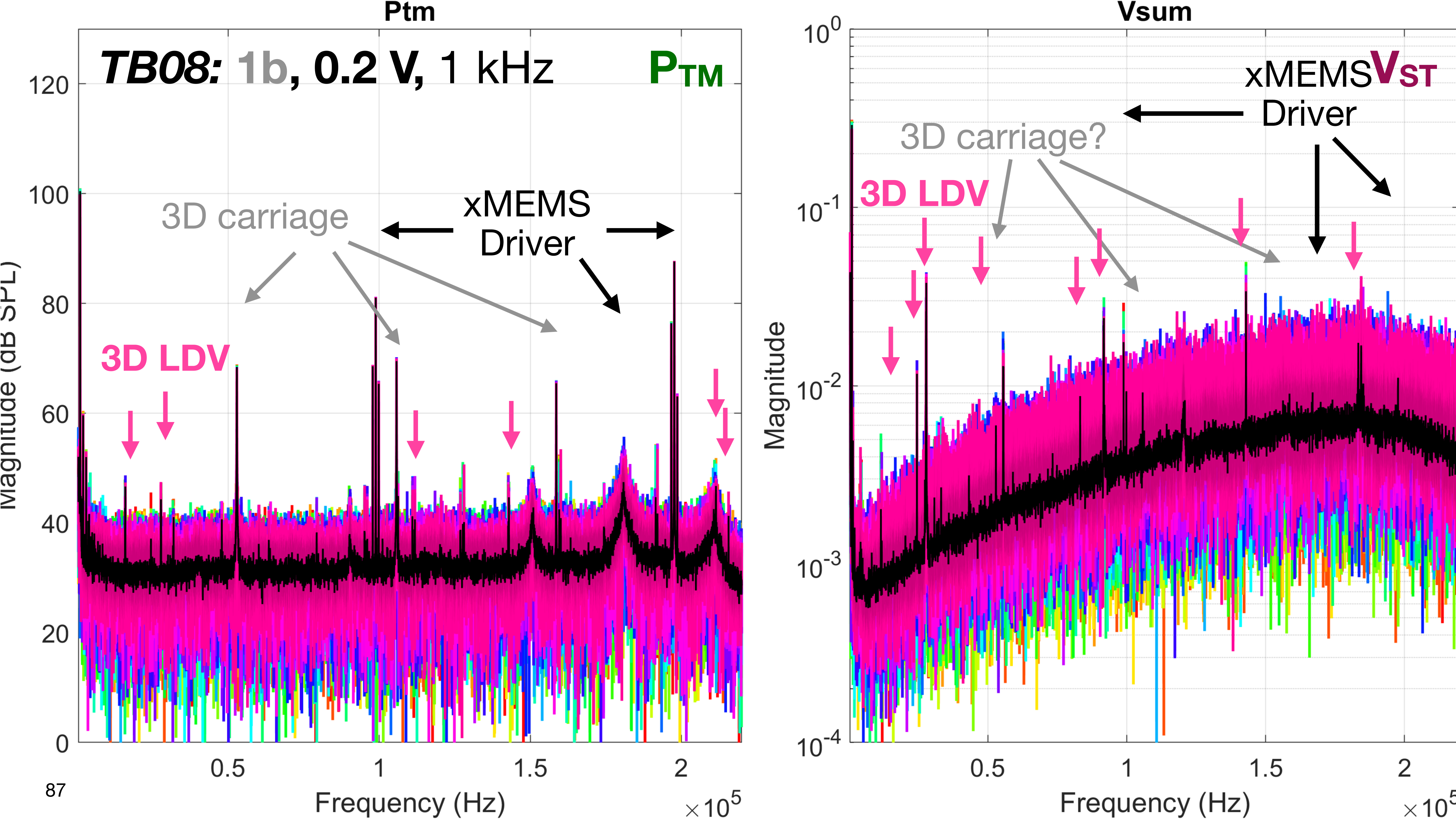

Ptm
TB08: 1b, 0.2 V, 1 kHz
PTM
3D carriage
xMEMS
Driver
3D LDV
Magnitude (dB SPL)
Frequency (Hz)
×10^5
Vsum
xMEMS VST
Driver
3D carriage?
3D LDV
Magnitude
Frequency (Hz)

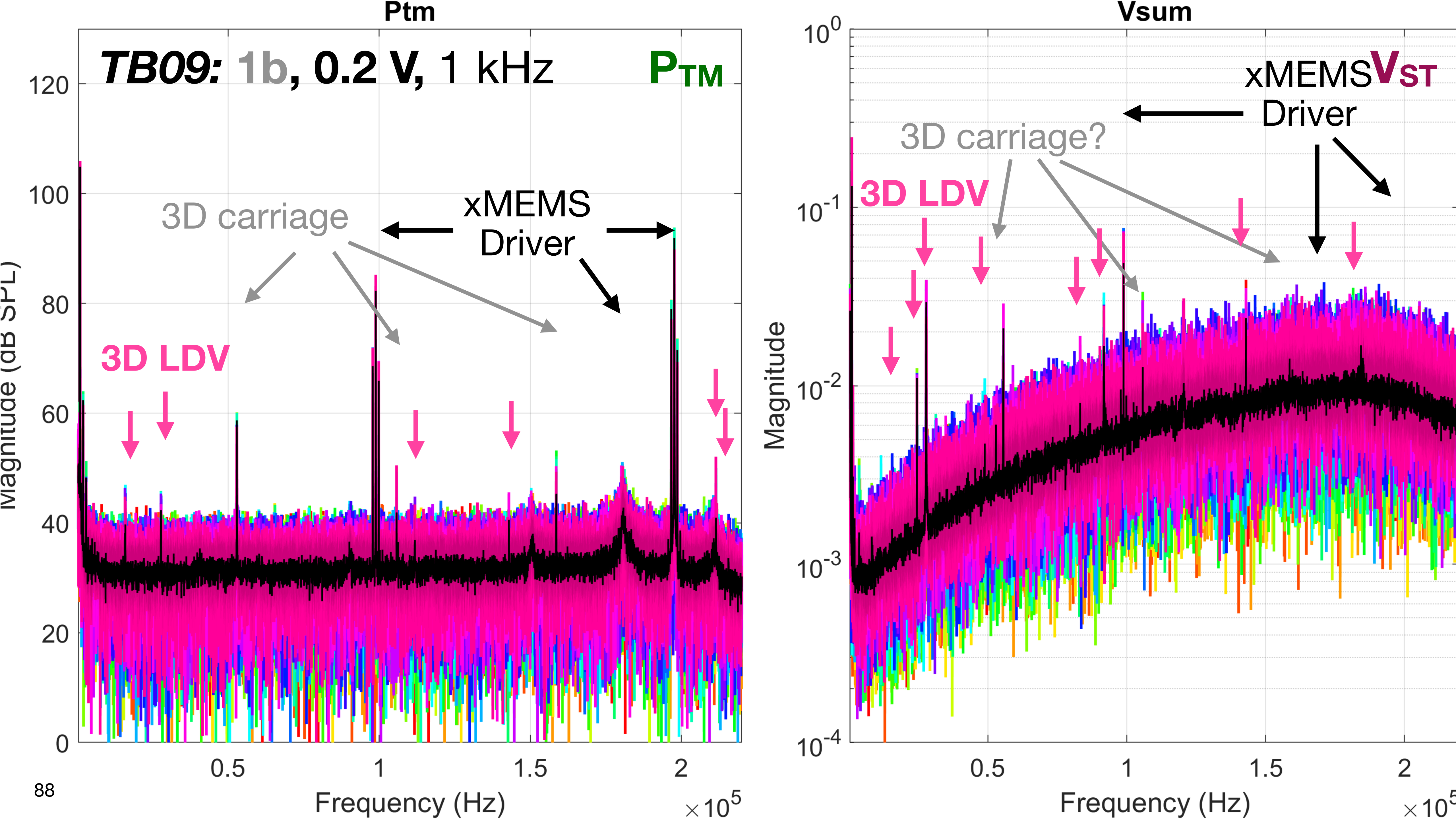

Ptm
TB09: 1b, 0.2 V, 1 kHz
PTM
3D carriage
xMEMS
Driver
3D LDV
Magnitude (dB SPL)
Frequency (Hz)
×10^5
Vsum
xMEMS VST
Driver
3D carriage?
3D LDV
Magnitude
Frequency (Hz)
×10^5

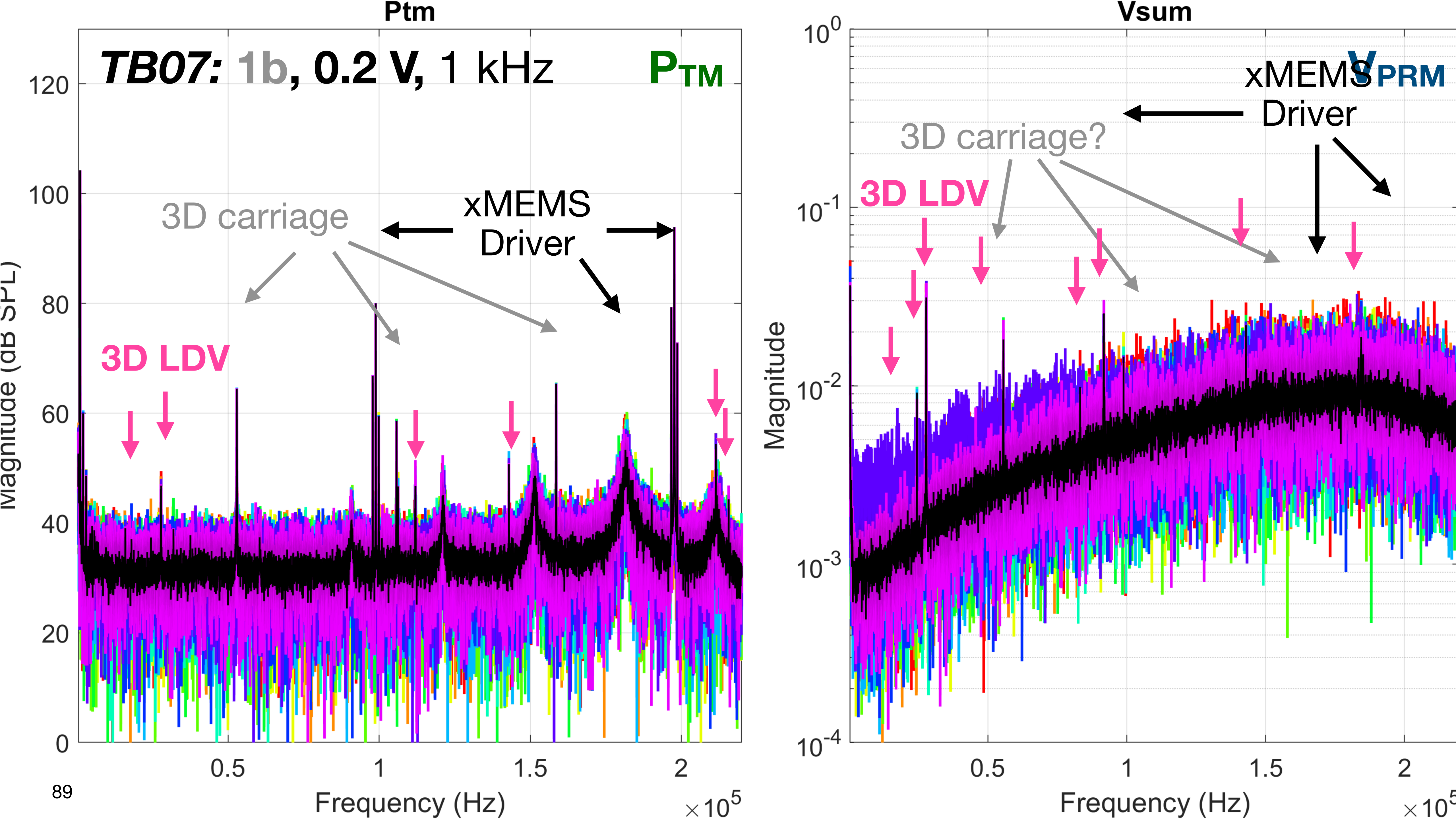
Ptm
TB07: 1b, 0.2 V, 1 kHz
PTM
3D carriage
xMEMS
Driver
3D LDV
Frequency (Hz)
×10^5
Vsum
VPRM
xMEMS
Driver
3D carriage?
3D LDV
Magnitude
Frequency (Hz)

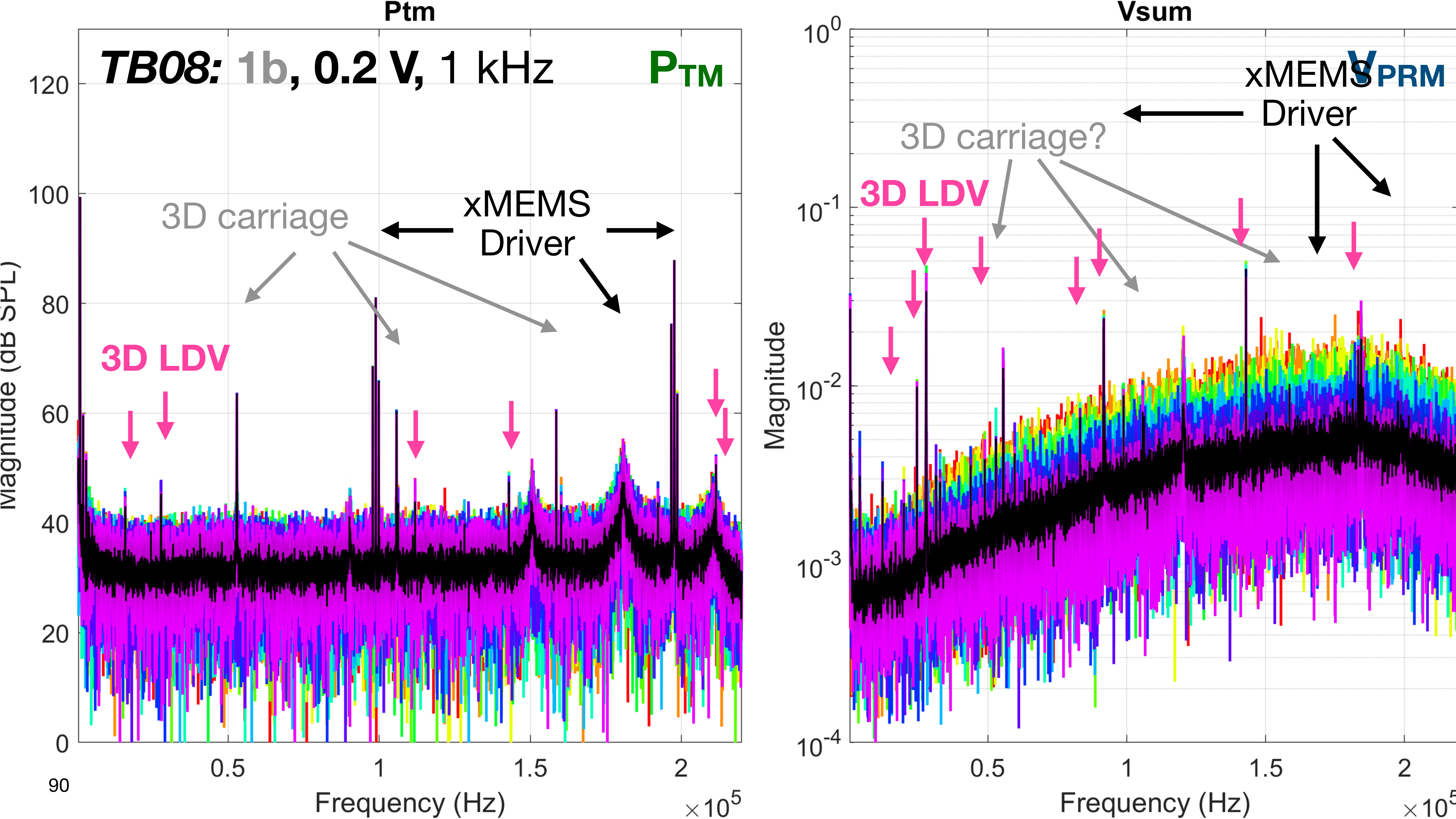

Ptm
TB08: 1b, 0.2 V, 1 kHz
PTM
3D carriage
xMEMS
Driver
3D LDV
Magnitude (dB SPL)
Frequency (Hz)
×10^5
Vsum
VPRM
xMEMS
Driver
3D carriage?
3D LDV
Magnitude
Frequency (Hz)

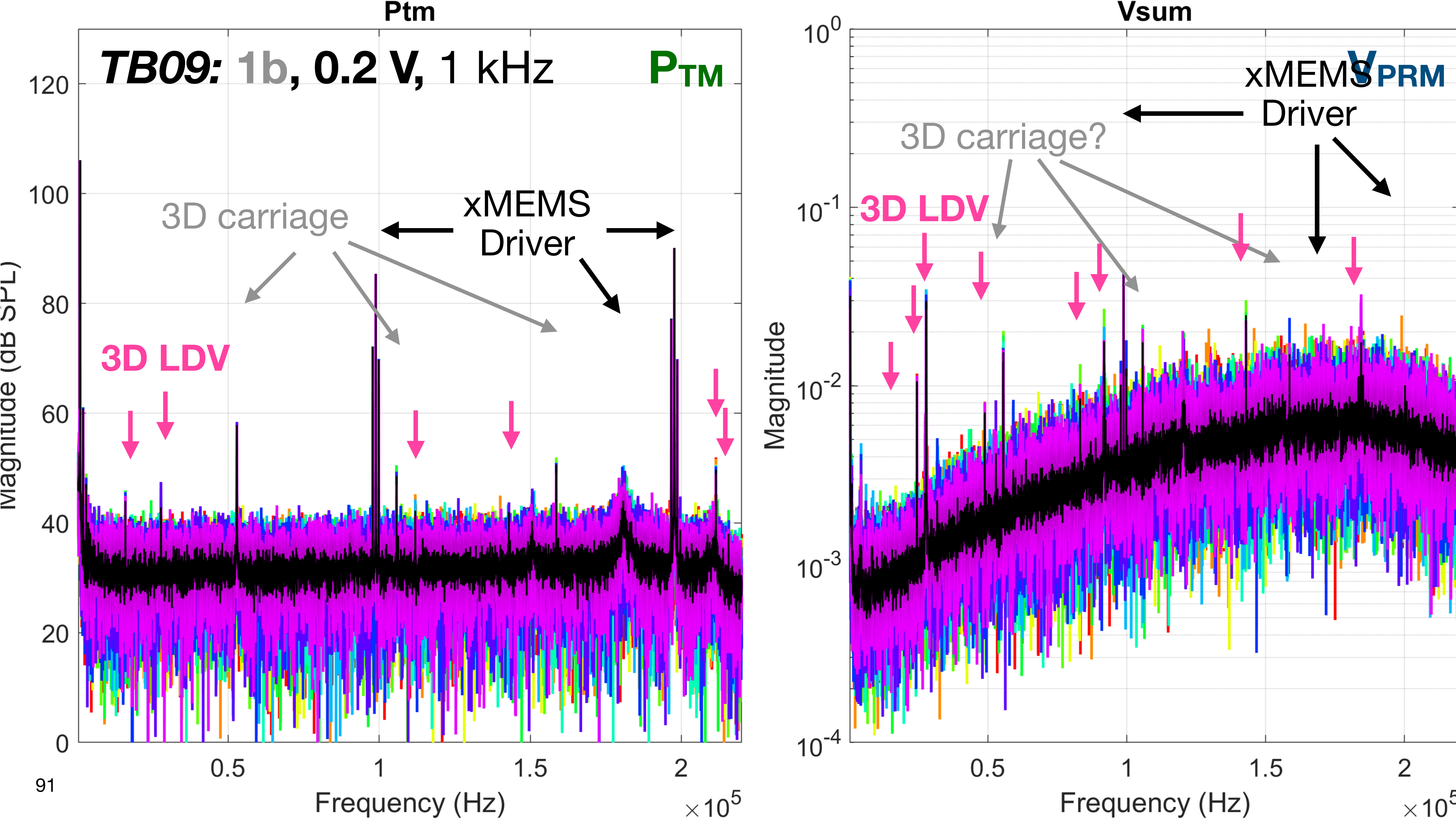

Ptm
TB09: 1b, 0.2 V, 1 kHz
P_TM
3D carriage
xMEMS Driver
3D LDV
Magnitude (dB SPL)
Frequency (Hz)
×10^5
Vsum
V_PRM
xMEMS Driver
3D carriage?
3D LDV
Magnitude
Frequency (Hz)

### Conclusions

- The 3D Carriage Controller introduces 3 peaks that are picked up electrically, mostly on the pressure channel: a 52.881 kHz fundamental and its 2nd (105.762 kHz) and 3rd (158.643 kHz) harmonics.
  - Peaks at similar frequencies sometimes appear in the velocity response.
- The 3D LDV introduces numerous peaks in the pressure and velocity responses and raises the velocity noise floor.
  - The peaks appear in the pressure at: 16.3, 27.8, 111.8, 143.2, 211.6, and 215.5 kHz
  - The peaks appear in the velocity at: 16.4, 24.4, 27.8, 48.9, 83.5, 92, 143.2, and 184 kHz
- The xMEMS speaker driver introduces sharp peaks at the valve, carrier, and modulation frequencies, plus the drive frequency; as well as a shorter wide peak around 180 kHz that mostly appears in the pressure.
  - Temporal-bone measurements can exhibit additional wide peaks below 180 kHz.
